# Deep learning based behavioral profiling of rodent stroke recovery

**DOI:** 10.1101/2021.08.11.455647

**Authors:** Rebecca Z Weber, Geertje Mulders, Julia Kaiser, Christian Tackenberg, Ruslan Rust

## Abstract

Stroke research heavily relies on rodent behavior when assessing underlying disease mechanisms and treatment efficacy. Although functional motor recovery is considered the primary targeted outcome, tests in rodents are still poorly reproducible, and often unsuitable for unraveling the complex behavior after injury. Here, we provide a comprehensive 3D gait analysis of mice after focal cerebral ischemia based on the new deep learning-based software (DeepLabCut, DLC) that only requires basic behavioral equipment. We demonstrate a high precision 3D tracking of 10 body parts (including all relevant joints and reference landmarks) in several mouse strains with an accuracy of 99.4%. Building on this rigor motion tracking, a comprehensive post-analysis (with >100 parameters) unveils biologically relevant differences in locomotor profiles after a stroke over a time course of three weeks. We further refine the widely used ladder rung test using deep learning and compare its performance to human annotators. The generated DLC-assisted tests were then benchmarked to five widely used conventional behavioral set-ups (neurological scoring, rotarod, ladder rung walk, cylinder test, and single-pellet grasping) regarding sensitivity, accuracy, time use and costs. We conclude that deep learning-based motion tracking with comprehensive post-analysis provides accurate and sensitive data to describe the complex recovery of rodents following a stroke. The experimental set-up and analysis can also benefit a range of other neurological injuries that affect locomotion.

## Introduction

Stroke is a leading cause of disability and death worldwide. Over 13.7 million strokes occur each year, and one in four people over 25 years of age will experience a stroke in their lifetime^1^. The presence of life-saving medicines allows timely intervention, which has significantly decreased mortality following a stroke^2,3^. However, acute treatments are not applicable in most patients, mainly because of the narrow therapeutic time window, leaving five million patients permanently disabled every year^4,5^. To promote recovery outside the confines of conventional therapies, a variety of experimental treatments in rodents have emerged targeting neuroprotection^6^, therapeutic angiogenesis^7–9^, axonal sprouting^10^, or cell-based therapies^11,12^. In most of these studies, behavioral evaluation is the primary outcome and ultimately provides evidence that functional impairment can be corrected by the experimental treatment. However, behavioral tests in rodents have proved difficult: (1) test results are often poorly reproducible and (2) the task is limited to a specific sensorimotor outcome, thus ignoring most of the other biologically relevant parameters of functional recovery after stroke^13^.

Advances in high-speed video equipment have enabled scientists to record massive datasets of animal behavior in exquisite detail, and commercial software solutions including Ethovision (Noldus), AnyMaze (Stoelting Co.), and Top Scan (CleverSys Inc.) have assisted with vision-based tracking and analysis. However, these technologies offer little methodological transparency, are not affordable for many laboratories^14^, and are often designed to study pre-specified modules within one particular paradigm (e.g., the Morris water maze or the open field test) rather than discover new behavioral patterns. The introduction of machine learning algorithms has recently permeated various sectors of life and provided a new set of tools ideally suited for behavior analysis. These algorithms, referred to as deep learning models, offer user-defined feature tracking with greater flexibility, as well as reduced software and hardware acquisition costs^15^. One of the latest contributions to this toolbox is the open-source software DeepLabCut (DLC)^16^, which uses convolutional neural networks to automatically capture movements and postures directly from images and without requiring active or passive markers. DLC is a modified version of a state-of-the-art algorithm for tracking human movement, DeeperCut^17^ and can be used in a broad range of study systems with near human-level accuracy^18,19^. Typically, such algorithms are seen as “data-hungry”; algorithms must be trained first by showing thousands of hand-labeled frames, an effort that requires an enormous amount of time. DLC, however, is pre-trained on ImageNet, a large database of images used for image recognition research^20^. With that pretraining in place, DLC only needs a few training examples (typically 50 - 200 frames) to achieve human-level accuracy, making it a highly data-efficient software^16,21^. DLC has already been implemented in different research fields including neuroscience^22–24^.

In this study, we developed a modular experimental set-up to identify biologically relevant parameters to reveal gait abnormalities and motor deficits after a focal ischemic stroke. We trained the neural networks to recognize mice of different fur colors from three perspectives (left, bottom, and right) and to label 10 body parts with high accuracy. A detailed comprehensive post-hoc script allows analysis of a wide range of anatomical features within basic locomotor functions, vertical and horizontal limb movements, and coordinative features using the freeware software environment R. We detect distinct changes in the overall mouse gait affecting e.g., step synchronization, limb trajectories and joint angles after ischemia. These changes are distinct at acute and chronic time points and primarily (but not exclusively) affect the body parts contralateral to the lesion. We further refine the conventional ladder rung tests with DLC (e.g., for detection of foot placements) and compare the deep learning-assisted analysis with widely used behavioral tests for stroke recovery that use human annotations, the gold standard. We detect similar levels of accuracy, less variation, and a considerable reduction in time using the DLC-based approach. The findings are valuable to the stroke field to develop more reliable behavioral readouts and can be applied to other neurological disorders in rodents involving gait abnormalities.

## Results

### Generation of a comprehensive locomotor profile using deep learning-based tracking

Our aim was to develop a sensitive and reliable profiling of functional motor recovery in mice after stroke using open-access deep learning software, DLC. Unraveling the complexity of changes in locomotion is best approached via generation of gait parameters^25^. Therefore, we customized a free walking runway with two mirrors that allowed 3D recording of the mice from the lateral/side and down perspectives. The runway can be exchanged with an irregular ladder rung to identify fine-motor impairments by paw placement (Fig. 1A). The dimensions of the set-up were adapted from the routinely used MotoRater (TSE Systems)^26^. After adaptation to the set-up, non-injured mice were recorded from below with a conventional GoPro Hero 8 camera during the behavioral tasks. The DLC networks were trained based on ResNet-50 by manually labeling 120 frames from randomly selected videos of different mice. Individual body parts were selected according to previous guidelines to enable a comprehensive analysis of coordination, movement, and relative positioning of the mouse joints from all three perspectives, and included: tail base, iliac crest, hip, back ankles, back toe tip, shoulder, wrist, elbow, front toe tip, and head (Fig. 1B, C)^27^.

**Fig. 1:**
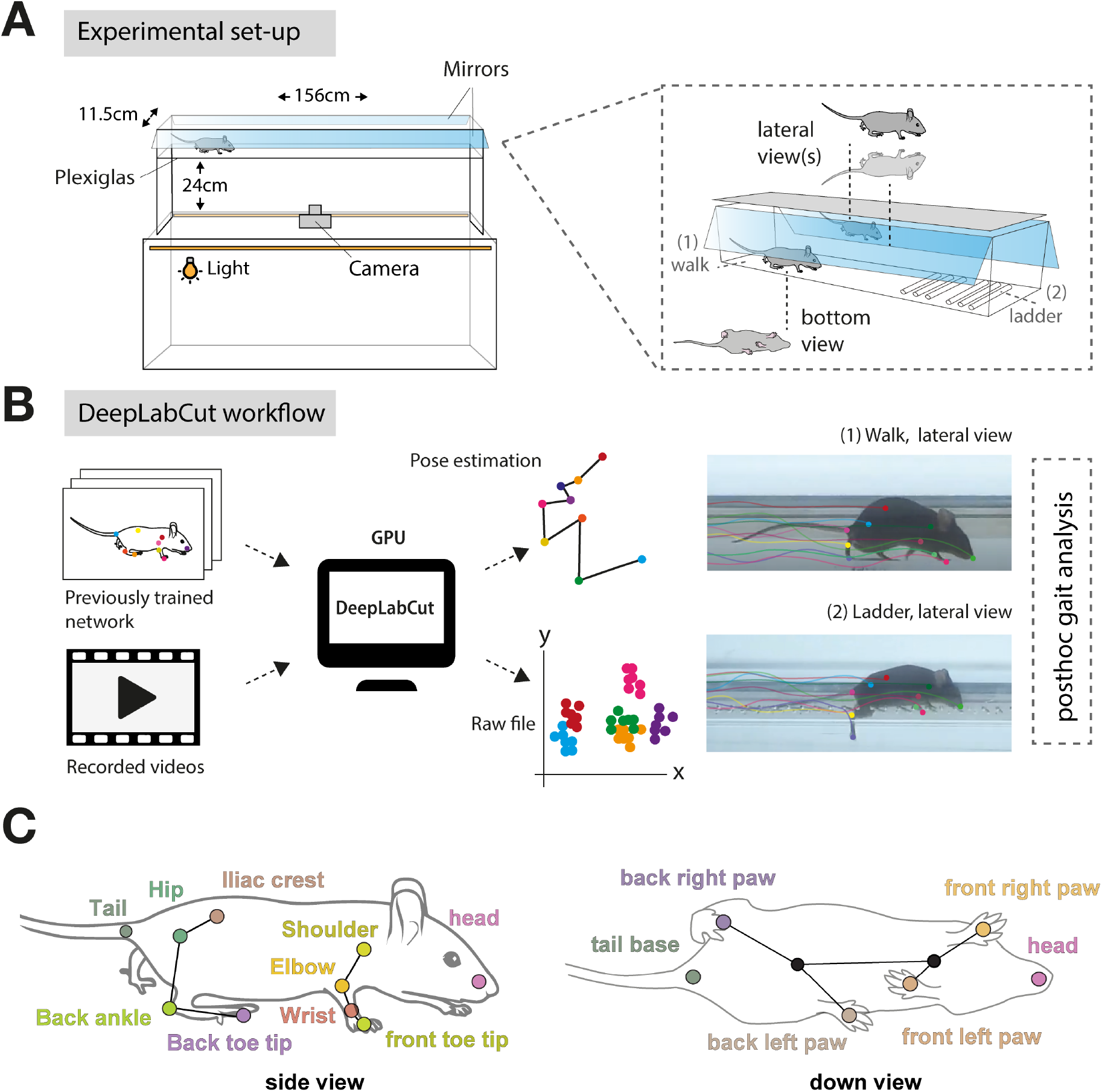
Experimental workflow to perform deep learning-based gait analysis. **(A)** Schematic view of the dimensions of experimental set-up. **(B)** Workflow to identify and label anatomical landmarks of mice for pose estimation. **(C)** Overview of labeled body parts from side and down perspective.

Next, we applied the neural network model to detect and extract the relevant body coordinates in each frame of all recorded videos. A training set of six videos proved sufficient to achieve a cross-entropy loss of < 0.1% indicating a marginal predicted divergences from the actual label after 500’000 iterations (Fig. 2A). A more detailed analysis revealed that all selected body part labels could be tracked to >99% (Fig. 2B). The ratio of confident labels (>95% likelihood) to total labels ranged from 96-100% for the runway and between 89-100% for the rung walk (Fig. 2B, C). In both set-ups, we observed the highest variability for the front and back toe tips. For further analysis all data points that did not pass the likelihood of detection threshold of 95% were excluded. The remaining data generated a full 3D profile of each animal during the behavioral task (Suppl. Fig 1A, B).

**Fig. 2:**
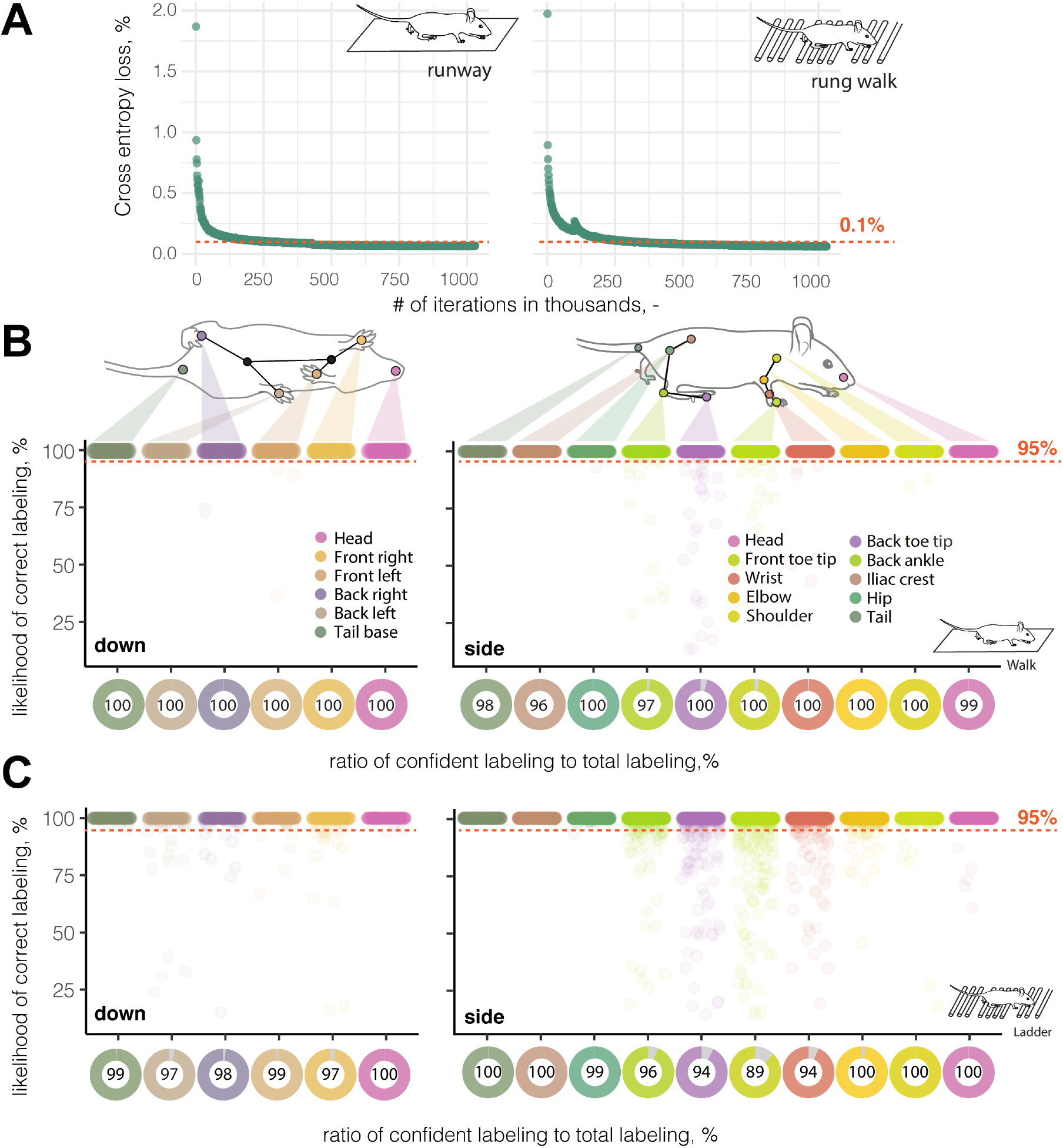
DeepLabCut enables markerless 3D tracking of mouse body parts with high levels of accuracy. **(A)** Training efficiency of neural networks. (**B)** Likelihood of a confident labeling for individual body parts from down view (left) side view (right) in the runway and **(C)** during the ladder rung walk. Each dot represents an anatomical landmark of individual image frames in a video. The red dotted line represents the confidence threshold of 95% likelihood for reliable labeling.

Next, we used the same trained networks to reliably label body parts of a) the same mice three days after stroke, b) different mice with the same genetic background (C57BL/6J, black fur) and c) mice with a different genetic background (NOD, white fur). We achieved similar confidence in labeling for mice after stroke (95–100%) and mice with the same genotype (97–100%) after minor refinement of the network (see Methods, Suppl. Fig. 2A, B). However, we were unable to successfully refine the pre-existing network to track mice of a different strain with white fur (0-41%). We then created an entirely new training set for these mice with the same training parameters and reached similar levels of confidence (94–100%) to the original training set (Suppl. Fig. 2C, D).

Overall, we demonstrated successful labeling and generation of 3D locomotor coordinates in non-injured and injured mice of different genetic backgrounds and fur colors for both the runway and ladder rung walk using deep learning.

### Deep learning trained networks detect distinct gait abnormalities following stroke

To identify stroke-related gait abnormalities across a specific time period, we induced a photothrombotic stroke in the right hemisphere of the sensorimotor cortex (Fig. 3A, B) ^7,28^. We confirmed successful induction of ischemia with a reduction of cerebral blood flow only in the ipsilesional right side (right: –72.1 ± 11.5%, p < 0.0001, left: –3.2 ± 8.6%, p = 0.872) using Laser doppler imaging 24 h after injury (Fig. 3C). Three weeks after injury, mice had histological damage in all cortical layers, which was accompanied by a microglial activation and glial scar formation on the ipsilesional hemisphere while sparing subcortical regions and the contralesional side. The injured tissue extended from +2 mm to –2 mm anterior posterior related to bregma, and the average stroke volume was 1.3 ± 0.2 mm^3^ (Fig. 3D, E).

**Fig. 3:**
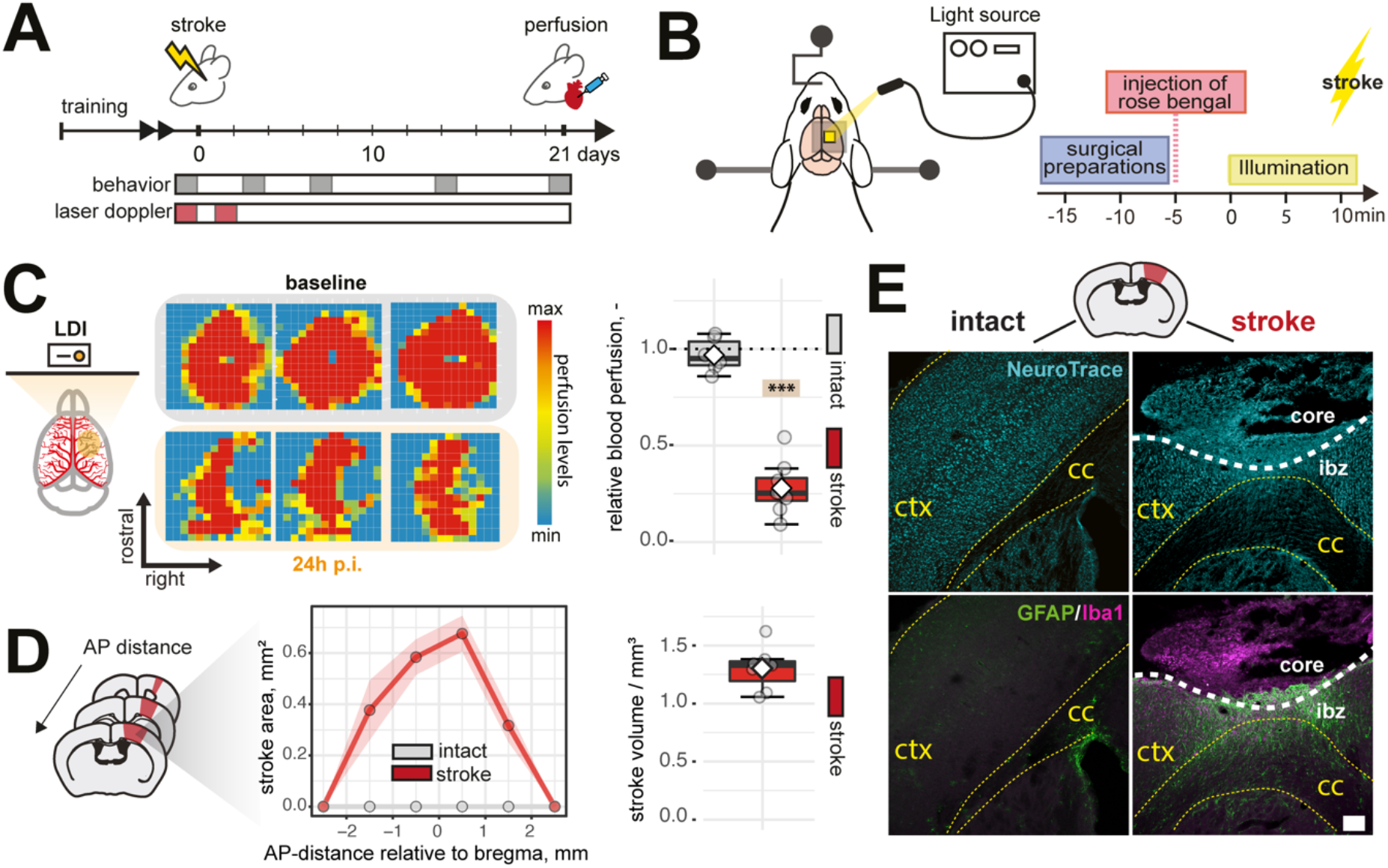
Induction of photothrombotic stroke leads to permanent focal ischemia in the cortex. **(A)** Schematic time course of experimental interventions **(B)** Schematic representation of stroke procedure **(C)** Laser Doppler Imaging (LDI) of three representative baseline and stroked brains 24 hours after injury. **D)** Quantification of stroke area and stroke volume at 21 dpi. **(E)** Representative histological overview of cortical damage (Neurons, cyan), inflammatory infiltration (Iba1+, magenta) and scar formation (GFAP^+^, green) at 21 dpi, scale bar: 100 µm. Data are shown as mean distributions where the white dot represents the mean. Boxplots indicate the 25% to 75% quartiles of the data. For boxplots: each dot in the plots represents one animal. Line graphs are plotted as mean ± sem. Significance of mean differences between the groups (baseline hemisphere, contralesional hemisphere, and ipsilesional hemisphere) was assessed using Tukey’s HSD. Asterisks indicate significance: ^***^ P < 0.001. ctx: cortex, cc: corpus callosum, ap: anterior posterior, p.i.: post injury, ibz: ischemic border zone.

We began the motion tracking analysis by assessing the overall gait at baseline and after injury over a three-week period. Individual steps were identified by the movement speed of each limb between frames as filmed from below (Fig. 4A, B). In uninjured animals the footfall pattern showed a typical gait synchronization^29^ of opposing front and back paws (Fig. 4C). Normalizing the data to a single step cycle revealed that this pattern was severely altered acutely after injury as shown by single-animal data (Fig. 4D). We noticed that the asynchronization between the paws is acutely increased after injury (p < 0.001, 3 dpi (days post injury); p = 0.029, 7 dpi; p = 0.031, 14 dpi) but recovered to baseline over 21 days (p = 0.81, 21 dpi; Fig. 4E). Furthermore, acutely injured mice walked slower as the step cycle duration was increased compared to intact mice (p = 0.05, 3 dpi, Fig. 4F). While the swing duration did not differ at any time point (all p > 0.05), stroked mice had a longer stance duration (p = 0.04, 3dpi, Fig. 4G, H). These alterations in the footfall pattern were associated with changes in the positioning of the paws during a step (Fig. 4I, J). The angle amplitude of the ipsilesional hindlimb relative to the body center increased acutely after injury (p = 0.003, 3dpi) while the angle of the front limbs remained unchanged (all p > 0.05, Fig. 4K).

**Fig. 4:**
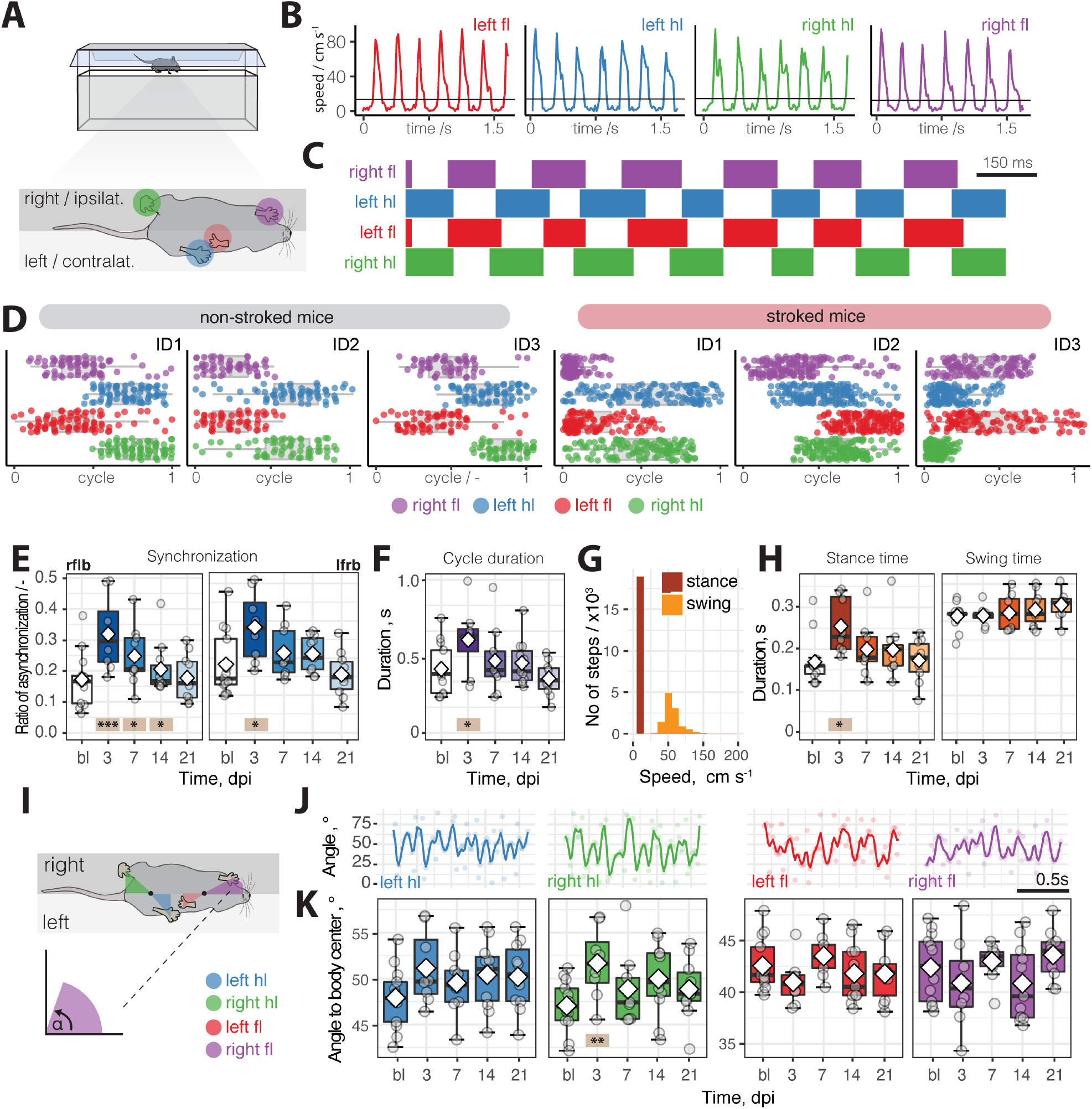
Gait changes in footfall pattern in spontaneous walk after stroke. (**A**) Schematic set-up of runway walks from bottom perspective. (**B**) Movement speed of individual fore- and hindlimbs during spontaneous walk. (**C)** Footfall profile of single mouse without injury. (**D**) Footfall profiles of a normalized locomotor cycle showing the stance and phase start and end of three individual control mice (left) and stroked mice (right). (**E**) Ratio of asynchronization at baseline, 3,7,14,21 dpi. (**F**) Duration of a cycle. (**G, H**) Comparison of cycle duration between stance and swing time in a time course. (**I**) Schematic view on analysis of positioning paws to body centers. (**J**) Profile of paw angles relative to body center of an individual animal. (**K**) Comparison of angles of individual paws to body center in a time course. Data are shown as mean distributions where the white dot represents the mean. Boxplots indicate the 25% to 75% quartiles of the data. Each dot in the plots represents one animal and significance of mean differences between the groups was assessed using repeated ANOVA with post-hoc analysis. Asterisks indicate significance: ^*^ P < 0.05, ^**^ P < 0.01, ^***^ P < 0.001

Overall, the synchronization of the footfall pattern was severely altered during the acute phase of stroke but returned to a normal pattern in the long-term

The kinematics of a spontaneous walk were then compared by tracking the fore- and hindlimb joints from the left and right-side perspectives (Fig. 5A, B). First, we analyzed the average height and total vertical movement of each joint involved in the hindlimb (iliac crest, hip, back ankle, and back toe tip) and forelimb movement (shoulder, elbow, wrist, and front toe tip, Fig. 5C, D). We identified alterations in the total vertical movement and average height of the fore- and hindlimb joints that was, as expected, more prominent on the contralateral left side. Most notably, the total vertical movement decreased in the contralateral left back and front toe tips, shoulder, and wrist at 3 dpi (all p < 0.05) with a partial but incomplete recovery over time (Fig. 5E, Suppl. Fig. 3). Interestingly, we also observe compensatory changes in the vertical movement on the ipsilateral right side most prominent in the back toe tip, back ankle, elbow, and wrist (Fig. 5E, Suppl. Fig 3). Next, we checked for alterations in the horizontal movement determining the average step length, as well as protraction, and retraction of the individual paws. At 3 dpi both retraction and protraction length are reduced in stroked mice. These changes remained more pronounced in the hind limbs during retraction whereas protractive changes returned to normal throughout the time course (Fig. 5F). Like the vertical movement, we also observed compensatory changes in protraction in the ipsilateral right hindlimb at later time points. Then, the joint positions were used to extract the angles of the hindlimbs (iliac crest-hip-ankle; hip-ankle-toe tip) and forelimbs (shoulder-elbow-wrist; elbow-wrist-toe tip). The angular variations were acutely unchanged after stroke and showed a similar profile throughout the time course (Fig. 5G, H, Suppl. Fig. 4).

**Fig. 5:**
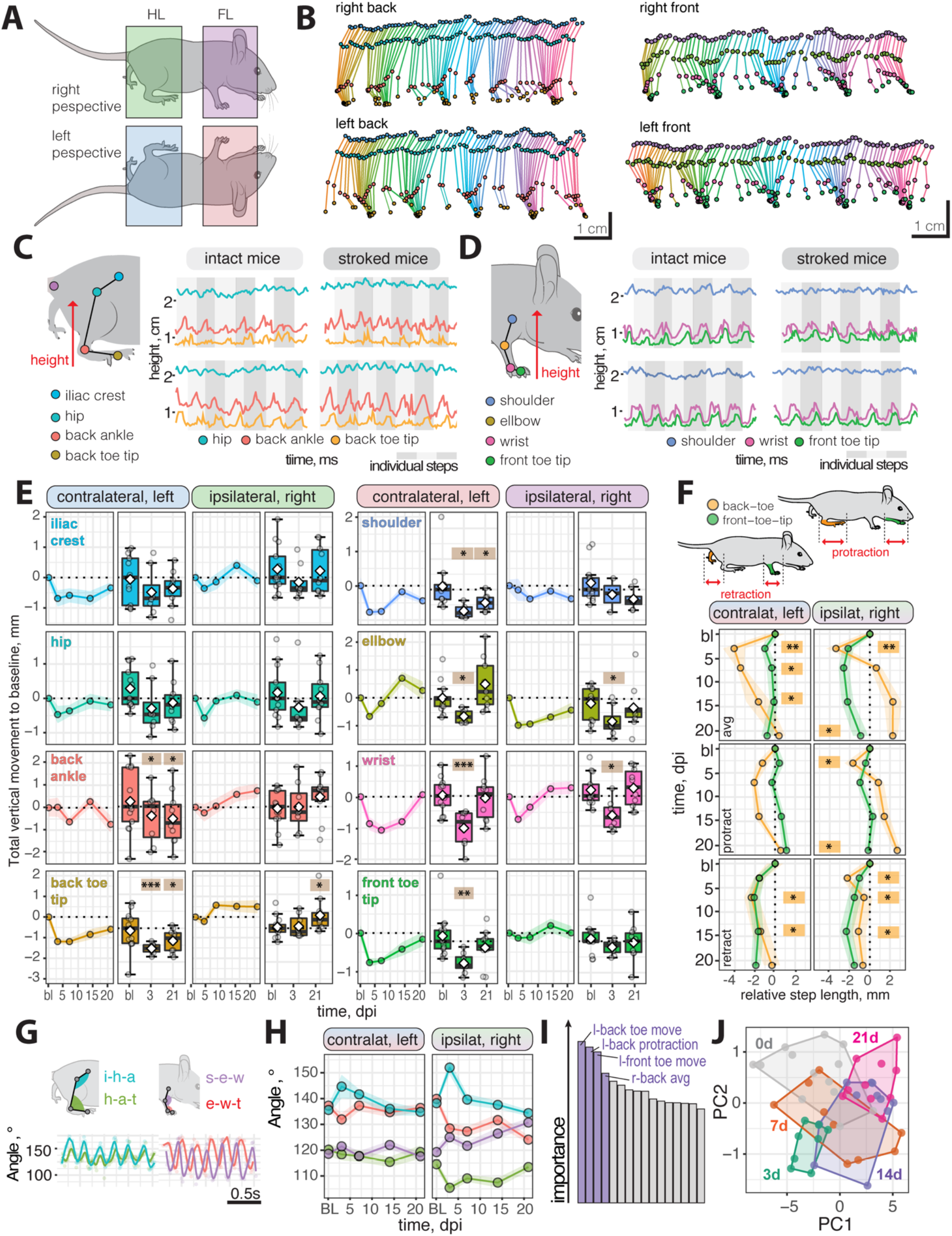
Kinematic changes in runway walk after stroke. (**A**) Schematic overview of analysis from the sides (**B**) Stick profile of fore and hindlimb movement in individual mouse (**C, D**) Profile of hindlimb and forelimb joints in intact and stroked mice (**E**) Absolute height of selected joints at baseline and 3,7,14,21 dpi. (**F**) Protraction and retraction of joints throughout a time course (**G, H**) Angular variability between front and hindlimb joints (**I**) Random Forest classification of most important parameters (**J**) Principal component analysis of baseline 3, 7, 14, 21 dpi. Data are shown as mean distributions where the white dot represents the mean. Boxplots indicate the 25% to 75% quartiles of the data. For boxplots: each dot in the plots represents one animal. Line graphs are plotted as mean ± sem. For line graphs: the dots represent the mean of the data. Significance of mean differences between the groups was assessed using repeated ANOVA with post-hoc analysis. Asterisks indicate significance: ^*^ P < 0.05, ^**^ P < 0.01, ^***^ P < 0.001. i-h-a: iliac crest-hip-ankle, h-a-t: hip-ankle-toe, s-e-w: shoulder-elbow-wrist, e-w-t: elbow-wrist-toe, PC: principal component.

To understand the individual importance of the >100 measured parameters (Table 1) during kinematics analysis, we applied a random forest classification to all animals throughout the entire time course (Fig. 5I). The most important parameters between the groups were: left front and back toe heights as well as protractive and total horizontal back toe movements. A separate analysis was performed between baseline and acutely injured mice at 3 dpi and mice with long-term deficits at 21 dpi (Suppl. Fig. 5). In these subgroup analyses, we were able to predict the acute injury status with 90% accuracy and long-term deficits with 85% accuracy using a confusion matrix. The overlap between the most important 20 parameters (top 10% of all measured parameters) in acute and chronic time points was only 20% further confirming the need to consider the entirety of the gait to understand the complexity of functional recovery over time (Suppl. Fig. 5). We then used a principal component analysis (PCA) to reduce the dimensions of our data and determine the differences between the groups. We found that data from later time points after injury cluster closer to the baseline suggesting that the recovery effects can be ascertained based on kinematic parameters. The separation expand when comparing only data from 3 dpi and 21 dpi to baseline (Suppl. Fig. 5). Importantly, these changes were not observed in non-stroked control mice throughout the time course (Suppl. Fig. 6).

Next, we considered whether deep networks can also be applied to conventional behavioral tests to detect fine motor deficits in a ladder rung test (Fig. 6A). Tracking the fore- and hindlimbs during the ladder rung recordings enabled the identification of stepping errors in the side view (Fig. 6 B, C). We identified a 106% increase of the overall error rate in injured animals compared with their intact controls (intact 5.27 ± 8.4%, stroked: 10.9 ± 12.6%, p < 0.001) at 3 dpi. This increased error rate after acute stroke was most pronounced on the contralesional side (left front paw: +182%; left back paw: +142%, both p < 0.001) but also marginally detectable in the ipsilesional site (right front paw: +21%, p = 0.423; right back paw: +17%, p < 0.001; Fig 6 D). In a time course of three weeks, we detected a marked increase of footfall errors in both the contralateral left front and back toe (all p < 0.001) compared to baseline. Although the error rate returned to baseline for the back paw (p = 0.397), it does not fully recover for front paw (p = 0.017), as previously observed ^7^ (Fig. 6E).

**Fig. 6:**
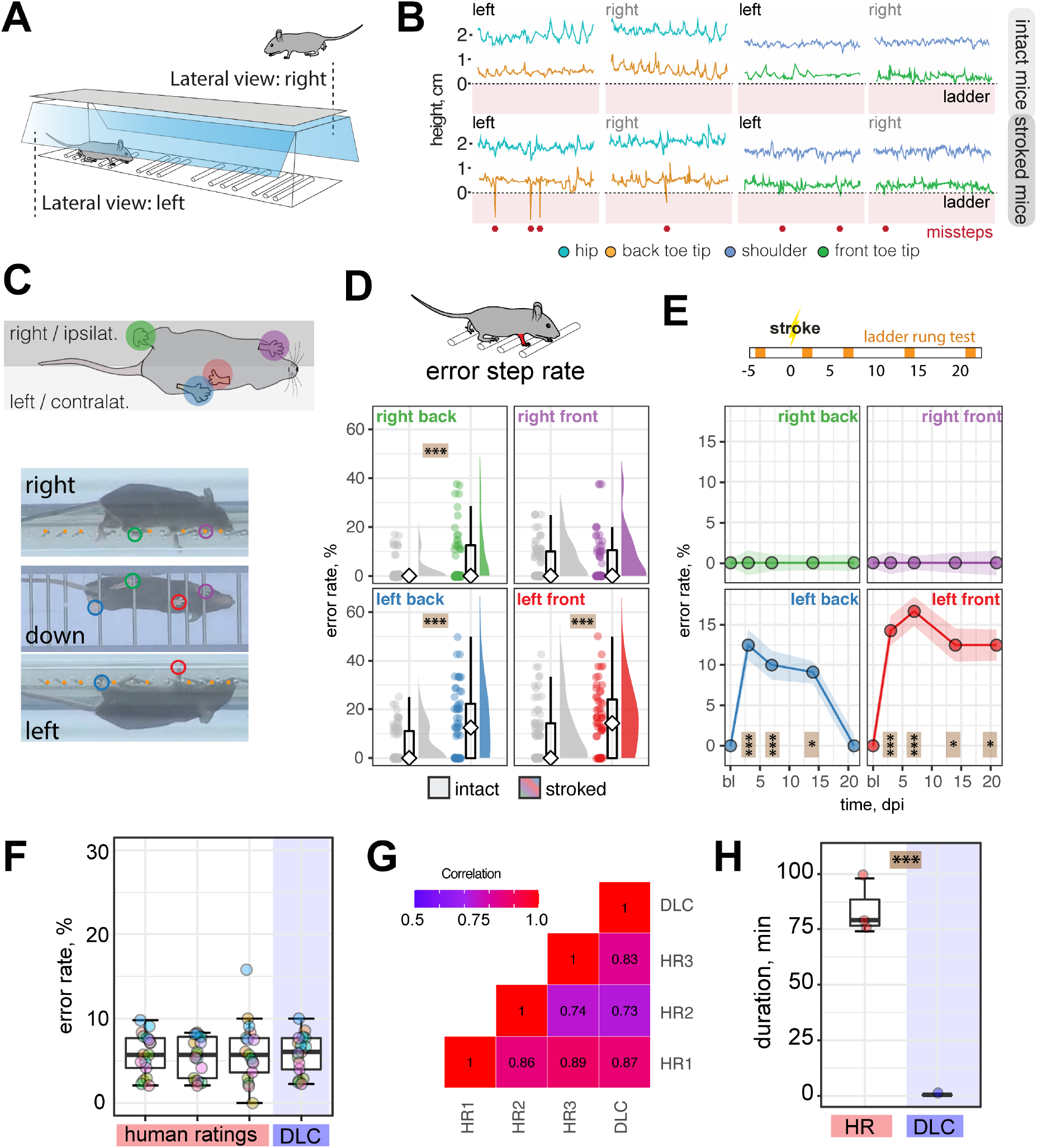
DeepLabCut assisted analysis of horizonal rung test after stroke. (**A**) Schematic view of ladder rung test. (**B**) Side view of step profile of hips, back toes, shoulder, and front toes in individual animals at baseline and 3 dpi. (**C**) Photographs of mouse from three perspectives. (**D**) Overall success and error rate in the contralesional and ipsilesional hemisphere of all paws. (**E**) Time course of error rate during ladder rung test in the individual paws. (**F**) Comparison of error rate scores in selected videos between three human observers and DLC. (**G**) Correlation matrix between human observers and DLC. (**H**) Duration of analysis for ladder rung test. Data are shown as mean distributions where the white dot represents the mean. Boxplots indicate the 25% to 75% quartiles of the data. For boxplots: each dot in the plots represents one animal. Line graphs are plotted as mean ± sem. For line graphs: the dots represent the mean of the data. Significance of mean differences between the groups was assessed using repeated ANOVA with post-hoc analysis. Asterisks indicate significance: * P < 0.05, ** P < 0.01, *** P < 0.001.

In a subset of 20 randomly selected videos, we cross verified the error rates by a blinded observer and compared the variability between the DLC-approach and the manual assessment of the parameters regarding (1) variability of the analysis and (2) duration of the analysis. We did not detect a difference in the scoring accuracy between the manual assessment and DLC-assisted analysis, but manual assessment required 200 times as more time (human: 4.18 ± 0.63 min; DLC: 0.02 min, p < 0.0001; Fig. 6F-H, Suppl. Fig. 7).

Overall, these results suggest that DLC-assisted analysis of the ladder rung test achieves human-level accuracy, while saving time and avoiding variability between human observers.

### Comparison of deep learning-based tracking to conventional behavioral tests for stroke-related functional recovery

Finally, we benchmarked DLC-tracking performance against popular functional tests to detect stroke-related functional deficits. We performed a rotarod test with the same set of animals and analyzed previously acquired data from a broad variety of behavioral tasks routinely used in stroke research including neurological scoring, cylinder test, the irregular ladder rung walk, and single pellet grasping (Fig. 7A, B).

**Fig. 7:**
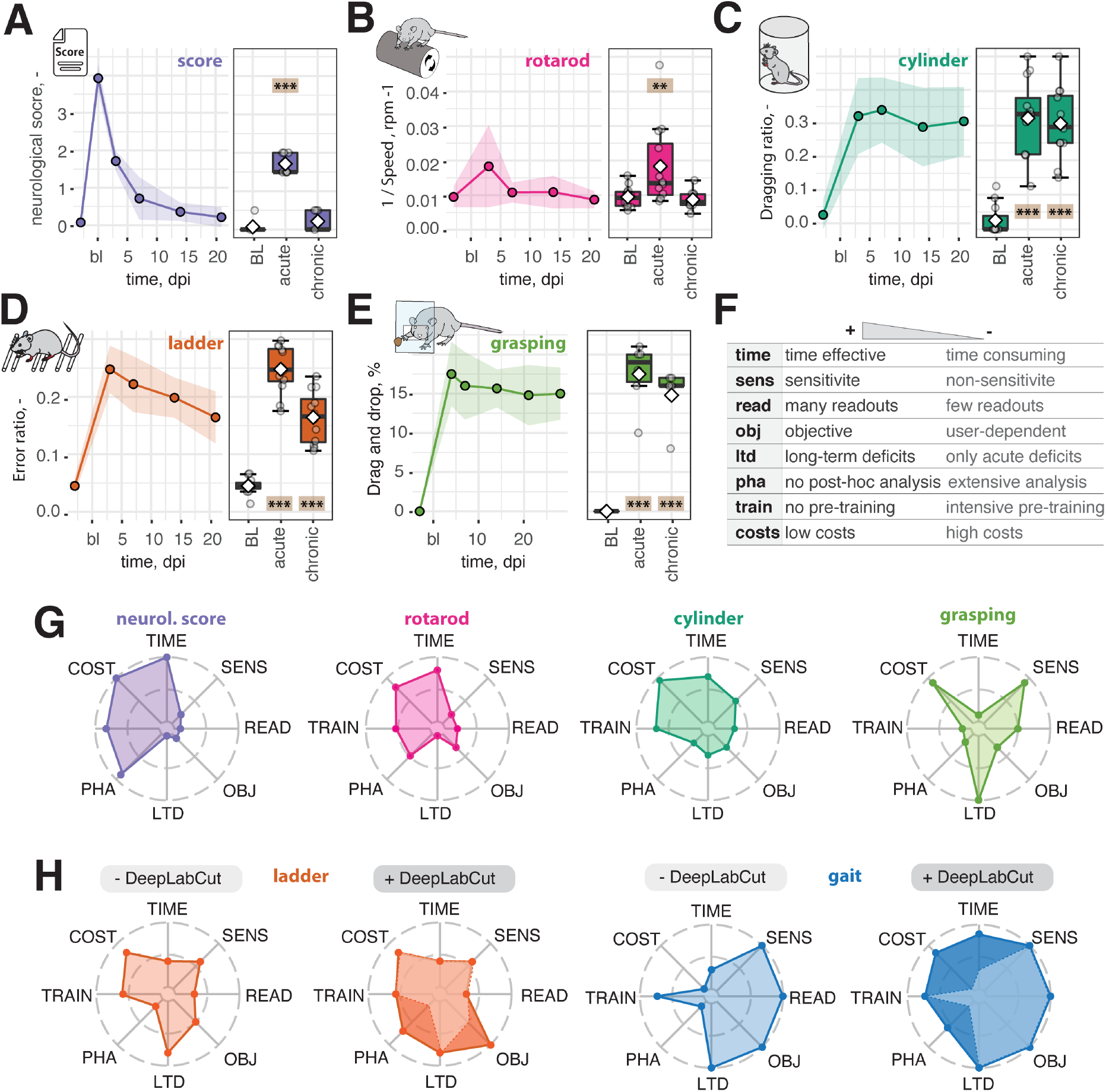
Functional assessment of recovery after stroke using conventional behavioral tests. (**A**) Neurological score, (**B**) Rotarod test, (**C**) Dragging during cylinder test, (**D**) Missteps in ladder rung test, (**E**) Drag and drop in single pallet grasping. (**F**) Semi-quantitative measure of relevant parameters for behavioral tests (time, sensitivity, readouts, objectivity, long-term deficits, post-hoc analysis, pre-training, and costs). (**G**) Spider chart of conventional behavioral tests (**H**) Spider chart of behavioral tests without and with DLC assistance. Dara are shown as line graphs and are plotted as mean ± sem. Significance of mean differences between the groups was assessed using repeated ANOVA with post-hoc analysis. Asterisks indicate significance: ** P < 0.01, *** P < 0.001.

In all behavioral set-ups, we identified initial deficits after stroke (rotarod: p= 0.006, all other tests: p < 0.001). While the neurological deficit score (21 dpi, p = 0.97) and the rotarod (21 dpi, p = 0.99) did not provide much sensitivity beyond the acute phase (Fig. 7A, B), the ladder rung test, cylinder test, and single pellet grasping were suitable to reveal long-term impairments in mouse stroke models (21 dpi, all p < 0.001, Fig. 7C-E, Suppl. Fig. 8A).

These functional tests were then further compared in a semi-quantitative spider diagram regarding (1) duration to perform the task, (2) objectivity, (3) post-hoc analysis, (4) requirement of pre-training, and (5) costs (Fig. 7G, Suppl. Fig. 8B, C). Despite the simple performance, the neurological scoring, rotarod and cylinder tests have the drawback of a relatively low sensitivity and objectivity. On the other hand, more sensitive tests such as the pellet grasping test require intense pre-training of the animals, or the manual post-analysis of a ladder rung test can be tedious and suffer from variability between investigators. All these conventional tests only provide a very low number of readouts, which may not capture the entire complexity of the acute injury and subsequent recovery.

More advanced analysis including kinematic tracking offers the advantage of generating a variety of parameters but the high costs for the set-up and the commercial software are disadvantages (Fig 7H). The DLC-assisted tracking presented here provides an open-source solution that is available at negligible costs and can be set up easily. The experiment duration is shortened, and animal welfare is improved since the test does not require marking the mouse joints beforehand. Most importantly, using our comprehensive post-analysis, the set-up reduces analysis time while minimizing observer biases during the evaluation.

## Discussion

Preclinical stroke research heavily relies on rodent behavior when assessing functional recovery or treatment efficacy. Nonetheless, there is an unmet demand for comprehensive unbiased tools to capture the complex gait alterations after stroke; many conventional methods either do not have much sensitivity aside from identifying initial injures; or require many resources and a time-consuming analysis. In this study, we used deep learning to refine 3D gait analysis of mice after stroke. We performed markerless labeling of 10 body parts in uninjured control mice of different strains and fur color with 99% accuracy. This allowed us to describe a set of >100 biologically meaningful parameters for examining e.g., synchronization, spatial variability, and joint angles during a spontaneous walk; and that showed differential importance for acute and long-term deficits. We refined our deep learning analysis for use with the ladder rung test, which achieved outcomes comparable to manual scoring accuracy. We found that our DLC-assisted tracking approach, when benchmarked to other conventionally used behavior tests in preclinical stroke research, outperformed those based on measures of sensitivity, time-demand, and resources.

The use of machine learning approaches has dramatically increased in life sciences and will likely gain importance in the future. The introduction of DeepLabCut considerably facilitated the markerless labeling of mice and expanded the scope of kinematic tracking software^16,21^. Although commercial attempts to automate behavioral tests eliminated observer bias, the analyzed parameters are often pre-defined and cannot be altered. Especially for customized set-ups, DLC has been shown to reach human-level accuracy while outperforming commercial systems (e.g. EthoVision, TSE Multi Conditioning system) at a fraction of the cost ^19^. These advantages may become more apparent in the future since unsupervised machine learning is beginning to reveal the true complexity of animal behavior and may allow recognition of behavioral sequences not detectable by humans. On the other hand, execution and interpretation of unsupervised tracking is often beyond the reach of many basic research labs and requires the necessary machine learning knowledge^13^.

Many neurological disorders (e.g., multiple sclerosis, Huntington’s, and spinal cord injury) result in pronounced motor deficits in patients, as well as in mice models, with alterations in the general locomotor pattern. These alterations are usually readily identifiable, especially in the acute phase, and excellent automated tools have recently been developed to track the motor impairments ^30^. In contrast, deficits following cortical stroke in mice often do not reveal such clear signs of injury and require higher levels of sensitivity to identify the motor impairment^7,8,31^. The degree of functional motor deficits after stroke is highly dependent on corticospinal tract lesions that often result in specific deficits e.g., impairment of fine motor skills^32^. Moreover, a stroke most commonly affects only one body side; therefore, an experimental set-up that contains 3D information is highly valuable, as it enables the detection of contra- and ipsilateral trajectories of each anatomical landmark. Accordingly, our set-ups enabled us to also detect intra-animal differences that may be important to distinguish between normal and compensatory movements throughout the recovery time course^33^.

Compensatory strategies (e.g. avoiding the use of the impaired limb or relying on the intact limb) are highly prevalent in rodents and in humans^34,35^. Although functional recovery is generally observed in a variety of tests, it is important to distinguish between compensatory responses and “true” recovery. These mechanisms are hard to dissect in specific trained tasks (e.g., reaching during single pellet grasping). Therefore, tasks of spontaneous limb movements and many kinematic parameters are valuable to distinguish these two recovery mechanisms^25^. Interestingly, we observed alterations in several ipsilateral trajectories during the runway walk affecting the vertical positioning as well as protractive and retractive movement (although less prominent than in the contralateral paws) that suggest a compensatory movement. Similar gait alterations have been previously reported in a mouse model of distal middle cerebral artery occlusion, another common model of ischemia in mice^36^. These compensatory movements are predominantly caused by either plastic change by the adjacent areas of cortex or through support from anatomical reorganization of the contralesional hemisphere^37–39^ and, therefore, could provide valuable information about the therapeutic effects of a drug or a treatment.

Apart from general kinematic gait analysis, variations of the horizontal ladder rung/foot fault or grid tests remain one of the most reproducible tasks to assess motor skill in rodents after injury, including stroke ^31^. However, these tests often remain unused in many experimental stroke studies, most likely due to the associated time-consuming analysis. DLC-assisted refinement might allow future studies to incorporate this important assessment into their analyses given the striking decrease in time investment. We have demonstrated that DLC-assisted refinement of these conventional tests represents a striking decrease in time-consumption. Therefore, it is conceivable that some of these conventional tests and others assessment methods (e.g., single pellet grasping) may profit from the advancements of deep learning and will not be fully replaced by kinematic gait analysis.

Interestingly, some of the assessed parameters showed an impairment after stroke only in the acute phase (e.g., synchronization, cycle duration, hip movement), while some parameters showed an initial impairment after injury followed by a partial or full recovery (e.g., wrist height, toe movement, and retraction) and others showed no recovery in the time course of this study. Given the number of parameters raised in this setting, this approach might be particularly suited to assess treatment efficacy of drug interventions in preclinical stroke research. Overall, we found a strong separation of parameters in the acute vs. chronic phase in the PCA and random forest analysis, making this approach suitable to assess both the acute phase as well as the chronic phase. It will be of interest in the future to assess the presented approach in different models of stroke as well as in additional neurological conditions such as spinal cord injury, ALS, cerebral palsy or others.^25^.

Notably, a detailed kinematic analysis required optional recording settings to generate a high contrast between animal and background. In our experience these parameters needed to be adapted to the fur color in the animals. Although we reached almost equivalent tracking accuracy of 99.4% (equivalent to losing 6 in every 1’000 recorded frames), mice with black and white fur could not be tracked based on the same neural network and required two training sessions, which may show slight differences in the analysis. Moreover, the high accuracy in our experimental set-up was achieved by recording only mice with smooth runs without longer interruptions. In the future, these limitations could be overcome by combining DLC-tracking with a recently developed unsupervised clustering approach to reveal grooming or other unpredictable stops during a run^40,41^.

Taken together, in this study, we developed a comprehensive gait analysis to assess stroke impairments in mice using deep learning. The developed set-up requires minimal resources and generates characteristic multifaceted outcomes for acute and chronic phases after stroke. Moreover, we refined conventional behavioral tests used in stroke assessment at human-level accuracy that may be expanded for other behavioral tests for stroke and other neurological diseases affecting locomotion.

## Materials and Methods

### Animals

All procedures were conducted in accordance with governmental, institutional (University of Zurich), and ARRIVE guidelines and had been approved by the Veterinarian Office of the Canton of Zurich (license: 209/2019). In total 25 wildtype (WT) mice with a C57BL/6 background mice and 12 non-obese diabetic SCID gamma (NSG) mice were used (female and male, 3 months of age). Mice were housed in standard type II/III cages on a 12h day/light cycle (6:00 A.M. lights on) with food and water ad libitum. All mice were acclimatized for at least a week to environmental conditions before set into experiment.

### Photothrombotic lesion

Mice were anesthetized using isoflurane (3% induction, 1.5% maintenance, Attane, Provet AG). Analgesic (Novalgin, Sanofi) was administered 24 h prior to the start of the procedure via drinking water. A photothrombotic stroke to unilaterally lesion the sensorimotor cortex was induced on the right hemisphere, as previously described (Labat-gest and Tomasi, 2013; Rust et al., 2019b). Briefly, animals were placed in a stereotactic frame (David Kopf Instruments), the surgical area was sanitized, and the skull was exposed through a midline skin incision. A cold light source (Olympus KL 1,500LCS, 150W, 3,000K) was positioned over the right forebrain cortex (anterior/posterior: −1.5– +1.5 mm and medial/lateral 0 mm to +2 mm relative to Bregma). Rose Bengal (15 mg/ml, in 0.9% NaCl, Sigma) was injected intraperitoneally 5 min prior to illumination and the region of interest was subsequently illuminated through the intact skull for 12 min. To restrict the illuminated area, an opaque template with an opening of 3 × 4 mm was placed directly on the skull. The wound was closed using a 6/0 silk suture and animals were allowed to recover. For postoperative care, all animals received analgesics (Novalgin, Sanofi) for at least 3 days after surgery.

### Blood Perfusion by Laser Doppler Imaging

Cerebral blood flow (CBF) was measured using Laser Doppler Imaging (Moor Instruments, MOORLDI2-IR). Animals were placed in a stereotactic frame; the surgical area was sanitized and the skull was exposed through a midline skin incision. The brain was scanned using the repeat image measurement mode. All data were exported and quantified in terms of flux in the ROI using Fiji (ImageJ).

### Immunofluorescence

Brain sections were washed with 0.1M phosphate buffer (PB) and incubated with blocking solution containing donkey serum (5%) in PB for 30 min at room temperature. Sections were incubated with primary antibodies (rb-GFAP 1:200, Dako, gt-Iba1, 1:500 Wako, NeuroTrace™ 1:200, Thermo Fischer) overnight at 4°C. The next day, sections were washed and incubated with corresponding secondary antibodies (1:500, Thermo Fischer Scientific). Sections were mounted in 0.1 M PB on Superfrost PlusTM microscope slides and coverslipped using Mowiol.

### Behavioral studies

Animal were subjected to a series of behavioral tests at different time points. The here used tests included the (1) runway, (2) ladder rung test, (3) the rotarod test, (4) neurological scoring, (5) cylinder test, (6) single pallet grasping. All tests were evaluated at baseline and 3, 7, 14 and 21 after stroke induction. Animals used for deep learning-assisted tests (runway, ladder rung) represent a different cohort of animals to the remaining behavior tasks.

#### Runway test

A runway walk was performed to assess whole body coordination during overground locomotion. The walking apparatus consisted of a clear Plexiglas basin, 156 cm long, 11.5 cm wide and 11.5 cm high (Fig. 1). The basin was equipped with two ∼ 45° mirrors (perpendicularly arranged) to allow simultaneous collection of side and bottom views to generate three-dimensional tracking data. Mice were recorded crossing the runway with a high-definition video camera (GoPro Hero 7) at a resolution of 4000 × 3000 and a rate of 60 frames per second. Lighting consisted of warm background light and cool white LED stripes positioned to maximize contrast and reduce reflection. After acclimatization to the apparatus, mice were trained in two daily sessions until they crossed the runway at constant speed and voluntarily (without external intervention). Each animal was placed individually on one end of the basin and was allowed to walk for 3 minutes.

#### Ladder rung test

The same set up as in the runway was used for the ladder rung test, to assess skilled locomotion. We replaced the Plexiglas runway with a horizontal ladder (length: 113 cm, width: 7 cm, distance to ground: 15 cm). To prevent habituation to a specific bar distance, bars were irregularly spaced (1-4 cm). For behavioral testing, a total of at least three runs per animal were recorded. Kinematic analysis of both tasks was based exclusively on video recordings and only passages with similar and constant movement velocities and without lateral instability were used. A misstep was defined when the mouse toe tips reached 0.5 cm below the ladder height. The error rate was calculated by errors / total steps* 100.

#### Rotarod test

The rotarod test is a standard sensory-motor test to investigate the animals’ ability to stay and run on an accelerated rod (Ugo Basile, Gemonio, Italy). All animals were pre-trained to stay on the accelerating rotarod (slowly increasing from 5 to 50 rpm in 300s) until they could remain on the rod for > 60 s. During the performance, the time and speed was measured until the animals fell or started to rotate with the rod without running. The test was always performed three times and means were used for statistical analysis. The recovery phase between the trials was at least 10 min.

#### Neurological score / Bederson score

We used a modified version of the Bederson (0-5) score to evaluate neurological deficits after stroke. The task was adapted from Biebet et al. ^42^ The following scoring was applied; (0) no observable deficit; (1) forelimb flexion; (2) forelimb flexion and decreased resistance to lateral push; (3) circling; (4) circling and spinning around the cranial-caudal axis; (5) no spontaneous movement/ death.

#### Cylinder Test

To evaluate locomotor asymmetry mice were placed in an opentop, clear plastic cylinder for about 10 min to record their forelimb activity while rearing against the wall of the arena. The task was adapted from Roome et al. ^43^. Forelimb use is defined by the placement of the whole palm on the wall of the arena, which indicates its use for body support. Forelimb contacts while rearing were scored with a total of 20 contacts recorded for each animal. Three parameters were analysed which include paw preference, symmetry and paw dragging. Paw preference was assessed by the number of impaired forelimb contacts to the total forelimb contacts. Symmetry was calculated by the ratio of asymmetrical paw contacts to total paw contacts. Paw dragging was assessed by the ratio of the number of dragged impaired forelimb contacts to total impaired forelimb contacts.

#### Single pellet grasping

All animals were trained to reach with their right paw for 14 days prior to stroke induction over the left motor cortex. Baseline measurements were taken on the day before surgery (0dpo) and test days started at 4 dpo and were conducted weekly thereafter (7, 14, 21, 28 dpo). For the duration of behavioral training and test periods, animals were food restricted, except for 1 day prior to 3 days post injury. Body weights were kept above 80% of initial weight. The single pellet reaching task was adapted from Chen et al. ^44^. Mice were trained to reach through a 0.5-cm-wide vertical slot on the right side of the acrylic box to obtain a food pellet (Bio-Serv, Dustless Precision Pellets, 20 mg) following the guidelines of the original protocol. To motivate the mice to not drop the pellet, we additionally added a grid floor to the box, resulting in the dropped pellets to be out of reach for the animals. Mice were further trained to walk to the back of the box in between grasps to reposition themselves as well as to calm them down in between unsuccessful grasping attempts. Mice that did not successfully learn the task during the two weeks of shaping were excluded from the task (n=2). During each experiment session, the grasping success was scored for 30 reaching attempts or for a maximum of 20 minutes. Scores for the grasp were as follows: “1” for a successful grasp and retrieval of the pellet (either on first attempt or after several attempts); “0” for a complete miss in which the pellet was misplaced and not retrieved into the box; “0.5” for drag or drops, in which the animal successfully grasped the pellet but dropped it during the retrieval phase. The success rate was calculated for each animal as end score = (total score/number of attempts*100).

### DeepLabCut (DLC)

Video recordings were processed by DeepLabCut (DLC, v. 2.1.5), a computer vision algorithm that allows automatic and markerless tracking of user-defined features. A relatively small subset of camera frames (training dataset) is manually labelled as accurately as possible (for each task and strain, respectively). Those frames are then used to train an artificial network. Once the network is sufficiently trained, different videos can then be directly input to DLC for automated tracking. The network predicts the marker locations in the new videos, and the 2D points can be further processed for performance evaluation and 3D reconstruction.

#### Dataset preparation

Each video was migrated to Adobe Premiere (v. 15.4) and optimized for image quality (color correction and sharpness). Videos were split into short one-run-sequences (left to right or right to left), cropped to remove background and exported/compressed in H.264 format.

#### Training

The general networks for both behavioral tests were trained based on ResNet-50 by manually labelling 120 frames selected using k-means clustering from multiple videos of different mice (N = 6 videos/network). We labelled 10 distinct body parts (head, front toe tip, wrist, shoulder, elbow, back toe, back ankle, iliac crest, hip, tail) in all videos of mice recorded from side views (left, right) and 8 body parts (head, right front toe, left front toe, center front, right back toe, left back toe, center back, tail base) in all videos showing the bottom view, respectively (for details, see Suppl. Fig. 1, 2). Using these hand labelled data, we allowed training to run for 1’030’000 iterations (DLC’s native cross-entropy loss function plateaued between 100’000 and 300’000 iterations). Evaluation of labelling accuracy was assessed using the ‘evaluate network’ function provided by DLC. This function computes the Euclidean error between the manual labels and the ones predicted by DLC averaged over the hand locations and test images (mean absolute error, proportional to the average root mean square error).

#### Refinement

20 outlier frames from each of the training videos where manually corrected and then added to the training dataset. Locations with a p <0.9 were relabeled. The network was then refined using the same numbers of iterations (1’030’000).

All experiments were performed inside the Anaconda environment (Python 3.7.8) provided by DLC using NVIDIA GeForce RTX 2060.

### Data processing with R

Video pixel coordinates for the labels produced by DLC were imported into R Studio (Version 4.04 (2021-02-15) and processed with custom scrips that can be assessed here: https://github.com/rustlab1/DLC-Gait-Analysis. Briefly, accuracy values of individual videos were evaluated and data points with a low likelihood were removed. Representative videos were chosen to plot a general overview of the gait. Next, individual steps were identified within the run by the speed of the paws to identify the “stance” and “swing” phase. These steps were analyzed for synchronization, speed, length, and duration from the down view over a time course. Additionally, the angular positioning between the body center and the individual paws was measured. From the lateral/side view, we next measured average and total height differences of individual joins (y-coordinates) and the total movement, protraction, and retraction changes per step (x-coordinates) over the time course. Next, we measured angular variability (max, average, min) between neighboring joints including (hip-ankle-toe, iliac crest-hip-back-ankle, elbow-wrist-front toe, shoulder-elbow-wrist).

All >100 generated parameters were extracted to perform a random forest classification (ntree = 100) to determine the importance (Gini impurity-based feature importance) for determining accuracy of the injury status. This was performed for all time points (bl, 3, 7, 14,21 dpi) and in a subgroup analysis between baseline vs. 3 dpi and baseline vs. 21 dpi. A confusion matrix was performed for determining the prediction accuracy of this model. The most five important parameters were used to perform a principal component analysis to demonstrate separation of these parameters.

### Statistical Analysis

Statistical analysis was performed using RStudio (4.04 (2021-02-15). Sample sizes were designed with adequate power according to our previous studies ^7,28,45^ and to the literature ^8,11^. Overview of sample sizes can be found in Suppl. Table 1. All data were tested for normal distribution by using the Shapiro-Wilk test. Normally distributed data were tested for differences with a two-tailed unpaired one-sample *t*-test to compare changes between two groups (differences between ipsi- and contralesional sides). Multiple comparisons were initially tested for normal distribution with the Shapiro-Wilk test. The significance of mean differences between normally distributed multiple comparisons was assessed using repeated measures ANOVA with post-hoc analysis (p adjustment method = holm). Variables exhibiting a skewed distribution were transformed, using natural logarithms before the tests to satisfy the prerequisite assumptions of normality. An overview of all raw data and tests can be found in Suppl. Table 3. Data are expressed as means ± SD, and statistical significance was defined as \**p* < 0.05, \*\**p* < 0.01, and \*\*\**p* < 0.001. Raw data, summarized data and statistical evaluation can be found in the supplementary information (Suppl. Table 2-37).

### Code availability

The code with sample data are available at https://github.com/rustlab1/DLC-Gait-Analysis

