## Supplementary material for "Deep learning based behavioral profiling of rodent stroke recovery": Table 1

**Table 1: Name and description of all generated parameters following runway performance**

| No | Prameter Cluster | Parameter Name | Parameter Description |
|---|---|---|---|
| 1 | synchronization and temporal features | average duration | average duration of a total step with all four paws, (s) |
| 2 | | average stance time | average stance time during a step, (s) |
| 3 | | average swing time | average swing time during a step, (s) |
| 4 | | left back duration | average duration of a step in the left back paw, (s) |
| 5 | | left front duration | average duration of a step in the left front paw, (s) |
| 6 | | right back duration | average duration of a step in the right back paw, (s) |
| 7 | | right front duration | average duration of a step in the right front paw, (s) |
| 8 | | average stride length | average length of a step from down view, (mm) |
| 9 | | left front back right synchronization | asynchronization ratio between left front paw and right back paw, 0 = complete symmetry, normal gait (-) |
| 10 | | right front back left synchronization | asynchronization ratio between right front paw and left back paw, 0 = complete symmetry, normal gait (-) |
| 11 | vertical joint movement and heights | left back ankle average height | average height changes during a step of the left back ankle compared to baseline = 0 , (mm) |
| 12 | | left back ankle vertical movement | total vertical movement changes during a step of the left back ankle compared to baseline = 0, (mm) |
| 13 | | left back toe tip average height | average height changes during a step of the left back toe tip compared to baseline = 0, (mm) |
| 14 | | left back toe tip vertical movement | total vertical movement changes during a step of the left back toe tip compared to baseline = 0, (mm) |
| 15 | | left front toe tip average height | average height changes during a step of the left front toe tip compared to baseline = 0, (mm) |
| 16 | | left front toe tip vertical movement | total vertical movement changes during a step of the left front toe tip compared to baseline = 0, (mm) |
| 17 | | left head average height | average height changes during a step of the head from left side compared to baseline = 0, (mm) |
| 18 | | left head vertical movement | total vertical movement changes during a step of the head from left side compared to baseline = 0, (mm) |
| 19 | | left hip average height | average height changes during a step of the left hip compared to baseline = 0, (mm) |
| 20 | | left hip vertical movement | total vertical movement changes during a step of the left hip compared to baseline = 0, (mm) |
| 21 | | left iliac crest average height | average height changes during a step of the left iliac crest compared to baseline = 0, (mm) |
| 22 | | left iliac crest vertical movement | total vertical movement changes during a step of the left iliac crest compared to baseline = 0, (mm) |
| 23 | | left tail base average height | average height changes during a step of the left tail base compared to baseline = 0, (mm) |
| 24 | | left tail base vertical movement | total vertical movement changes during a step of the left tail base compared to baseline = 0, (mm) |
| 25 | | left wrist average height | average height changes during a step of the left wrist compared to baseline = 0, (mm) |
| 26 | | left wrist vertical movement | total vertical movement changes during a step of the left wrist compared to baseline = 0, (mm) |

| | | | |
|---|---|---|---|
| 27 | vertical joint movement and heights | right back ankle average height | average height changes during a step of the right back ankle compared to baseline = 0, (mm) |
| 28 | | right back ankle vertical movement | total vertical movement changes during a step of the left back ankle compared to baseline = 0, (mm) |
| 29 | | right back toe tip average height | average height changes during a step of the right back toe tip compared to baseline = 0, (mm) |
| 30 | | right back toe tip vertical movement | total vertical movement changes during a step of the right back toe tip compared to baseline = 0, (mm) |
| 31 | | right front toe tip average height | average height changes during a step of the right front toe tip compared to baseline = 0, (mm) |
| 32 | | right front toe tip vertical movement | total vertical movement changes during a step of the right front toe tip compared to baseline = 0, (mm) |
| 33 | | right head average height | average height changes during a step of the head from right side compared to baseline = 0, (mm) |
| 34 | | right head vertical movement | total vertical movement changes during a step of the head from right side compared to baseline = 0, (mm) |
| 35 | | right hip average height | average height changes during a step of the right hip compared to baseline = 0, (mm) |
| 36 | | right hip vertical movement | total vertical movement changes during a step of the right hip compared to baseline = 0, (mm) |
| 37 | | right iliac crest average height | average height changes during a step of the right iliac crest compared to baseline = 0, (mm) |
| 38 | | right iliac crest vertical movement | total vertical movement changes during a step of the right iliac crest compared to baseline = 0, (mm) |
| 39 | | right tail base average height | average height changes during a step of the right tail base compared to baseline = 0, (mm) |
| 40 | | right tail base vertical movement | total vertical movement changes during a step of the right tail base compared to baseline = 0, (mm) |
| 41 | | right wrist average height | average height changes during a step of the right wrist compared to baseline = 0, (mm) |
| 42 | | right wrist vertical movement | total vertical movement changes during a step of the right wrist compared to baseline = 0, (mm) |
| 43 | step length, protraction and retraction | left back average length | average length of a step of the left back paw compared to baseline = 0, (mm) |
| 44 | | left back median length | median length of a step of the left back paw compared to baseline = 0, (mm) |
| 45 | | left back total horizontal movement | total horizontal movement during a step of the left back paw compared to baseline = 0, (mm) |
| 46 | | left back protraction max | maximum protractive movement during a step of the left back paw compared to baseline = 0, (mm) |
| 47 | | left back retraction max | maximum retractive movement during a step of the left back paw compared to baseline = 0, (mm) |
| 48 | | left front average length | average length of a step of the left front paw compared to baseline = 0, (mm) |
| 49 | | left front median length | median length of a step of the left front paw compared to baseline = 0, (mm) |
| 50 | | left front total horizontal movement | total horizontal movement during a step of the left front paw compared to baseline = 0, (mm) |
| 51 | | left front protraction max | maximum protractive movement during a step of the left front paw compared to baseline = 0, (mm) |
| 52 | | left front retraction max | maximum retractive movement during a step of the left front paw compared to baseline = 0, (mm) |
| 53 | | right back average length | average length of a step of the right back paw compared to baseline = 0, (mm) |
| 54 | | right back median length | median length of a step of the right back paw compared to baseline = 0, (mm) |

| | | | |
|---|---|---|---|
| 55 | | right back total horizontal movement | total horizontal movement during a step of the right back paw compared to baseline = 0, (mm) |
| 56 | | right back protraction max | maximum protractive movement during a step of the right back paw compared to baseline = 0, (mm) |
| 57 | | right back retraction max | maximum retractive movement during a step of the right back paw compared to baseline = 0, (mm) |
| 58 | | right front average length | average length of a step of the right front paw compared to baseline = 0, (mm) |
| 59 | | right front median length | median length of a step of the right front paw compared to baseline = 0, (mm) |
| 60 | | right front total horizontal movement | total horizontal movement during a step of the right front paw compared to baseline = 0, (mm) |
| 61 | | right front protraction max | maximum protractive movement during a step of the right front paw compared to baseline = 0, (mm) |
| 62 | step length, protraction and retraction | right front retraction max | maximum retractive movement during a step of the right front paw compared to baseline = 0, (mm) |
| 63 | | left hip total horizontal movement | total horizontal movement during a step of the left hip compared to baseline = 0, (mm) |
| 64 | | left hip average length | average distance covered by the left hip during a step compared to baseline = 0, (mm) |
| 64 | | left tail base total horizontal movement | total horizontal movement during a step of the left tail base compared to baseline = 0, (mm) |
| 64 | | left tail base average length | average distance covered by the left tail base during a step compared to baseline = 0, (mm) |
| 64 | | right hip total horizontal movement | total horizontal movement during a step of the right hip compared to baseline = 0, (mm) |
| 64 | | right hip average length | average distance covered by the right hip during a step compared to baseline = 0, (mm) |
| 64 | | right tail base total horizontal movement | total horizontal movement during a step of the right tail base compared to baseline = 0, (mm) |
| 64 | | right tail base average length | average distance covered by the right tail base during a step compared to baseline = 0, (mm) |
| 64 | | left back average angle | average angle between body center and left back paw during a step from down view, (°) |
| 64 | | left back max angle | maximum angle between body center and left back paw during a step from down view, (°) |
| 64 | | left back min angle | minimum angle between body center and left back paw during a step from down view, (°) |
| 64 | | left front average angle | average angle between body center and left front paw during a step from down view, (°) |
| 64 | | left front max angle | maximum angle between body center and left front paw during a step from down view, (°) |
| 64 | paw to body center angles | left front min angle | minimum angle between body center and left front paw during a step from down view, (°) |
| 64 | | right back average angle | average angle between body center and right back paw during a step from down view, (°) |
| 64 | | right back max angle | maximum angle between body center and right back paw during a step from down view, (°) |
| 64 | | right back min angle | minimum angle between body center and right back paw during a step from down view, (°) |
| 64 | | right front average angle | average angle between body center and right front paw during a step from down view, (°) |
| 64 | | right front max angle | maximum angle between body center and right front paw during a step from down view, (°) |
| 64 | | right front min angle | minimum angle between body center and right front paw during a step from down view, (°) |

| | | | |
|---|---|---|---|
| 64 | | average angle left back hip ankle toe | average angle between left hip, left back ankle and left back toe tip during a step from side view, (°) |
| 64 | | average angle left back iliac hip ankle | average angle between left iliac crest, left hip and left back ankle during a step from side view, (°) |
| 64 | | average angle left front elbow wrist toetip | average angle between left elbow, left wrist and left front toe tip during a step from side view, (°) |
| 64 | | average angle left front shoulder elbow wrist | average angle between left shuolder, left elbow and left wrist during a step from side view, (°) |
| 64 | | max left back hip ankle toe | maximum angle between left hip, left back ankle and left back toe tip during a step from side view, (°) |
| 64 | | max left back iliac hip ankle | maximum angle between left iliac crest, left hip and left back ankle during a step from side view, (°) |
| 64 | | max left front elbow wrist toetip | maximum angle between left elbow, left wrist and left front toe tip during a step from side view, (°) |
| 64 | | max left front shoulder elbow wrist | maximum angle between left shuolder, left elbow and left wrist during a step from side view, (°) |
| 64 | | min left back hip ankle toe | minimum angle between left hip, left back ankle and left back toe tip during a step from side view, (°) |
| 64 | | min left back iliac hip ankle | minimum angle between left iliac crest, left hip and left back ankle during a step from side view, (°) |
| 64 | | min left front elbow wrist toetip | minimum angle between left elbow, left wrist and left front toe tip during a step from side view, (°) |
| 64 | joint angles | min left front shoulder elbow wrist | minimum angle between left shuolder, left elbow and left wrist during a step from side view, (°) |
| 64 | | average angle right back hip ankle toe | average angle between right hip, right back ankle and right back toe tip during a step from side view, (°) |
| 64 | | average angle right back iliac hip ankle | average angle between right iliac crest, right hip and right back ankle during a step from side view, (°) |
| 64 | | average angle right front elbow wrist toetip | average angle between right elbow, right wrist and right front toe tip during a step from side view, (°) |
| 64 | | average angle right front shoulder elbow wrist | average angle between right shuolder, right elbow and right wrist during a step from side view, (°) |
| 64 | | max right back hip ankle toe | maximum angle between right hip, right back ankle and right back toe tip during a step from side view, (°) |
| 64 | | max right back iliac hip ankle | maximum angle between right iliac crest, right hip and right back ankle during a step from side view, (°) |
| 64 | | max right front elbow wrist toetip | maximum angle between right elbow, right wrist and right front toe tip during a step from side view, (°) |
| 64 | | max right front shoulder elbow wrist | maximum angle between right shuolder, right elbow and right wrist during a step from side view, (°) |
| 64 | | min right back hip ankle toe | minimum angle between right hip, right back ankle and right back toe tip during a step from side view, (°) |
| 64 | | min right back iliac hip ankle | minimum angle between right iliac crest, right hip and right back ankle during a step from side view, (°) |
| 64 | | min right front elbow wrist toetip | minimum angle between right elbow, right wrist and right front toe tip during a step from side view, (°) |
| 64 | | min right front shoulder elbow wrist | minimum angle between right shuolder, right elbow and right wrist during a step from side view, (°) |
