## Supplementary Figures for "Deep learning based behavioral profiling of rodent stroke recovery"

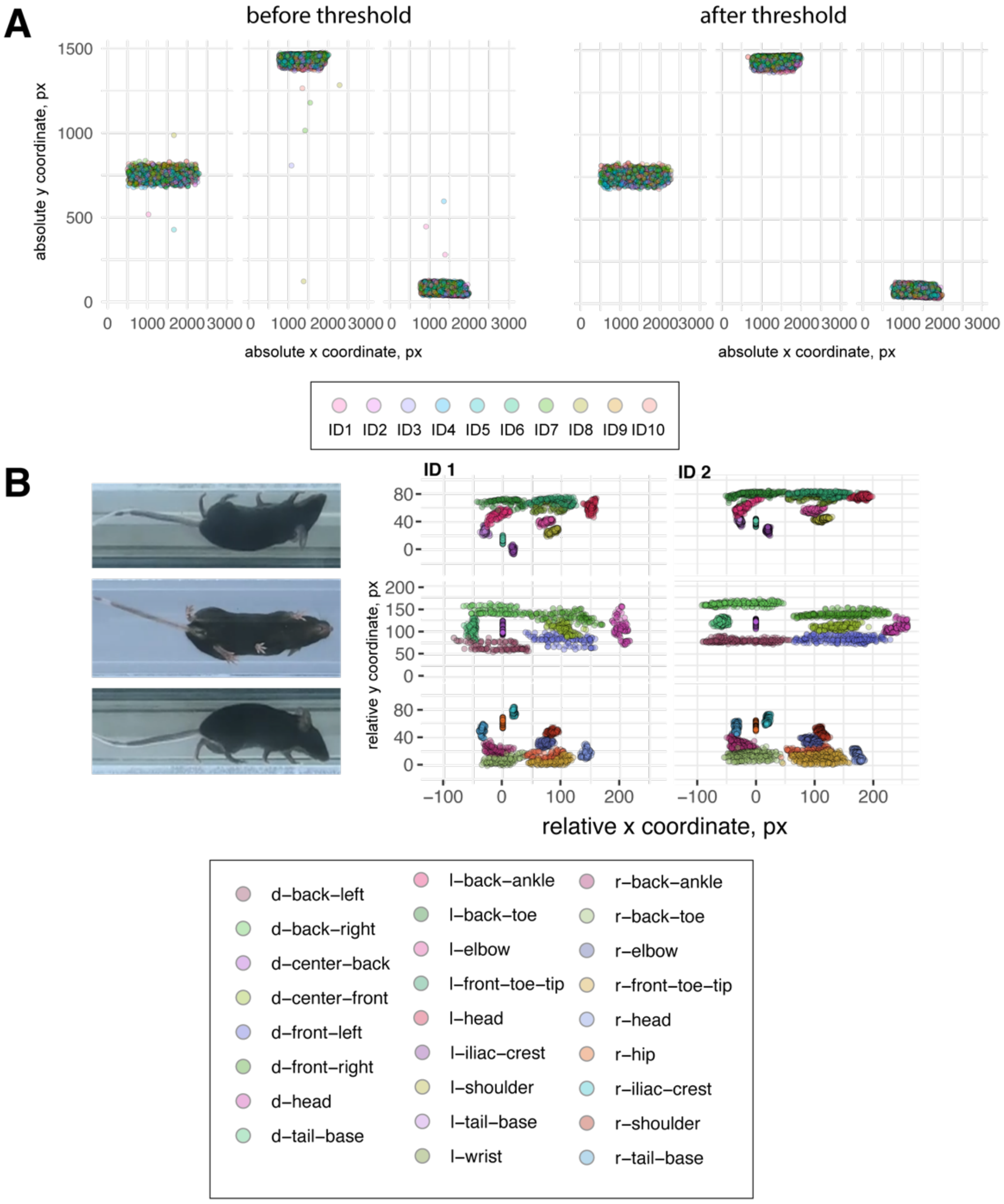


**Suppl. Fig. 1: Walking profile of mice following DeepLabCut tracking.** (**A**) Randomly sampled raw data plot of original x and y coordinates before pre-processing and after pre-processing. (**B**) Walking profile normalized to the hip coordinates of randomly selected non-injured mice (ID 1, ID2). Each dot represents an anatomical landmark.

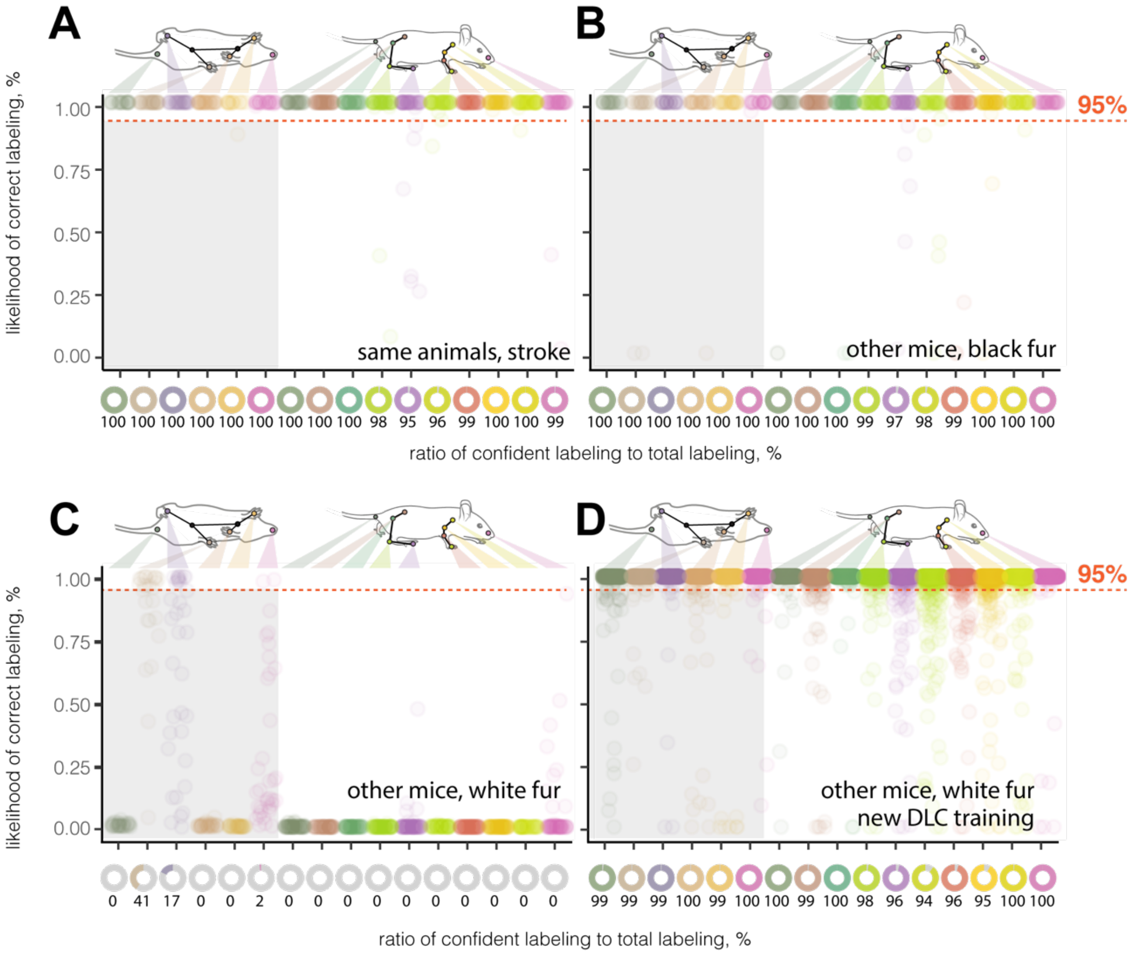


**Suppl. Fig. 2: Tracking of body parts in injured mice and mice with different genotypes. (A)** Likelihood of a confident labeling for individual body parts in the runway in stroked mice **(B)** stains with a different genotype but same (black) fur color and **(C)** different genotype but white fur color **(D)** Likelihood of a confident labeling for new training set in mice with different genotype and white fur. Each dot represents an anatomical landmark of individual image frames in a video. The red dotted line represents the confidence threshold of 95% likelihood for reliable labeling.

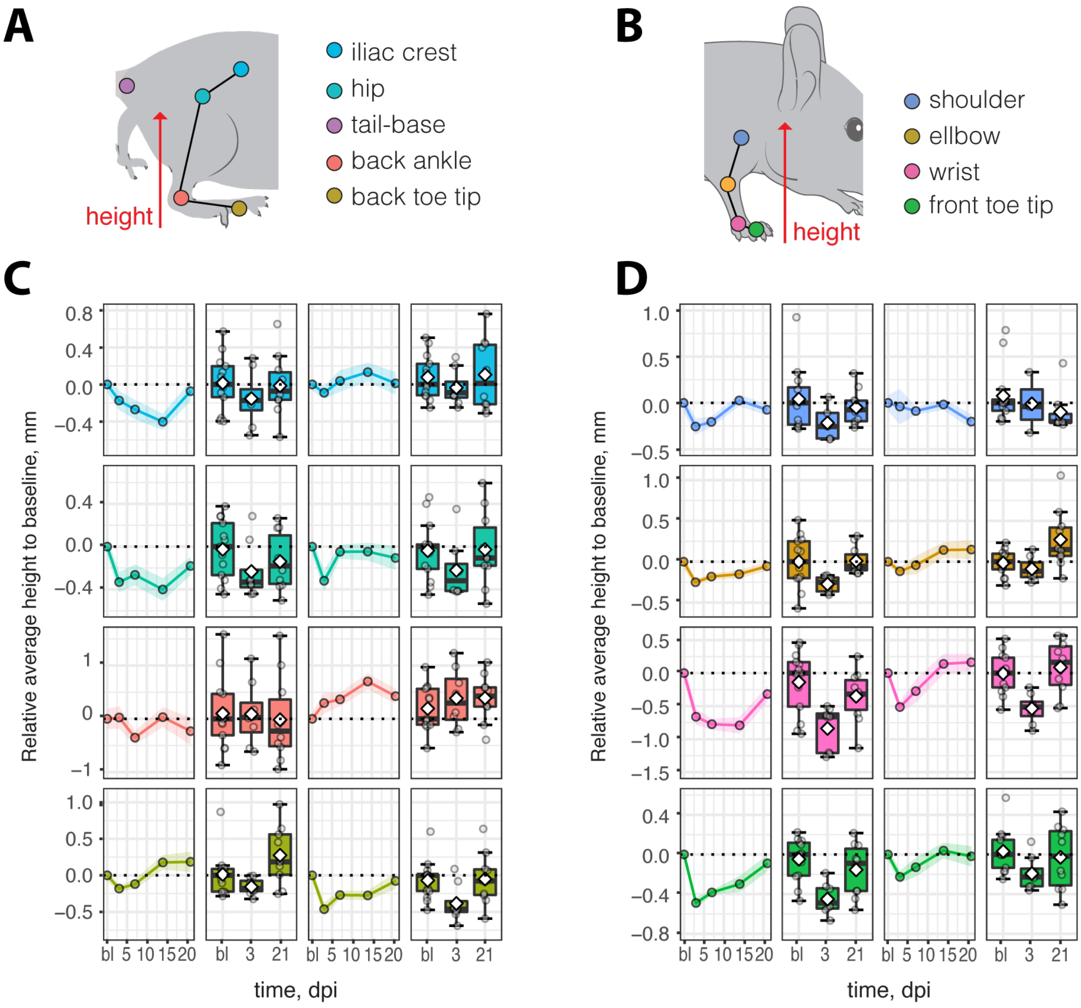


**Suppl. Fig 3: Kinematic changes in spontaneous walk after stroke**. (A, B) Schematic overview of tracked parameters (C, D) Average height of selected joints at 3,7, 14, 21 dpi compared to baseline. Data are shown as mean distributions where the white dot represents the mean. Boxplots indicate the 25% to 75% quartiles of the data. For boxplots: each dot in the plots represents one animal. Line graphs are plotted as mean ± sem. For line graphs: the dots represent the mean of the data. Significance of mean differences between the groups was assessed using Tukey's HSD. Asterisks indicate significance: $^{*}$ $P < 0.05$, $^{**}$ $P < 0.01$, $^{***}$ $P < 0.001$

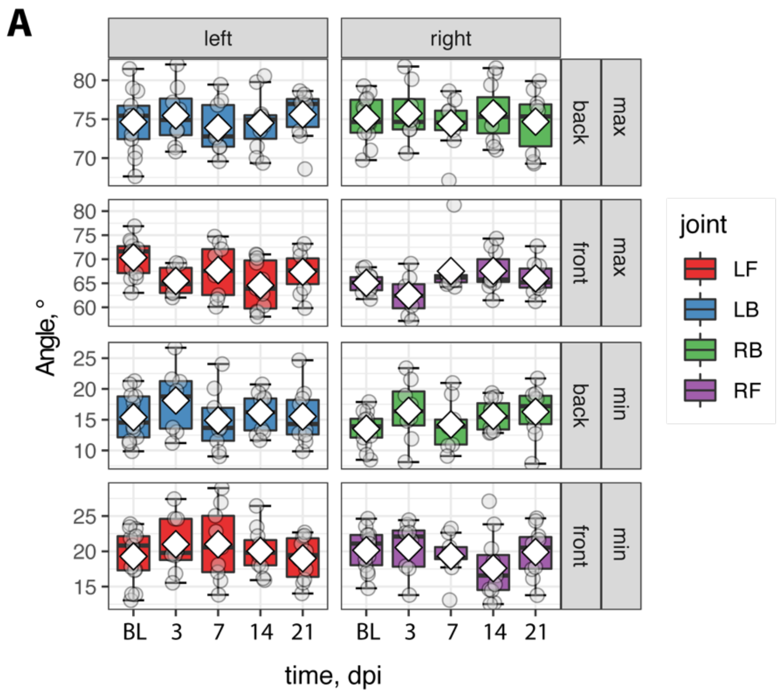


**Suppl. Fig 4: Angular variability between body center and front and hind paws**. (A) Comparison of angles of individual paws to body center in a time course. Data are shown as mean distributions where the white dot represents the mean. Boxplots indicate the 25% to 75% quartiles of the data. Each dot in the plots represents one animal and significance of mean differences between the groups was assessed using Tukey's HSD.

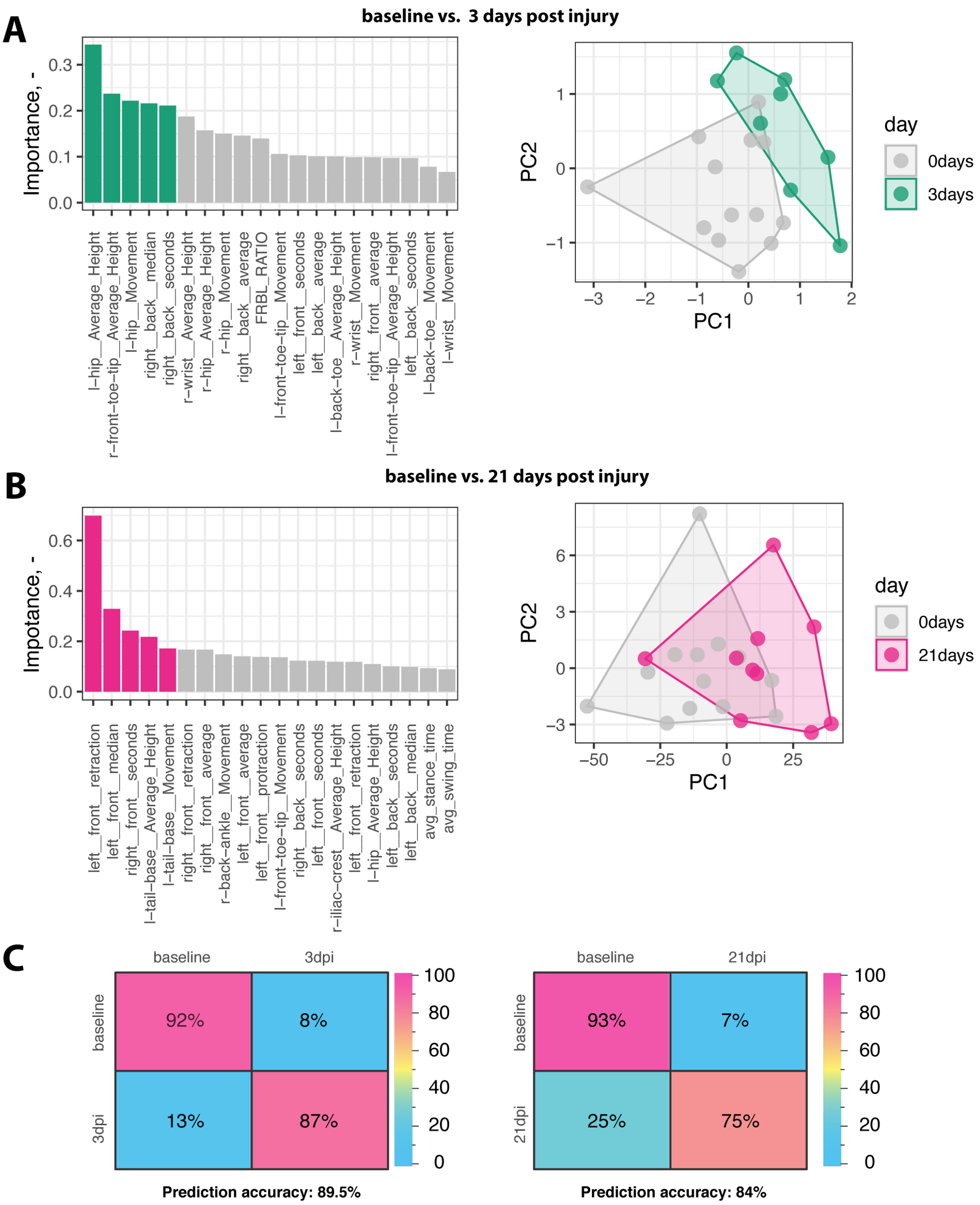


**Suppl. Fig 5: Subgroup analysis for random forest classification and principal component analysis. (A)** Random Forest classification of most important parameters (Gini impurity-based feature importance) between baseline and 3 dpi and principal component plot. (**B**) Random Forest classification of most important parameters Gini impurity-based feature importance between baseline and 21 dpi and principal component plot. (C) Confusion matrix and prediction accuracy of these models. Each dot in principal component analysis represents a video of individual animals.

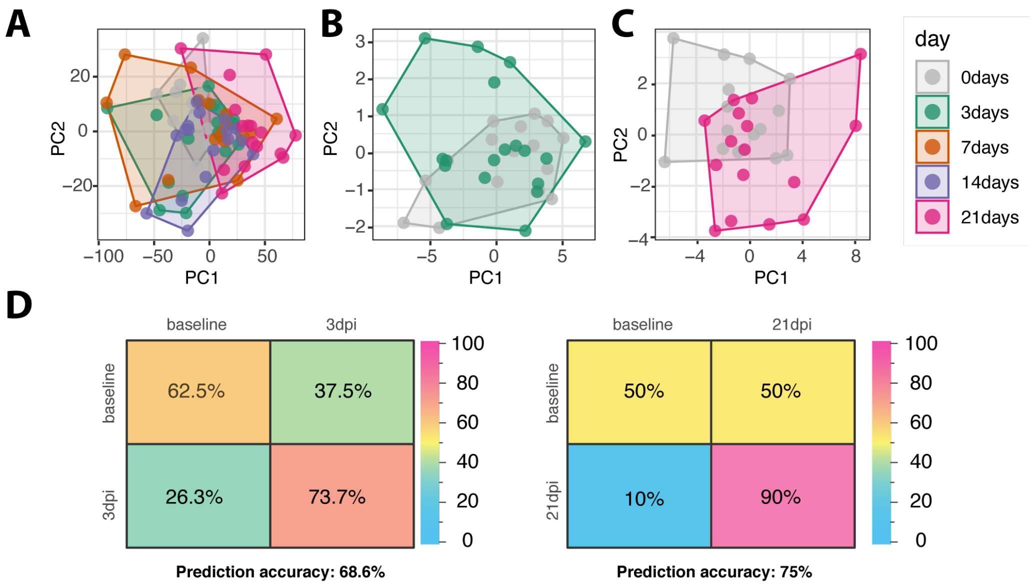


**Suppl. Fig 6: Principal component analysis and random forest classification of uninjured control mice.** (**A**) Principal component analysis of intact mice at day 0, 3, 7, 14, 21 and subgroup analysis of (B) 0 days and 3 days or (**C**) 0 days and 21 days with parameters previously generated from stroked mice. (**D**) Confusion matrix and prediction accuracy of a random forest classification in uninjured mice from 0 days and 3 days (left) and 0 days and 21 days (right). Each dot in principal component analysis represents a video of individual animals.

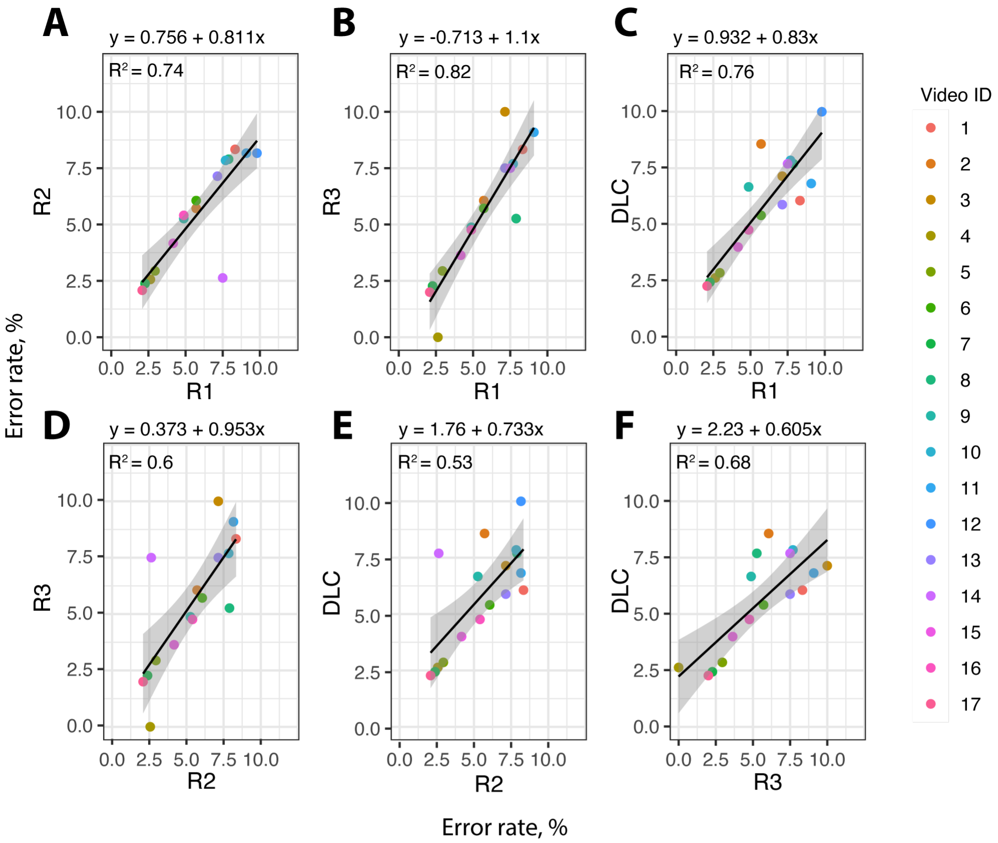


**Suppl. Fig 7: Correlation plots between human annotators and DeepLabCut evaluation of the ladder rung test**. (A) Correlation plot between Rater 1 and Rater 2. (B) Correlation plot between rater 1 and rater 3. (C) Correlation plot between rater 1 and rater 3. (D) Correlation plot between rater 2 and rater 3. (E) Correlation plot between DLC and rater 2. (F) Correlation plot between rater 3 and rater DLC. Individual dots represent randomly selected videos of both injured and non-injured mice.

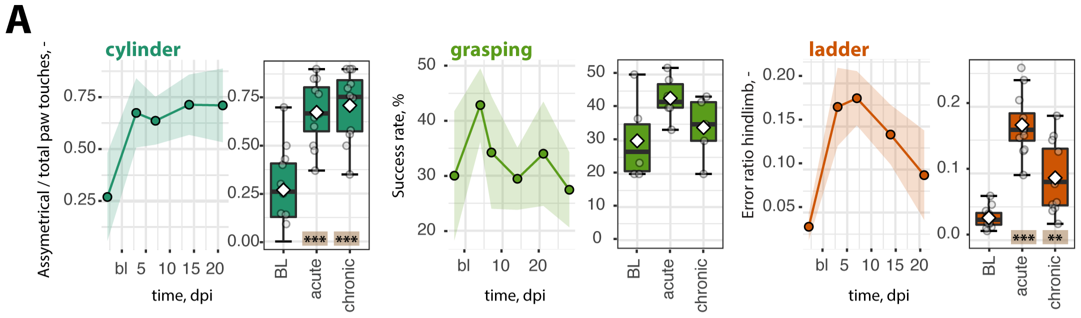


**B**

| Parameter \ Criteria | neurol. score | rota-rod | cylinder test | pellet grasping | ladder rung test | ladder + DLC | runway test | runway + DLC |
|---|---|---|---|---|---|---|---|---|
| duration to perform task | 10 | 8 | 7 | 1 | 4 | 4 | 3 | 8 |
| sensitivity of task | 2 | 2 | 5 | 8 | 6 | 6 | 10 | 10 |
| # of readouts | 1 | 2 | 3 | 5 | 3 | 3 | 10 | 10 |
| objectivity / reproducibility | 1 | 3 | 3 | 3 | 5 | 9.5 | 6 | 6 |
| detection of long-term deficits | 0 | 0 | 3 | 10 | 8 | 8 | 10 | 10 |
| duration of post-hoc analysis | 10 | 5 | 2 | 3 | 3 | 7 | 1 | 10 |
| intensity of pre-training | 10 | 3 | 7 | 1 | 5 | 5 | 9 | 9 |
| low costs for set-up | 10 | 8 | 9.5 | 9 | 8 | 8 | 0.5 | 8 |

**C**

| Parameter \ Criteria | low (0-2) | average (3-7) | high (8-10) |
|---|---|---|---|
| duration to perform task | > 10 min / mouse | 2 - 10 min / mouse | < 2 min / mouse |
| sensitivity of task | obvious deficits, only | moderate deficits | fine-motor deficits |
| # of readouts | 1 readout | 1-5 readout(s) | > 5 readouts |
| objectivity / reproducibility | highly variable, user-dependent | requires precise guidelines to ensure reproducibility | user-independent |
| detection of long-term deficits | no | only in big strokes, some animals | yes |
| duration of post-hoc analysis | > 10 min / mouse | 2 - 10 min / mouse | < 2 min / mouse |
| intensity of pre-training | days-weeks of pre-training | several, short pre-trainings | no pre-training required |
| low costs for set-up | > 1500 USD | 500 - 1500 USD | < 500 USD |

**Suppl. Fig 8: Functional assessment of recovery after stroke using conventional behavioral tests** (**A**) Time course of recovery after stroke for asymmetry in the cylinder test (left), successful grasp and retrieval of the pellet in the single pellet grasping task (middle) and hindlimb errors in the ladder rung test (right). (**B**) Semi-quantitative scoring of assessment criteria including duration, sensitivity, number of readouts, reproducibility, detection of long-term deficits, duration of post-hoc analysis, intensity of pre-training and overall costs. (**C**) Description of scoring criteria for behavioral test comparison.
