## Supplementary Tables for "Deep learning based behavioral profiling of rodent stroke recovery"

**Suppl. Table 1: Overview of sample size for each experiment**

| Overview | Desction of experiment | Sample size N |
|---|---|---|
| **Validation of DeepLabCut set-up** | C57Bl/6 (1st batch) for training | 12 |
| | C57Bl/6 (2nd batch) for validation | 8 |
| | NSG (white fur) | 14 |
| **Stroke confirmation and tissue analysis** | Laser Doppler Imaging | 7-8 |
| | Stroke size analysis | 7 |
| **DeepLabCut analysis** | runway analysis | 8-13 |
| | ladder rung analysis | 8-13 |
| **Conventional tests** | Neurological score | 7 |
| | Rotarod | 12 |
| | Cylinder test | 12 |
| | ladder rung test | 12 |
| | single pellet grasping | 6 |

**Suppl. Table 2: Accuracy of labeling different mice during runway test**

| Group | Parameter | Likeliehood | Training Session | Setup |
|---|---|---|---|---|
| Bl6, original mice | d-back-left | 100 | Original Session 1 | runway |
| Bl6, original mice | d-back-right | 100 | Original Session 1 | runway |
| Bl6, original mice | d-front-left | 100 | Original Session 1 | runway |
| Bl6, original mice | d-front-right | 100 | Original Session 1 | runway |
| Bl6, original mice | d-head | 100 | Original Session 1 | runway |
| Bl6, original mice | d-tail-base | 100 | Original Session 1 | runway |
| Bl6, original mice | l-back-ankle | 96.77419355 | Original Session 1 | runway |
| Bl6, original mice | l-back-toe | 93.63057325 | Original Session 1 | runway |
| Bl6, original mice | l-elbow | 100 | Original Session 1 | runway |
| Bl6, original mice | l-front-toe-tip | 98.125 | Original Session 1 | runway |
| Bl6, original mice | l-head | 100 | Original Session 1 | runway |
| Bl6, original mice | l-hip | 100 | Original Session 1 | runway |
| Bl6, original mice | l-iliac-crest | 100 | Original Session 1 | runway |
| Bl6, original mice | l-shoulder | 99.35483871 | Original Session 1 | runway |
| Bl6, original mice | l-tail-base | 100 | Original Session 1 | runway |
| Bl6, original mice | l-wrist | 100 | Original Session 1 | runway |
| Bl6, original mice | r-back-ankle | 99.35483871 | Original Session 1 | runway |
| Bl6, original mice | r-back-toe | 100 | Original Session 1 | runway |
| Bl6, original mice | r-elbow | 100 | Original Session 1 | runway |
| Bl6, original mice | r-front-toe-tip | 96.73202614 | Original Session 1 | runway |
| Bl6, original mice | r-head | 98.03921569 | Original Session 1 | runway |
| Bl6, original mice | r-hip | 100 | Original Session 1 | runway |
| Bl6, original mice | r-iliac-crest | 100 | Original Session 1 | runway |
| Bl6, original mice | r-shoulder | 100 | Original Session 1 | runway |
| Bl6, original mice | r-tail-base | 100 | Original Session 1 | runway |
| Bl6, original mice | r-wrist | 99.35897436 | Original Session 1 | runway |
| Bl6, new mice | d-back-left | 100 | Original Session 1 | runway |
| Bl6, new mice | d-back-right | 100 | Original Session 1 | runway |
| Bl6, new mice | d-front-left | 100 | Original Session 1 | runway |
| Bl6, new mice | d-front-right | 100 | Original Session 1 | runway |
| Bl6, new mice | d-head | 100 | Original Session 1 | runway |
| Bl6, new mice | d-tail-base | 100 | Original Session 1 | runway |
| Bl6, new mice | l-back-ankle | 97.63313609 | Original Session 1 | runway |
| Bl6, new mice | l-back-toe | 92.89940828 | Original Session 1 | runway |
| Bl6, new mice | l-elbow | 100 | Original Session 1 | runway |
| Bl6, new mice | l-front-toe-tip | 98.25581395 | Original Session 1 | runway |
| Bl6, new mice | l-head | 100 | Original Session 1 | runway |
| Bl6, new mice | l-hip | 100 | Original Session 1 | runway |
| Bl6, new mice | l-iliac-crest | 100 | Original Session 1 | runway |
| Bl6, new mice | l-shoulder | 100 | Original Session 1 | runway |
| Bl6, new mice | l-tail-base | 100 | Original Session 1 | runway |
| Bl6, new mice | l-wrist | 100 | Original Session 1 | runway |
| Bl6, new mice | r-back-ankle | 100 | Original Session 1 | runway |
| Bl6, new mice | r-back-toe | 100 | Original Session 1 | runway |
| Bl6, new mice | r-elbow | 100 | Original Session 1 | runway |
| Bl6, new mice | r-front-toe-tip | 96.49122807 | Original Session 1 | runway |

| | | | | |
|---|---|---|---|---|
| Bl6, new mice | r-head | 100 | Original Session 1 | runway |
| Bl6, new mice | r-hip | 100 | Original Session 1 | runway |
| Bl6, new mice | r-iliac-crest | 100 | Original Session 1 | runway |
| Bl6, new mice | r-shoulder | 100 | Original Session 1 | runway |
| Bl6, new mice | r-tail-base | 100 | Original Session 1 | runway |
| Bl6, new mice | r-wrist | 99.42196532 | Original Session 1 | runway |
| NOD, white fur | d-back-left | 41.11111111 | Original Session 1 | runway |
| NOD, white fur | d-back-right | 17.87709497 | Original Session 1 | runway |
| NOD, white fur | d-front-left | 0 | Original Session 1 | runway |
| NOD, white fur | d-front-right | 0 | Original Session 1 | runway |
| NOD, white fur | d-head | 1.886792453 | Original Session 1 | runway |
| NOD, white fur | d-tail-base | 0 | Original Session 1 | runway |
| NOD, white fur | l-back-ankle | 0 | Original Session 1 | runway |
| NOD, white fur | l-back-toe | 0 | Original Session 1 | runway |
| NOD, white fur | l-elbow | 0 | Original Session 1 | runway |
| NOD, white fur | l-front-toe-tip | 0 | Original Session 1 | runway |
| NOD, white fur | l-head | 0 | Original Session 1 | runway |
| NOD, white fur | l-hip | 0 | Original Session 1 | runway |
| NOD, white fur | l-iliac-crest | 0 | Original Session 1 | runway |
| NOD, white fur | l-shoulder | 0 | Original Session 1 | runway |
| NOD, white fur | l-tail-base | 0 | Original Session 1 | runway |
| NOD, white fur | l-wrist | 0 | Original Session 1 | runway |
| NOD, white fur | r-back-ankle | 0 | Original Session 1 | runway |
| NOD, white fur | r-back-toe | 0 | Original Session 1 | runway |
| NOD, white fur | r-elbow | 0 | Original Session 1 | runway |
| NOD, white fur | r-front-toe-tip | 0 | Original Session 1 | runway |
| NOD, white fur | r-head | 0 | Original Session 1 | runway |
| NOD, white fur | r-hip | 0 | Original Session 1 | runway |
| NOD, white fur | r-iliac-crest | 0 | Original Session 1 | runway |
| NOD, white fur | r-shoulder | 0 | Original Session 1 | runway |
| NOD, white fur | r-tail-base | 0 | Original Session 1 | runway |
| NOD, white fur | r-wrist | 0 | Original Session 1 | runway |
| NOD, white fur | d-back-left | 99.25925926 | Modified Session 2 | runway |
| NOD, white fur | d-back-right | 99.25925926 | Modified Session 2 | runway |
| NOD, white fur | d-front-left | 99.23664122 | Modified Session 2 | runway |
| NOD, white fur | d-front-right | 99.24242424 | Modified Session 2 | runway |
| NOD, white fur | d-head | 100 | Modified Session 2 | runway |
| NOD, white fur | d-tail-base | 98.48484848 | Modified Session 2 | runway |
| NOD, white fur | l-back-ankle | 99.25925926 | Modified Session 2 | runway |
| NOD, white fur | l-back-toe | 95.45454545 | Modified Session 2 | runway |
| NOD, white fur | l-elbow | 95.45454545 | Modified Session 2 | runway |
| NOD, white fur | l-front-toe-tip | 96.92307692 | Modified Session 2 | runway |
| NOD, white fur | l-head | 100 | Modified Session 2 | runway |
| NOD, white fur | l-hip | 100 | Modified Session 2 | runway |
| NOD, white fur | l-iliac-crest | 99.23664122 | Modified Session 2 | runway |
| NOD, white fur | l-shoulder | 98.4962406 | Modified Session 2 | runway |
| NOD, white fur | l-tail-base | 100 | Modified Session 2 | runway |
| NOD, white fur | l-wrist | 96.96969697 | Modified Session 2 | runway |

| | | | | |
|---|---|---|---|---|
| NOD, white fur | r-back-ankle | 96.2962963 | Modified Session 2 | runway |
| NOD, white fur | r-back-toe | 96.26865672 | Modified Session 2 | runway |
| NOD, white fur | r-elbow | 89.3129771 | Modified Session 2 | runway |
| NOD, white fur | r-front-toe-tip | 83.07692308 | Modified Session 2 | runway |
| NOD, white fur | r-head | 100 | Modified Session 2 | runway |
| NOD, white fur | r-hip | 100 | Modified Session 2 | runway |
| NOD, white fur | r-iliac-crest | 98.46153846 | Modified Session 2 | runway |
| NOD, white fur | r-shoulder | 99.23664122 | Modified Session 2 | runway |
| NOD, white fur | r-tail-base | 100 | Modified Session 2 | runway |
| NOD, white fur | r-wrist | 93.89312977 | Modified Session 2 | runway |

**Suppl. Table 3: Summary of accuracy during runway**

| Group | Likelihood | Training |
|---|---|---|
| Bl6, original mic | 99.28344848 | Original Session 1 |
| Bl6, new mice | 99.41159814 | Original Session 1 |
| NOD, white fur | 2.341346098 | Original Session 1 |
| NOD, white fur | 97.45471542 | Modified Session 2 |

**Suppl Table 4: Accuracy of labeling different mice during runway test**

| Group | Parameter | Likelihood | Training Session | Setup |
|---|---|---|---|---|
| Bl6, original mice | d-back-left | 96.969697 | Original Session 1 | ladder rung |
| Bl6, original mice | d-back-right | 98.2758621 | Original Session 1 | ladder rung |
| Bl6, original mice | d-front-left | 98.6998917 | Original Session 1 | ladder rung |
| Bl6, original mice | d-front-right | 95.9821429 | Original Session 1 | ladder rung |
| Bl6, original mice | d-head | 99.6760259 | Original Session 1 | ladder rung |
| Bl6, original mice | d-tail-base | 99.3576017 | Original Session 1 | ladder rung |
| Bl6, original mice | l-back-ankle | 94.5790081 | Original Session 1 | ladder rung |
| Bl6, original mice | l-back-toe | 91.1235955 | Original Session 1 | ladder rung |
| Bl6, original mice | l-elbow | 99.1351351 | Original Session 1 | ladder rung |
| Bl6, original mice | l-front-toe-tip | 81.6826411 | Original Session 1 | ladder rung |
| Bl6, original mice | l-head | 99.6499417 | Original Session 1 | ladder rung |
| Bl6, original mice | l-hip | 100 | Original Session 1 | ladder rung |
| Bl6, original mice | l-iliac-crest | 100 | Original Session 1 | ladder rung |
| Bl6, original mice | l-shoulder | 100 | Original Session 1 | ladder rung |
| Bl6, original mice | l-tail-base | 100 | Original Session 1 | ladder rung |
| Bl6, original mice | l-wrist | 89.5744681 | Original Session 1 | ladder rung |
| Bl6, original mice | r-back-ankle | 88.3886256 | Original Session 1 | ladder rung |
| Bl6, original mice | r-back-toe | 90.0816803 | Original Session 1 | ladder rung |
| Bl6, original mice | r-elbow | 95.3362256 | Original Session 1 | ladder rung |
| Bl6, original mice | r-front-toe-tip | 89.9022801 | Original Session 1 | ladder rung |
| Bl6, original mice | r-head | 100 | Original Session 1 | ladder rung |
| Bl6, original mice | r-hip | 100 | Original Session 1 | ladder rung |
| Bl6, original mice | r-iliac-crest | 100 | Original Session 1 | ladder rung |
| Bl6, original mice | r-shoulder | 97.4566474 | Original Session 1 | ladder rung |
| Bl6, original mice | r-tail-base | 100 | Original Session 1 | ladder rung |
| Bl6, original mice | r-wrist | 95.2329361 | Original Session 1 | ladder rung |
| Bl6, original mice | d-back-left | 100 | Original Session 1 | ladder rung |
| Bl6, new mice | d-back-right | 95.6018519 | Original Session 1 | ladder rung |
| Bl6, new mice | d-front-left | 97.752809 | Original Session 1 | ladder rung |
| Bl6, new mice | d-front-right | 94.5454545 | Original Session 1 | ladder rung |
| Bl6, new mice | d-head | 99.7722096 | Original Session 1 | ladder rung |
| Bl6, new mice | d-tail-base | 98.3758701 | Original Session 1 | ladder rung |
| Bl6, new mice | l-back-ankle | 87.5 | Original Session 1 | ladder rung |
| Bl6, new mice | l-back-toe | 82.5287356 | Original Session 1 | ladder rung |
| Bl6, new mice | l-elbow | 95.1388889 | Original Session 1 | ladder rung |
| Bl6, new mice | l-front-toe-tip | 74.0909091 | Original Session 1 | ladder rung |
| Bl6, new mice | l-head | 98.6238532 | Original Session 1 | ladder rung |
| Bl6, new mice | l-hip | 100 | Original Session 1 | ladder rung |
| Bl6, new mice | l-iliac-crest | 100 | Original Session 1 | ladder rung |
| Bl6, new mice | l-shoulder | 98.5781991 | Original Session 1 | ladder rung |
| Bl6, new mice | l-tail-base | 100 | Original Session 1 | ladder rung |
| Bl6, new mice | l-wrist | 86.0411899 | Original Session 1 | ladder rung |
| Bl6, new mice | r-back-ankle | 87.0748299 | Original Session 1 | ladder rung |
| Bl6, new mice | r-back-toe | 85.778781 | Original Session 1 | ladder rung |
| Bl6, new mice | r-elbow | 82.4884793 | Original Session 1 | ladder rung |
| Bl6, new mice | r-front-toe-tip | 86.4367816 | Original Session 1 | ladder rung |

| | | | | |
|---|---|---|---|---|
| Bl6, new mice | r-head | 99.3055556 | Original Session 1 | ladder rung |
| Bl6, new mice | r-hip | 99.3243243 | Original Session 1 | ladder rung |
| Bl6, new mice | r-iliac-crest | 99.7727273 | Original Session 1 | ladder rung |
| Bl6, new mice | r-shoulder | 76.443418 | Original Session 1 | ladder rung |
| Bl6, new mice | r-tail-base | 100 | Original Session 1 | ladder rung |
| Bl6, new mice | r-wrist | 92.9864253 | Original Session 1 | ladder rung |
| NOD, white fur | d-back-left | 0 | Original Session 1 | ladder rung |
| NOD, white fur | d-back-right | 0 | Original Session 1 | ladder rung |
| NOD, white fur | d-front-left | 0 | Original Session 1 | ladder rung |
| NOD, white fur | d-front-right | 0 | Original Session 1 | ladder rung |
| NOD, white fur | d-head | 0 | Original Session 1 | ladder rung |
| NOD, white fur | d-tail-base | 0 | Original Session 1 | ladder rung |
| NOD, white fur | l-back-ankle | 0 | Original Session 1 | ladder rung |
| NOD, white fur | l-back-toe | 0 | Original Session 1 | ladder rung |
| NOD, white fur | l-elbow | 0 | Original Session 1 | ladder rung |
| NOD, white fur | l-front-toe-tip | 0 | Original Session 1 | ladder rung |
| NOD, white fur | l-head | 0 | Original Session 1 | ladder rung |
| NOD, white fur | l-hip | 0 | Original Session 1 | ladder rung |
| NOD, white fur | l-iliac-crest | 0 | Original Session 1 | ladder rung |
| NOD, white fur | l-shoulder | 0 | Original Session 1 | ladder rung |
| NOD, white fur | l-tail-base | 0 | Original Session 1 | ladder rung |
| NOD, white fur | l-wrist | 0 | Original Session 1 | ladder rung |
| NOD, white fur | r-back-ankle | 0 | Original Session 1 | ladder rung |
| NOD, white fur | r-back-toe | 0 | Original Session 1 | ladder rung |
| NOD, white fur | r-elbow | 0 | Original Session 1 | ladder rung |
| NOD, white fur | r-front-toe-tip | 0 | Original Session 1 | ladder rung |
| NOD, white fur | r-head | 0 | Original Session 1 | ladder rung |
| NOD, white fur | r-hip | 0 | Original Session 1 | ladder rung |
| NOD, white fur | r-iliac-crest | 0 | Original Session 1 | ladder rung |
| NOD, white fur | r-shoulder | 0 | Original Session 1 | ladder rung |
| NOD, white fur | r-tail-base | 0 | Original Session 1 | ladder rung |
| NOD, white fur | r-wrist | 0 | Original Session 1 | ladder rung |
| NOD, white fur | d-back-left | 95.959596 | Modified Session 2 | ladder rung |
| NOD, white fur | d-back-right | 96.7576792 | Modified Session 2 | ladder rung |
| NOD, white fur | d-front-left | 89.2307692 | Modified Session 2 | ladder rung |
| NOD, white fur | d-front-right | 88.0749574 | Modified Session 2 | ladder rung |
| NOD, white fur | d-head | 99.6705107 | Modified Session 2 | ladder rung |
| NOD, white fur | d-tail-base | 82 | Modified Session 2 | ladder rung |
| NOD, white fur | l-back-ankle | 94.5392491 | Modified Session 2 | ladder rung |
| NOD, white fur | l-back-toe | 90.6354515 | Modified Session 2 | ladder rung |
| NOD, white fur | l-elbow | 98.6394558 | Modified Session 2 | ladder rung |
| NOD, white fur | l-front-toe-tip | 80.952381 | Modified Session 2 | ladder rung |
| NOD, white fur | l-head | 99.6557659 | Modified Session 2 | ladder rung |
| NOD, white fur | l-hip | 100 | Modified Session 2 | ladder rung |
| NOD, white fur | l-iliac-crest | 99.323181 | Modified Session 2 | ladder rung |
| NOD, white fur | l-shoulder | 99.6539792 | Modified Session 2 | ladder rung |
| NOD, white fur | l-tail-base | 99.6557659 | Modified Session 2 | ladder rung |
| NOD, white fur | l-wrist | 86.0306644 | Modified Session 2 | ladder rung |

| | | | | |
|---|---|---|---|---|
| NOD, white fur | r-back-ankle | 94.8892675 | Modified Session 2 | ladder rung |
| NOD, white fur | r-back-toe | 93.2773109 | Modified Session 2 | ladder rung |
| NOD, white fur | r-elbow | 93.814433 | Modified Session 2 | ladder rung |
| NOD, white fur | r-front-toe-tip | 75 | Modified Session 2 | ladder rung |
| NOD, white fur | r-head | 99.6533795 | Modified Session 2 | ladder rung |
| NOD, white fur | r-hip | 99.6655518 | Modified Session 2 | ladder rung |
| NOD, white fur | r-iliac-crest | 98.3305509 | Modified Session 2 | ladder rung |
| NOD, white fur | r-shoulder | 94.6826758 | Modified Session 2 | ladder rung |
| NOD, white fur | r-tail-base | 98.9813243 | Modified Session 2 | ladder rung |
| NOD, white fur | r-wrist | 80.4123711 | Modified Session 2 | ladder rung |

**Suppl. Table 5: Summary of accuracy during rung test**

| Group | Likelihood | Training |
|---|---|---|
| Bl6, original mice | 96.3372002 | Original Session 1 |
| Bl6, new mice | 93.0062036 | Original Session 1 |
| NOD, white fur | 0 | Original Session 1 |
| NOD, white fur | 93.4417797 | Modified Session 2 |

**Suppl Table 6: Relative laser Doppler intensities 24h after stroke injury, baseline is 1**

| Mouse_ID | Side | Relative LDI |
|---|---|---|
| ID_1 | L | 0.96964282 |
| ID_2 | L | 0.85884438 |
| ID_3 | L | 0.94832747 |
| ID_4 | L | 0.93208403 |
| ID_5 | L | 1.07793734 |
| ID_6 | L | 1.06103588 |
| ID_7 | L | 0.91017473 |
| ID_1 | R | 0.38120362 |
| ID_2 | R | 0.17035167 |
| ID_3 | R | 0.31565163 |
| ID_4 | R | 0.25427492 |
| ID_5 | R | 0.25176975 |
| ID_6 | R | 0.54153461 |
| ID_7 | R | 0.22838815 |
| ID_1 | baseline | 1.04581865 |
| ID_2 | baseline | 0.87407882 |
| ID_3 | baseline | 0.87236997 |
| ID_4 | baseline | 1.00907829 |
| ID_5 | baseline | 0.95524939 |
| ID_6 | baseline | 0.97361957 |
| ID_7 | baseline | 1.03513831 |
| ID_8 | baseline | 1.23464701 |

**Suppl. Table 7: Summary of LDI**

| Side | LDI | SD |
|---|---|---|
| L | 0.96543524 | 0.0791983 |
| R | 0.30616776 | 0.12333259 |
| baseline | 1 | 0.11546168 |

**Suppl Table 8: Statistical evaluation of data**

| Group1 | Group2 | padj | Sig | Stat test |
|---|---|---|---|---|
| Left side | baseline | 0.872246 | ns | Tukey multiple comparison |
| right side | baseline | 5.2007E-10 | **** | Tukey multiple comparison |
| right side | left side | 4.0138E-09 | **** | Tukey multiple comparison |

**Suppl Table 9:Stroke volume on left and right hemisphere at 21 dpi**

| Mouse_ID | Side | Storke Volume in mm3 |
|---|---|---|
| ID_1 | L | 0 |
| ID_2 | L | 0 |
| ID_3 | L | 0 |
| ID_4 | L | 0 |
| ID_5 | L | 0 |
| ID_6 | L | 0 |
| ID_7 | L | 0 |
| ID_1 | R | 1.299355407 |
| ID_2 | R | 1.089375951 |
| ID_3 | R | 1.329701471 |
| ID_4 | R | 1.056641243 |
| ID_5 | R | 1.382891652 |
| ID_6 | R | 1.621320592 |
| ID_7 | R | 1.34263095 |

**Suppl Table 10: Summary of stroke volume on left and right hemisphere at 21 dpi**

| Side | Storke Volume in mm3 | SD |
|---|---|---|
| L | 0 | 0 |
| R | 1.303131038 | 0.189883864 |

**Suppl Table 11: Stroke areas in cortical sections 21 dpi**

| Mouse_ID | Side | A-P axis relative to bregma in mm | Stroke area in mm2 |
|---|---|---|---|
| ID_1 | L | -2.5 | 0 |
| ID_2 | L | -2.5 | 0 |
| ID_3 | L | -2.5 | 0 |
| ID_4 | L | -2.5 | 0 |
| ID_5 | L | -2.5 | 0 |
| ID_6 | L | -2.5 | 0 |
| ID_7 | L | -2.5 | 0 |
| ID_1 | R | -2.5 | 0 |
| ID_2 | R | -2.5 | 0 |
| ID_3 | R | -2.5 | 0 |
| ID_4 | R | -2.5 | 0 |
| ID_5 | R | -2.5 | 0 |
| ID_6 | R | -2.5 | 0 |
| ID_7 | R | -2.5 | 0 |
| ID_1 | L | -1.5 | 0 |
| ID_2 | L | -1.5 | 0 |
| ID_3 | L | -1.5 | 0 |
| ID_4 | L | -1.5 | 0 |
| ID_5 | L | -1.5 | 0 |
| ID_6 | L | -1.5 | 0 |
| ID_7 | L | -1.5 | 0 |
| ID_1 | R | -1.5 | 0 |
| ID_2 | R | -1.5 | 0.62052749 |
| ID_3 | R | -1.5 | 0.395874101 |

| | | | |
|---|---|---|---|
| ID_4 | R | -1.5 | 0.269451822 |
| ID_5 | R | -1.5 | 0 |
| ID_6 | R | -1.5 | 0.807621874 |
| ID_7 | R | -1.5 | 0.546754634 |
| ID_1 | L | -0.5 | 0 |
| ID_2 | L | -0.5 | 0 |
| ID_3 | L | -0.5 | 0 |
| ID_4 | L | -0.5 | 0 |
| ID_5 | L | -0.5 | 0 |
| ID_6 | L | -0.5 | 0 |
| ID_7 | L | -0.5 | 0 |
| ID_1 | R | -0.5 | 0.237142277 |
| ID_2 | R | -0.5 | 0.722405361 |
| ID_3 | R | -0.5 | 0.4974167 |
| ID_4 | R | -0.5 | 0.590926211 |
| ID_5 | R | -0.5 | 0.548070937 |
| ID_6 | R | -0.5 | 0.757378657 |
| ID_7 | R | -0.5 | 0.736623045 |
| ID_1 | L | 0.5 | 0 |
| ID_2 | L | 0.5 | 0 |
| ID_3 | L | 0.5 | 0 |
| ID_4 | L | 0.5 | 0 |
| ID_5 | L | 0.5 | 0 |
| ID_6 | L | 0.5 | 0 |
| ID_7 | L | 0.5 | 0 |
| ID_1 | R | 0.5 | 0.913724914 |
| ID_2 | R | 0.5 | 0.473883078 |
| ID_3 | R | 0.5 | 0.768881997 |
| ID_4 | R | 0.5 | 0.72641693 |
| ID_5 | R | 0.5 | 0.838932186 |
| ID_6 | R | 0.5 | 0.578163799 |
| ID_7 | R | 0.5 | 0.434000051 |
| ID_1 | L | 1.5 | 0 |
| ID_2 | L | 1.5 | 0 |
| ID_3 | L | 1.5 | 0 |
| ID_4 | L | 1.5 | 0 |
| ID_5 | L | 1.5 | 0 |
| ID_6 | L | 1.5 | 0 |
| ID_7 | L | 1.5 | 0 |
| ID_1 | R | 1.5 | 0.434094674 |
| ID_2 | R | 1.5 | 0.257521549 |
| ID_3 | R | 1.5 | 0.286860314 |
| ID_4 | R | 1.5 | 0.047268964 |
| ID_5 | R | 1.5 | 0.607549084 |
| ID_6 | R | 1.5 | 0.288816558 |
| ID_7 | R | 1.5 | 0.296568696 |
| ID_1 | L | 2.5 | 0 |
| ID_2 | L | 2.5 | 0 |

| | | | |
|---|---|---|---|
| ID_3 | L | 2.5 | 0 |
| ID_4 | L | 2.5 | 0 |
| ID_5 | L | 2.5 | 0 |
| ID_6 | L | 2.5 | 0 |
| ID_7 | L | 2.5 | 0 |
| ID_1 | R | 2.5 | 0 |
| ID_2 | R | 2.5 | 0 |
| ID_3 | R | 2.5 | 0 |
| ID_4 | R | 2.5 | 0 |
| ID_5 | R | 2.5 | 0 |
| ID_6 | R | 2.5 | 0 |
| ID_7 | R | 2.5 | 0 |

**Suppl Table 12: Summary of stroke areas in cortical sections 21 dpi**

| Side | A-P axis relative to bregma | Stroke area in mm2 | SD |
|---|---|---|---|
| L | -2.5 | 0 | 0 |
| R | -2.5 | 0 | 0 |
| L | -1.5 | 0 | 0 |
| R | -1.5 | 0.377175703 | 0.308149882 |
| L | -0.5 | 0 | 0 |
| R | -0.5 | 0.584280455 | 0.18332815 |
| L | 0.5 | 0 | 0 |
| R | 0.5 | 0.676286136 | 0.18403602 |
| L | 1.5 | 0 | 0 |
| R | 1.5 | 0.316954263 | 0.171533564 |
| L | 2.5 | 0 | 0 |
| R | 2.5 | 0 | 0 |

**Suppl. Table 13: Raw data bottom analysis**

| day | ID | Parameter | Value |
|---|---|---|---|
| baseline | ID_1 | average duration | 0.25833333 |
| baseline | ID_2 | average duration | 0.54166667 |
| baseline | ID_3 | average duration | 0.36666667 |
| baseline | ID_4 | average duration | 0.425 |
| baseline | ID_5 | average duration | 0.59166667 |
| baseline | ID_6 | average duration | 0.61666667 |
| baseline | ID_7 | average duration | 0.25 |
| baseline | ID_8 | average duration | 0.40833333 |
| baseline | ID_9 | average duration | 0.33333333 |
| baseline | ID_10 | average duration | 0.23333333 |
| baseline | ID_11 | average duration | 0.75 |
| baseline | ID_12 | average duration | 0.38333333 |
| baseline | ID_13 | average duration | 0.23333333 |
| 3days | ID_1 | average duration | 0.61666667 |
| 3days | ID_2 | average duration | 0.33333333 |
| 3days | ID_3 | average duration | 0.3 |
| 3days | ID_4 | average duration | 0.6 |
| 3days | ID_5 | average duration | 0.71666667 |
| 3days | ID_6 | average duration | 0.65 |
| 3days | ID_7 | average duration | 0.99166667 |
| 3days | ID_8 | average duration | 0.66666667 |
| 7days | ID_1 | average duration | 0.38333333 |
| 7days | ID_2 | average duration | 0.53333333 |
| 7days | ID_3 | average duration | 0.36666667 |
| 7days | ID_4 | average duration | 0.23333333 |
| 7days | ID_5 | average duration | 0.25 |
| 7days | ID_6 | average duration | 0.40833333 |
| 7days | ID_7 | average duration | 0.65833333 |
| 7days | ID_8 | average duration | 0.45 |
| 7days | ID_9 | average duration | 0.95 |
| 14days | ID_1 | average duration | 0.35 |
| 14days | ID_2 | average duration | 0.3 |
| 14days | ID_3 | average duration | 0.38333333 |
| 14days | ID_4 | average duration | 0.48333333 |
| 14days | ID_5 | average duration | 0.31666667 |
| 14days | ID_6 | average duration | 0.56666667 |
| 14days | ID_7 | average duration | 0.4 |
| 14days | ID_8 | average duration | 0.50833333 |
| 14days | ID_9 | average duration | 0.56666667 |
| 14days | ID_10 | average duration | 0.8 |
| 14days | ID_11 | average duration | 0.33333333 |
| 21days | ID_1 | average duration | 0.48333333 |
| 21days | ID_2 | average duration | 0.425 |
| 21days | ID_3 | average duration | 0.40833333 |
| 21days | ID_4 | average duration | 0.16666667 |
| 21days | ID_5 | average duration | 0.36666667 |
| 21days | ID_6 | average duration | 0.31666667 |
| 21days | ID_7 | average duration | 0.24166667 |
| 21days | ID_8 | average duration | 0.525 |
| 21days | ID_9 | average duration | 0.31666667 |
| 21days | ID_10 | average duration | 0.275 |
| baseline | ID_1 | average stance time | 0.13333333 |
| baseline | ID_2 | average stance time | 0.15 |
| baseline | ID_3 | average stance time | 0.13333333 |

| | | | |
|---|---|---|---|
| baseline | ID_4 | average stance time | 0.13333333 |
| baseline | ID_5 | average stance time | 0.26666667 |
| baseline | ID_6 | average stance time | 0.13333333 |
| baseline | ID_7 | average stance time | 0.11666667 |
| baseline | ID_8 | average stance time | 0.1 |
| baseline | ID_9 | average stance time | 0.13333333 |
| baseline | ID_10 | average stance time | 0.1 |
| baseline | ID_11 | average stance time | 0.21666667 |
| baseline | ID_12 | average stance time | 0.11666667 |
| baseline | ID_13 | average stance time | 0.1 |
| 3days | ID_1 | average stance time | 0.28333333 |
| 3days | ID_2 | average stance time | 0.16666667 |
| 3days | ID_3 | average stance time | 0.15 |
| 3days | ID_4 | average stance time | 0.28333333 |
| 3days | ID_5 | average stance time | 0.26666667 |
| 3days | ID_6 | average stance time | 0.18333333 |
| 3days | ID_7 | average stance time | 0.2 |
| 3days | ID_8 | average stance time | 0.16666667 |
| 7days | ID_1 | average stance time | 0.15 |
| 7days | ID_2 | average stance time | 0.14166667 |
| 7days | ID_3 | average stance time | 0.15 |
| 7days | ID_4 | average stance time | 0.11666667 |
| 7days | ID_5 | average stance time | 0.1 |
| 7days | ID_6 | average stance time | 0.2 |
| 7days | ID_7 | average stance time | 0.28333333 |
| 7days | ID_8 | average stance time | 0.16666667 |
| 7days | ID_9 | average stance time | 0.19166667 |
| 14days | ID_1 | average stance time | 0.1 |
| 14days | ID_2 | average stance time | 0.13333333 |
| 14days | ID_3 | average stance time | 0.13333333 |
| 14days | ID_4 | average stance time | 0.16666667 |
| 14days | ID_5 | average stance time | 0.13333333 |
| 14days | ID_6 | average stance time | 0.3 |
| 14days | ID_7 | average stance time | 0.19166667 |
| 14days | ID_8 | average stance time | 0.18333333 |
| 14days | ID_9 | average stance time | 0.16666667 |
| 14days | ID_10 | average stance time | 0.16666667 |
| 14days | ID_11 | average stance time | 0.15 |
| 21days | ID_1 | average stance time | 0.18333333 |
| 21days | ID_2 | average stance time | 0.15 |
| 21days | ID_3 | average stance time | 0.2 |
| 21days | ID_4 | average stance time | 0.06666667 |
| 21days | ID_5 | average stance time | 0.18333333 |
| 21days | ID_6 | average stance time | 0.15833333 |
| 21days | ID_7 | average stance time | 0.11666667 |
| 21days | ID_8 | average stance time | 0.1 |
| 21days | ID_9 | average stance time | 0.15 |
| 21days | ID_10 | average stance time | 0.13333333 |
| baseline | ID_1 | average swing time | 0.125 |
| baseline | ID_2 | average swing time | 0.13333333 |
| baseline | ID_3 | average swing time | 0.13333333 |
| baseline | ID_4 | average swing time | 0.10833333 |
| baseline | ID_5 | average swing time | 0.15 |
| baseline | ID_6 | average swing time | 0.15 |
| baseline | ID_7 | average swing time | 0.13333333 |
| baseline | ID_8 | average swing time | 0.1 |

| | | | |
|---|---|---|---|
| baseline | ID_9 | average swing time | 0.13333333 |
| baseline | ID_10 | average swing time | 0.13333333 |
| baseline | ID_11 | average swing time | 0.13333333 |
| baseline | ID_12 | average swing time | 0.13333333 |
| baseline | ID_13 | average swing time | 0.13333333 |
| 3days | ID_1 | average swing time | 0.13333333 |
| 3days | ID_2 | average swing time | 0.13333333 |
| 3days | ID_3 | average swing time | 0.13333333 |
| 3days | ID_4 | average swing time | 0.11666667 |
| 3days | ID_5 | average swing time | 0.13333333 |
| 3days | ID_6 | average swing time | 0.13333333 |
| 3days | ID_7 | average swing time | 0.15 |
| 3days | ID_8 | average swing time | 0.11666667 |
| 7days | ID_1 | average swing time | 0.11666667 |
| 7days | ID_2 | average swing time | 0.11666667 |
| 7days | ID_3 | average swing time | 0.11666667 |
| 7days | ID_4 | average swing time | 0.13333333 |
| 7days | ID_5 | average swing time | 0.11666667 |
| 7days | ID_6 | average swing time | 0.15833333 |
| 7days | ID_7 | average swing time | 0.15 |
| 7days | ID_8 | average swing time | 0.16666667 |
| 7days | ID_9 | average swing time | 0.13333333 |
| 14days | ID_1 | average swing time | 0.11666667 |
| 14days | ID_2 | average swing time | 0.13333333 |
| 14days | ID_3 | average swing time | 0.11666667 |
| 14days | ID_4 | average swing time | 0.13333333 |
| 14days | ID_5 | average swing time | 0.13333333 |
| 14days | ID_6 | average swing time | 0.13333333 |
| 14days | ID_7 | average swing time | 0.16666667 |
| 14days | ID_8 | average swing time | 0.16666667 |
| 14days | ID_9 | average swing time | 0.11666667 |
| 14days | ID_10 | average swing time | 0.15 |
| 14days | ID_11 | average swing time | 0.15 |
| 21days | ID_1 | average swing time | 0.11666667 |
| 21days | ID_2 | average swing time | 0.16666667 |
| 21days | ID_3 | average swing time | 0.15 |
| 21days | ID_4 | average swing time | 0.13333333 |
| 21days | ID_5 | average swing time | 0.15 |
| 21days | ID_6 | average swing time | 0.14166667 |
| 21days | ID_7 | average swing time | 0.15 |
| 21days | ID_8 | average swing time | 0.14166667 |
| 21days | ID_9 | average swing time | 0.15 |
| 21days | ID_10 | average swing time | 0.13333333 |
| baseline | ID_1 | left front right back synchronization | 0.10416667 |
| baseline | ID_2 | left front right back synchronization | 0.17931034 |
| baseline | ID_3 | left front right back synchronization | 0.15789474 |
| baseline | ID_4 | left front right back synchronization | 0.28030303 |
| baseline | ID_5 | left front right back synchronization | 0.28497409 |
| baseline | ID_6 | left front right back synchronization | 0.10434783 |
| baseline | ID_7 | left front right back synchronization | 0.12844037 |
| baseline | ID_8 | left front right back synchronization | 0.22834646 |
| baseline | ID_9 | left front right back synchronization | 0.17213115 |
| baseline | ID_10 | left front right back synchronization | 0.0990099 |
| baseline | ID_11 | left front right back synchronization | 0.37820513 |
| baseline | ID_12 | left front right back synchronization | 0.13114754 |
| baseline | ID_13 | left front right back synchronization | 0.13684211 |

| | | | |
|---|---|---|---|
| 3days | ID_1 | left front right back synchronization | 0.33163265 |
| 3days | ID_2 | left front right back synchronization | 0.21126761 |
| 3days | ID_3 | left front right back synchronization | 0.18656716 |
| 3days | ID_4 | left front right back synchronization | 0.4 |
| 3days | ID_5 | left front right back synchronization | 0.36065574 |
| 3days | ID_6 | left front right back synchronization | 0.16564417 |
| 3days | ID_7 | left front right back synchronization | 0.30120482 |
| 3days | ID_8 | left front right back synchronization | 0.26 |
| 7days | ID_1 | left front right back synchronization | 0.1884058 |
| 7days | ID_2 | left front right back synchronization | 0.1986755 |
| 7days | ID_3 | left front right back synchronization | 0.28985507 |
| 7days | ID_4 | left front right back synchronization | 0.20535714 |
| 7days | ID_5 | left front right back synchronization | 0.14159292 |
| 7days | ID_6 | left front right back synchronization | 0.17037037 |
| 7days | ID_7 | left front right back synchronization | 0.34027778 |
| 7days | ID_8 | left front right back synchronization | 0.15503876 |
| 7days | ID_9 | left front right back synchronization | 0.27333333 |
| 14days | ID_1 | left front right back synchronization | 0.24576271 |
| 14days | ID_2 | left front right back synchronization | 0.15 |
| 14days | ID_3 | left front right back synchronization | 0.15909091 |
| 14days | ID_4 | left front right back synchronization | 0.19708029 |
| 14days | ID_5 | left front right back synchronization | 0.15315315 |
| 14days | ID_6 | left front right back synchronization | 0.27338129 |
| 14days | ID_7 | left front right back synchronization | 0.15 |
| 14days | ID_8 | left front right back synchronization | 0.16666667 |
| 14days | ID_9 | left front right back synchronization | 0.26206897 |
| 14days | ID_10 | left front right back synchronization | 0.18 |
| 14days | ID_11 | left front right back synchronization | 0.15 |
| 21days | ID_1 | left front right back synchronization | 0.26751592 |
| 21days | ID_2 | left front right back synchronization | 0.14655172 |
| 21days | ID_3 | left front right back synchronization | 0.21481481 |
| 21days | ID_4 | left front right back synchronization | 0.09589041 |
| 21days | ID_5 | left front right back synchronization | 0.17391304 |
| 21days | ID_6 | left front right back synchronization | 0.15151515 |
| 21days | ID_7 | left front right back synchronization | 0.09345794 |
| 21days | ID_8 | left front right back synchronization | 0.16666667 |
| 21days | ID_9 | left front right back synchronization | 0.1171875 |
| 21days | ID_10 | left front right back synchronization | 0.15044248 |
| baseline | ID_1 | right front left back synchronization | 0.0625 |
| baseline | ID_2 | right front left back synchronization | 0.14482759 |
| baseline | ID_3 | right front left back synchronization | 0.09022556 |
| baseline | ID_4 | right front left back synchronization | 0.28030303 |
| baseline | ID_5 | right front left back synchronization | 0.37823834 |
| baseline | ID_6 | right front left back synchronization | 0.16521739 |
| baseline | ID_7 | right front left back synchronization | 0.18348624 |
| baseline | ID_8 | right front left back synchronization | 0.2519685 |
| baseline | ID_9 | right front left back synchronization | 0.16393443 |
| baseline | ID_10 | right front left back synchronization | 0.07920792 |
| baseline | ID_11 | right front left back synchronization | 0.17307692 |
| baseline | ID_12 | right front left back synchronization | 0.14754098 |
| baseline | ID_13 | right front left back synchronization | 0.09473684 |
| 3days | ID_1 | right front left back synchronization | 0.48979592 |
| 3days | ID_2 | right front left back synchronization | 0.26760563 |
| 3days | ID_3 | right front left back synchronization | 0.17910448 |
| 3days | ID_4 | right front left back synchronization | 0.48717949 |
| 3days | ID_5 | right front left back synchronization | 0.40983607 |

| | | | |
|---|---|---|---|
| 3days | ID_6 | right front left back synchronization | 0.32515337 |
| 3days | ID_7 | right front left back synchronization | 0.21084337 |
| 3days | ID_8 | right front left back synchronization | 0.23333333 |
| 7days | ID_1 | right front left back synchronization | 0.31884058 |
| 7days | ID_2 | right front left back synchronization | 0.18543046 |
| 7days | ID_3 | right front left back synchronization | 0.28985507 |
| 7days | ID_4 | right front left back synchronization | 0.16071429 |
| 7days | ID_5 | right front left back synchronization | 0.20353982 |
| 7days | ID_6 | right front left back synchronization | 0.2 |
| 7days | ID_7 | right front left back synchronization | 0.43055556 |
| 7days | ID_8 | right front left back synchronization | 0.10852713 |
| 7days | ID_9 | right front left back synchronization | 0.30666667 |
| 14days | ID_1 | right front left back synchronization | 0.23728814 |
| 14days | ID_2 | right front left back synchronization | 0.1625 |
| 14days | ID_3 | right front left back synchronization | 0.16666667 |
| 14days | ID_4 | right front left back synchronization | 0.24087591 |
| 14days | ID_5 | right front left back synchronization | 0.17117117 |
| 14days | ID_6 | right front left back synchronization | 0.41726619 |
| 14days | ID_7 | right front left back synchronization | 0.2 |
| 14days | ID_8 | right front left back synchronization | 0.20138889 |
| 14days | ID_9 | right front left back synchronization | 0.27586207 |
| 14days | ID_10 | right front left back synchronization | 0.26666667 |
| 14days | ID_11 | right front left back synchronization | 0.15 |
| 21days | ID_1 | right front left back synchronization | 0.29936306 |
| 21days | ID_2 | right front left back synchronization | 0.11206897 |
| 21days | ID_3 | right front left back synchronization | 0.27407407 |
| 21days | ID_4 | right front left back synchronization | 0.10958904 |
| 21days | ID_5 | right front left back synchronization | 0.23188406 |
| 21days | ID_6 | right front left back synchronization | 0.13636364 |
| 21days | ID_7 | right front left back synchronization | 0.09345794 |
| 21days | ID_8 | right front left back synchronization | 0.06862745 |
| 21days | ID_9 | right front left back synchronization | 0.1953125 |
| 21days | ID_10 | right front left back synchronization | 0.22123894 |

**Suppl. Table 14: Summary of bottom analysis**

| day | Parameter | Mean | SD |
|---|---|---|---|
| baseline | average dur | 0.41474359 | 0.16534734 |
| 3days | average dur | 0.609375 | 0.21864652 |
| 7days | average dur | 0.47037037 | 0.22267749 |
| 14days | average dur | 0.45530303 | 0.14992001 |
| 21days | average dur | 0.3525 | 0.11097561 |
| baseline | average star | 0.141025641 | 0.04840769 |
| 3days | average star | 0.2125 | 0.05616727 |
| 7days | average star | 0.166666667 | 0.05416667 |
| 14days | average star | 0.165909091 | 0.05170087 |
| 21days | average star | 0.144166667 | 0.04121047 |
| baseline | average swi | 0.130769231 | 0.01377087 |
| 3days | average swi | 0.13125 | 0.01068117 |
| 7days | average swi | 0.134259259 | 0.01973982 |
| 14days | average swi | 0.137878788 | 0.01839521 |
| 21days | average swi | 0.143333333 | 0.0134944 |
| baseline | left front rig | 0.183470719 | 0.0856535 |
| 3days | left front rig | 0.277121519 | 0.08526072 |
| 7days | left front rig | 0.218100741 | 0.06769096 |
| 14days | left front rig | 0.189745818 | 0.04802373 |
| 21days | left front rig | 0.157795566 | 0.05319644 |
| baseline | right front l | 0.170404904 | 0.08919253 |
| 3days | right front l | 0.325356458 | 0.12344471 |
| 7days | right front l | 0.244903286 | 0.09919638 |
| 14days | right front l | 0.226335063 | 0.07660734 |
| 21days | right front l | 0.174197967 | 0.08079999 |

**Suppl. Table 15: Statistical test of bottom analysis**

| Measure | group1 | group2 | p.adj | sig | Test |
|---|---|---|---|---|---|
| average dura | baseline | 3days | 0.04648307 | * | Repeated ANOVA, PH test* |
| average dura | baseline | 7days | 1 | ns | Repeated ANOVA, PH test* |
| average dura | baseline | 14days | 1 | ns | Repeated ANOVA, PH test* |
| average dura | baseline | 21days | 1 | ns | Repeated ANOVA, PH test* |
| average stan | baseline | 3days | 0.03937093 | * | Repeated ANOVA, PH test* |
| average stan | baseline | 7days | 0.77236619 | ns | Repeated ANOVA, PH test* |
| average stan | baseline | 14days | 0.77236619 | ns | Repeated ANOVA, PH test* |
| average stan | baseline | 21days | 0.89167767 | ns | Repeated ANOVA, PH test* |
| duration_sd_ | baseline | 3days | 0.94665102 | ns | Repeated ANOVA, PH test* |
| duration_sd_ | baseline | 7days | 1 | ns | Repeated ANOVA, PH test* |
| duration_sd_ | baseline | 14days | 1 | ns | Repeated ANOVA, PH test* |
| duration_sd_ | baseline | 21days | 1 | ns | Repeated ANOVA, PH test* |
| LFRB synch | baseline | 3days | 0.02116338 | * | Repeated ANOVA, PH test* |
| LFRB synch | baseline | 7days | 0.47641714 | ns | Repeated ANOVA, PH test* |
| LFRB synch | baseline | 14days | 0.93621763 | ns | Repeated ANOVA, PH test* |
| LFRB synch | baseline | 21days | 0.93621763 | ns | Repeated ANOVA, PH test* |
| RFBL synch | baseline | 3days | 8.8073E-05 | **** | Repeated ANOVA, PH test* |
| RFBL synch | baseline | 7days | 0.0288736 | * | Repeated ANOVA, PH test* |
| RFBL synch | baseline | 14days | 0.03061719 | * | Repeated ANOVA, PH test* |
| RFBL synch | baseline | 21days | 0.80513565 | ns | Repeated ANOVA, PH test* |

* p.adjust.method = "holm", ref group = baseline (for more details see Methods)

**Suppl. Table 16: Raw data bottom angle analysis**

| day | Measure | Angle, ° | ID |
|---|---|---|---|
| 0days | LB__avg_Angle | 50.4077012 | ID_1 |
| 0days | LB__avg_Angle | 46.2632675 | ID_2 |
| 0days | LB__avg_Angle | 51.8477229 | ID_3 |
| 0days | LB__avg_Angle | 43.6508697 | ID_4 |
| 0days | LB__avg_Angle | 54.2814501 | ID_5 |
| 0days | LB__avg_Angle | 47.6089647 | ID_6 |
| 0days | LB__avg_Angle | 47.3941644 | ID_7 |
| 0days | LB__avg_Angle | 42.1677037 | ID_8 |
| 0days | LB__avg_Angle | 48.8915014 | ID_9 |
| 0days | LB__avg_Angle | 49.6482824 | ID_10 |
| 0days | LB__avg_Angle | 48.4205062 | ID_11 |
| 0days | LB__avg_Angle | 44.9161935 | ID_12 |
| 0days | LB__avg_Angle | 45.3047359 | ID_13 |
| 3days | LB__avg_Angle | 56.7093179 | ID_1 |
| 3days | LB__avg_Angle | 49.7875511 | ID_2 |
| 3days | LB__avg_Angle | 49.6046545 | ID_3 |
| 3days | LB__avg_Angle | 48.5468763 | ID_4 |
| 3days | LB__avg_Angle | 45.5417375 | ID_5 |
| 3days | LB__avg_Angle | 46.4329682 | ID_6 |
| 3days | LB__avg_Angle | 56.8399674 | ID_7 |
| 3days | LB__avg_Angle | 47.7674786 | ID_8 |
| 7days | LB__avg_Angle | 47.4811774 | ID_1 |
| 7days | LB__avg_Angle | 45.6543481 | ID_2 |
| 7days | LB__avg_Angle | 52.4828104 | ID_3 |
| 7days | LB__avg_Angle | 48.945709 | ID_4 |
| 7days | LB__avg_Angle | 50.8862455 | ID_5 |
| 7days | LB__avg_Angle | 51.4478357 | ID_6 |
| 7days | LB__avg_Angle | 48.9761677 | ID_7 |
| 7days | LB__avg_Angle | 50.5034691 | ID_8 |
| 7days | LB__avg_Angle | 55.529131 | ID_9 |
| 14days | LB__avg_Angle | 50.3122777 | ID_1 |
| 14days | LB__avg_Angle | 51.9261004 | ID_2 |
| 14days | LB__avg_Angle | 47.8045234 | ID_3 |
| 14days | LB__avg_Angle | 49.6406405 | ID_4 |
| 14days | LB__avg_Angle | 52.3945749 | ID_5 |
| 14days | LB__avg_Angle | 55.6305889 | ID_6 |
| 14days | LB__avg_Angle | 48.4529191 | ID_7 |
| 14days | LB__avg_Angle | 48.9886127 | ID_8 |
| 14days | LB__avg_Angle | 48.0132903 | ID_9 |
| 14days | LB__avg_Angle | 52.781057 | ID_10 |
| 14days | LB__avg_Angle | 55.0773354 | ID_11 |
| 21days | LB__avg_Angle | 47.672133 | ID_1 |
| 21days | LB__avg_Angle | 54.1935514 | ID_2 |
| 21days | LB__avg_Angle | 55.6359358 | ID_3 |
| 21days | LB__avg_Angle | 53.8303683 | ID_4 |
| 21days | LB__avg_Angle | 51.5771797 | ID_5 |
| 21days | LB__avg_Angle | 47.7049767 | ID_6 |
| 21days | LB__avg_Angle | 49.3323145 | ID_7 |
| 21days | LB__avg_Angle | 42.3500748 | ID_8 |
| 21days | LB__avg_Angle | 48.4011887 | ID_9 |
| 21days | LB__avg_Angle | 46.693582 | ID_10 |
| 0days | LB__max | 75.6027361 | ID_1 |
| 0days | LB__max | 81.4634481 | ID_2 |
| 0days | LB__max | 74.069999 | ID_3 |

| | | | |
|---|---|---|---|
| 0days | LB__max | 70.774883 | ID_4 |
| 0days | LB__max | 72.4356346 | ID_5 |
| 0days | LB__max | 69.9449045 | ID_6 |
| 0days | LB__max | 78.6149721 | ID_7 |
| 0days | LB__max | 78.9854971 | ID_8 |
| 0days | LB__max | 73.0850971 | ID_9 |
| 0days | LB__max | 75.4969288 | ID_10 |
| 0days | LB__max | 67.6386185 | ID_11 |
| 0days | LB__max | 76.7281341 | ID_12 |
| 0days | LB__max | 75.4352644 | ID_13 |
| 3days | LB__max | 71.3561627 | ID_1 |
| 3days | LB__max | 73.4782816 | ID_2 |
| 3days | LB__max | 73.5486086 | ID_3 |
| 3days | LB__max | 70.8213052 | ID_4 |
| 3days | LB__max | 79.0749323 | ID_5 |
| 3days | LB__max | 76.2572046 | ID_6 |
| 3days | LB__max | 82.0311242 | ID_7 |
| 3days | LB__max | 77.1708438 | ID_8 |
| 7days | LB__max | 72.7784758 | ID_1 |
| 7days | LB__max | 76.8154719 | ID_2 |
| 7days | LB__max | 71.4535317 | ID_3 |
| 7days | LB__max | 73.435902 | ID_4 |
| 7days | LB__max | 79.4444907 | ID_5 |
| 7days | LB__max | 72.6221119 | ID_6 |
| 7days | LB__max | 69.5756072 | ID_7 |
| 7days | LB__max | 71.4153506 | ID_8 |
| 7days | LB__max | 77.3774398 | ID_9 |
| 14days | LB__max | 74.9646969 | ID_1 |
| 14days | LB__max | 71.473176 | ID_2 |
| 14days | LB__max | 74.1434788 | ID_3 |
| 14days | LB__max | 70.0281556 | ID_4 |
| 14days | LB__max | 75.6343805 | ID_5 |
| 14days | LB__max | 69.3589935 | ID_6 |
| 14days | LB__max | 79.7671195 | ID_7 |
| 14days | LB__max | 80.5561463 | ID_8 |
| 14days | LB__max | 73.4745868 | ID_9 |
| 14days | LB__max | 74.9809401 | ID_10 |
| 14days | LB__max | 75.3733273 | ID_11 |
| 21days | LB__max | 77.1749442 | ID_1 |
| 21days | LB__max | 68.5800134 | ID_2 |
| 21days | LB__max | 72.8488419 | ID_3 |
| 21days | LB__max | 77.1250917 | ID_4 |
| 21days | LB__max | 77.724208 | ID_5 |
| 21days | LB__max | 78.6170218 | ID_6 |
| 21days | LB__max | 76.8056567 | ID_7 |
| 21days | LB__max | 75.4825825 | ID_8 |
| 21days | LB__max | 78.1885294 | ID_9 |
| 21days | LB__max | 73.484921 | ID_10 |
| 0days | LB__min | 18.779951 | ID_1 |
| 0days | LB__min | 12.1393046 | ID_2 |
| 0days | LB__min | 14.5346713 | ID_3 |
| 0days | LB__min | 9.86731573 | ID_4 |
| 0days | LB__min | 21.2852704 | ID_5 |
| 0days | LB__min | 17.7882452 | ID_6 |
| 0days | LB__min | 13.2839772 | ID_7 |
| 0days | LB__min | 11.1572571 | ID_8 |

| | | | |
|---|---|---|---|
| 0days | LB__min | 20.7572374 | ID_9 |
| 0days | LB__min | 11.512426 | ID_10 |
| 0days | LB__min | 16.4456199 | ID_11 |
| 0days | LB__min | 13.8128615 | ID_12 |
| 0days | LB__min | 18.9384396 | ID_13 |
| 3days | LB__min | 26.6972495 | ID_1 |
| 3days | LB__min | 21.1645706 | ID_2 |
| 3days | LB__min | 21.601163 | ID_3 |
| 3days | LB__min | 13.0184654 | ID_4 |
| 3days | LB__min | 11.2072338 | ID_5 |
| 3days | LB__min | 13.7395223 | ID_6 |
| 3days | LB__min | 19.7260928 | ID_7 |
| 3days | LB__min | 17.8103329 | ID_8 |
| 7days | LB__min | 15.6142006 | ID_1 |
| 7days | LB__min | 9.04592594 | ID_2 |
| 7days | LB__min | 13.3531695 | ID_3 |
| 7days | LB__min | 11.5769437 | ID_4 |
| 7days | LB__min | 16.9383934 | ID_5 |
| 7days | LB__min | 20.0917916 | ID_6 |
| 7days | LB__min | 13.6707086 | ID_7 |
| 7days | LB__min | 9.80482036 | ID_8 |
| 7days | LB__min | 24.0422522 | ID_9 |
| 14days | LB__min | 18.2062696 | ID_1 |
| 14days | LB__min | 20.739789 | ID_2 |
| 14days | LB__min | 12.5164845 | ID_3 |
| 14days | LB__min | 13.4960615 | ID_4 |
| 14days | LB__min | 16.2081826 | ID_5 |
| 14days | LB__min | 18.3561095 | ID_6 |
| 14days | LB__min | 13.0310458 | ID_7 |
| 14days | LB__min | 11.6624493 | ID_8 |
| 14days | LB__min | 18.5569438 | ID_9 |
| 14days | LB__min | 15.700712 | ID_10 |
| 14days | LB__min | 19.3793992 | ID_11 |
| 21days | LB__min | 12.8352746 | ID_1 |
| 21days | LB__min | 19.2433292 | ID_2 |
| 21days | LB__min | 17.5211671 | ID_3 |
| 21days | LB__min | 24.6634923 | ID_4 |
| 21days | LB__min | 13.3747976 | ID_5 |
| 21days | LB__min | 9.86454066 | ID_6 |
| 21days | LB__min | 18.4372497 | ID_7 |
| 21days | LB__min | 11.4983433 | ID_8 |
| 21days | LB__min | 12.5371903 | ID_9 |
| 21days | LB__min | 15.2239172 | ID_10 |
| 0days | LF__avg_Angle | 44.3766867 | ID_1 |
| 0days | LF__avg_Angle | 45.6786656 | ID_2 |
| 0days | LF__avg_Angle | 45.6025225 | ID_3 |
| 0days | LF__avg_Angle | 40.0442004 | ID_4 |
| 0days | LF__avg_Angle | 41.658499 | ID_5 |
| 0days | LF__avg_Angle | 45.8373681 | ID_6 |
| 0days | LF__avg_Angle | 40.9825227 | ID_7 |
| 0days | LF__avg_Angle | 46.3197746 | ID_8 |
| 0days | LF__avg_Angle | 39.7405324 | ID_9 |
| 0days | LF__avg_Angle | 41.446467 | ID_10 |
| 0days | LF__avg_Angle | 47.9112928 | ID_11 |
| 0days | LF__avg_Angle | 42.0872957 | ID_12 |
| 0days | LF__avg_Angle | 40.4297428 | ID_13 |

| | | | |
|---|---|---|---|
| 3days | LF__avg_Angle | 40.1017464 | ID_1 |
| 3days | LF__avg_Angle | 38.7049219 | ID_2 |
| 3days | LF__avg_Angle | 38.4627874 | ID_3 |
| 3days | LF__avg_Angle | 45.6049819 | ID_4 |
| 3days | LF__avg_Angle | 44.6651621 | ID_5 |
| 3days | LF__avg_Angle | 41.476749 | ID_6 |
| 3days | LF__avg_Angle | 41.566284 | ID_7 |
| 3days | LF__avg_Angle | 41.9276646 | ID_8 |
| 7days | LF__avg_Angle | 43.3831178 | ID_1 |
| 7days | LF__avg_Angle | 43.9012933 | ID_2 |
| 7days | LF__avg_Angle | 45.2450563 | ID_3 |
| 7days | LF__avg_Angle | 44.252576 | ID_4 |
| 7days | LF__avg_Angle | 47.1048525 | ID_5 |
| 7days | LF__avg_Angle | 38.8874691 | ID_6 |
| 7days | LF__avg_Angle | 43.4536674 | ID_7 |
| 7days | LF__avg_Angle | 41.652114 | ID_8 |
| 7days | LF__avg_Angle | 44.5867547 | ID_9 |
| 14days | LF__avg_Angle | 38.9225394 | ID_1 |
| 14days | LF__avg_Angle | 45.8809814 | ID_2 |
| 14days | LF__avg_Angle | 45.7993107 | ID_3 |
| 14days | LF__avg_Angle | 38.4470849 | ID_4 |
| 14days | LF__avg_Angle | 39.600663 | ID_5 |
| 14days | LF__avg_Angle | 38.4059727 | ID_6 |
| 14days | LF__avg_Angle | 37.2801076 | ID_7 |
| 14days | LF__avg_Angle | 37.5934035 | ID_8 |
| 14days | LF__avg_Angle | 39.6993253 | ID_9 |
| 14days | LF__avg_Angle | 43.2303069 | ID_10 |
| 14days | LF__avg_Angle | 39.7132891 | ID_11 |
| 21days | LF__avg_Angle | 41.9922658 | ID_1 |
| 21days | LF__avg_Angle | 38.1049868 | ID_2 |
| 21days | LF__avg_Angle | 38.8365056 | ID_3 |
| 21days | LF__avg_Angle | 44.9341325 | ID_4 |
| 21days | LF__avg_Angle | 45.5550475 | ID_5 |
| 21days | LF__avg_Angle | 41.8214176 | ID_6 |
| 21days | LF__avg_Angle | 42.9731244 | ID_7 |
| 21days | LF__avg_Angle | 43.8879138 | ID_8 |
| 21days | LF__avg_Angle | 40.3193466 | ID_9 |
| 21days | LF__avg_Angle | 45.9123759 | ID_10 |
| 0days | LF__max | 71.3555474 | ID_1 |
| 0days | LF__max | 73.6205435 | ID_2 |
| 0days | LF__max | 72.7194728 | ID_3 |
| 0days | LF__max | 73.9135913 | ID_4 |
| 0days | LF__max | 72.1734714 | ID_5 |
| 0days | LF__max | 76.8629446 | ID_6 |
| 0days | LF__max | 65.9668713 | ID_7 |
| 0days | LF__max | 71.6323368 | ID_8 |
| 0days | LF__max | 68.1159155 | ID_9 |
| 0days | LF__max | 63.0155705 | ID_10 |
| 0days | LF__max | 71.6518054 | ID_11 |
| 0days | LF__max | 66.2048668 | ID_12 |
| 0days | LF__max | 67.1179287 | ID_13 |
| 3days | LF__max | 62.8944716 | ID_1 |
| 3days | LF__max | 62.0243492 | ID_2 |
| 3days | LF__max | 64.4625047 | ID_3 |
| 3days | LF__max | 65.1996682 | ID_4 |
| 3days | LF__max | 63.0171567 | ID_5 |

| | | | |
|---|---|---|---|
| 3days | LF__max | 67.9348204 | ID_6 |
| 3days | LF__max | 69.0442159 | ID_7 |
| 3days | LF__max | 69.2306117 | ID_8 |
| 7days | LF__max | 74.7220877 | ID_1 |
| 7days | LF__max | 60.1052282 | ID_2 |
| 7days | LF__max | 73.5050624 | ID_3 |
| 7days | LF__max | 66.6297859 | ID_4 |
| 7days | LF__max | 70.7693029 | ID_5 |
| 7days | LF__max | 62.5451288 | ID_6 |
| 7days | LF__max | 72.0979674 | ID_7 |
| 7days | LF__max | 67.5696475 | ID_8 |
| 7days | LF__max | 61.6892277 | ID_9 |
| 14days | LF__max | 70.8544514 | ID_1 |
| 14days | LF__max | 69.2382287 | ID_2 |
| 14days | LF__max | 71.0093515 | ID_3 |
| 14days | LF__max | 59.1540156 | ID_4 |
| 14days | LF__max | 58.9617768 | ID_5 |
| 14days | LF__max | 65.8127948 | ID_6 |
| 14days | LF__max | 60.3439419 | ID_7 |
| 14days | LF__max | 58.0746393 | ID_8 |
| 14days | LF__max | 63.8947623 | ID_9 |
| 14days | LF__max | 62.8024434 | ID_10 |
| 14days | LF__max | 70.2428672 | ID_11 |
| 21days | LF__max | 67.897832 | ID_1 |
| 21days | LF__max | 68.5109692 | ID_2 |
| 21days | LF__max | 72.0326392 | ID_3 |
| 21days | LF__max | 70.721441 | ID_4 |
| 21days | LF__max | 67.4516664 | ID_5 |
| 21days | LF__max | 62.7378209 | ID_6 |
| 21days | LF__max | 67.4799931 | ID_7 |
| 21days | LF__max | 63.9467885 | ID_8 |
| 21days | LF__max | 59.830988 | ID_9 |
| 21days | LF__max | 73.2086895 | ID_10 |
| 0days | LF__min | 23.8666621 | ID_1 |
| 0days | LF__min | 22.012441 | ID_2 |
| 0days | LF__min | 23.2117812 | ID_3 |
| 0days | LF__min | 17.4185718 | ID_4 |
| 0days | LF__min | 13.0684365 | ID_5 |
| 0days | LF__min | 13.8849846 | ID_6 |
| 0days | LF__min | 17.3052319 | ID_7 |
| 0days | LF__min | 21.8392712 | ID_8 |
| 0days | LF__min | 17.7712817 | ID_9 |
| 0days | LF__min | 13.9475869 | ID_10 |
| 0days | LF__min | 20.8303449 | ID_11 |
| 0days | LF__min | 23.4446216 | ID_12 |
| 0days | LF__min | 22.143269 | ID_13 |
| 3days | LF__min | 20.0237983 | ID_1 |
| 3days | LF__min | 16.8376944 | ID_2 |
| 3days | LF__min | 19.4127383 | ID_3 |
| 3days | LF__min | 24.6163476 | ID_4 |
| 3days | LF__min | 27.4194864 | ID_5 |
| 3days | LF__min | 24.5957825 | ID_6 |
| 3days | LF__min | 19.5470476 | ID_7 |
| 3days | LF__min | 15.5388239 | ID_8 |
| 7days | LF__min | 20.5571831 | ID_1 |
| 7days | LF__min | 25.0332006 | ID_2 |

| | | | |
|---|---|---|---|
| 7days | LF__min | 22.8107086 | ID_3 |
| 7days | LF__min | 28.962962 | ID_4 |
| 7days | LF__min | 26.9066641 | ID_5 |
| 7days | LF__min | 15.8483888 | ID_6 |
| 7days | LF__min | 17.052464 | ID_7 |
| 7days | LF__min | 18.0987147 | ID_8 |
| 7days | LF__min | 13.8332881 | ID_9 |
| 14days | LF__min | 18.1671132 | ID_1 |
| 14days | LF__min | 21.0296377 | ID_2 |
| 14days | LF__min | 26.4345983 | ID_3 |
| 14days | LF__min | 23.4241709 | ID_4 |
| 14days | LF__min | 19.4164151 | ID_5 |
| 14days | LF__min | 17.8566158 | ID_6 |
| 14days | LF__min | 20.1419256 | ID_7 |
| 14days | LF__min | 22.1649327 | ID_8 |
| 14days | LF__min | 15.9152806 | ID_9 |
| 14days | LF__min | 19.8013228 | ID_10 |
| 14days | LF__min | 16.1697534 | ID_11 |
| 21days | LF__min | 22.1063674 | ID_1 |
| 21days | LF__min | 14.0338723 | ID_2 |
| 21days | LF__min | 15.5453273 | ID_3 |
| 21days | LF__min | 21.0632339 | ID_4 |
| 21days | LF__min | 18.8898574 | ID_5 |
| 21days | LF__min | 22.6806905 | ID_6 |
| 21days | LF__min | 20.0356047 | ID_7 |
| 21days | LF__min | 16.0254051 | ID_8 |
| 21days | LF__min | 22.2348411 | ID_9 |
| 21days | LF__min | 17.4894865 | ID_10 |
| 0days | RB__avg_Angle | 42.8575696 | ID_1 |
| 0days | RB__avg_Angle | 47.8907736 | ID_2 |
| 0days | RB__avg_Angle | 45.4616962 | ID_3 |
| 0days | RB__avg_Angle | 46.4126777 | ID_4 |
| 0days | RB__avg_Angle | 51.1492484 | ID_5 |
| 0days | RB__avg_Angle | 48.938546 | ID_6 |
| 0days | RB__avg_Angle | 50.3347242 | ID_7 |
| 0days | RB__avg_Angle | 42.5578142 | ID_8 |
| 0days | RB__avg_Angle | 45.3973929 | ID_9 |
| 0days | RB__avg_Angle | 48.2625869 | ID_10 |
| 0days | RB__avg_Angle | 49.9375309 | ID_11 |
| 0days | RB__avg_Angle | 47.5111561 | ID_12 |
| 0days | RB__avg_Angle | 46.988602 | ID_13 |
| 3days | RB__avg_Angle | 56.6963994 | ID_1 |
| 3days | RB__avg_Angle | 50.0101968 | ID_2 |
| 3days | RB__avg_Angle | 48.2409531 | ID_3 |
| 3days | RB__avg_Angle | 56.5917773 | ID_4 |
| 3days | RB__avg_Angle | 53.4249406 | ID_5 |
| 3days | RB__avg_Angle | 50.6791906 | ID_6 |
| 3days | RB__avg_Angle | 53.1195539 | ID_7 |
| 3days | RB__avg_Angle | 52.5745364 | ID_8 |
| 7days | RB__avg_Angle | 50.1109123 | ID_1 |
| 7days | RB__avg_Angle | 43.3631402 | ID_2 |
| 7days | RB__avg_Angle | 47.2184043 | ID_3 |
| 7days | RB__avg_Angle | 47.295858 | ID_4 |
| 7days | RB__avg_Angle | 47.4237984 | ID_5 |
| 7days | RB__avg_Angle | 49.2825584 | ID_6 |
| 7days | RB__avg_Angle | 45.6127664 | ID_7 |

| | | | |
|---|---|---|---|
| 7days | RB__avg_Angle | 45.5638926 | ID_8 |
| 7days | RB__avg_Angle | 57.9434523 | ID_9 |
| 14days | RB__avg_Angle | 52.7787434 | ID_1 |
| 14days | RB__avg_Angle | 44.1025601 | ID_2 |
| 14days | RB__avg_Angle | 44.1475928 | ID_3 |
| 14days | RB__avg_Angle | 46.4501452 | ID_4 |
| 14days | RB__avg_Angle | 51.0710783 | ID_5 |
| 14days | RB__avg_Angle | 54.6343834 | ID_6 |
| 14days | RB__avg_Angle | 52.0154762 | ID_7 |
| 14days | RB__avg_Angle | 52.7532014 | ID_8 |
| 14days | RB__avg_Angle | 54.9479005 | ID_9 |
| 14days | RB__avg_Angle | 47.1932394 | ID_10 |
| 14days | RB__avg_Angle | 43.3683346 | ID_11 |
| 21days | RB__avg_Angle | 46.5341358 | ID_1 |
| 21days | RB__avg_Angle | 49.6472984 | ID_2 |
| 21days | RB__avg_Angle | 53.0770775 | ID_3 |
| 21days | RB__avg_Angle | 47.5182862 | ID_4 |
| 21days | RB__avg_Angle | 51.0303299 | ID_5 |
| 21days | RB__avg_Angle | 51.8459975 | ID_6 |
| 21days | RB__avg_Angle | 47.550468 | ID_7 |
| 21days | RB__avg_Angle | 43.8839373 | ID_8 |
| 21days | RB__avg_Angle | 53.6451286 | ID_9 |
| 21days | RB__avg_Angle | 48.3182814 | ID_10 |
| 0days | RB__max | 79.2524699 | ID_1 |
| 0days | RB__max | 76.5545904 | ID_2 |
| 0days | RB__max | 76.2632394 | ID_3 |
| 0days | RB__max | 74.4156858 | ID_4 |
| 0days | RB__max | 69.7314665 | ID_5 |
| 0days | RB__max | 73.1861504 | ID_6 |
| 0days | RB__max | 78.4634387 | ID_7 |
| 0days | RB__max | 77.4835416 | ID_8 |
| 0days | RB__max | 74.4808528 | ID_9 |
| 0days | RB__max | 77.6515358 | ID_10 |
| 0days | RB__max | 70.9886417 | ID_11 |
| 0days | RB__max | 73.2560674 | ID_12 |
| 0days | RB__max | 74.6354537 | ID_13 |
| 3days | RB__max | 74.4457326 | ID_1 |
| 3days | RB__max | 73.6827252 | ID_2 |
| 3days | RB__max | 80.1357593 | ID_3 |
| 3days | RB__max | 73.6737137 | ID_4 |
| 3days | RB__max | 74.8518291 | ID_5 |
| 3days | RB__max | 76.7463005 | ID_6 |
| 3days | RB__max | 70.605494 | ID_7 |
| 3days | RB__max | 81.7661769 | ID_8 |
| 7days | RB__max | 78.607353 | ID_1 |
| 7days | RB__max | 73.582011 | ID_2 |
| 7days | RB__max | 76.0878981 | ID_3 |
| 7days | RB__max | 75.9765241 | ID_4 |
| 7days | RB__max | 77.4873907 | ID_5 |
| 7days | RB__max | 74.3065407 | ID_6 |
| 7days | RB__max | 67.1127163 | ID_7 |
| 7days | RB__max | 74.5862763 | ID_8 |
| 7days | RB__max | 72.2565461 | ID_9 |
| 14days | RB__max | 80.9040511 | ID_1 |
| 14days | RB__max | 71.0466587 | ID_2 |
| 14days | RB__max | 75.2850996 | ID_3 |

| | | | |
|---|---|---|---|
| 14days | RB__max | 79.4131704 | ID_4 |
| 14days | RB__max | 76.2390665 | ID_5 |
| 14days | RB__max | 72.1987127 | ID_6 |
| 14days | RB__max | 75.1817158 | ID_7 |
| 14days | RB__max | 75.8606938 | ID_8 |
| 14days | RB__max | 81.5700622 | ID_9 |
| 14days | RB__max | 71.4506763 | ID_10 |
| 14days | RB__max | 74.1709864 | ID_11 |
| 21days | RB__max | 77.0178975 | ID_1 |
| 21days | RB__max | 74.621803 | ID_2 |
| 21days | RB__max | 74.5669036 | ID_3 |
| 21days | RB__max | 69.2736063 | ID_4 |
| 21days | RB__max | 76.0697052 | ID_5 |
| 21days | RB__max | 76.5524895 | ID_6 |
| 21days | RB__max | 78.7967047 | ID_7 |
| 21days | RB__max | 69.5051639 | ID_8 |
| 21days | RB__max | 79.9013704 | ID_9 |
| 21days | RB__max | 70.4869638 | ID_10 |
| 0days | RB__min | 11.0013328 | ID_1 |
| 0days | RB__min | 16.3580449 | ID_2 |
| 0days | RB__min | 12.0878673 | ID_3 |
| 0days | RB__min | 14.0298275 | ID_4 |
| 0days | RB__min | 12.3896594 | ID_5 |
| 0days | RB__min | 15.1524297 | ID_6 |
| 0days | RB__min | 13.928863 | ID_7 |
| 0days | RB__min | 17.8776828 | ID_8 |
| 0days | RB__min | 8.49166957 | ID_9 |
| 0days | RB__min | 13.9296039 | ID_10 |
| 0days | RB__min | 16.3993358 | ID_11 |
| 0days | RB__min | 9.32792728 | ID_12 |
| 0days | RB__min | 15.0074391 | ID_13 |
| 3days | RB__min | 18.9303141 | ID_1 |
| 3days | RB__min | 8.110522 | ID_2 |
| 3days | RB__min | 14.7929504 | ID_3 |
| 3days | RB__min | 23.3820277 | ID_4 |
| 3days | RB__min | 21.5492284 | ID_5 |
| 3days | RB__min | 15.2845306 | ID_6 |
| 3days | RB__min | 11.8865389 | ID_7 |
| 3days | RB__min | 17.4785325 | ID_8 |
| 7days | RB__min | 10.9708971 | ID_1 |
| 7days | RB__min | 13.8859988 | ID_2 |
| 7days | RB__min | 14.3423761 | ID_3 |
| 7days | RB__min | 15.0188234 | ID_4 |
| 7days | RB__min | 13.8104608 | ID_5 |
| 7days | RB__min | 19.1767414 | ID_6 |
| 7days | RB__min | 10.7747456 | ID_7 |
| 7days | RB__min | 9.09689337 | ID_8 |
| 7days | RB__min | 20.9777243 | ID_9 |
| 14days | RB__min | 13.6894278 | ID_1 |
| 14days | RB__min | 19.3527546 | ID_2 |
| 14days | RB__min | 16.8424035 | ID_3 |
| 14days | RB__min | 13.6571717 | ID_4 |
| 14days | RB__min | 12.8456176 | ID_5 |
| 14days | RB__min | 18.6861882 | ID_6 |
| 14days | RB__min | 12.9638358 | ID_7 |
| 14days | RB__min | 13.1649191 | ID_8 |

| | | | |
|---|---|---|---|
| 14days | RB__min | 18.5242693 | ID_9 |
| 14days | RB__min | 16.5177571 | ID_10 |
| 14days | RB__min | 15.4198488 | ID_11 |
| 21days | RB__min | 13.8780983 | ID_1 |
| 21days | RB__min | 18.4699646 | ID_2 |
| 21days | RB__min | 18.3929991 | ID_3 |
| 21days | RB__min | 21.705148 | ID_4 |
| 21days | RB__min | 16.0258867 | ID_5 |
| 21days | RB__min | 19.026834 | ID_6 |
| 21days | RB__min | 15.4930936 | ID_7 |
| 21days | RB__min | 7.86374811 | ID_8 |
| 21days | RB__min | 12.7132641 | ID_9 |
| 21days | RB__min | 19.236803 | ID_10 |
| 0days | RF__avg_Angle | 42.5470983 | ID_1 |
| 0days | RF__avg_Angle | 42.0275756 | ID_2 |
| 0days | RF__avg_Angle | 44.9028249 | ID_3 |
| 0days | RF__avg_Angle | 47.1531512 | ID_4 |
| 0days | RF__avg_Angle | 44.5139313 | ID_5 |
| 0days | RF__avg_Angle | 38.5663288 | ID_6 |
| 0days | RF__avg_Angle | 41.2785736 | ID_7 |
| 0days | RF__avg_Angle | 41.1782994 | ID_8 |
| 0days | RF__avg_Angle | 40.1662729 | ID_9 |
| 0days | RF__avg_Angle | 39.1084113 | ID_10 |
| 0days | RF__avg_Angle | 44.0872109 | ID_11 |
| 0days | RF__avg_Angle | 38.1311052 | ID_12 |
| 0days | RF__avg_Angle | 38.6904266 | ID_13 |
| 3days | RF__avg_Angle | 41.245129 | ID_1 |
| 3days | RF__avg_Angle | 39.4679567 | ID_2 |
| 3days | RF__avg_Angle | 42.4764923 | ID_3 |
| 3days | RF__avg_Angle | 48.3882549 | ID_4 |
| 3days | RF__avg_Angle | 40.0920999 | ID_5 |
| 3days | RF__avg_Angle | 38.9129161 | ID_6 |
| 3days | RF__avg_Angle | 34.324454 | ID_7 |
| 3days | RF__avg_Angle | 37.616126 | ID_8 |
| 7days | RF__avg_Angle | 42.4987069 | ID_1 |
| 7days | RF__avg_Angle | 42.3441563 | ID_2 |
| 7days | RF__avg_Angle | 43.7749879 | ID_3 |
| 7days | RF__avg_Angle | 43.2740301 | ID_4 |
| 7days | RF__avg_Angle | 43.4596765 | ID_5 |
| 7days | RF__avg_Angle | 40.4807501 | ID_6 |
| 7days | RF__avg_Angle | 41.7278437 | ID_7 |
| 7days | RF__avg_Angle | 43.4188312 | ID_8 |
| 7days | RF__avg_Angle | 44.9213287 | ID_9 |
| 14days | RF__avg_Angle | 44.0113542 | ID_1 |
| 14days | RF__avg_Angle | 46.7872042 | ID_2 |
| 14days | RF__avg_Angle | 39.700803 | ID_3 |
| 14days | RF__avg_Angle | 43.9078137 | ID_4 |
| 14days | RF__avg_Angle | 46.5437653 | ID_5 |
| 14days | RF__avg_Angle | 42.4092405 | ID_6 |
| 14days | RF__avg_Angle | 40.5157696 | ID_7 |
| 14days | RF__avg_Angle | 42.6896769 | ID_8 |
| 14days | RF__avg_Angle | 36.7752737 | ID_9 |
| 14days | RF__avg_Angle | 43.9457101 | ID_10 |
| 14days | RF__avg_Angle | 37.4580785 | ID_11 |
| 21days | RF__avg_Angle | 42.4759573 | ID_1 |
| 21days | RF__avg_Angle | 44.5900231 | ID_2 |

| | | | |
|---|---|---|---|
| 21days | RF__avg_Angle | 47.919704 | ID_3 |
| 21days | RF__avg_Angle | 41.8289005 | ID_4 |
| 21days | RF__avg_Angle | 40.3849643 | ID_5 |
| 21days | RF__avg_Angle | 42.0178862 | ID_6 |
| 21days | RF__avg_Angle | 44.1430753 | ID_7 |
| 21days | RF__avg_Angle | 39.145023 | ID_8 |
| 21days | RF__avg_Angle | 41.2880727 | ID_9 |
| 21days | RF__avg_Angle | 45.7266488 | ID_10 |
| 0days | RF__max | 65.1650482 | ID_1 |
| 0days | RF__max | 63.5633778 | ID_2 |
| 0days | RF__max | 66.2751629 | ID_3 |
| 0days | RF__max | 63.3982971 | ID_4 |
| 0days | RF__max | 68.1431847 | ID_5 |
| 0days | RF__max | 63.1142133 | ID_6 |
| 0days | RF__max | 68.3333809 | ID_7 |
| 0days | RF__max | 65.4583984 | ID_8 |
| 0days | RF__max | 66.4922866 | ID_9 |
| 0days | RF__max | 63.9437223 | ID_10 |
| 0days | RF__max | 65.649095 | ID_11 |
| 0days | RF__max | 61.7136828 | ID_12 |
| 0days | RF__max | 64.8635144 | ID_13 |
| 3days | RF__max | 58.1757834 | ID_1 |
| 3days | RF__max | 57.18388 | ID_2 |
| 3days | RF__max | 61.873653 | ID_3 |
| 3days | RF__max | 60.3414347 | ID_4 |
| 3days | RF__max | 64.3680365 | ID_5 |
| 3days | RF__max | 62.9111936 | ID_6 |
| 3days | RF__max | 69.0835872 | ID_7 |
| 3days | RF__max | 66.1051816 | ID_8 |
| 7days | RF__max | 64.2548686 | ID_1 |
| 7days | RF__max | 66.1595638 | ID_2 |
| 7days | RF__max | 67.2837534 | ID_3 |
| 7days | RF__max | 66.3447234 | ID_4 |
| 7days | RF__max | 66.70868 | ID_5 |
| 7days | RF__max | 64.8272284 | ID_6 |
| 7days | RF__max | 65.2065834 | ID_7 |
| 7days | RF__max | 65.6265089 | ID_8 |
| 7days | RF__max | 81.2978772 | ID_9 |
| 14days | RF__max | 65.7958842 | ID_1 |
| 14days | RF__max | 65.0531998 | ID_2 |
| 14days | RF__max | 68.1754157 | ID_3 |
| 14days | RF__max | 71.5300217 | ID_4 |
| 14days | RF__max | 68.488436 | ID_5 |
| 14days | RF__max | 61.4620513 | ID_6 |
| 14days | RF__max | 64.3952247 | ID_7 |
| 14days | RF__max | 72.7651226 | ID_8 |
| 14days | RF__max | 65.5283881 | ID_9 |
| 14days | RF__max | 74.306757 | ID_10 |
| 14days | RF__max | 65.7601374 | ID_11 |
| 21days | RF__max | 63.3070223 | ID_1 |
| 21days | RF__max | 65.2948342 | ID_2 |
| 21days | RF__max | 72.7203099 | ID_3 |
| 21days | RF__max | 68.5884133 | ID_4 |
| 21days | RF__max | 61.235668 | ID_5 |
| 21days | RF__max | 63.9349587 | ID_6 |
| 21days | RF__max | 66.5295257 | ID_7 |

| | | | |
|---|---|---|---|
| 21days | RF__max | 68.8908511 | ID_8 |
| 21days | RF__max | 64.5858355 | ID_9 |
| 21days | RF__max | 65.0675158 | ID_10 |
| 0days | RF__min | 24.6319339 | ID_1 |
| 0days | RF__min | 14.7688315 | ID_2 |
| 0days | RF__min | 18.0309036 | ID_3 |
| 0days | RF__min | 21.744939 | ID_4 |
| 0days | RF__min | 19.2623516 | ID_5 |
| 0days | RF__min | 16.7684322 | ID_6 |
| 0days | RF__min | 21.5467565 | ID_7 |
| 0days | RF__min | 17.1549487 | ID_8 |
| 0days | RF__min | 18.826316 | ID_9 |
| 0days | RF__min | 23.5426925 | ID_10 |
| 0days | RF__min | 22.3413851 | ID_11 |
| 0days | RF__min | 22.3218924 | ID_12 |
| 0days | RF__min | 21.0932269 | ID_13 |
| 3days | RF__min | 23.8001133 | ID_1 |
| 3days | RF__min | 21.5756968 | ID_2 |
| 3days | RF__min | 22.6118278 | ID_3 |
| 3days | RF__min | 24.4239223 | ID_4 |
| 3days | RF__min | 17.9108196 | ID_5 |
| 3days | RF__min | 22.6727528 | ID_6 |
| 3days | RF__min | 13.8024176 | ID_7 |
| 3days | RF__min | 17.578919 | ID_8 |
| 7days | RF__min | 22.6544294 | ID_1 |
| 7days | RF__min | 18.8966195 | ID_2 |
| 7days | RF__min | 18.9814378 | ID_3 |
| 7days | RF__min | 17.7187437 | ID_4 |
| 7days | RF__min | 20.6005688 | ID_5 |
| 7days | RF__min | 13.0939727 | ID_6 |
| 7days | RF__min | 23.2448505 | ID_7 |
| 7days | RF__min | 19.0907271 | ID_8 |
| 7days | RF__min | 19.7519316 | ID_9 |
| 14days | RF__min | 27.0984077 | ID_1 |
| 14days | RF__min | 23.841235 | ID_2 |
| 14days | RF__min | 12.7051403 | ID_3 |
| 14days | RF__min | 14.6091791 | ID_4 |
| 14days | RF__min | 16.563378 | ID_5 |
| 14days | RF__min | 14.4490435 | ID_6 |
| 14days | RF__min | 17.6629117 | ID_7 |
| 14days | RF__min | 18.5965583 | ID_8 |
| 14days | RF__min | 15.3036826 | ID_9 |
| 14days | RF__min | 12.5284398 | ID_10 |
| 14days | RF__min | 20.3577251 | ID_11 |
| 21days | RF__min | 24.6862726 | ID_1 |
| 21days | RF__min | 19.9209133 | ID_2 |
| 21days | RF__min | 22.1483525 | ID_3 |
| 21days | RF__min | 16.1469054 | ID_4 |
| 21days | RF__min | 21.1234341 | ID_5 |
| 21days | RF__min | 17.4537046 | ID_6 |
| 21days | RF__min | 21.6309296 | ID_7 |
| 21days | RF__min | 13.7761458 | ID_8 |
| 21days | RF__min | 17.7483715 | ID_9 |
| 21days | RF__min | 24.29959 | ID_10 |

**Suppl. Table 17: Summary of bottom angle analysis**

| day | Measure | Mean | SD |
|---|---|---|---|
| 0days | LB__avg_Angle | 47.75408182 | 3.3690005 |
| 3days | LB__avg_Angle | 50.15381894 | 4.33437741 |
| 7days | LB__avg_Angle | 50.21187711 | 2.8908405 |
| 14days | LB__avg_Angle | 51.00199275 | 2.75730422 |
| 21days | LB__avg_Angle | 49.7391305 | 4.06844479 |
| 0days | LB__max | 74.63662441 | 3.88910043 |
| 3days | LB__max | 75.46730787 | 3.88677426 |
| 7days | LB__max | 73.87982019 | 3.265263 |
| 14days | LB__max | 74.52318194 | 3.51678681 |
| 21days | LB__max | 75.60318105 | 3.12537027 |
| 0days | LB__min | 15.40789053 | 3.83166016 |
| 3days | LB__min | 18.12057878 | 5.21855366 |
| 7days | LB__min | 14.90424509 | 4.87824285 |
| 14days | LB__min | 16.16849516 | 3.10961442 |
| 21days | LB__min | 15.5199302 | 4.45887447 |
| 0days | LF__avg_Angle | 43.23965924 | 2.78945572 |
| 3days | LF__avg_Angle | 41.56378715 | 2.56394347 |
| 7days | LF__avg_Angle | 43.60743345 | 2.30625999 |
| 14days | LF__avg_Angle | 40.41572586 | 3.10462092 |
| 21days | LF__avg_Angle | 42.43371163 | 2.73292041 |
| 0days | LF__max | 70.334682 | 3.92349668 |
| 3days | LF__max | 65.47597481 | 2.89340866 |
| 7days | LF__max | 67.73704872 | 5.4002759 |
| 14days | LF__max | 64.580843 | 5.10775037 |
| 21days | LF__max | 67.38188278 | 4.19884365 |
| 0days | LF__min | 19.28803726 | 3.91814991 |
| 3days | LF__min | 20.99896488 | 4.13953225 |
| 7days | LF__min | 21.01150822 | 5.24331798 |
| 14days | LF__min | 20.04743327 | 3.13943973 |
| 21days | LF__min | 19.0104686 | 3.10427479 |
| 0days | RB__avg_Angle | 47.20771682 | 2.66120631 |
| 3days | RB__avg_Angle | 52.66719351 | 3.00065815 |
| 7days | RB__avg_Angle | 48.20164253 | 4.17102957 |
| 14days | RB__avg_Angle | 49.40569593 | 4.42802183 |
| 21days | RB__avg_Angle | 49.30509406 | 3.10780087 |
| 0days | RB__max | 75.10485648 | 2.85876929 |
| 3days | RB__max | 75.7384664 | 3.66409559 |
| 7days | RB__max | 74.44480626 | 3.37197267 |
| 14days | RB__max | 75.75644489 | 3.61049547 |
| 21days | RB__max | 74.6792608 | 3.78181284 |
| 0days | RB__min | 13.53705253 | 2.78885881 |
| 3days | RB__min | 16.42683058 | 5.01051585 |
| 7days | RB__min | 14.22829565 | 3.87040931 |
| 14days | RB__min | 15.60583578 | 2.49934934 |
| 21days | RB__min | 16.28058394 | 4.01370633 |
| 0days | RF__avg_Angle | 41.71932383 | 2.822009 |
| 3days | RF__avg_Angle | 40.31542859 | 4.08163483 |
| 7days | RF__avg_Angle | 42.87781238 | 1.28772978 |
| 14days | RF__avg_Angle | 42.24951723 | 3.32277835 |
| 21days | RF__avg_Angle | 42.95202551 | 2.63924393 |
| 0days | RF__max | 65.08564342 | 1.94780001 |
| 3days | RF__max | 62.50534375 | 4.00019755 |
| 7days | RF__max | 67.52330968 | 5.25149054 |
| 14days | RF__max | 67.56914894 | 3.91888757 |

| | | | |
|---|---|---|---|
| 21days | RF__max | 66.01549344 | 3.30116977 |
| 0days | RF__min | 20.15650845 | 2.92822757 |
| 3days | RF__min | 20.54705866 | 3.71679916 |
| 7days | RF__min | 19.33703122 | 2.95847428 |
| 14days | RF__min | 17.6105183 | 4.61167268 |
| 21days | RF__min | 19.89346194 | 3.5592443 |

**Suppl. Table 18: Statistical test of bottom angle analysis**

| Measure | group1 | group2 | p.adj | sig | Test* |
|---|---|---|---|---|---|
| LB__avg_Ang | 0days | 3days | 0.35602904 | ns | Repeat ANOVA + PH test |
| LB__avg_Ang | 0days | 7days | 0.35602904 | ns | Repeat ANOVA + PH test |
| LB__avg_Ang | 0days | 14days | 0.1169572 | ns | Repeat ANOVA + PH test |
| LB__avg_Ang | 0days | 21days | 0.35602904 | ns | Repeat ANOVA + PH test |
| LB__max2 | 0days | 3days | 1 | ns | Repeat ANOVA + PH test |
| LB__max2 | 0days | 7days | 1 | ns | Repeat ANOVA + PH test |
| LB__max2 | 0days | 14days | 1 | ns | Repeat ANOVA + PH test |
| LB__max2 | 0days | 21days | 1 | ns | Repeat ANOVA + PH test |
| LB__min2 | 0days | 3days | 0.3863356 | ns | Repeat ANOVA + PH test |
| LB__min2 | 0days | 7days | 1 | ns | Repeat ANOVA + PH test |
| LB__min2 | 0days | 14days | 1 | ns | Repeat ANOVA + PH test |
| LB__min2 | 0days | 21days | 1 | ns | Repeat ANOVA + PH test |
| LF__avg_Ang | 0days | 3days | 0.91277765 | ns | Repeat ANOVA + PH test |
| LF__avg_Ang | 0days | 7days | 1 | ns | Repeat ANOVA + PH test |
| LF__avg_Ang | 0days | 14days | 0.23150504 | ns | Repeat ANOVA + PH test |
| LF__avg_Ang | 0days | 21days | 1 | ns | Repeat ANOVA + PH test |
| LF__max2 | 0days | 3days | 0.00899179 | ** | Repeat ANOVA + PH test |
| LF__max2 | 0days | 7days | 0.10685635 | ns | Repeat ANOVA + PH test |
| LF__max2 | 0days | 14days | 0.00048251 | *** | Repeat ANOVA + PH test |
| LF__max2 | 0days | 21days | 0.10685635 | ns | Repeat ANOVA + PH test |
| LF__min2 | 0days | 3days | 1 | ns | Repeat ANOVA + PH test |
| LF__min2 | 0days | 7days | 1 | ns | Repeat ANOVA + PH test |
| LF__min2 | 0days | 14days | 1 | ns | Repeat ANOVA + PH test |
| LF__min2 | 0days | 21days | 1 | ns | Repeat ANOVA + PH test |
| RB__avg_Ang | 0days | 3days | 0.0034547 | ** | Repeat ANOVA + PH test |
| RB__avg_Ang | 0days | 7days | 0.52765777 | ns | Repeat ANOVA + PH test |
| RB__avg_Ang | 0days | 14days | 0.41888756 | ns | Repeat ANOVA + PH test |
| RB__avg_Ang | 0days | 21days | 0.41888756 | ns | Repeat ANOVA + PH test |
| RB__max2 | 0days | 3days | 1 | ns | Repeat ANOVA + PH test |
| RB__max2 | 0days | 7days | 1 | ns | Repeat ANOVA + PH test |
| RB__max2 | 0days | 14days | 1 | ns | Repeat ANOVA + PH test |
| RB__max2 | 0days | 21days | 1 | ns | Repeat ANOVA + PH test |
| RB__min2 | 0days | 3days | 0.29063093 | ns | Repeat ANOVA + PH test |
| RB__min2 | 0days | 7days | 0.6604527 | ns | Repeat ANOVA + PH test |
| RB__min2 | 0days | 14days | 0.32875256 | ns | Repeat ANOVA + PH test |
| RB__min2 | 0days | 21days | 0.29063093 | ns | Repeat ANOVA + PH test |
| RF__avg_Ang | 0days | 3days | 1 | ns | Repeat ANOVA + PH test |
| RF__avg_Ang | 0days | 7days | 1 | ns | Repeat ANOVA + PH test |
| RF__avg_Ang | 0days | 14days | 1 | ns | Repeat ANOVA + PH test |
| RF__avg_Ang | 0days | 21days | 1 | ns | Repeat ANOVA + PH test |
| RF__max2 | 0days | 3days | 0.38078015 | ns | Repeat ANOVA + PH test |
| RF__max2 | 0days | 7days | 0.38078015 | ns | Repeat ANOVA + PH test |
| RF__max2 | 0days | 14days | 0.38078015 | ns | Repeat ANOVA + PH test |
| RF__max2 | 0days | 21days | 0.5424264 | ns | Repeat ANOVA + PH test |
| RF__min2 | 0days | 3days | 1 | ns | Repeat ANOVA + PH test |
| RF__min2 | 0days | 7days | 1 | ns | Repeat ANOVA + PH test |
| RF__min2 | 0days | 14days | 0.34870829 | ns | Repeat ANOVA + PH test |
| RF__min2 | 0days | 21days | 1 | ns | Repeat ANOVA + PH test |

**Suppl. Table 19: Raw data side perspective height analysis**

| day | Measure | Value | ID |
|---|---|---|---|
| 0days | l-back-ankle__Average_Heig | -0.558372 | ID_1 |
| 0days | l-back-ankle__Average_Heig | -0.0928311 | ID_2 |
| 0days | l-back-ankle__Average_Heig | -0.0743401 | ID_3 |
| 0days | l-back-ankle__Average_Heig | 0 | ID_4 |
| 0days | l-back-ankle__Average_Heig | 1.04525235 | ID_5 |
| 0days | l-back-ankle__Average_Heig | 0.46756773 | ID_6 |
| 0days | l-back-ankle__Average_Heig | -0.306591 | ID_7 |
| 0days | l-back-ankle__Average_Heig | -0.8661789 | ID_8 |
| 0days | l-back-ankle__Average_Heig | 0.49640185 | ID_9 |
| 0days | l-back-ankle__Average_Heig | -0.5475319 | ID_10 |
| 0days | l-back-ankle__Average_Heig | 1.5837189 | ID_11 |
| 0days | l-back-ankle__Average_Heig | 0.12513895 | ID_12 |
| 0days | l-back-ankle__Average_Heig | 0.04543175 | ID_13 |
| 3days | l-back-ankle__Average_Heig | 1.12716089 | ID_1 |
| 3days | l-back-ankle__Average_Heig | 0.01360356 | ID_2 |
| 3days | l-back-ankle__Average_Heig | 0.17252705 | ID_3 |
| 3days | l-back-ankle__Average_Heig | 0.0378154 | ID_4 |
| 3days | l-back-ankle__Average_Heig | -0.5515274 | ID_5 |
| 3days | l-back-ankle__Average_Heig | -0.614162 | ID_6 |
| 3days | l-back-ankle__Average_Heig | 0.68601229 | ID_7 |
| 3days | l-back-ankle__Average_Heig | -0.1506365 | ID_8 |
| 7days | l-back-ankle__Average_Heig | -0.5815902 | ID_1 |
| 7days | l-back-ankle__Average_Heig | -0.6634506 | ID_2 |
| 7days | l-back-ankle__Average_Heig | -0.4829293 | ID_3 |
| 7days | l-back-ankle__Average_Heig | -0.3621267 | ID_4 |
| 7days | l-back-ankle__Average_Heig | -0.1706575 | ID_5 |
| 7days | l-back-ankle__Average_Heig | 0.13531241 | ID_6 |
| 7days | l-back-ankle__Average_Heig | 0.49527216 | ID_7 |
| 7days | l-back-ankle__Average_Heig | -0.351378 | ID_8 |
| 7days | l-back-ankle__Average_Heig | 0.06333767 | ID_9 |
| 14days | l-back-ankle__Average_Heig | -0.076288 | ID_1 |
| 14days | l-back-ankle__Average_Heig | 0.15133667 | ID_2 |
| 14days | l-back-ankle__Average_Heig | -0.1810717 | ID_3 |
| 14days | l-back-ankle__Average_Heig | 0.15781205 | ID_4 |
| 14days | l-back-ankle__Average_Heig | 0.0320646 | ID_5 |
| 14days | l-back-ankle__Average_Heig | 1.71953848 | ID_6 |
| 14days | l-back-ankle__Average_Heig | -0.032379 | ID_7 |
| 14days | l-back-ankle__Average_Heig | 0.19315873 | ID_8 |
| 14days | l-back-ankle__Average_Heig | -0.499656 | ID_9 |
| 14days | l-back-ankle__Average_Heig | 0.50124279 | ID_10 |
| 14days | l-back-ankle__Average_Heig | -0.5566281 | ID_11 |
| 21days | l-back-ankle__Average_Heig | -0.0485212 | ID_1 |
| 21days | l-back-ankle__Average_Heig | 1.00486578 | ID_2 |
| 21days | l-back-ankle__Average_Heig | 1.55559221 | ID_3 |
| 21days | l-back-ankle__Average_Heig | -0.1152101 | ID_4 |
| 21days | l-back-ankle__Average_Heig | -0.345291 | ID_5 |
| 21days | l-back-ankle__Average_Heig | -0.527633 | ID_6 |
| 21days | l-back-ankle__Average_Heig | -0.9426903 | ID_7 |
| 21days | l-back-ankle__Average_Heig | -0.4602843 | ID_8 |
| 21days | l-back-ankle__Average_Heig | -0.7781565 | ID_9 |
| 21days | l-back-ankle__Average_Heig | 0.49855778 | ID_10 |
| 0days | l-back-ankle__Movement | -0.9914844 | ID_1 |
| 0days | l-back-ankle__Movement | 0.02306694 | ID_2 |
| 0days | l-back-ankle__Movement | -0.2829376 | ID_3 |

| | | | |
|---|---|---|---|
| 0days | l-back-ankle__Movement | 0.74780835 | ID_4 |
| 0days | l-back-ankle__Movement | 1.85085199 | ID_5 |
| 0days | l-back-ankle__Movement | 2.32714528 | ID_6 |
| 0days | l-back-ankle__Movement | -0.681279 | ID_7 |
| 0days | l-back-ankle__Movement | -2.3847816 | ID_8 |
| 0days | l-back-ankle__Movement | 1.82626031 | ID_9 |
| 0days | l-back-ankle__Movement | -1.0159964 | ID_10 |
| 0days | l-back-ankle__Movement | 1.79036629 | ID_11 |
| 0days | l-back-ankle__Movement | 0 | ID_12 |
| 0days | l-back-ankle__Movement | -0.1161506 | ID_13 |
| 3days | l-back-ankle__Movement | 0.21187297 | ID_1 |
| 3days | l-back-ankle__Movement | 0.2124552 | ID_2 |
| 3days | l-back-ankle__Movement | 0.27917367 | ID_3 |
| 3days | l-back-ankle__Movement | -1.2343567 | ID_4 |
| 3days | l-back-ankle__Movement | -2.0757384 | ID_5 |
| 3days | l-back-ankle__Movement | -1.9038635 | ID_6 |
| 3days | l-back-ankle__Movement | 1.32226482 | ID_7 |
| 3days | l-back-ankle__Movement | -0.1723685 | ID_8 |
| 7days | l-back-ankle__Movement | -0.9425015 | ID_1 |
| 7days | l-back-ankle__Movement | -0.9437131 | ID_2 |
| 7days | l-back-ankle__Movement | -0.8073341 | ID_3 |
| 7days | l-back-ankle__Movement | -0.6768367 | ID_4 |
| 7days | l-back-ankle__Movement | -0.6315821 | ID_5 |
| 7days | l-back-ankle__Movement | 0.10237663 | ID_6 |
| 7days | l-back-ankle__Movement | 0.62162624 | ID_7 |
| 7days | l-back-ankle__Movement | -0.1545461 | ID_8 |
| 7days | l-back-ankle__Movement | -0.0778493 | ID_9 |
| 14days | l-back-ankle__Movement | 1.28366887 | ID_1 |
| 14days | l-back-ankle__Movement | 0.20422954 | ID_2 |
| 14days | l-back-ankle__Movement | -0.2875744 | ID_3 |
| 14days | l-back-ankle__Movement | 0.37955496 | ID_4 |
| 14days | l-back-ankle__Movement | 0.60992437 | ID_5 |
| 14days | l-back-ankle__Movement | 2.32838002 | ID_6 |
| 14days | l-back-ankle__Movement | 0.2006356 | ID_7 |
| 14days | l-back-ankle__Movement | 0.27062048 | ID_8 |
| 14days | l-back-ankle__Movement | -0.9489289 | ID_9 |
| 14days | l-back-ankle__Movement | 0.48324093 | ID_10 |
| 14days | l-back-ankle__Movement | -0.0334763 | ID_11 |
| 21days | l-back-ankle__Movement | -1.1024072 | ID_1 |
| 21days | l-back-ankle__Movement | 0.82531936 | ID_2 |
| 21days | l-back-ankle__Movement | 1.64836129 | ID_3 |
| 21days | l-back-ankle__Movement | -0.6164052 | ID_4 |
| 21days | l-back-ankle__Movement | -0.8705098 | ID_5 |
| 21days | l-back-ankle__Movement | -1.5564573 | ID_6 |
| 21days | l-back-ankle__Movement | -2.2947539 | ID_7 |
| 21days | l-back-ankle__Movement | -0.1685381 | ID_8 |
| 21days | l-back-ankle__Movement | -1.5300792 | ID_9 |
| 21days | l-back-ankle__Movement | 0.1785303 | ID_10 |
| 0days | l-back-toe__Average_Height | -0.2007638 | ID_1 |
| 0days | l-back-toe__Average_Height | -0.0731845 | ID_2 |
| 0days | l-back-toe__Average_Height | -0.1845622 | ID_3 |
| 0days | l-back-toe__Average_Height | 0.1274839 | ID_4 |
| 0days | l-back-toe__Average_Height | -0.3949807 | ID_5 |
| 0days | l-back-toe__Average_Height | 0.14552974 | ID_6 |
| 0days | l-back-toe__Average_Height | 0.25121786 | ID_7 |
| 0days | l-back-toe__Average_Height | -0.5706094 | ID_8 |

| | | | |
|---|---|---|---|
| 0days | l-back-toe__Average_Height | -0.2047977 | ID_9 |
| 0days | l-back-toe__Average_Height | 0 | ID_10 |
| 0days | l-back-toe__Average_Height | 0.27136664 | ID_11 |
| 0days | l-back-toe__Average_Height | 0.50930984 | ID_12 |
| 0days | l-back-toe__Average_Height | 0.33806656 | ID_13 |
| 3days | l-back-toe__Average_Height | -0.1667614 | ID_1 |
| 3days | l-back-toe__Average_Height | -0.2929109 | ID_2 |
| 3days | l-back-toe__Average_Height | -0.2101754 | ID_3 |
| 3days | l-back-toe__Average_Height | -0.3827026 | ID_4 |
| 3days | l-back-toe__Average_Height | -0.3570359 | ID_5 |
| 3days | l-back-toe__Average_Height | -0.1997423 | ID_6 |
| 3days | l-back-toe__Average_Height | -0.4044975 | ID_7 |
| 3days | l-back-toe__Average_Height | -0.1764409 | ID_8 |
| 7days | l-back-toe__Average_Height | -0.3629967 | ID_1 |
| 7days | l-back-toe__Average_Height | -0.3695886 | ID_2 |
| 7days | l-back-toe__Average_Height | -0.149925 | ID_3 |
| 7days | l-back-toe__Average_Height | -0.2441127 | ID_4 |
| 7days | l-back-toe__Average_Height | -0.1580983 | ID_5 |
| 7days | l-back-toe__Average_Height | -0.0772535 | ID_6 |
| 7days | l-back-toe__Average_Height | -0.2933143 | ID_7 |
| 7days | l-back-toe__Average_Height | -0.1037771 | ID_8 |
| 7days | l-back-toe__Average_Height | -0.18197 | ID_9 |
| 14days | l-back-toe__Average_Height | -0.1052261 | ID_1 |
| 14days | l-back-toe__Average_Height | -0.0422381 | ID_2 |
| 14days | l-back-toe__Average_Height | -0.2627759 | ID_3 |
| 14days | l-back-toe__Average_Height | -0.2959257 | ID_4 |
| 14days | l-back-toe__Average_Height | -0.4000789 | ID_5 |
| 14days | l-back-toe__Average_Height | -0.2026806 | ID_6 |
| 14days | l-back-toe__Average_Height | -0.1521897 | ID_7 |
| 14days | l-back-toe__Average_Height | -0.0435699 | ID_8 |
| 14days | l-back-toe__Average_Height | -0.2636383 | ID_9 |
| 14days | l-back-toe__Average_Height | -0.1224169 | ID_10 |
| 14days | l-back-toe__Average_Height | 0.21770285 | ID_11 |
| 21days | l-back-toe__Average_Height | -0.0608787 | ID_1 |
| 21days | l-back-toe__Average_Height | 0.27231807 | ID_2 |
| 21days | l-back-toe__Average_Height | -0.0697866 | ID_3 |
| 21days | l-back-toe__Average_Height | -0.0390446 | ID_4 |
| 21days | l-back-toe__Average_Height | -0.0497526 | ID_5 |
| 21days | l-back-toe__Average_Height | -0.1145241 | ID_6 |
| 21days | l-back-toe__Average_Height | -0.1393812 | ID_7 |
| 21days | l-back-toe__Average_Height | -0.1410475 | ID_8 |
| 21days | l-back-toe__Average_Height | 0.13321495 | ID_9 |
| 21days | l-back-toe__Average_Height | 0.31476302 | ID_10 |
| 0days | l-back-toe__Movement | -0.891783 | ID_1 |
| 0days | l-back-toe__Movement | -0.912764 | ID_2 |
| 0days | l-back-toe__Movement | -0.6658001 | ID_3 |
| 0days | l-back-toe__Movement | 1.48389972 | ID_4 |
| 0days | l-back-toe__Movement | 1.2754539 | ID_5 |
| 0days | l-back-toe__Movement | 0 | ID_6 |
| 0days | l-back-toe__Movement | 0.19085782 | ID_7 |
| 0days | l-back-toe__Movement | -2.7356388 | ID_8 |
| 0days | l-back-toe__Movement | -0.6101814 | ID_9 |
| 0days | l-back-toe__Movement | -0.934677 | ID_10 |
| 0days | l-back-toe__Movement | 0.57495879 | ID_11 |
| 0days | l-back-toe__Movement | 0.70774739 | ID_12 |
| 0days | l-back-toe__Movement | 0.51821809 | ID_13 |

| | | | |
|---|---|---|---|
| 3days | l-back-toe__Movement | -1.3070101 | ID_1 |
| 3days | l-back-toe__Movement | -0.950261 | ID_2 |
| 3days | l-back-toe__Movement | -1.375857 | ID_3 |
| 3days | l-back-toe__Movement | -1.0712494 | ID_4 |
| 3days | l-back-toe__Movement | -1.4527488 | ID_5 |
| 3days | l-back-toe__Movement | -0.7754522 | ID_6 |
| 3days | l-back-toe__Movement | -1.0120016 | ID_7 |
| 3days | l-back-toe__Movement | -1.684607 | ID_8 |
| 7days | l-back-toe__Movement | -0.9393896 | ID_1 |
| 7days | l-back-toe__Movement | -1.9767143 | ID_2 |
| 7days | l-back-toe__Movement | -0.5922708 | ID_3 |
| 7days | l-back-toe__Movement | -1.1917231 | ID_4 |
| 7days | l-back-toe__Movement | -1.3039235 | ID_5 |
| 7days | l-back-toe__Movement | -0.5461987 | ID_6 |
| 7days | l-back-toe__Movement | -1.1634658 | ID_7 |
| 7days | l-back-toe__Movement | -1.44731 | ID_8 |
| 7days | l-back-toe__Movement | -1.3614519 | ID_9 |
| 14days | l-back-toe__Movement | 0.41138077 | ID_1 |
| 14days | l-back-toe__Movement | 0.33586402 | ID_2 |
| 14days | l-back-toe__Movement | -1.4108377 | ID_3 |
| 14days | l-back-toe__Movement | -1.6355577 | ID_4 |
| 14days | l-back-toe__Movement | -1.5974239 | ID_5 |
| 14days | l-back-toe__Movement | -0.0730451 | ID_6 |
| 14days | l-back-toe__Movement | -0.7894132 | ID_7 |
| 14days | l-back-toe__Movement | -1.0419263 | ID_8 |
| 14days | l-back-toe__Movement | -2.0363561 | ID_9 |
| 14days | l-back-toe__Movement | -0.8431799 | ID_10 |
| 14days | l-back-toe__Movement | -0.2961896 | ID_11 |
| 21days | l-back-toe__Movement | -0.7334589 | ID_1 |
| 21days | l-back-toe__Movement | 0.16867439 | ID_2 |
| 21days | l-back-toe__Movement | -0.3295037 | ID_3 |
| 21days | l-back-toe__Movement | -1.3069666 | ID_4 |
| 21days | l-back-toe__Movement | -0.4545942 | ID_5 |
| 21days | l-back-toe__Movement | -1.5645656 | ID_6 |
| 21days | l-back-toe__Movement | -1.7102679 | ID_7 |
| 21days | l-back-toe__Movement | -1.3163913 | ID_8 |
| 21days | l-back-toe__Movement | -0.2712721 | ID_9 |
| 21days | l-back-toe__Movement | -0.007773 | ID_10 |
| 0days | l-elbow__Average_Height | -0.2393984 | ID_1 |
| 0days | l-elbow__Average_Height | -0.0606596 | ID_2 |
| 0days | l-elbow__Average_Height | 0.05889582 | ID_3 |
| 0days | l-elbow__Average_Height | 0.05348408 | ID_4 |
| 0days | l-elbow__Average_Height | -0.2840598 | ID_5 |
| 0days | l-elbow__Average_Height | 0.08663327 | ID_6 |
| 0days | l-elbow__Average_Height | 0.86839951 | ID_7 |
| 0days | l-elbow__Average_Height | 0.13338148 | ID_8 |
| 0days | l-elbow__Average_Height | 0.10046729 | ID_9 |
| 0days | l-elbow__Average_Height | -0.2225656 | ID_10 |
| 0days | l-elbow__Average_Height | -0.2256471 | ID_11 |
| 0days | l-elbow__Average_Height | -0.1043292 | ID_12 |
| 0days | l-elbow__Average_Height | 0 | ID_13 |
| 3days | l-elbow__Average_Height | -0.0945403 | ID_1 |
| 3days | l-elbow__Average_Height | -0.2066039 | ID_2 |
| 3days | l-elbow__Average_Height | -0.3239991 | ID_3 |
| 3days | l-elbow__Average_Height | -0.2082555 | ID_4 |
| 3days | l-elbow__Average_Height | -0.0155349 | ID_5 |

| | | | |
|---|---|---|---|
| 3days | l-elbow__Average_Height | -0.2428056 | ID_6 |
| 3days | l-elbow__Average_Height | -0.1569429 | ID_7 |
| 3days | l-elbow__Average_Height | -0.0066386 | ID_8 |
| 7days | l-elbow__Average_Height | 0.16589829 | ID_1 |
| 7days | l-elbow__Average_Height | -0.0325917 | ID_2 |
| 7days | l-elbow__Average_Height | -0.2381256 | ID_3 |
| 7days | l-elbow__Average_Height | -0.1196973 | ID_4 |
| 7days | l-elbow__Average_Height | -0.3797344 | ID_5 |
| 7days | l-elbow__Average_Height | 0.06673845 | ID_6 |
| 7days | l-elbow__Average_Height | 0.16629383 | ID_7 |
| 7days | l-elbow__Average_Height | -0.1595892 | ID_8 |
| 7days | l-elbow__Average_Height | -0.2987942 | ID_9 |
| 14days | l-elbow__Average_Height | -0.1311944 | ID_1 |
| 14days | l-elbow__Average_Height | -0.2710208 | ID_2 |
| 14days | l-elbow__Average_Height | 0.21045108 | ID_3 |
| 14days | l-elbow__Average_Height | 0.51704939 | ID_4 |
| 14days | l-elbow__Average_Height | 0.33233005 | ID_5 |
| 14days | l-elbow__Average_Height | -0.1913757 | ID_6 |
| 14days | l-elbow__Average_Height | 0.7371687 | ID_7 |
| 14days | l-elbow__Average_Height | 0.51850075 | ID_8 |
| 14days | l-elbow__Average_Height | -0.1202261 | ID_9 |
| 14days | l-elbow__Average_Height | 0.17608252 | ID_10 |
| 14days | l-elbow__Average_Height | 0.14184654 | ID_11 |
| 21days | l-elbow__Average_Height | -0.22229 | ID_1 |
| 21days | l-elbow__Average_Height | -0.2523522 | ID_2 |
| 21days | l-elbow__Average_Height | 0.00876878 | ID_3 |
| 21days | l-elbow__Average_Height | 0.97198922 | ID_4 |
| 21days | l-elbow__Average_Height | 0.45282924 | ID_5 |
| 21days | l-elbow__Average_Height | 0.12554679 | ID_6 |
| 21days | l-elbow__Average_Height | 0.63911243 | ID_7 |
| 21days | l-elbow__Average_Height | 0.5608876 | ID_8 |
| 21days | l-elbow__Average_Height | 0.18487568 | ID_9 |
| 0days | l-elbow__Movement | -0.3591778 | ID_1 |
| 0days | l-elbow__Movement | -0.1485998 | ID_2 |
| 0days | l-elbow__Movement | -0.250601 | ID_3 |
| 0days | l-elbow__Movement | 0.06269647 | ID_4 |
| 0days | l-elbow__Movement | -0.5882387 | ID_5 |
| 0days | l-elbow__Movement | 0.36670743 | ID_6 |
| 0days | l-elbow__Movement | 1.46395694 | ID_7 |
| 0days | l-elbow__Movement | 0.72611005 | ID_8 |
| 0days | l-elbow__Movement | 0.07244689 | ID_9 |
| 0days | l-elbow__Movement | 0 | ID_10 |
| 0days | l-elbow__Movement | -0.6767126 | ID_11 |
| 0days | l-elbow__Movement | -0.2700936 | ID_12 |
| 0days | l-elbow__Movement | 0.06863677 | ID_13 |
| 3days | l-elbow__Movement | -0.4701718 | ID_1 |
| 3days | l-elbow__Movement | -0.869143 | ID_2 |
| 3days | l-elbow__Movement | -0.874188 | ID_3 |
| 3days | l-elbow__Movement | -0.5699819 | ID_4 |
| 3days | l-elbow__Movement | -0.4102257 | ID_5 |
| 3days | l-elbow__Movement | -0.7628235 | ID_6 |
| 3days | l-elbow__Movement | -0.8789411 | ID_7 |
| 3days | l-elbow__Movement | -0.3000482 | ID_8 |
| 7days | l-elbow__Movement | -0.110808 | ID_1 |
| 7days | l-elbow__Movement | 0.02819215 | ID_2 |
| 7days | l-elbow__Movement | -0.8159358 | ID_3 |

| | | | |
|---|---|---|---|
| 7days | l-elbow__Movement | -0.2017046 | ID_4 |
| 7days | l-elbow__Movement | -0.8158182 | ID_5 |
| 7days | l-elbow__Movement | 1.26708913 | ID_6 |
| 7days | l-elbow__Movement | 0.06676947 | ID_7 |
| 7days | l-elbow__Movement | -0.5380342 | ID_8 |
| 7days | l-elbow__Movement | -0.5854277 | ID_9 |
| 14days | l-elbow__Movement | -0.3594977 | ID_1 |
| 14days | l-elbow__Movement | -0.5746941 | ID_2 |
| 14days | l-elbow__Movement | 0.70404172 | ID_3 |
| 14days | l-elbow__Movement | 1.48744661 | ID_4 |
| 14days | l-elbow__Movement | 0.74460189 | ID_5 |
| 14days | l-elbow__Movement | -0.4975155 | ID_6 |
| 14days | l-elbow__Movement | 1.19959825 | ID_7 |
| 14days | l-elbow__Movement | 0.91943454 | ID_8 |
| 14days | l-elbow__Movement | -0.4305603 | ID_9 |
| 14days | l-elbow__Movement | 0.77137131 | ID_10 |
| 14days | l-elbow__Movement | 0.12683158 | ID_11 |
| 21days | l-elbow__Movement | -0.4734081 | ID_1 |
| 21days | l-elbow__Movement | -0.2018055 | ID_2 |
| 21days | l-elbow__Movement | -0.4027433 | ID_3 |
| 21days | l-elbow__Movement | 2.29973789 | ID_4 |
| 21days | l-elbow__Movement | 0.62506938 | ID_5 |
| 21days | l-elbow__Movement | 0.26455313 | ID_6 |
| 21days | l-elbow__Movement | 1.56354487 | ID_7 |
| 21days | l-elbow__Movement | 1.2242387 | ID_8 |
| 21days | l-elbow__Movement | 0.02430408 | ID_9 |
| 0days | l-front-toe-tip__Average_Hei | -0.0147911 | ID_1 |
| 0days | l-front-toe-tip__Average_Hei | -0.4672062 | ID_2 |
| 0days | l-front-toe-tip__Average_Hei | 0.13754486 | ID_3 |
| 0days | l-front-toe-tip__Average_Hei | -0.2127102 | ID_4 |
| 0days | l-front-toe-tip__Average_Hei | -0.2197684 | ID_5 |
| 0days | l-front-toe-tip__Average_Hei | 0.11758264 | ID_6 |
| 0days | l-front-toe-tip__Average_Hei | 0.09879332 | ID_7 |
| 0days | l-front-toe-tip__Average_Hei | 0.01056145 | ID_8 |
| 0days | l-front-toe-tip__Average_Hei | -0.0771106 | ID_9 |
| 0days | l-front-toe-tip__Average_Hei | 0 | ID_10 |
| 0days | l-front-toe-tip__Average_Hei | -0.3761377 | ID_11 |
| 0days | l-front-toe-tip__Average_Hei | 0.15583334 | ID_12 |
| 0days | l-front-toe-tip__Average_Hei | 0.2226021 | ID_13 |
| 3days | l-front-toe-tip__Average_Hei | -0.5414452 | ID_1 |
| 3days | l-front-toe-tip__Average_Hei | -0.3469393 | ID_2 |
| 3days | l-front-toe-tip__Average_Hei | -0.188103 | ID_3 |
| 3days | l-front-toe-tip__Average_Hei | -0.6663616 | ID_4 |
| 3days | l-front-toe-tip__Average_Hei | -0.5568749 | ID_5 |
| 3days | l-front-toe-tip__Average_Hei | -0.3228904 | ID_6 |
| 3days | l-front-toe-tip__Average_Hei | -0.5251257 | ID_7 |
| 3days | l-front-toe-tip__Average_Hei | -0.4537633 | ID_8 |
| 7days | l-front-toe-tip__Average_Hei | -0.3615072 | ID_1 |
| 7days | l-front-toe-tip__Average_Hei | -0.4075054 | ID_2 |
| 7days | l-front-toe-tip__Average_Hei | -0.3830309 | ID_3 |
| 7days | l-front-toe-tip__Average_Hei | -0.3858967 | ID_4 |
| 7days | l-front-toe-tip__Average_Hei | -0.269788 | ID_5 |
| 7days | l-front-toe-tip__Average_Hei | -0.4060665 | ID_6 |
| 7days | l-front-toe-tip__Average_Hei | -0.4506956 | ID_7 |
| 7days | l-front-toe-tip__Average_Hei | -0.2462014 | ID_8 |
| 7days | l-front-toe-tip__Average_Hei | -0.3652331 | ID_9 |

| | | | |
|---|---|---|---|
| 14days | l-front-toe-tip__Average_Hei | -0.2869397 | ID_1 |
| 14days | l-front-toe-tip__Average_Hei | -0.0884651 | ID_2 |
| 14days | l-front-toe-tip__Average_Hei | -0.4246284 | ID_3 |
| 14days | l-front-toe-tip__Average_Hei | -0.4654027 | ID_4 |
| 14days | l-front-toe-tip__Average_Hei | 0.13222713 | ID_5 |
| 14days | l-front-toe-tip__Average_Hei | -0.646435 | ID_6 |
| 14days | l-front-toe-tip__Average_Hei | -0.2675328 | ID_7 |
| 14days | l-front-toe-tip__Average_Hei | -0.2986062 | ID_8 |
| 14days | l-front-toe-tip__Average_Hei | -0.7209941 | ID_9 |
| 14days | l-front-toe-tip__Average_Hei | -0.4827357 | ID_10 |
| 14days | l-front-toe-tip__Average_Hei | 0.03143259 | ID_11 |
| 21days | l-front-toe-tip__Average_Hei | -0.3688372 | ID_1 |
| 21days | l-front-toe-tip__Average_Hei | 0.21574139 | ID_2 |
| 21days | l-front-toe-tip__Average_Hei | -0.0419055 | ID_3 |
| 21days | l-front-toe-tip__Average_Hei | 0.1045365 | ID_4 |
| 21days | l-front-toe-tip__Average_Hei | -0.0544842 | ID_5 |
| 21days | l-front-toe-tip__Average_Hei | -0.5595758 | ID_6 |
| 21days | l-front-toe-tip__Average_Hei | 0.09082528 | ID_7 |
| 21days | l-front-toe-tip__Average_Hei | -0.4457039 | ID_8 |
| 21days | l-front-toe-tip__Average_Hei | -0.3679859 | ID_9 |
| 21days | l-front-toe-tip__Average_Hei | -0.1242323 | ID_10 |
| 0days | l-front-toe-tip__Movement | -0.1104458 | ID_1 |
| 0days | l-front-toe-tip__Movement | -0.4465396 | ID_2 |
| 0days | l-front-toe-tip__Movement | 0.23189562 | ID_3 |
| 0days | l-front-toe-tip__Movement | 0.5418076 | ID_4 |
| 0days | l-front-toe-tip__Movement | -0.2357491 | ID_5 |
| 0days | l-front-toe-tip__Movement | 0.34702465 | ID_6 |
| 0days | l-front-toe-tip__Movement | 0.57335941 | ID_7 |
| 0days | l-front-toe-tip__Movement | 2.08940108 | ID_8 |
| 0days | l-front-toe-tip__Movement | 0 | ID_9 |
| 0days | l-front-toe-tip__Movement | -0.5582958 | ID_10 |
| 0days | l-front-toe-tip__Movement | -0.3919764 | ID_11 |
| 0days | l-front-toe-tip__Movement | -0.1937157 | ID_12 |
| 0days | l-front-toe-tip__Movement | 0.3474683 | ID_13 |
| 3days | l-front-toe-tip__Movement | -1.0127956 | ID_1 |
| 3days | l-front-toe-tip__Movement | -0.5421167 | ID_2 |
| 3days | l-front-toe-tip__Movement | -0.177845 | ID_3 |
| 3days | l-front-toe-tip__Movement | -0.7988863 | ID_4 |
| 3days | l-front-toe-tip__Movement | -0.7890427 | ID_5 |
| 3days | l-front-toe-tip__Movement | -0.3779705 | ID_6 |
| 3days | l-front-toe-tip__Movement | -0.7549858 | ID_7 |
| 3days | l-front-toe-tip__Movement | -1.1554824 | ID_8 |
| 7days | l-front-toe-tip__Movement | -0.6854878 | ID_1 |
| 7days | l-front-toe-tip__Movement | -0.6253576 | ID_2 |
| 7days | l-front-toe-tip__Movement | -0.5864964 | ID_3 |
| 7days | l-front-toe-tip__Movement | -0.7229588 | ID_4 |
| 7days | l-front-toe-tip__Movement | -0.8602315 | ID_5 |
| 7days | l-front-toe-tip__Movement | -0.6352478 | ID_6 |
| 7days | l-front-toe-tip__Movement | -0.9297532 | ID_7 |
| 7days | l-front-toe-tip__Movement | -0.7220486 | ID_8 |
| 7days | l-front-toe-tip__Movement | -1.0779973 | ID_9 |
| 14days | l-front-toe-tip__Movement | -0.4652339 | ID_1 |
| 14days | l-front-toe-tip__Movement | -0.419892 | ID_2 |
| 14days | l-front-toe-tip__Movement | -0.4886277 | ID_3 |
| 14days | l-front-toe-tip__Movement | -0.4357413 | ID_4 |
| 14days | l-front-toe-tip__Movement | 0.58281312 | ID_5 |

| | | | |
|---|---|---|---|
| 14days | l-front-toe-tip__Movement | -0.927662 | ID_6 |
| 14days | l-front-toe-tip__Movement | 0.16564881 | ID_7 |
| 14days | l-front-toe-tip__Movement | -0.1091076 | ID_8 |
| 14days | l-front-toe-tip__Movement | -1.4602212 | ID_9 |
| 14days | l-front-toe-tip__Movement | -0.0918883 | ID_10 |
| 14days | l-front-toe-tip__Movement | 0.38813104 | ID_11 |
| 21days | l-front-toe-tip__Movement | -0.1569518 | ID_1 |
| 21days | l-front-toe-tip__Movement | -0.1643208 | ID_2 |
| 21days | l-front-toe-tip__Movement | 0.27387035 | ID_3 |
| 21days | l-front-toe-tip__Movement | 0.71802332 | ID_4 |
| 21days | l-front-toe-tip__Movement | -0.2800393 | ID_5 |
| 21days | l-front-toe-tip__Movement | -1.1417461 | ID_6 |
| 21days | l-front-toe-tip__Movement | 0.08672806 | ID_7 |
| 21days | l-front-toe-tip__Movement | 0.04293131 | ID_8 |
| 21days | l-front-toe-tip__Movement | -1.1301173 | ID_9 |
| 21days | l-front-toe-tip__Movement | -0.2582717 | ID_10 |
| 0days | l-head__Average_Height | -0.0399667 | ID_1 |
| 0days | l-head__Average_Height | 0.44070507 | ID_2 |
| 0days | l-head__Average_Height | 1.04710834 | ID_3 |
| 0days | l-head__Average_Height | 1.78224115 | ID_4 |
| 0days | l-head__Average_Height | 0.87841401 | ID_5 |
| 0days | l-head__Average_Height | -0.2647838 | ID_6 |
| 0days | l-head__Average_Height | 0.03996673 | ID_7 |
| 0days | l-head__Average_Height | 0.52762914 | ID_8 |
| 0days | l-head__Average_Height | -0.6174724 | ID_9 |
| 0days | l-head__Average_Height | -0.1490404 | ID_10 |
| 0days | l-head__Average_Height | -0.1169046 | ID_11 |
| 0days | l-head__Average_Height | -0.3830891 | ID_12 |
| 3days | l-head__Average_Height | -0.6172452 | ID_1 |
| 3days | l-head__Average_Height | -0.7234231 | ID_2 |
| 3days | l-head__Average_Height | -0.6205401 | ID_3 |
| 3days | l-head__Average_Height | -0.4518147 | ID_4 |
| 3days | l-head__Average_Height | -0.7060306 | ID_5 |
| 3days | l-head__Average_Height | -0.3735361 | ID_6 |
| 3days | l-head__Average_Height | -0.9765378 | ID_7 |
| 3days | l-head__Average_Height | -0.1629469 | ID_8 |
| 7days | l-head__Average_Height | 0.81704756 | ID_1 |
| 7days | l-head__Average_Height | -0.630912 | ID_2 |
| 7days | l-head__Average_Height | -0.8995152 | ID_3 |
| 7days | l-head__Average_Height | -1.103179 | ID_4 |
| 7days | l-head__Average_Height | -1.1233793 | ID_5 |
| 7days | l-head__Average_Height | -0.7840083 | ID_6 |
| 7days | l-head__Average_Height | -0.7860921 | ID_7 |
| 7days | l-head__Average_Height | -0.4804879 | ID_8 |
| 7days | l-head__Average_Height | -0.6033511 | ID_9 |
| 14days | l-head__Average_Height | 0.1112093 | ID_1 |
| 14days | l-head__Average_Height | -0.3559697 | ID_2 |
| 14days | l-head__Average_Height | -0.5079412 | ID_3 |
| 14days | l-head__Average_Height | 1.54130125 | ID_4 |
| 14days | l-head__Average_Height | 0.42462363 | ID_5 |
| 14days | l-head__Average_Height | -0.2031307 | ID_6 |
| 14days | l-head__Average_Height | -0.6163382 | ID_7 |
| 14days | l-head__Average_Height | -0.6897253 | ID_8 |
| 14days | l-head__Average_Height | -0.0913937 | ID_9 |
| 14days | l-head__Average_Height | 0.23132581 | ID_10 |
| 14days | l-head__Average_Height | -0.1329957 | ID_11 |

| | | | |
|---|---|---|---|
| 21days | l-head__Average_Height | 1.36458986 | ID_1 |
| 21days | l-head__Average_Height | 0.04135524 | ID_2 |
| 21days | l-head__Average_Height | -0.6929751 | ID_3 |
| 21days | l-head__Average_Height | -0.2387244 | ID_4 |
| 21days | l-head__Average_Height | -0.325672 | ID_5 |
| 21days | l-head__Average_Height | -0.579733 | ID_6 |
| 21days | l-head__Average_Height | -0.441725 | ID_7 |
| 21days | l-head__Average_Height | -0.0284937 | ID_8 |
| 21days | l-head__Average_Height | -1.0952739 | ID_9 |
| 21days | l-head__Average_Height | -0.6293497 | ID_10 |
| 0days | l-head__Movement | -0.2423463 | ID_1 |
| 0days | l-head__Movement | 2.99432841 | ID_2 |
| 0days | l-head__Movement | 0.27923577 | ID_3 |
| 0days | l-head__Movement | 4.02776935 | ID_4 |
| 0days | l-head__Movement | 1.75034694 | ID_5 |
| 0days | l-head__Movement | -0.6740571 | ID_6 |
| 0days | l-head__Movement | 0.05166077 | ID_7 |
| 0days | l-head__Movement | 0.94065384 | ID_8 |
| 0days | l-head__Movement | -1.9499972 | ID_9 |
| 0days | l-head__Movement | -0.0516608 | ID_10 |
| 0days | l-head__Movement | -1.0674218 | ID_11 |
| 0days | l-head__Movement | -1.4630043 | ID_12 |
| 3days | l-head__Movement | -2.8558414 | ID_1 |
| 3days | l-head__Movement | -2.5586508 | ID_2 |
| 3days | l-head__Movement | -2.3237033 | ID_3 |
| 3days | l-head__Movement | -2.0557824 | ID_4 |
| 3days | l-head__Movement | -2.5158933 | ID_5 |
| 3days | l-head__Movement | -1.0614751 | ID_6 |
| 3days | l-head__Movement | -2.6521061 | ID_7 |
| 3days | l-head__Movement | -0.46177 | ID_8 |
| 7days | l-head__Movement | 1.62925398 | ID_1 |
| 7days | l-head__Movement | -1.5110295 | ID_2 |
| 7days | l-head__Movement | -2.673798 | ID_3 |
| 7days | l-head__Movement | -2.8680211 | ID_4 |
| 7days | l-head__Movement | -3.2296941 | ID_5 |
| 7days | l-head__Movement | -1.8488808 | ID_6 |
| 7days | l-head__Movement | -2.3014372 | ID_7 |
| 7days | l-head__Movement | -2.0484046 | ID_8 |
| 7days | l-head__Movement | -1.9557916 | ID_9 |
| 14days | l-head__Movement | -0.1129699 | ID_1 |
| 14days | l-head__Movement | -0.9414617 | ID_2 |
| 14days | l-head__Movement | -2.2444634 | ID_3 |
| 14days | l-head__Movement | 2.33652881 | ID_4 |
| 14days | l-head__Movement | 2.5672687 | ID_5 |
| 14days | l-head__Movement | -1.7412132 | ID_6 |
| 14days | l-head__Movement | -2.1233444 | ID_7 |
| 14days | l-head__Movement | -2.1287506 | ID_8 |
| 14days | l-head__Movement | -0.4323665 | ID_9 |
| 14days | l-head__Movement | -1.3024088 | ID_10 |
| 14days | l-head__Movement | -0.9574556 | ID_11 |
| 21days | l-head__Movement | 2.49882834 | ID_1 |
| 21days | l-head__Movement | -0.1015888 | ID_2 |
| 21days | l-head__Movement | -2.488392 | ID_3 |
| 21days | l-head__Movement | -2.0341705 | ID_4 |
| 21days | l-head__Movement | -1.1551673 | ID_5 |
| 21days | l-head__Movement | -2.6573545 | ID_6 |

| | | | |
|---|---|---|---|
| 21days | l-head__Movement | -1.6932782 | ID_7 |
| 21days | l-head__Movement | -0.9107645 | ID_8 |
| 21days | l-head__Movement | -2.940025 | ID_9 |
| 21days | l-head__Movement | -2.0159836 | ID_10 |
| 0days | l-hip__Average_Height | -0.44478 | ID_1 |
| 0days | l-hip__Average_Height | -0.0779021 | ID_2 |
| 0days | l-hip__Average_Height | 0.22000872 | ID_3 |
| 0days | l-hip__Average_Height | 0.37772816 | ID_4 |
| 0days | l-hip__Average_Height | 0.06173439 | ID_5 |
| 0days | l-hip__Average_Height | 0.28655259 | ID_6 |
| 0days | l-hip__Average_Height | 0 | ID_7 |
| 0days | l-hip__Average_Height | 0.28380395 | ID_8 |
| 0days | l-hip__Average_Height | 0.09884662 | ID_9 |
| 0days | l-hip__Average_Height | -0.3924345 | ID_10 |
| 0days | l-hip__Average_Height | -0.2661656 | ID_11 |
| 0days | l-hip__Average_Height | -0.1642707 | ID_12 |
| 0days | l-hip__Average_Height | -0.3138523 | ID_13 |
| 3days | l-hip__Average_Height | -0.4424023 | ID_1 |
| 3days | l-hip__Average_Height | -0.3450636 | ID_2 |
| 3days | l-hip__Average_Height | -0.318778 | ID_3 |
| 3days | l-hip__Average_Height | -0.3109952 | ID_4 |
| 3days | l-hip__Average_Height | -0.3595845 | ID_5 |
| 3days | l-hip__Average_Height | -0.4389856 | ID_6 |
| 3days | l-hip__Average_Height | 0.06458484 | ID_7 |
| 3days | l-hip__Average_Height | 0.28541868 | ID_8 |
| 7days | l-hip__Average_Height | -0.5057745 | ID_1 |
| 7days | l-hip__Average_Height | 0.01702515 | ID_2 |
| 7days | l-hip__Average_Height | -0.5614886 | ID_3 |
| 7days | l-hip__Average_Height | -0.2643041 | ID_4 |
| 7days | l-hip__Average_Height | 0.17797797 | ID_5 |
| 7days | l-hip__Average_Height | -0.4972085 | ID_6 |
| 7days | l-hip__Average_Height | -0.1133062 | ID_7 |
| 7days | l-hip__Average_Height | -0.4577949 | ID_8 |
| 7days | l-hip__Average_Height | 0.11693554 | ID_9 |
| 14days | l-hip__Average_Height | 0.04016729 | ID_1 |
| 14days | l-hip__Average_Height | -0.1670369 | ID_2 |
| 14days | l-hip__Average_Height | 0.33361269 | ID_3 |
| 14days | l-hip__Average_Height | -0.5270553 | ID_4 |
| 14days | l-hip__Average_Height | -0.4989137 | ID_5 |
| 14days | l-hip__Average_Height | -0.5581158 | ID_6 |
| 14days | l-hip__Average_Height | -0.5762918 | ID_7 |
| 14days | l-hip__Average_Height | -0.739587 | ID_8 |
| 14days | l-hip__Average_Height | 0.21458182 | ID_9 |
| 14days | l-hip__Average_Height | -0.4010188 | ID_10 |
| 14days | l-hip__Average_Height | -0.3005449 | ID_11 |
| 21days | l-hip__Average_Height | -0.1404379 | ID_1 |
| 21days | l-hip__Average_Height | -0.1076918 | ID_2 |
| 21days | l-hip__Average_Height | 0.17621642 | ID_3 |
| 21days | l-hip__Average_Height | -0.3595326 | ID_4 |
| 21days | l-hip__Average_Height | -0.2215566 | ID_5 |
| 21days | l-hip__Average_Height | -0.3710596 | ID_6 |
| 21days | l-hip__Average_Height | 0.26581864 | ID_7 |
| 21days | l-hip__Average_Height | -0.5029991 | ID_8 |
| 21days | l-hip__Average_Height | -0.3125981 | ID_9 |
| 21days | l-hip__Average_Height | 0.17901183 | ID_10 |
| 0days | l-hip__Movement | -0.449144 | ID_1 |

| | | | |
|---|---|---|---|
| 0days | l-hip__Movement | 0.3985917 | ID_2 |
| 0days | l-hip__Movement | 0 | ID_3 |
| 0days | l-hip__Movement | 1.12620366 | ID_4 |
| 0days | l-hip__Movement | -0.1733888 | ID_5 |
| 0days | l-hip__Movement | 1.14533099 | ID_6 |
| 0days | l-hip__Movement | 1.09662952 | ID_7 |
| 0days | l-hip__Movement | 0.53047927 | ID_8 |
| 0days | l-hip__Movement | 0.75835218 | ID_9 |
| 0days | l-hip__Movement | -0.1286163 | ID_10 |
| 0days | l-hip__Movement | -0.1952401 | ID_11 |
| 0days | l-hip__Movement | -0.2297682 | ID_12 |
| 0days | l-hip__Movement | -0.163768 | ID_13 |
| 3days | l-hip__Movement | -1.1155466 | ID_1 |
| 3days | l-hip__Movement | -0.6857796 | ID_2 |
| 3days | l-hip__Movement | -0.4757547 | ID_3 |
| 3days | l-hip__Movement | -0.3524353 | ID_4 |
| 3days | l-hip__Movement | -0.8004561 | ID_5 |
| 3days | l-hip__Movement | -0.4620348 | ID_6 |
| 3days | l-hip__Movement | 0.95327937 | ID_7 |
| 3days | l-hip__Movement | 0.5827101 | ID_8 |
| 7days | l-hip__Movement | -0.5792133 | ID_1 |
| 7days | l-hip__Movement | -0.0686231 | ID_2 |
| 7days | l-hip__Movement | -1.2099425 | ID_3 |
| 7days | l-hip__Movement | -0.577758 | ID_4 |
| 7days | l-hip__Movement | 0.17055567 | ID_5 |
| 7days | l-hip__Movement | -0.4865024 | ID_6 |
| 7days | l-hip__Movement | 0.15170056 | ID_7 |
| 7days | l-hip__Movement | -0.3605221 | ID_8 |
| 7days | l-hip__Movement | 0.36856975 | ID_9 |
| 14days | l-hip__Movement | 0.25712958 | ID_1 |
| 14days | l-hip__Movement | 0.42699317 | ID_2 |
| 14days | l-hip__Movement | -0.0199676 | ID_3 |
| 14days | l-hip__Movement | -0.964921 | ID_4 |
| 14days | l-hip__Movement | -0.444228 | ID_5 |
| 14days | l-hip__Movement | -1.1068478 | ID_6 |
| 14days | l-hip__Movement | -0.8293296 | ID_7 |
| 14days | l-hip__Movement | -0.6747009 | ID_8 |
| 14days | l-hip__Movement | 1.24007211 | ID_9 |
| 14days | l-hip__Movement | -0.0778198 | ID_10 |
| 14days | l-hip__Movement | 0.20605933 | ID_11 |
| 21days | l-hip__Movement | 0.00652735 | ID_1 |
| 21days | l-hip__Movement | 0.12645404 | ID_2 |
| 21days | l-hip__Movement | -0.126433 | ID_3 |
| 21days | l-hip__Movement | -0.6851101 | ID_4 |
| 21days | l-hip__Movement | -0.3674863 | ID_5 |
| 21days | l-hip__Movement | -0.2466754 | ID_6 |
| 21days | l-hip__Movement | 0.46760155 | ID_7 |
| 21days | l-hip__Movement | -0.6284351 | ID_8 |
| 21days | l-hip__Movement | -0.9116065 | ID_9 |
| 21days | l-hip__Movement | 1.08235313 | ID_10 |
| 0days | l-iliac-crest__Average_Height | -0.3628712 | ID_1 |
| 0days | l-iliac-crest__Average_Height | 0.13439664 | ID_2 |
| 0days | l-iliac-crest__Average_Height | 0.38431837 | ID_3 |
| 0days | l-iliac-crest__Average_Height | 0.57175904 | ID_4 |
| 0days | l-iliac-crest__Average_Height | -0.3900182 | ID_5 |
| 0days | l-iliac-crest__Average_Height | 0 | ID_6 |

| | | | |
|---|---|---|---|
| 0days | l-iliac-crest__Average_Heigh | 0.19718105 | ID_7 |
| 0days | l-iliac-crest__Average_Heigh | -0.136695 | ID_8 |
| 0days | l-iliac-crest__Average_Heigh | 0.23736064 | ID_9 |
| 0days | l-iliac-crest__Average_Heigh | -0.031464 | ID_10 |
| 0days | l-iliac-crest__Average_Heigh | -0.3975253 | ID_11 |
| 0days | l-iliac-crest__Average_Heigh | -0.0896213 | ID_12 |
| 0days | l-iliac-crest__Average_Heigh | 0.08435949 | ID_13 |
| 3days | l-iliac-crest__Average_Heigh | -0.5494283 | ID_1 |
| 3days | l-iliac-crest__Average_Heigh | -0.4668579 | ID_2 |
| 3days | l-iliac-crest__Average_Heigh | -0.2158797 | ID_3 |
| 3days | l-iliac-crest__Average_Heigh | -0.1701912 | ID_4 |
| 3days | l-iliac-crest__Average_Heigh | -0.1780643 | ID_5 |
| 3days | l-iliac-crest__Average_Heigh | -0.1401275 | ID_6 |
| 3days | l-iliac-crest__Average_Heigh | 0.21172895 | ID_7 |
| 3days | l-iliac-crest__Average_Heigh | 0.28233014 | ID_8 |
| 7days | l-iliac-crest__Average_Heigh | -0.3104302 | ID_1 |
| 7days | l-iliac-crest__Average_Heigh | -0.2572337 | ID_2 |
| 7days | l-iliac-crest__Average_Heigh | -0.5490395 | ID_3 |
| 7days | l-iliac-crest__Average_Heigh | -0.4375251 | ID_4 |
| 7days | l-iliac-crest__Average_Heigh | -0.0306272 | ID_5 |
| 7days | l-iliac-crest__Average_Heigh | -0.285001 | ID_6 |
| 7days | l-iliac-crest__Average_Heigh | 0.36952528 | ID_7 |
| 7days | l-iliac-crest__Average_Heigh | 0.00995395 | ID_8 |
| 14days | l-iliac-crest__Average_Heigh | -0.2170776 | ID_9 |
| 14days | l-iliac-crest__Average_Heigh | 0.00879214 | ID_1 |
| 14days | l-iliac-crest__Average_Heigh | -0.4679329 | ID_2 |
| 14days | l-iliac-crest__Average_Heigh | -0.5622044 | ID_3 |
| 14days | l-iliac-crest__Average_Heigh | -0.442494 | ID_4 |
| 14days | l-iliac-crest__Average_Heigh | -0.3832791 | ID_5 |
| 14days | l-iliac-crest__Average_Heigh | -0.4039169 | ID_6 |
| 14days | l-iliac-crest__Average_Heigh | -0.5298417 | ID_7 |
| 14days | l-iliac-crest__Average_Heigh | 0.38699294 | ID_8 |
| 14days | l-iliac-crest__Average_Heigh | -0.5183808 | ID_9 |
| 14days | l-iliac-crest__Average_Heigh | -0.1409395 | ID_10 |
| 21days | l-iliac-crest__Average_Heigh | 0.15711757 | ID_11 |
| 21days | l-iliac-crest__Average_Heigh | -0.0382242 | ID_1 |
| 21days | l-iliac-crest__Average_Heigh | -0.2682869 | ID_2 |
| 21days | l-iliac-crest__Average_Heigh | -0.1676372 | ID_3 |
| 21days | l-iliac-crest__Average_Heigh | 0.05191179 | ID_4 |
| 21days | l-iliac-crest__Average_Heigh | -0.1616574 | ID_5 |
| 21days | l-iliac-crest__Average_Heigh | 0.30549848 | ID_6 |
| 21days | l-iliac-crest__Average_Heigh | -0.1079949 | ID_7 |
| 21days | l-iliac-crest__Average_Heigh | -0.5673959 | ID_8 |
| 21days | l-iliac-crest__Average_Heigh | 0.65186404 | ID_9 |
| 0days | l-iliac-crest__Movement | -0.9264023 | ID_10 |
| 0days | l-iliac-crest__Movement | -0.2966488 | ID_1 |
| 0days | l-iliac-crest__Movement | 0 | ID_2 |
| 0days | l-iliac-crest__Movement | 0.96675868 | ID_3 |
| 0days | l-iliac-crest__Movement | -0.9852611 | ID_4 |
| 0days | l-iliac-crest__Movement | 0.91055 | ID_5 |
| 0days | l-iliac-crest__Movement | 0.63231218 | ID_6 |
| 0days | l-iliac-crest__Movement | -1.0445128 | ID_7 |
| 0days | l-iliac-crest__Movement | 0.80678632 | ID_8 |
| 0days | l-iliac-crest__Movement | 0.47001061 | ID_9 |
| 0days | l-iliac-crest__Movement | -1.0238567 | ID_10 |
| 0days | l-iliac-crest__Movement | -0.3866982 | ID_11 |

| | | | |
|---|---|---|---|
| 0days | l-iliac-crest__Movement | 0.08397723 | ID_12 |
| 3days | l-iliac-crest__Movement | -1.31425 | ID_13 |
| 3days | l-iliac-crest__Movement | -1.1431332 | ID_1 |
| 3days | l-iliac-crest__Movement | -0.648544 | ID_2 |
| 3days | l-iliac-crest__Movement | -0.4993067 | ID_3 |
| 3days | l-iliac-crest__Movement | -0.8674843 | ID_4 |
| 3days | l-iliac-crest__Movement | -0.7261842 | ID_5 |
| 3days | l-iliac-crest__Movement | 0.81199157 | ID_6 |
| 3days | l-iliac-crest__Movement | 0.47738176 | ID_7 |
| 7days | l-iliac-crest__Movement | -0.3812341 | ID_8 |
| 7days | l-iliac-crest__Movement | -1.2193037 | ID_1 |
| 7days | l-iliac-crest__Movement | -1.261826 | ID_2 |
| 7days | l-iliac-crest__Movement | -0.7895844 | ID_3 |
| 7days | l-iliac-crest__Movement | -0.3242382 | ID_4 |
| 7days | l-iliac-crest__Movement | -0.8407973 | ID_5 |
| 7days | l-iliac-crest__Movement | 0.71135336 | ID_6 |
| 7days | l-iliac-crest__Movement | -0.0333127 | ID_7 |
| 14days | l-iliac-crest__Movement | 0.06016784 | ID_8 |
| 14days | l-iliac-crest__Movement | 0.36320639 | ID_9 |
| 14days | l-iliac-crest__Movement | -1.5307449 | ID_1 |
| 14days | l-iliac-crest__Movement | -1.408449 | ID_2 |
| 14days | l-iliac-crest__Movement | -1.4738531 | ID_3 |
| 14days | l-iliac-crest__Movement | -1.4955974 | ID_4 |
| 14days | l-iliac-crest__Movement | -0.5683011 | ID_5 |
| 14days | l-iliac-crest__Movement | -0.6846737 | ID_6 |
| 14days | l-iliac-crest__Movement | 0.74989581 | ID_7 |
| 14days | l-iliac-crest__Movement | -0.8236276 | ID_8 |
| 14days | l-iliac-crest__Movement | -0.5397105 | ID_9 |
| 21days | l-iliac-crest__Movement | -0.1947859 | ID_10 |
| 21days | l-iliac-crest__Movement | -0.2728709 | ID_11 |
| 21days | l-iliac-crest__Movement | -0.6709748 | ID_1 |
| 21days | l-iliac-crest__Movement | -0.406304 | ID_2 |
| 21days | l-iliac-crest__Movement | -0.450563 | ID_3 |
| 21days | l-iliac-crest__Movement | -0.9414072 | ID_4 |
| 21days | l-iliac-crest__Movement | 0.18955474 | ID_5 |
| 21days | l-iliac-crest__Movement | -0.0912889 | ID_6 |
| 21days | l-iliac-crest__Movement | -1.4079614 | ID_7 |
| 21days | l-iliac-crest__Movement | 0.56774494 | ID_8 |
| 0days | l-shoulder__Average_Height | -0.2639678 | ID_1 |
| 0days | l-shoulder__Average_Height | -0.2569669 | ID_2 |
| 0days | l-shoulder__Average_Height | -0.2782261 | ID_3 |
| 0days | l-shoulder__Average_Height | 0.06219628 | ID_4 |
| 0days | l-shoulder__Average_Height | 0.33123189 | ID_5 |
| 0days | l-shoulder__Average_Height | 0.03136468 | ID_6 |
| 0days | l-shoulder__Average_Height | 0.14397045 | ID_7 |
| 0days | l-shoulder__Average_Height | 0.92531116 | ID_8 |
| 0days | l-shoulder__Average_Height | 0.24068171 | ID_9 |
| 0days | l-shoulder__Average_Height | -0.0313647 | ID_10 |
| 0days | l-shoulder__Average_Height | -0.227171 | ID_11 |
| 0days | l-shoulder__Average_Height | -0.1833853 | ID_12 |
| 3days | l-shoulder__Average_Height | -0.3847011 | ID_1 |
| 3days | l-shoulder__Average_Height | -0.2549553 | ID_2 |
| 3days | l-shoulder__Average_Height | -0.3930354 | ID_3 |
| 3days | l-shoulder__Average_Height | -0.1082167 | ID_4 |
| 3days | l-shoulder__Average_Height | 0.06229494 | ID_5 |
| 7days | l-shoulder__Average_Height | -0.1833509 | ID_1 |

| | | | |
|---|---|---|---|
| 7days | l-shoulder__Average_Height | -0.2264867 | ID_2 |
| 7days | l-shoulder__Average_Height | -0.1820852 | ID_3 |
| 7days | l-shoulder__Average_Height | -0.3811283 | ID_4 |
| 7days | l-shoulder__Average_Height | -0.3155338 | ID_5 |
| 7days | l-shoulder__Average_Height | -0.1605345 | ID_6 |
| 7days | l-shoulder__Average_Height | 0.14939833 | ID_7 |
| 7days | l-shoulder__Average_Height | -0.2664628 | ID_8 |
| 14days | l-shoulder__Average_Height | -0.0337489 | ID_1 |
| 14days | l-shoulder__Average_Height | -0.3586667 | ID_2 |
| 14days | l-shoulder__Average_Height | 0.02657494 | ID_3 |
| 14days | l-shoulder__Average_Height | 0.22786212 | ID_4 |
| 14days | l-shoulder__Average_Height | 0.00729058 | ID_5 |
| 14days | l-shoulder__Average_Height | -0.0236413 | ID_6 |
| 14days | l-shoulder__Average_Height | 0.11387469 | ID_7 |
| 14days | l-shoulder__Average_Height | 0.18105948 | ID_8 |
| 14days | l-shoulder__Average_Height | 0.2490173 | ID_9 |
| 21days | l-shoulder__Average_Height | -0.2036713 | ID_1 |
| 21days | l-shoulder__Average_Height | -0.2287081 | ID_2 |
| 21days | l-shoulder__Average_Height | -0.0517059 | ID_3 |
| 21days | l-shoulder__Average_Height | 0.17344833 | ID_4 |
| 21days | l-shoulder__Average_Height | -0.2653139 | ID_5 |
| 21days | l-shoulder__Average_Height | -0.177478 | ID_6 |
| 21days | l-shoulder__Average_Height | -0.0956549 | ID_7 |
| 21days | l-shoulder__Average_Height | 0.31921705 | ID_8 |
| 21days | l-shoulder__Average_Height | 0.01745091 | ID_9 |
| 21days | l-shoulder__Average_Height | 0.02632362 | ID_10 |
| 0days | l-shoulder__Movement | -0.4993784 | ID_1 |
| 0days | l-shoulder__Movement | -0.3641267 | ID_2 |
| 0days | l-shoulder__Movement | -0.3023594 | ID_3 |
| 0days | l-shoulder__Movement | 0.16173306 | ID_4 |
| 0days | l-shoulder__Movement | -0.0467876 | ID_5 |
| 0days | l-shoulder__Movement | 0.1527792 | ID_6 |
| 0days | l-shoulder__Movement | 0.54277713 | ID_7 |
| 0days | l-shoulder__Movement | 2.37741519 | ID_8 |
| 0days | l-shoulder__Movement | 0.0526326 | ID_9 |
| 0days | l-shoulder__Movement | 0.04678756 | ID_10 |
| 0days | l-shoulder__Movement | -0.5659054 | ID_11 |
| 0days | l-shoulder__Movement | -0.3075269 | ID_12 |
| 3days | l-shoulder__Movement | -0.9211351 | ID_1 |
| 3days | l-shoulder__Movement | -0.8850393 | ID_2 |
| 3days | l-shoulder__Movement | -0.7711989 | ID_3 |
| 3days | l-shoulder__Movement | -0.574643 | ID_4 |
| 3days | l-shoulder__Movement | -0.358047 | ID_5 |
| 7days | l-shoulder__Movement | -0.8068592 | ID_1 |
| 7days | l-shoulder__Movement | -0.8889002 | ID_2 |
| 7days | l-shoulder__Movement | -0.690238 | ID_3 |
| 7days | l-shoulder__Movement | -0.9319632 | ID_4 |
| 7days | l-shoulder__Movement | -0.8581543 | ID_5 |
| 7days | l-shoulder__Movement | -0.6503535 | ID_6 |
| 7days | l-shoulder__Movement | -0.4490889 | ID_7 |
| 7days | l-shoulder__Movement | -0.4100919 | ID_8 |
| 14days | l-shoulder__Movement | -0.4189732 | ID_1 |
| 14days | l-shoulder__Movement | -0.908218 | ID_2 |
| 14days | l-shoulder__Movement | -0.1772168 | ID_3 |
| 14days | l-shoulder__Movement | 0.10042066 | ID_4 |
| 14days | l-shoulder__Movement | -0.5304521 | ID_5 |

| | | | |
|---|---|---|---|
| 14days | l-shoulder__Movement | -0.3574914 | ID_6 |
| 14days | l-shoulder__Movement | -0.1376612 | ID_7 |
| 14days | l-shoulder__Movement | 0.0846998 | ID_8 |
| 14days | l-shoulder__Movement | 0.50607172 | ID_9 |
| 21days | l-shoulder__Movement | -0.2942903 | ID_1 |
| 21days | l-shoulder__Movement | -0.4887408 | ID_2 |
| 21days | l-shoulder__Movement | -0.6515596 | ID_3 |
| 21days | l-shoulder__Movement | -0.2397791 | ID_4 |
| 21days | l-shoulder__Movement | -0.7298592 | ID_5 |
| 21days | l-shoulder__Movement | -0.7385215 | ID_6 |
| 21days | l-shoulder__Movement | -0.3908298 | ID_7 |
| 21days | l-shoulder__Movement | 0.04704945 | ID_8 |
| 21days | l-shoulder__Movement | -0.5734602 | ID_9 |
| 21days | l-shoulder__Movement | -0.251682 | ID_10 |
| 0days | l-tail-base__Average_Height | -0.3192629 | ID_1 |
| 0days | l-tail-base__Average_Height | 0.06050594 | ID_2 |
| 0days | l-tail-base__Average_Height | -0.2867967 | ID_3 |
| 0days | l-tail-base__Average_Height | 0.70787679 | ID_4 |
| 0days | l-tail-base__Average_Height | -0.3067845 | ID_5 |
| 0days | l-tail-base__Average_Height | 0 | ID_6 |
| 0days | l-tail-base__Average_Height | 0.02259488 | ID_7 |
| 0days | l-tail-base__Average_Height | 0.32524338 | ID_8 |
| 0days | l-tail-base__Average_Height | 0.02259161 | ID_9 |
| 0days | l-tail-base__Average_Height | -0.3950668 | ID_10 |
| 0days | l-tail-base__Average_Height | -0.3294701 | ID_11 |
| 0days | l-tail-base__Average_Height | -0.6811874 | ID_12 |
| 0days | l-tail-base__Average_Height | 0.21650675 | ID_13 |
| 3days | l-tail-base__Average_Height | -0.2851315 | ID_1 |
| 3days | l-tail-base__Average_Height | -0.3042199 | ID_2 |
| 3days | l-tail-base__Average_Height | -0.324553 | ID_3 |
| 3days | l-tail-base__Average_Height | -0.0835919 | ID_4 |
| 3days | l-tail-base__Average_Height | 0.11254996 | ID_5 |
| 3days | l-tail-base__Average_Height | -0.3287968 | ID_6 |
| 3days | l-tail-base__Average_Height | 0.64616202 | ID_7 |
| 3days | l-tail-base__Average_Height | 0.98984473 | ID_8 |
| 7days | l-tail-base__Average_Height | -0.19543 | ID_1 |
| 7days | l-tail-base__Average_Height | -0.2014654 | ID_2 |
| 7days | l-tail-base__Average_Height | -0.6513686 | ID_3 |
| 7days | l-tail-base__Average_Height | 0.08705057 | ID_4 |
| 7days | l-tail-base__Average_Height | -0.5485664 | ID_5 |
| 7days | l-tail-base__Average_Height | -0.1128758 | ID_6 |
| 7days | l-tail-base__Average_Height | 0.05543214 | ID_7 |
| 7days | l-tail-base__Average_Height | -0.4786034 | ID_8 |
| 7days | l-tail-base__Average_Height | 0.25057599 | ID_9 |
| 14days | l-tail-base__Average_Height | 0.60247966 | ID_1 |
| 14days | l-tail-base__Average_Height | -0.5164752 | ID_2 |
| 14days | l-tail-base__Average_Height | 1.14574511 | ID_3 |
| 14days | l-tail-base__Average_Height | -0.339392 | ID_4 |
| 14days | l-tail-base__Average_Height | -0.1848235 | ID_5 |
| 14days | l-tail-base__Average_Height | -0.4406353 | ID_6 |
| 14days | l-tail-base__Average_Height | -0.8135411 | ID_7 |
| 14days | l-tail-base__Average_Height | -0.535126 | ID_8 |
| 14days | l-tail-base__Average_Height | 0.96679538 | ID_9 |
| 14days | l-tail-base__Average_Height | -0.0303554 | ID_10 |
| 14days | l-tail-base__Average_Height | -0.3294469 | ID_11 |
| 21days | l-tail-base__Average_Height | 0.04145865 | ID_1 |

| | | | |
|---|---|---|---|
| 21days | l-tail-base__Average_Height | -0.5190112 | ID_2 |
| 21days | l-tail-base__Average_Height | -0.352422 | ID_3 |
| 21days | l-tail-base__Average_Height | 0.19250263 | ID_4 |
| 21days | l-tail-base__Average_Height | -0.3555155 | ID_5 |
| 21days | l-tail-base__Average_Height | -0.1853749 | ID_6 |
| 21days | l-tail-base__Average_Height | -0.2815834 | ID_7 |
| 21days | l-tail-base__Average_Height | -0.192806 | ID_8 |
| 21days | l-tail-base__Average_Height | -0.6894058 | ID_9 |
| 21days | l-tail-base__Average_Height | -0.1881053 | ID_10 |
| 0days | l-tail-base__Movement | -1.4445186 | ID_1 |
| 0days | l-tail-base__Movement | 0.82770614 | ID_2 |
| 0days | l-tail-base__Movement | -1.3344519 | ID_3 |
| 0days | l-tail-base__Movement | 1.09139114 | ID_4 |
| 0days | l-tail-base__Movement | -1.266019 | ID_5 |
| 0days | l-tail-base__Movement | 0.06382176 | ID_6 |
| 0days | l-tail-base__Movement | 0.12528136 | ID_7 |
| 0days | l-tail-base__Movement | 0 | ID_8 |
| 0days | l-tail-base__Movement | 0.07114391 | ID_9 |
| 0days | l-tail-base__Movement | -1.0333938 | ID_10 |
| 0days | l-tail-base__Movement | -1.6426021 | ID_11 |
| 0days | l-tail-base__Movement | -2.000292 | ID_12 |
| 0days | l-tail-base__Movement | 0.45592032 | ID_13 |
| 3days | l-tail-base__Movement | -1.3518024 | ID_1 |
| 3days | l-tail-base__Movement | -1.026158 | ID_2 |
| 3days | l-tail-base__Movement | -1.0562546 | ID_3 |
| 3days | l-tail-base__Movement | -0.7828295 | ID_4 |
| 3days | l-tail-base__Movement | -0.7770649 | ID_5 |
| 3days | l-tail-base__Movement | -0.9997812 | ID_6 |
| 3days | l-tail-base__Movement | 0.71321414 | ID_7 |
| 3days | l-tail-base__Movement | 1.08829781 | ID_8 |
| 7days | l-tail-base__Movement | -1.437235 | ID_1 |
| 7days | l-tail-base__Movement | 0.20246082 | ID_2 |
| 7days | l-tail-base__Movement | -1.6303672 | ID_3 |
| 7days | l-tail-base__Movement | -0.5314102 | ID_4 |
| 7days | l-tail-base__Movement | -0.9261191 | ID_5 |
| 7days | l-tail-base__Movement | 0.2067443 | ID_6 |
| 7days | l-tail-base__Movement | -0.4474122 | ID_7 |
| 7days | l-tail-base__Movement | -1.3791844 | ID_8 |
| 7days | l-tail-base__Movement | 0.51120546 | ID_9 |
| 14days | l-tail-base__Movement | 0.77962477 | ID_1 |
| 14days | l-tail-base__Movement | -0.9008344 | ID_2 |
| 14days | l-tail-base__Movement | 0.57259416 | ID_3 |
| 14days | l-tail-base__Movement | -1.7896185 | ID_4 |
| 14days | l-tail-base__Movement | -1.5304171 | ID_5 |
| 14days | l-tail-base__Movement | -1.5906238 | ID_6 |
| 14days | l-tail-base__Movement | -2.0533953 | ID_7 |
| 14days | l-tail-base__Movement | -1.7541873 | ID_8 |
| 14days | l-tail-base__Movement | 1.02715145 | ID_9 |
| 14days | l-tail-base__Movement | -0.063508 | ID_10 |
| 14days | l-tail-base__Movement | -0.7408715 | ID_11 |
| 21days | l-tail-base__Movement | -0.5553334 | ID_1 |
| 21days | l-tail-base__Movement | -1.6746836 | ID_2 |
| 21days | l-tail-base__Movement | -1.4230839 | ID_3 |
| 21days | l-tail-base__Movement | -0.7162338 | ID_4 |
| 21days | l-tail-base__Movement | -1.0745174 | ID_5 |
| 21days | l-tail-base__Movement | 0.12446746 | ID_6 |

| | | | |
|---|---|---|---|
| 21days | l-tail-base__Movement | -0.8875311 | ID_7 |
| 21days | l-tail-base__Movement | -1.0737186 | ID_8 |
| 21days | l-tail-base__Movement | -1.550028 | ID_9 |
| 21days | l-tail-base__Movement | -0.6307086 | ID_10 |
| 0days | l-wrist__Average_Height | 0 | ID_1 |
| 0days | l-wrist__Average_Height | -0.9168882 | ID_2 |
| 0days | l-wrist__Average_Height | 0.13156143 | ID_3 |
| 0days | l-wrist__Average_Height | -0.0089616 | ID_4 |
| 0days | l-wrist__Average_Height | -0.8736788 | ID_5 |
| 0days | l-wrist__Average_Height | 0.42761381 | ID_6 |
| 0days | l-wrist__Average_Height | 0.16573231 | ID_7 |
| 0days | l-wrist__Average_Height | -0.5073333 | ID_8 |
| 0days | l-wrist__Average_Height | 0.26751428 | ID_9 |
| 0days | l-wrist__Average_Height | -0.2163956 | ID_10 |
| 0days | l-wrist__Average_Height | -0.7771338 | ID_11 |
| 0days | l-wrist__Average_Height | 0.05496432 | ID_12 |
| 0days | l-wrist__Average_Height | 0.46029332 | ID_13 |
| 3days | l-wrist__Average_Height | -1.2033035 | ID_1 |
| 3days | l-wrist__Average_Height | -0.5027445 | ID_2 |
| 3days | l-wrist__Average_Height | -0.6602885 | ID_3 |
| 3days | l-wrist__Average_Height | -1.2235549 | ID_4 |
| 3days | l-wrist__Average_Height | -1.2686132 | ID_5 |
| 3days | l-wrist__Average_Height | -0.6346261 | ID_6 |
| 3days | l-wrist__Average_Height | -0.6605776 | ID_7 |
| 3days | l-wrist__Average_Height | -0.5429727 | ID_8 |
| 7days | l-wrist__Average_Height | -0.3656664 | ID_1 |
| 7days | l-wrist__Average_Height | -0.7288855 | ID_2 |
| 7days | l-wrist__Average_Height | -0.4997624 | ID_3 |
| 7days | l-wrist__Average_Height | -0.9261974 | ID_4 |
| 7days | l-wrist__Average_Height | -0.4650491 | ID_5 |
| 7days | l-wrist__Average_Height | -0.8274723 | ID_6 |
| 7days | l-wrist__Average_Height | -1.107149 | ID_7 |
| 7days | l-wrist__Average_Height | -0.7732756 | ID_8 |
| 7days | l-wrist__Average_Height | -0.9342893 | ID_9 |
| 14days | l-wrist__Average_Height | -0.3065015 | ID_1 |
| 14days | l-wrist__Average_Height | -0.2901254 | ID_2 |
| 14days | l-wrist__Average_Height | -0.7515281 | ID_3 |
| 14days | l-wrist__Average_Height | -0.8692448 | ID_4 |
| 14days | l-wrist__Average_Height | -0.6732098 | ID_5 |
| 14days | l-wrist__Average_Height | -1.476333 | ID_6 |
| 14days | l-wrist__Average_Height | -0.7906645 | ID_7 |
| 14days | l-wrist__Average_Height | -0.8622238 | ID_8 |
| 14days | l-wrist__Average_Height | -1.4346124 | ID_9 |
| 14days | l-wrist__Average_Height | -1.0778146 | ID_10 |
| 14days | l-wrist__Average_Height | -0.333383 | ID_11 |
| 21days | l-wrist__Average_Height | -0.2890956 | ID_1 |
| 21days | l-wrist__Average_Height | 0.25069776 | ID_2 |
| 21days | l-wrist__Average_Height | -0.0317026 | ID_3 |
| 21days | l-wrist__Average_Height | -0.3461518 | ID_4 |
| 21days | l-wrist__Average_Height | -0.3719616 | ID_5 |
| 21days | l-wrist__Average_Height | -1.134017 | ID_6 |
| 21days | l-wrist__Average_Height | -0.068622 | ID_7 |
| 21days | l-wrist__Average_Height | -0.6158714 | ID_8 |
| 21days | l-wrist__Average_Height | -0.6861222 | ID_9 |
| 21days | l-wrist__Average_Height | -0.2163796 | ID_10 |
| 0days | l-wrist__Movement | -0.0668485 | ID_1 |

| | | | |
|---|---|---|---|
| 0days | l-wrist__Movement | -0.8233552 | ID_2 |
| 0days | l-wrist__Movement | 0.43410383 | ID_3 |
| 0days | l-wrist__Movement | 0.65900356 | ID_4 |
| 0days | l-wrist__Movement | -0.7015359 | ID_5 |
| 0days | l-wrist__Movement | 1.0313634 | ID_6 |
| 0days | l-wrist__Movement | 0 | ID_7 |
| 0days | l-wrist__Movement | -0.7123789 | ID_8 |
| 0days | l-wrist__Movement | 0.36823713 | ID_9 |
| 0days | l-wrist__Movement | -0.3149964 | ID_10 |
| 0days | l-wrist__Movement | -0.2060994 | ID_11 |
| 0days | l-wrist__Movement | 0.29965421 | ID_12 |
| 0days | l-wrist__Movement | 0.32131534 | ID_13 |
| 3days | l-wrist__Movement | -1.5042083 | ID_1 |
| 3days | l-wrist__Movement | -0.5815651 | ID_2 |
| 3days | l-wrist__Movement | -0.5748475 | ID_3 |
| 3days | l-wrist__Movement | -1.5834412 | ID_4 |
| 3days | l-wrist__Movement | -2.1141262 | ID_5 |
| 3days | l-wrist__Movement | -0.5199515 | ID_6 |
| 3days | l-wrist__Movement | -0.7188211 | ID_7 |
| 3days | l-wrist__Movement | -1.0053682 | ID_8 |
| 7days | l-wrist__Movement | -0.040705 | ID_1 |
| 7days | l-wrist__Movement | -1.3582428 | ID_2 |
| 7days | l-wrist__Movement | -0.9271556 | ID_3 |
| 7days | l-wrist__Movement | -1.6903592 | ID_4 |
| 7days | l-wrist__Movement | -0.9693707 | ID_5 |
| 7days | l-wrist__Movement | -0.876172 | ID_6 |
| 7days | l-wrist__Movement | -1.0666631 | ID_7 |
| 7days | l-wrist__Movement | -1.122757 | ID_8 |
| 7days | l-wrist__Movement | -1.4632493 | ID_9 |
| 14days | l-wrist__Movement | -0.026349 | ID_1 |
| 14days | l-wrist__Movement | -0.5308143 | ID_2 |
| 14days | l-wrist__Movement | -1.1437958 | ID_3 |
| 14days | l-wrist__Movement | -1.1734528 | ID_4 |
| 14days | l-wrist__Movement | -0.6209772 | ID_5 |
| 14days | l-wrist__Movement | -1.5635549 | ID_6 |
| 14days | l-wrist__Movement | -0.7988941 | ID_7 |
| 14days | l-wrist__Movement | -0.8620086 | ID_8 |
| 14days | l-wrist__Movement | -0.6140472 | ID_9 |
| 14days | l-wrist__Movement | -0.8267626 | ID_10 |
| 14days | l-wrist__Movement | 0.43595011 | ID_11 |
| 21days | l-wrist__Movement | -0.4633034 | ID_1 |
| 21days | l-wrist__Movement | 1.32532468 | ID_2 |
| 21days | l-wrist__Movement | 0.48366636 | ID_3 |
| 21days | l-wrist__Movement | 0.19644942 | ID_4 |
| 21days | l-wrist__Movement | -0.7893995 | ID_5 |
| 21days | l-wrist__Movement | -1.0992442 | ID_6 |
| 21days | l-wrist__Movement | -0.0645873 | ID_7 |
| 21days | l-wrist__Movement | 0.23430611 | ID_8 |
| 21days | l-wrist__Movement | -0.832874 | ID_9 |
| 21days | l-wrist__Movement | 0.31235869 | ID_10 |
| 0days | r-back-ankle__Average_Heig | -0.1468829 | ID_1 |
| 0days | r-back-ankle__Average_Heig | -0.0401207 | ID_2 |
| 0days | r-back-ankle__Average_Heig | 0.39735087 | ID_3 |
| 0days | r-back-ankle__Average_Heig | 0.37653903 | ID_4 |
| 0days | r-back-ankle__Average_Heig | 0.96955425 | ID_5 |
| 0days | r-back-ankle__Average_Heig | 0 | ID_6 |

| | | | |
|---|---|---|---|
| 0days | r-back-ankle__Average_Heig | -0.0345592 | ID_7 |
| 0days | r-back-ankle__Average_Heig | -0.5484057 | ID_8 |
| 0days | r-back-ankle__Average_Heig | -0.1130645 | ID_9 |
| 0days | r-back-ankle__Average_Heig | -0.1082404 | ID_10 |
| 0days | r-back-ankle__Average_Heig | 0.60831657 | ID_11 |
| 0days | r-back-ankle__Average_Heig | 0.58211529 | ID_12 |
| 0days | r-back-ankle__Average_Heig | 0.56271777 | ID_13 |
| 3days | r-back-ankle__Average_Heig | 0.47113181 | ID_1 |
| 3days | r-back-ankle__Average_Heig | -0.2493458 | ID_2 |
| 3days | r-back-ankle__Average_Heig | -0.1913517 | ID_3 |
| 3days | r-back-ankle__Average_Heig | 0.67381952 | ID_4 |
| 3days | r-back-ankle__Average_Heig | 1.23222732 | ID_5 |
| 3days | r-back-ankle__Average_Heig | 0.0424854 | ID_6 |
| 3days | r-back-ankle__Average_Heig | 0.98055707 | ID_7 |
| 3days | r-back-ankle__Average_Heig | 0.11957431 | ID_8 |
| 7days | r-back-ankle__Average_Heig | 0.07067285 | ID_1 |
| 7days | r-back-ankle__Average_Heig | -0.1828693 | ID_2 |
| 7days | r-back-ankle__Average_Heig | 0.36533421 | ID_3 |
| 7days | r-back-ankle__Average_Heig | 0.46978811 | ID_4 |
| 7days | r-back-ankle__Average_Heig | 0.22741484 | ID_5 |
| 7days | r-back-ankle__Average_Heig | 0.96930615 | ID_6 |
| 7days | r-back-ankle__Average_Heig | 0.87412391 | ID_7 |
| 7days | r-back-ankle__Average_Heig | 0.09158385 | ID_8 |
| 7days | r-back-ankle__Average_Heig | 0.60109886 | ID_9 |
| 14days | r-back-ankle__Average_Heig | -0.1909383 | ID_1 |
| 14days | r-back-ankle__Average_Heig | 0.47264935 | ID_2 |
| 14days | r-back-ankle__Average_Heig | 0.62618015 | ID_3 |
| 14days | r-back-ankle__Average_Heig | 0.70032853 | ID_4 |
| 14days | r-back-ankle__Average_Heig | 0.88070319 | ID_5 |
| 14days | r-back-ankle__Average_Heig | 0.82596298 | ID_6 |
| 14days | r-back-ankle__Average_Heig | 0.96959819 | ID_7 |
| 14days | r-back-ankle__Average_Heig | 0.86704915 | ID_8 |
| 14days | r-back-ankle__Average_Heig | -0.2693921 | ID_9 |
| 14days | r-back-ankle__Average_Heig | 1.01156415 | ID_10 |
| 14days | r-back-ankle__Average_Heig | -0.1367473 | ID_11 |
| 21days | r-back-ankle__Average_Heig | -0.3918919 | ID_1 |
| 21days | r-back-ankle__Average_Heig | 0.53226777 | ID_2 |
| 21days | r-back-ankle__Average_Heig | 0.58103652 | ID_3 |
| 21days | r-back-ankle__Average_Heig | 1.05763863 | ID_4 |
| 21days | r-back-ankle__Average_Heig | 0.31793073 | ID_5 |
| 21days | r-back-ankle__Average_Heig | 0.85203847 | ID_6 |
| 21days | r-back-ankle__Average_Heig | -0.1132625 | ID_7 |
| 21days | r-back-ankle__Average_Heig | 0.22886673 | ID_8 |
| 21days | r-back-ankle__Average_Heig | 0.59331043 | ID_9 |
| 21days | r-back-ankle__Average_Heig | 0.23823543 | ID_10 |
| 0days | r-back-ankle__Movement | -0.53666 | ID_1 |
| 0days | r-back-ankle__Movement | -0.2437109 | ID_2 |
| 0days | r-back-ankle__Movement | 0.12589186 | ID_3 |
| 0days | r-back-ankle__Movement | -0.2556339 | ID_4 |
| 0days | r-back-ankle__Movement | 0.9582799 | ID_5 |
| 0days | r-back-ankle__Movement | 0.72373607 | ID_6 |
| 0days | r-back-ankle__Movement | 0.1745796 | ID_7 |
| 0days | r-back-ankle__Movement | -1.7483562 | ID_8 |
| 0days | r-back-ankle__Movement | -1.2817317 | ID_9 |
| 0days | r-back-ankle__Movement | -0.5547742 | ID_10 |
| 0days | r-back-ankle__Movement | 0 | ID_11 |

| | | | |
|---|---|---|---|
| 0days | r-back-ankle__Movement | 0.78925078 | ID_12 |
| 0days | r-back-ankle__Movement | 0.77578419 | ID_13 |
| 3days | r-back-ankle__Movement | -0.4562464 | ID_1 |
| 3days | r-back-ankle__Movement | -1.2200183 | ID_2 |
| 3days | r-back-ankle__Movement | -1.312673 | ID_3 |
| 3days | r-back-ankle__Movement | -0.2749098 | ID_4 |
| 3days | r-back-ankle__Movement | 1.11911827 | ID_5 |
| 3days | r-back-ankle__Movement | -0.2213618 | ID_6 |
| 3days | r-back-ankle__Movement | 1.81552971 | ID_7 |
| 3days | r-back-ankle__Movement | 0.43921921 | ID_8 |
| 7days | r-back-ankle__Movement | -1.0519767 | ID_1 |
| 7days | r-back-ankle__Movement | -0.5108035 | ID_2 |
| 7days | r-back-ankle__Movement | -0.9018289 | ID_3 |
| 7days | r-back-ankle__Movement | 0.0665612 | ID_4 |
| 7days | r-back-ankle__Movement | -0.6195518 | ID_5 |
| 7days | r-back-ankle__Movement | 2.01364032 | ID_6 |
| 7days | r-back-ankle__Movement | 0.83046006 | ID_7 |
| 7days | r-back-ankle__Movement | 0.05341691 | ID_8 |
| 7days | r-back-ankle__Movement | 0.67465299 | ID_9 |
| 14days | r-back-ankle__Movement | -1.1212053 | ID_1 |
| 14days | r-back-ankle__Movement | 0.29950301 | ID_2 |
| 14days | r-back-ankle__Movement | 0.33130637 | ID_3 |
| 14days | r-back-ankle__Movement | 1.03161695 | ID_4 |
| 14days | r-back-ankle__Movement | 1.0908857 | ID_5 |
| 14days | r-back-ankle__Movement | 0.33265384 | ID_6 |
| 14days | r-back-ankle__Movement | 1.24265452 | ID_7 |
| 14days | r-back-ankle__Movement | 1.28354218 | ID_8 |
| 14days | r-back-ankle__Movement | 0.57465742 | ID_9 |
| 14days | r-back-ankle__Movement | 3.04106747 | ID_10 |
| 14days | r-back-ankle__Movement | -0.2951909 | ID_11 |
| 21days | r-back-ankle__Movement | -1.5107487 | ID_1 |
| 21days | r-back-ankle__Movement | 0.78949516 | ID_2 |
| 21days | r-back-ankle__Movement | 0.81447419 | ID_3 |
| 21days | r-back-ankle__Movement | 1.56851356 | ID_4 |
| 21days | r-back-ankle__Movement | 0.25379168 | ID_5 |
| 21days | r-back-ankle__Movement | 1.47900434 | ID_6 |
| 21days | r-back-ankle__Movement | -1.0127802 | ID_7 |
| 21days | r-back-ankle__Movement | 0.68665494 | ID_8 |
| 21days | r-back-ankle__Movement | 0.91334627 | ID_9 |
| 21days | r-back-ankle__Movement | 0.51075904 | ID_10 |
| 0days | r-back-toe__Average_Height | 0.16428184 | ID_1 |
| 0days | r-back-toe__Average_Height | 0.23238301 | ID_2 |
| 0days | r-back-toe__Average_Height | -0.205038 | ID_3 |
| 0days | r-back-toe__Average_Height | -0.0667881 | ID_4 |
| 0days | r-back-toe__Average_Height | -0.2077944 | ID_5 |
| 0days | r-back-toe__Average_Height | 0 | ID_6 |
| 0days | r-back-toe__Average_Height | 0.09773425 | ID_7 |
| 0days | r-back-toe__Average_Height | 0.10768269 | ID_8 |
| 0days | r-back-toe__Average_Height | 0.02231382 | ID_9 |
| 0days | r-back-toe__Average_Height | 0.07480318 | ID_10 |
| 0days | r-back-toe__Average_Height | -0.2910118 | ID_11 |
| 0days | r-back-toe__Average_Height | -0.0926191 | ID_12 |
| 0days | r-back-toe__Average_Height | -0.0277035 | ID_13 |
| 3days | r-back-toe__Average_Height | -0.1705249 | ID_1 |
| 3days | r-back-toe__Average_Height | -0.2552123 | ID_2 |
| 3days | r-back-toe__Average_Height | -0.0630304 | ID_3 |

| | | | |
|---|---|---|---|
| 3days | r-back-toe__Average_Height | -0.2271536 | ID_4 |
| 3days | r-back-toe__Average_Height | 0.06989828 | ID_5 |
| 3days | r-back-toe__Average_Height | -0.1739801 | ID_6 |
| 3days | r-back-toe__Average_Height | 0.1501916 | ID_7 |
| 3days | r-back-toe__Average_Height | -0.0341578 | ID_8 |
| 7days | r-back-toe__Average_Height | -0.0422763 | ID_1 |
| 7days | r-back-toe__Average_Height | -0.0695279 | ID_2 |
| 7days | r-back-toe__Average_Height | 0.34405031 | ID_3 |
| 7days | r-back-toe__Average_Height | 0.74579805 | ID_4 |
| 7days | r-back-toe__Average_Height | -0.037491 | ID_5 |
| 7days | r-back-toe__Average_Height | 0.41882362 | ID_6 |
| 7days | r-back-toe__Average_Height | -0.0918367 | ID_7 |
| 7days | r-back-toe__Average_Height | -0.0923621 | ID_8 |
| 7days | r-back-toe__Average_Height | -0.2709577 | ID_9 |
| 14days | r-back-toe__Average_Height | -0.2069974 | ID_1 |
| 14days | r-back-toe__Average_Height | -0.132112 | ID_2 |
| 14days | r-back-toe__Average_Height | 0.14321988 | ID_3 |
| 14days | r-back-toe__Average_Height | 0.21929715 | ID_4 |
| 14days | r-back-toe__Average_Height | 0.31739203 | ID_5 |
| 14days | r-back-toe__Average_Height | -0.0223594 | ID_6 |
| 14days | r-back-toe__Average_Height | 0.60258774 | ID_7 |
| 14days | r-back-toe__Average_Height | 0.77690214 | ID_8 |
| 14days | r-back-toe__Average_Height | -0.0537482 | ID_9 |
| 14days | r-back-toe__Average_Height | 0.29723644 | ID_10 |
| 14days | r-back-toe__Average_Height | -0.0556741 | ID_11 |
| 21days | r-back-toe__Average_Height | -0.204243 | ID_1 |
| 21days | r-back-toe__Average_Height | 0.16089253 | ID_2 |
| 21days | r-back-toe__Average_Height | 0.07056671 | ID_3 |
| 21days | r-back-toe__Average_Height | 1.0563191 | ID_4 |
| 21days | r-back-toe__Average_Height | -0.0126671 | ID_5 |
| 21days | r-back-toe__Average_Height | 0.13612435 | ID_6 |
| 21days | r-back-toe__Average_Height | 0.07347411 | ID_7 |
| 21days | r-back-toe__Average_Height | 0.60808882 | ID_8 |
| 21days | r-back-toe__Average_Height | 0.37612957 | ID_9 |
| 21days | r-back-toe__Average_Height | 0.43363305 | ID_10 |
| 0days | r-back-toe__Movement | 0.15088302 | ID_1 |
| 0days | r-back-toe__Movement | 0.84273158 | ID_2 |
| 0days | r-back-toe__Movement | -0.5728224 | ID_3 |
| 0days | r-back-toe__Movement | -0.8089688 | ID_4 |
| 0days | r-back-toe__Movement | -0.2289414 | ID_5 |
| 0days | r-back-toe__Movement | 0.20414094 | ID_6 |
| 0days | r-back-toe__Movement | 0.01306226 | ID_7 |
| 0days | r-back-toe__Movement | 1.49536378 | ID_8 |
| 0days | r-back-toe__Movement | -0.5327173 | ID_9 |
| 0days | r-back-toe__Movement | -0.0958501 | ID_10 |
| 0days | r-back-toe__Movement | -0.0451249 | ID_11 |
| 0days | r-back-toe__Movement | 0.47176219 | ID_12 |
| 0days | r-back-toe__Movement | 0 | ID_13 |
| 3days | r-back-toe__Movement | -0.413253 | ID_1 |
| 3days | r-back-toe__Movement | -0.4418956 | ID_2 |
| 3days | r-back-toe__Movement | -0.4449753 | ID_3 |
| 3days | r-back-toe__Movement | -0.2852193 | ID_4 |
| 3days | r-back-toe__Movement | 0.94427768 | ID_5 |
| 3days | r-back-toe__Movement | -0.1405329 | ID_6 |
| 3days | r-back-toe__Movement | 1.1655564 | ID_7 |
| 3days | r-back-toe__Movement | 0.4619632 | ID_8 |

| | | | |
|---|---|---|---|
| 7days | r-back-toe__Movement | 0.64306093 | ID_1 |
| 7days | r-back-toe__Movement | 0.88709069 | ID_2 |
| 7days | r-back-toe__Movement | 0.57444742 | ID_3 |
| 7days | r-back-toe__Movement | 0.76843434 | ID_4 |
| 7days | r-back-toe__Movement | -0.1996166 | ID_5 |
| 7days | r-back-toe__Movement | 1.23218581 | ID_6 |
| 7days | r-back-toe__Movement | 0.31819638 | ID_7 |
| 7days | r-back-toe__Movement | -0.2934807 | ID_8 |
| 7days | r-back-toe__Movement | -0.1789538 | ID_9 |
| 14days | r-back-toe__Movement | -0.8820356 | ID_1 |
| 14days | r-back-toe__Movement | -0.3334188 | ID_2 |
| 14days | r-back-toe__Movement | 1.73425966 | ID_3 |
| 14days | r-back-toe__Movement | 0.51693385 | ID_4 |
| 14days | r-back-toe__Movement | 0.87075354 | ID_5 |
| 14days | r-back-toe__Movement | -0.1281288 | ID_6 |
| 14days | r-back-toe__Movement | 1.32247091 | ID_7 |
| 14days | r-back-toe__Movement | 1.41374384 | ID_8 |
| 14days | r-back-toe__Movement | 0.04532501 | ID_9 |
| 14days | r-back-toe__Movement | 1.00426211 | ID_10 |
| 14days | r-back-toe__Movement | -0.5592575 | ID_11 |
| 21days | r-back-toe__Movement | -0.1767319 | ID_1 |
| 21days | r-back-toe__Movement | 0.55448603 | ID_2 |
| 21days | r-back-toe__Movement | 0.33671286 | ID_3 |
| 21days | r-back-toe__Movement | 3.09688759 | ID_4 |
| 21days | r-back-toe__Movement | 0.07650652 | ID_5 |
| 21days | r-back-toe__Movement | 0.44406029 | ID_6 |
| 21days | r-back-toe__Movement | -0.0751893 | ID_7 |
| 21days | r-back-toe__Movement | 0.99516644 | ID_8 |
| 21days | r-back-toe__Movement | 0.6239984 | ID_9 |
| 21days | r-back-toe__Movement | 1.74329356 | ID_10 |
| 0days | r-elbow__Average_Height | 0.00362726 | ID_1 |
| 0days | r-elbow__Average_Height | -0.4731648 | ID_2 |
| 0days | r-elbow__Average_Height | 0 | ID_3 |
| 0days | r-elbow__Average_Height | 0.59821548 | ID_4 |
| 0days | r-elbow__Average_Height | -0.1795274 | ID_5 |
| 0days | r-elbow__Average_Height | -0.2091248 | ID_6 |
| 0days | r-elbow__Average_Height | 0.06555415 | ID_7 |
| 0days | r-elbow__Average_Height | 0.0028623 | ID_8 |
| 0days | r-elbow__Average_Height | 0.00705776 | ID_9 |
| 0days | r-elbow__Average_Height | -0.3061086 | ID_10 |
| 0days | r-elbow__Average_Height | -0.3496225 | ID_11 |
| 0days | r-elbow__Average_Height | -0.2138599 | ID_12 |
| 0days | r-elbow__Average_Height | 0.15087725 | ID_13 |
| 3days | r-elbow__Average_Height | -0.4602182 | ID_1 |
| 3days | r-elbow__Average_Height | -0.44453 | ID_2 |
| 3days | r-elbow__Average_Height | -0.4661707 | ID_3 |
| 3days | r-elbow__Average_Height | -0.5317932 | ID_4 |
| 3days | r-elbow__Average_Height | -0.6897216 | ID_5 |
| 3days | r-elbow__Average_Height | -0.4917161 | ID_6 |
| 3days | r-elbow__Average_Height | 0.08653793 | ID_7 |
| 3days | r-elbow__Average_Height | -0.0810403 | ID_8 |
| 7days | r-elbow__Average_Height | -0.5885457 | ID_1 |
| 7days | r-elbow__Average_Height | -0.3345832 | ID_2 |
| 7days | r-elbow__Average_Height | -0.38448 | ID_3 |
| 7days | r-elbow__Average_Height | -0.0576795 | ID_4 |
| 7days | r-elbow__Average_Height | -0.0200633 | ID_5 |

| | | | |
|---|---|---|---|
| 7days | r-elbow__Average_Height | -0.3382561 | ID_6 |
| 7days | r-elbow__Average_Height | -0.1930686 | ID_7 |
| 7days | r-elbow__Average_Height | -0.1289549 | ID_8 |
| 7days | r-elbow__Average_Height | -0.269897 | ID_9 |
| 14days | r-elbow__Average_Height | -0.2779463 | ID_1 |
| 14days | r-elbow__Average_Height | 0.04722786 | ID_2 |
| 14days | r-elbow__Average_Height | -0.4490856 | ID_3 |
| 14days | r-elbow__Average_Height | -0.1542237 | ID_4 |
| 14days | r-elbow__Average_Height | -0.3414627 | ID_5 |
| 14days | r-elbow__Average_Height | -0.3675006 | ID_6 |
| 14days | r-elbow__Average_Height | -0.7401342 | ID_7 |
| 14days | r-elbow__Average_Height | -0.168529 | ID_8 |
| 14days | r-elbow__Average_Height | -0.272153 | ID_9 |
| 14days | r-elbow__Average_Height | -0.2597085 | ID_10 |
| 21days | r-elbow__Average_Height | -0.3408939 | ID_11 |
| 21days | r-elbow__Average_Height | 0.31149782 | ID_1 |
| 21days | r-elbow__Average_Height | -0.2743563 | ID_2 |
| 21days | r-elbow__Average_Height | -0.252446 | ID_3 |
| 21days | r-elbow__Average_Height | 0.06826335 | ID_4 |
| 21days | r-elbow__Average_Height | -0.5913508 | ID_5 |
| 21days | r-elbow__Average_Height | -0.0757413 | ID_6 |
| 21days | r-elbow__Average_Height | 0.08629657 | ID_7 |
| 21days | r-elbow__Average_Height | -0.0771913 | ID_8 |
| 21days | r-elbow__Average_Height | 0.63612734 | ID_9 |
| 0days | r-elbow__Movement | -0.1548557 | ID_10 |
| 0days | r-elbow__Movement | -0.9641164 | ID_1 |
| 0days | r-elbow__Movement | 0.02450275 | ID_2 |
| 0days | r-elbow__Movement | 0.57259644 | ID_3 |
| 0days | r-elbow__Movement | -0.1796794 | ID_4 |
| 0days | r-elbow__Movement | -0.7718495 | ID_5 |
| 0days | r-elbow__Movement | 0.36785776 | ID_6 |
| 0days | r-elbow__Movement | 0 | ID_7 |
| 0days | r-elbow__Movement | 0.37572922 | ID_8 |
| 0days | r-elbow__Movement | -1.1822798 | ID_9 |
| 0days | r-elbow__Movement | -0.8096325 | ID_10 |
| 0days | r-elbow__Movement | 0.11151942 | ID_11 |
| 0days | r-elbow__Movement | 0.46812633 | ID_12 |
| 3days | r-elbow__Movement | -0.9704 | ID_13 |
| 3days | r-elbow__Movement | -1.2574547 | ID_1 |
| 3days | r-elbow__Movement | -0.8054357 | ID_2 |
| 3days | r-elbow__Movement | -1.0453943 | ID_3 |
| 3days | r-elbow__Movement | -1.4619121 | ID_4 |
| 3days | r-elbow__Movement | -0.9935279 | ID_5 |
| 3days | r-elbow__Movement | -0.1506226 | ID_6 |
| 3days | r-elbow__Movement | 0.06173137 | ID_7 |
| 7days | r-elbow__Movement | -1.4591558 | ID_8 |
| 7days | r-elbow__Movement | -0.7326618 | ID_1 |
| 7days | r-elbow__Movement | -1.0562758 | ID_2 |
| 7days | r-elbow__Movement | -0.9081111 | ID_3 |
| 7days | r-elbow__Movement | -0.7497038 | ID_4 |
| 7days | r-elbow__Movement | -0.9367159 | ID_5 |
| 7days | r-elbow__Movement | -0.8367696 | ID_6 |
| 7days | r-elbow__Movement | -0.9526068 | ID_7 |
| 7days | r-elbow__Movement | -0.9394047 | ID_8 |
| 14days | r-elbow__Movement | -0.6687055 | ID_9 |
| 14days | r-elbow__Movement | -0.454514 | ID_1 |

| | | | |
|---|---|---|---|
| 14days | r-elbow__Movement | -1.1927558 | ID_2 |
| 14days | r-elbow__Movement | -0.4436888 | ID_3 |
| 14days | r-elbow__Movement | -0.957033 | ID_4 |
| 14days | r-elbow__Movement | -0.7357572 | ID_5 |
| 14days | r-elbow__Movement | -1.4834283 | ID_6 |
| 14days | r-elbow__Movement | 0.23674366 | ID_7 |
| 14days | r-elbow__Movement | -0.7070423 | ID_8 |
| 14days | r-elbow__Movement | -0.4223416 | ID_9 |
| 21days | r-elbow__Movement | -0.8425795 | ID_10 |
| 21days | r-elbow__Movement | 0.29334309 | ID_11 |
| 21days | r-elbow__Movement | -0.3426718 | ID_1 |
| 21days | r-elbow__Movement | -0.4123859 | ID_2 |
| 21days | r-elbow__Movement | -0.5608952 | ID_3 |
| 21days | r-elbow__Movement | -1.4820006 | ID_4 |
| 21days | r-elbow__Movement | -0.465817 | ID_5 |
| 21days | r-elbow__Movement | -0.3569001 | ID_6 |
| 21days | r-elbow__Movement | -0.7083302 | ID_7 |
| 21days | r-elbow__Movement | 1.64210311 | ID_8 |
| 0days | r-front-toe-tip__Average_Hei | 0.03135791 | ID_1 |
| 0days | r-front-toe-tip__Average_Hei | 0.18709357 | ID_2 |
| 0days | r-front-toe-tip__Average_Hei | 0.1476945 | ID_3 |
| 0days | r-front-toe-tip__Average_Hei | -0.001675 | ID_4 |
| 0days | r-front-toe-tip__Average_Hei | -0.1703256 | ID_5 |
| 0days | r-front-toe-tip__Average_Hei | 0.00514434 | ID_6 |
| 0days | r-front-toe-tip__Average_Hei | 0 | ID_7 |
| 0days | r-front-toe-tip__Average_Hei | 0.57169011 | ID_8 |
| 0days | r-front-toe-tip__Average_Hei | -0.0052955 | ID_9 |
| 0days | r-front-toe-tip__Average_Hei | -0.2480867 | ID_10 |
| 0days | r-front-toe-tip__Average_Hei | -0.1338386 | ID_11 |
| 0days | r-front-toe-tip__Average_Hei | -0.1716836 | ID_12 |
| 0days | r-front-toe-tip__Average_Hei | 0.20355628 | ID_13 |
| 3days | r-front-toe-tip__Average_Hei | -0.2158472 | ID_1 |
| 3days | r-front-toe-tip__Average_Hei | -0.3191591 | ID_2 |
| 3days | r-front-toe-tip__Average_Hei | -0.0324925 | ID_3 |
| 3days | r-front-toe-tip__Average_Hei | -0.2357843 | ID_4 |
| 3days | r-front-toe-tip__Average_Hei | -0.1801322 | ID_5 |
| 3days | r-front-toe-tip__Average_Hei | -0.3621174 | ID_6 |
| 3days | r-front-toe-tip__Average_Hei | -0.3246147 | ID_7 |
| 3days | r-front-toe-tip__Average_Hei | 0.12807963 | ID_8 |
| 7days | r-front-toe-tip__Average_Hei | -0.0384315 | ID_1 |
| 7days | r-front-toe-tip__Average_Hei | 0.40600335 | ID_2 |
| 7days | r-front-toe-tip__Average_Hei | -0.3368922 | ID_3 |
| 7days | r-front-toe-tip__Average_Hei | 0.25196417 | ID_4 |
| 7days | r-front-toe-tip__Average_Hei | -0.3000916 | ID_5 |
| 7days | r-front-toe-tip__Average_Hei | 0.27875009 | ID_6 |
| 7days | r-front-toe-tip__Average_Hei | -0.2231241 | ID_7 |
| 7days | r-front-toe-tip__Average_Hei | -0.3058005 | ID_8 |
| 7days | r-front-toe-tip__Average_Hei | -0.1284495 | ID_9 |
| 14days | r-front-toe-tip__Average_Hei | 0.17720183 | ID_1 |
| 14days | r-front-toe-tip__Average_Hei | -0.4238257 | ID_2 |
| 14days | r-front-toe-tip__Average_Hei | 0.03896691 | ID_3 |
| 14days | r-front-toe-tip__Average_Hei | 0.18950152 | ID_4 |
| 14days | r-front-toe-tip__Average_Hei | 0.17184373 | ID_5 |
| 14days | r-front-toe-tip__Average_Hei | -0.4353274 | ID_6 |
| 14days | r-front-toe-tip__Average_Hei | 0.1754423 | ID_7 |
| 14days | r-front-toe-tip__Average_Hei | 0.2353217 | ID_8 |

| | | | |
|---|---|---|---|
| 14days | r-front-toe-tip__Average_Hei | -0.3142367 | ID_9 |
| 14days | r-front-toe-tip__Average_Hei | -0.0381014 | ID_10 |
| 14days | r-front-toe-tip__Average_Hei | -0.1564677 | ID_11 |
| 21days | r-front-toe-tip__Average_Hei | -0.2037808 | ID_1 |
| 21days | r-front-toe-tip__Average_Hei | -0.3479303 | ID_2 |
| 21days | r-front-toe-tip__Average_Hei | -0.3511617 | ID_3 |
| 21days | r-front-toe-tip__Average_Hei | 0.43123269 | ID_4 |
| 21days | r-front-toe-tip__Average_Hei | -0.5066782 | ID_5 |
| 21days | r-front-toe-tip__Average_Hei | 0.24663604 | ID_6 |
| 21days | r-front-toe-tip__Average_Hei | -0.1644824 | ID_7 |
| 21days | r-front-toe-tip__Average_Hei | 0.27898279 | ID_8 |
| 21days | r-front-toe-tip__Average_Hei | 0.19914095 | ID_9 |
| 21days | r-front-toe-tip__Average_Hei | 0.12763178 | ID_10 |
| 0days | r-front-toe-tip__Movement | 0.38925413 | ID_1 |
| 0days | r-front-toe-tip__Movement | 0 | ID_2 |
| 0days | r-front-toe-tip__Movement | -0.012741 | ID_3 |
| 0days | r-front-toe-tip__Movement | -0.0011338 | ID_4 |
| 0days | r-front-toe-tip__Movement | 0.09993967 | ID_5 |
| 0days | r-front-toe-tip__Movement | 0.29299115 | ID_6 |
| 0days | r-front-toe-tip__Movement | -0.0045303 | ID_7 |
| 0days | r-front-toe-tip__Movement | 1.39466025 | ID_8 |
| 0days | r-front-toe-tip__Movement | -0.5692494 | ID_9 |
| 0days | r-front-toe-tip__Movement | -0.5764126 | ID_10 |
| 0days | r-front-toe-tip__Movement | 0.27938092 | ID_11 |
| 0days | r-front-toe-tip__Movement | -0.4648929 | ID_12 |
| 0days | r-front-toe-tip__Movement | 0.25553153 | ID_13 |
| 3days | r-front-toe-tip__Movement | -0.4652572 | ID_1 |
| 3days | r-front-toe-tip__Movement | 0.01358452 | ID_2 |
| 3days | r-front-toe-tip__Movement | -0.3115956 | ID_3 |
| 3days | r-front-toe-tip__Movement | -0.1114153 | ID_4 |
| 3days | r-front-toe-tip__Movement | -0.0541393 | ID_5 |
| 3days | r-front-toe-tip__Movement | -0.1504963 | ID_6 |
| 3days | r-front-toe-tip__Movement | -0.5226726 | ID_7 |
| 3days | r-front-toe-tip__Movement | 0.10113918 | ID_8 |
| 7days | r-front-toe-tip__Movement | 0.29063786 | ID_1 |
| 7days | r-front-toe-tip__Movement | 0.83210408 | ID_2 |
| 7days | r-front-toe-tip__Movement | -0.5529492 | ID_3 |
| 7days | r-front-toe-tip__Movement | 0.46573136 | ID_4 |
| 7days | r-front-toe-tip__Movement | -0.4383184 | ID_5 |
| 7days | r-front-toe-tip__Movement | 0.58447269 | ID_6 |
| 7days | r-front-toe-tip__Movement | -0.727144 | ID_7 |
| 7days | r-front-toe-tip__Movement | -0.1118996 | ID_8 |
| 7days | r-front-toe-tip__Movement | -0.2620417 | ID_9 |
| 14days | r-front-toe-tip__Movement | 0.4401529 | ID_1 |
| 14days | r-front-toe-tip__Movement | -0.146346 | ID_2 |
| 14days | r-front-toe-tip__Movement | -0.0910604 | ID_3 |
| 14days | r-front-toe-tip__Movement | 0.62465954 | ID_4 |
| 14days | r-front-toe-tip__Movement | 0.27596688 | ID_5 |
| 14days | r-front-toe-tip__Movement | -1.1249305 | ID_6 |
| 14days | r-front-toe-tip__Movement | 0.6489982 | ID_7 |
| 14days | r-front-toe-tip__Movement | 0.56490184 | ID_8 |
| 14days | r-front-toe-tip__Movement | -0.6545676 | ID_9 |
| 14days | r-front-toe-tip__Movement | 0.19486483 | ID_10 |
| 14days | r-front-toe-tip__Movement | -0.629778 | ID_11 |
| 21days | r-front-toe-tip__Movement | -0.065203 | ID_1 |
| 21days | r-front-toe-tip__Movement | -0.3671034 | ID_2 |

| | | | |
|---|---|---|---|
| 21days | r-front-toe-tip__Movement | -0.3778014 | ID_3 |
| 21days | r-front-toe-tip__Movement | 0.93774655 | ID_4 |
| 21days | r-front-toe-tip__Movement | -1.1437734 | ID_5 |
| 21days | r-front-toe-tip__Movement | 0.38580527 | ID_6 |
| 21days | r-front-toe-tip__Movement | -0.2891268 | ID_7 |
| 21days | r-front-toe-tip__Movement | 0.15692873 | ID_8 |
| 21days | r-front-toe-tip__Movement | 0.07725664 | ID_9 |
| 21days | r-front-toe-tip__Movement | 0.29202303 | ID_10 |
| 0days | r-hip__Average_Height | -0.4497609 | ID_1 |
| 0days | r-hip__Average_Height | -0.059288 | ID_2 |
| 0days | r-hip__Average_Height | 0 | ID_3 |
| 0days | r-hip__Average_Height | 0.46250577 | ID_4 |
| 0days | r-hip__Average_Height | -0.2079616 | ID_5 |
| 0days | r-hip__Average_Height | 0.01601584 | ID_6 |
| 0days | r-hip__Average_Height | 0.0162802 | ID_7 |
| 0days | r-hip__Average_Height | 0.19679759 | ID_8 |
| 0days | r-hip__Average_Height | -0.3757805 | ID_9 |
| 0days | r-hip__Average_Height | 0.02052768 | ID_10 |
| 0days | r-hip__Average_Height | -0.1778465 | ID_11 |
| 0days | r-hip__Average_Height | 0.40169812 | ID_12 |
| 0days | r-hip__Average_Height | -0.3324927 | ID_13 |
| 3days | r-hip__Average_Height | -0.4157003 | ID_1 |
| 3days | r-hip__Average_Height | -0.4197327 | ID_2 |
| 3days | r-hip__Average_Height | -0.2307823 | ID_3 |
| 3days | r-hip__Average_Height | -0.4071027 | ID_4 |
| 3days | r-hip__Average_Height | -0.2016535 | ID_5 |
| 3days | r-hip__Average_Height | -0.4136191 | ID_6 |
| 3days | r-hip__Average_Height | -0.0399806 | ID_7 |
| 3days | r-hip__Average_Height | 0.3522125 | ID_8 |
| 7days | r-hip__Average_Height | 0.01960609 | ID_1 |
| 7days | r-hip__Average_Height | -0.2428318 | ID_2 |
| 7days | r-hip__Average_Height | -0.0476098 | ID_3 |
| 7days | r-hip__Average_Height | 0.14324227 | ID_4 |
| 7days | r-hip__Average_Height | -0.0017086 | ID_5 |
| 7days | r-hip__Average_Height | -0.1109276 | ID_6 |
| 7days | r-hip__Average_Height | -0.411578 | ID_7 |
| 7days | r-hip__Average_Height | -0.190413 | ID_8 |
| 7days | r-hip__Average_Height | 0.22695951 | ID_9 |
| 14days | r-hip__Average_Height | 0.2049254 | ID_1 |
| 14days | r-hip__Average_Height | -0.0463459 | ID_2 |
| 14days | r-hip__Average_Height | -0.1501316 | ID_3 |
| 14days | r-hip__Average_Height | -0.1642908 | ID_4 |
| 14days | r-hip__Average_Height | 0.13371959 | ID_5 |
| 14days | r-hip__Average_Height | 0.217187 | ID_6 |
| 14days | r-hip__Average_Height | -0.5425727 | ID_7 |
| 14days | r-hip__Average_Height | -0.522632 | ID_8 |
| 14days | r-hip__Average_Height | 0.22694619 | ID_9 |
| 14days | r-hip__Average_Height | 0.00571137 | ID_10 |
| 14days | r-hip__Average_Height | -0.0915817 | ID_11 |
| 21days | r-hip__Average_Height | -0.0990266 | ID_1 |
| 21days | r-hip__Average_Height | -0.5339945 | ID_2 |
| 21days | r-hip__Average_Height | -0.4061764 | ID_3 |
| 21days | r-hip__Average_Height | 0.40818297 | ID_4 |
| 21days | r-hip__Average_Height | -0.1786207 | ID_5 |
| 21days | r-hip__Average_Height | -0.1757608 | ID_6 |
| 21days | r-hip__Average_Height | 0.24357971 | ID_7 |

| | | | |
|---|---|---|---|
| 21days | r-hip__Average_Height | 0.59612176 | ID_8 |
| 21days | r-hip__Average_Height | -0.0116845 | ID_9 |
| 21days | r-hip__Average_Height | -0.112731 | ID_10 |
| 0days | r-hip__Movement | -0.5543281 | ID_1 |
| 0days | r-hip__Movement | 0.97697186 | ID_2 |
| 0days | r-hip__Movement | 0.46183821 | ID_3 |
| 0days | r-hip__Movement | 1.71775673 | ID_4 |
| 0days | r-hip__Movement | -0.4879813 | ID_5 |
| 0days | r-hip__Movement | 0 | ID_6 |
| 0days | r-hip__Movement | -0.3338742 | ID_7 |
| 0days | r-hip__Movement | 1.13570277 | ID_8 |
| 0days | r-hip__Movement | -0.8249045 | ID_9 |
| 0days | r-hip__Movement | -0.1916942 | ID_10 |
| 0days | r-hip__Movement | -0.0039839 | ID_11 |
| 0days | r-hip__Movement | 0.26954998 | ID_12 |
| 0days | r-hip__Movement | 0.00253249 | ID_13 |
| 3days | r-hip__Movement | -0.8315936 | ID_1 |
| 3days | r-hip__Movement | -0.599086 | ID_2 |
| 3days | r-hip__Movement | -0.317279 | ID_3 |
| 3days | r-hip__Movement | -0.7174234 | ID_4 |
| 3days | r-hip__Movement | -0.5445019 | ID_5 |
| 3days | r-hip__Movement | -0.630397 | ID_6 |
| 3days | r-hip__Movement | -0.0882008 | ID_7 |
| 3days | r-hip__Movement | 1.62112633 | ID_8 |
| 7days | r-hip__Movement | 0.13201038 | ID_1 |
| 7days | r-hip__Movement | -0.076238 | ID_2 |
| 7days | r-hip__Movement | -0.5621799 | ID_3 |
| 7days | r-hip__Movement | 0.44731972 | ID_4 |
| 7days | r-hip__Movement | -0.2004937 | ID_5 |
| 7days | r-hip__Movement | 0.38042612 | ID_6 |
| 7days | r-hip__Movement | -0.6785643 | ID_7 |
| 7days | r-hip__Movement | -0.0853527 | ID_8 |
| 7days | r-hip__Movement | 0.52051755 | ID_9 |
| 14days | r-hip__Movement | 0.11939104 | ID_1 |
| 14days | r-hip__Movement | -0.0290794 | ID_2 |
| 14days | r-hip__Movement | 0.71295079 | ID_3 |
| 14days | r-hip__Movement | -0.3349939 | ID_4 |
| 14days | r-hip__Movement | 0.37541754 | ID_5 |
| 14days | r-hip__Movement | 0.08961341 | ID_6 |
| 14days | r-hip__Movement | -0.9189982 | ID_7 |
| 14days | r-hip__Movement | -0.7602581 | ID_8 |
| 14days | r-hip__Movement | 0.17726289 | ID_9 |
| 14days | r-hip__Movement | 0.82457658 | ID_10 |
| 14days | r-hip__Movement | -0.5056043 | ID_11 |
| 21days | r-hip__Movement | -0.0628528 | ID_1 |
| 21days | r-hip__Movement | -1.0330445 | ID_2 |
| 21days | r-hip__Movement | -0.2517236 | ID_3 |
| 21days | r-hip__Movement | 1.2426687 | ID_4 |
| 21days | r-hip__Movement | -0.6463288 | ID_5 |
| 21days | r-hip__Movement | 0.46505367 | ID_6 |
| 21days | r-hip__Movement | 0.26521156 | ID_7 |
| 21days | r-hip__Movement | 1.03866696 | ID_8 |
| 21days | r-hip__Movement | -0.1559217 | ID_9 |
| 21days | r-hip__Movement | -0.2562772 | ID_10 |
| 0days | r-iliac-crest__Average_Heigh | -0.2505019 | ID_1 |
| 0days | r-iliac-crest__Average_Heigh | 0.31183046 | ID_2 |

| | | | |
|---|---|---|---|
| 0days | r-iliac-crest__Average_Heigh | 0.16535694 | ID_3 |
| 0days | r-iliac-crest__Average_Heigh | 0.504612 | ID_4 |
| 0days | r-iliac-crest__Average_Heigh | 0 | ID_5 |
| 0days | r-iliac-crest__Average_Heigh | 0.22236175 | ID_6 |
| 0days | r-iliac-crest__Average_Heigh | -0.0271328 | ID_7 |
| 0days | r-iliac-crest__Average_Heigh | 0.44566443 | ID_8 |
| 0days | r-iliac-crest__Average_Heigh | -0.1167121 | ID_9 |
| 0days | r-iliac-crest__Average_Heigh | 0.12722814 | ID_10 |
| 0days | r-iliac-crest__Average_Heigh | -0.1694924 | ID_11 |
| 0days | r-iliac-crest__Average_Heigh | -0.0818991 | ID_12 |
| 0days | r-iliac-crest__Average_Heigh | -0.1263372 | ID_13 |
| 3days | r-iliac-crest__Average_Heigh | -0.2488331 | ID_1 |
| 3days | r-iliac-crest__Average_Heigh | -0.0575799 | ID_2 |
| 3days | r-iliac-crest__Average_Heigh | -0.1217178 | ID_3 |
| 3days | r-iliac-crest__Average_Heigh | -0.1231048 | ID_4 |
| 3days | r-iliac-crest__Average_Heigh | -0.028291 | ID_5 |
| 3days | r-iliac-crest__Average_Heigh | -0.1996952 | ID_6 |
| 3days | r-iliac-crest__Average_Heigh | 0.20389234 | ID_7 |
| 3days | r-iliac-crest__Average_Heigh | 0.29356764 | ID_8 |
| 7days | r-iliac-crest__Average_Heigh | -0.0558289 | ID_1 |
| 7days | r-iliac-crest__Average_Heigh | 0.11319651 | ID_2 |
| 7days | r-iliac-crest__Average_Heigh | 0.03684093 | ID_3 |
| 7days | r-iliac-crest__Average_Heigh | 0.30489547 | ID_4 |
| 7days | r-iliac-crest__Average_Heigh | 0.03775729 | ID_5 |
| 7days | r-iliac-crest__Average_Heigh | 0.52473411 | ID_6 |
| 7days | r-iliac-crest__Average_Heigh | 0.69938113 | ID_7 |
| 7days | r-iliac-crest__Average_Heigh | -0.2851213 | ID_8 |
| 7days | r-iliac-crest__Average_Heigh | -0.3502932 | ID_9 |
| 14days | r-iliac-crest__Average_Heigh | 0.23703158 | ID_1 |
| 14days | r-iliac-crest__Average_Heigh | 0.13260314 | ID_2 |
| 14days | r-iliac-crest__Average_Heigh | 0.40341246 | ID_3 |
| 14days | r-iliac-crest__Average_Heigh | 0.12627804 | ID_4 |
| 14days | r-iliac-crest__Average_Heigh | 0.6876098 | ID_5 |
| 14days | r-iliac-crest__Average_Heigh | -0.011419 | ID_6 |
| 14days | r-iliac-crest__Average_Heigh | -0.4675634 | ID_7 |
| 14days | r-iliac-crest__Average_Heigh | -0.4001553 | ID_8 |
| 14days | r-iliac-crest__Average_Heigh | 0.39091757 | ID_9 |
| 14days | r-iliac-crest__Average_Heigh | 0.18609424 | ID_10 |
| 14days | r-iliac-crest__Average_Heigh | 0.04278494 | ID_11 |
| 21days | r-iliac-crest__Average_Heigh | -0.2106064 | ID_1 |
| 21days | r-iliac-crest__Average_Heigh | -0.2841178 | ID_2 |
| 21days | r-iliac-crest__Average_Heigh | -0.3105821 | ID_3 |
| 21days | r-iliac-crest__Average_Heigh | 0.76170517 | ID_4 |
| 21days | r-iliac-crest__Average_Heigh | -0.2154079 | ID_5 |
| 21days | r-iliac-crest__Average_Heigh | 0.39869757 | ID_6 |
| 21days | r-iliac-crest__Average_Heigh | -0.1317501 | ID_7 |
| 21days | r-iliac-crest__Average_Heigh | 0.4514136 | ID_8 |
| 21days | r-iliac-crest__Average_Heigh | 0.43861571 | ID_9 |
| 21days | r-iliac-crest__Average_Heigh | 0.15495561 | ID_10 |
| 0days | r-iliac-crest__Movement | -0.5478296 | ID_1 |
| 0days | r-iliac-crest__Movement | 1.2688499 | ID_2 |
| 0days | r-iliac-crest__Movement | 0.37669238 | ID_3 |
| 0days | r-iliac-crest__Movement | 1.17802958 | ID_4 |
| 0days | r-iliac-crest__Movement | -0.0773928 | ID_5 |
| 0days | r-iliac-crest__Movement | 0.00950445 | ID_6 |
| 0days | r-iliac-crest__Movement | -0.0847591 | ID_7 |

| | | | |
|---|---|---|---|
| 0days | r-iliac-crest__Movement | 1.91939243 | ID_8 |
| 0days | r-iliac-crest__Movement | -0.2848737 | ID_9 |
| 0days | r-iliac-crest__Movement | 0 | ID_10 |
| 0days | r-iliac-crest__Movement | 0.69444858 | ID_11 |
| 0days | r-iliac-crest__Movement | -0.2142627 | ID_12 |
| 0days | r-iliac-crest__Movement | -0.6708985 | ID_13 |
| 3days | r-iliac-crest__Movement | -0.67236 | ID_1 |
| 3days | r-iliac-crest__Movement | -0.3112036 | ID_2 |
| 3days | r-iliac-crest__Movement | -0.5177308 | ID_3 |
| 3days | r-iliac-crest__Movement | -0.3426203 | ID_4 |
| 3days | r-iliac-crest__Movement | -0.4293133 | ID_5 |
| 3days | r-iliac-crest__Movement | -0.3657583 | ID_6 |
| 3days | r-iliac-crest__Movement | 0.46783034 | ID_7 |
| 3days | r-iliac-crest__Movement | 0.92146101 | ID_8 |
| 7days | r-iliac-crest__Movement | -0.6382622 | ID_1 |
| 7days | r-iliac-crest__Movement | 0.51034028 | ID_2 |
| 7days | r-iliac-crest__Movement | -0.1402113 | ID_3 |
| 7days | r-iliac-crest__Movement | 0.95080326 | ID_4 |
| 7days | r-iliac-crest__Movement | -0.1795549 | ID_5 |
| 7days | r-iliac-crest__Movement | 1.4135252 | ID_6 |
| 7days | r-iliac-crest__Movement | 1.12129972 | ID_7 |
| 7days | r-iliac-crest__Movement | -0.4370683 | ID_8 |
| 7days | r-iliac-crest__Movement | -0.4894146 | ID_9 |
| 14days | r-iliac-crest__Movement | 0.36710025 | ID_1 |
| 14days | r-iliac-crest__Movement | 0.21510089 | ID_2 |
| 14days | r-iliac-crest__Movement | 0.79584784 | ID_3 |
| 14days | r-iliac-crest__Movement | 0.39869722 | ID_4 |
| 14days | r-iliac-crest__Movement | 1.67559451 | ID_5 |
| 14days | r-iliac-crest__Movement | 0.06223912 | ID_6 |
| 14days | r-iliac-crest__Movement | -0.6342515 | ID_7 |
| 14days | r-iliac-crest__Movement | -0.4390243 | ID_8 |
| 14days | r-iliac-crest__Movement | 0.50996829 | ID_9 |
| 14days | r-iliac-crest__Movement | 1.15494125 | ID_10 |
| 14days | r-iliac-crest__Movement | 0.56894137 | ID_11 |
| 21days | r-iliac-crest__Movement | -0.6107009 | ID_1 |
| 21days | r-iliac-crest__Movement | -0.1710312 | ID_2 |
| 21days | r-iliac-crest__Movement | -0.4965877 | ID_3 |
| 21days | r-iliac-crest__Movement | 1.31765594 | ID_4 |
| 21days | r-iliac-crest__Movement | -0.4556228 | ID_5 |
| 21days | r-iliac-crest__Movement | 1.33111391 | ID_6 |
| 21days | r-iliac-crest__Movement | -0.4287663 | ID_7 |
| 21days | r-iliac-crest__Movement | 0.95618634 | ID_8 |
| 21days | r-iliac-crest__Movement | 0.77571354 | ID_9 |
| 21days | r-iliac-crest__Movement | -0.0321817 | ID_10 |
| 0days | r-shoulder__Average_Height | -0.0410086 | ID_1 |
| 0days | r-shoulder__Average_Height | -0.1994091 | ID_2 |
| 0days | r-shoulder__Average_Height | 0.00507822 | ID_3 |
| 0days | r-shoulder__Average_Height | 0.65070218 | ID_4 |
| 0days | r-shoulder__Average_Height | 0 | ID_5 |
| 0days | r-shoulder__Average_Height | 0.03556996 | ID_6 |
| 0days | r-shoulder__Average_Height | -0.161415 | ID_7 |
| 0days | r-shoulder__Average_Height | 0.78605563 | ID_8 |
| 0days | r-shoulder__Average_Height | 0.145472 | ID_9 |
| 0days | r-shoulder__Average_Height | -0.1727972 | ID_10 |
| 0days | r-shoulder__Average_Height | -0.001637 | ID_11 |
| 0days | r-shoulder__Average_Height | -0.0847022 | ID_12 |

| | | | |
|---|---|---|---|
| 0days | r-shoulder__Average_Height | 0.02663445 | ID_13 |
| 3days | r-shoulder__Average_Height | -0.3224065 | ID_1 |
| 3days | r-shoulder__Average_Height | -0.039671 | ID_2 |
| 3days | r-shoulder__Average_Height | 0.33247339 | ID_3 |
| 7days | r-shoulder__Average_Height | -0.0463295 | ID_1 |
| 7days | r-shoulder__Average_Height | 0.11004258 | ID_2 |
| 7days | r-shoulder__Average_Height | -0.0734568 | ID_3 |
| 7days | r-shoulder__Average_Height | -0.101915 | ID_4 |
| 7days | r-shoulder__Average_Height | -0.1659842 | ID_5 |
| 7days | r-shoulder__Average_Height | -0.1339037 | ID_6 |
| 7days | r-shoulder__Average_Height | -0.031272 | ID_7 |
| 7days | r-shoulder__Average_Height | -0.2939886 | ID_8 |
| 14days | r-shoulder__Average_Height | -0.1375322 | ID_1 |
| 14days | r-shoulder__Average_Height | -0.1722596 | ID_2 |
| 14days | r-shoulder__Average_Height | 0.00811027 | ID_3 |
| 14days | r-shoulder__Average_Height | -0.033674 | ID_4 |
| 14days | r-shoulder__Average_Height | -0.4199987 | ID_5 |
| 14days | r-shoulder__Average_Height | -0.016972 | ID_6 |
| 14days | r-shoulder__Average_Height | 0.03909872 | ID_7 |
| 14days | r-shoulder__Average_Height | 0.09131395 | ID_8 |
| 14days | r-shoulder__Average_Height | 0.18869352 | ID_9 |
| 21days | r-shoulder__Average_Height | -0.1975349 | ID_1 |
| 21days | r-shoulder__Average_Height | -0.2335856 | ID_2 |
| 21days | r-shoulder__Average_Height | -0.2111115 | ID_3 |
| 21days | r-shoulder__Average_Height | -0.0205237 | ID_4 |
| 21days | r-shoulder__Average_Height | -0.2132415 | ID_5 |
| 21days | r-shoulder__Average_Height | 0.42863289 | ID_6 |
| 21days | r-shoulder__Average_Height | -0.2062587 | ID_7 |
| 21days | r-shoulder__Average_Height | -0.1528395 | ID_8 |
| 0days | r-shoulder__Movement | 0.28274637 | ID_1 |
| 0days | r-shoulder__Movement | -0.5001339 | ID_2 |
| 0days | r-shoulder__Movement | 0.47819926 | ID_3 |
| 0days | r-shoulder__Movement | 1.48064408 | ID_4 |
| 0days | r-shoulder__Movement | 0.36421513 | ID_5 |
| 0days | r-shoulder__Movement | 0.37497867 | ID_6 |
| 0days | r-shoulder__Movement | -0.0597733 | ID_7 |
| 0days | r-shoulder__Movement | 1.38267606 | ID_8 |
| 0days | r-shoulder__Movement | 0 | ID_9 |
| 0days | r-shoulder__Movement | -0.2915515 | ID_10 |
| 0days | r-shoulder__Movement | -0.2088125 | ID_11 |
| 0days | r-shoulder__Movement | -0.3701008 | ID_12 |
| 0days | r-shoulder__Movement | -0.1323511 | ID_13 |
| 3days | r-shoulder__Movement | -0.7424135 | ID_1 |
| 3days | r-shoulder__Movement | -0.1246607 | ID_2 |
| 3days | r-shoulder__Movement | 0.36773151 | ID_3 |
| 7days | r-shoulder__Movement | -0.5371689 | ID_4 |
| 7days | r-shoulder__Movement | 0.04017472 | ID_5 |
| 7days | r-shoulder__Movement | -0.5611534 | ID_6 |
| 7days | r-shoulder__Movement | -0.4302203 | ID_7 |
| 7days | r-shoulder__Movement | -0.4650271 | ID_8 |
| 7days | r-shoulder__Movement | -0.3308973 | ID_1 |
| 7days | r-shoulder__Movement | -0.3384382 | ID_2 |
| 7days | r-shoulder__Movement | -0.2716098 | ID_3 |
| 14days | r-shoulder__Movement | -0.3166392 | ID_4 |
| 14days | r-shoulder__Movement | -0.4847344 | ID_5 |
| 14days | r-shoulder__Movement | 0.03914954 | ID_6 |

| | | | |
|---|---|---|---|
| 14days | r-shoulder__Movement | 0.53511622 | ID_7 |
| 14days | r-shoulder__Movement | -0.7019067 | ID_8 |
| 14days | r-shoulder__Movement | -0.4604283 | ID_9 |
| 14days | r-shoulder__Movement | -0.032709 | ID_1 |
| 14days | r-shoulder__Movement | 0.27185114 | ID_2 |
| 14days | r-shoulder__Movement | -0.1576145 | ID_3 |
| 21days | r-shoulder__Movement | -0.6614872 | ID_4 |
| 21days | r-shoulder__Movement | -0.4029086 | ID_5 |
| 21days | r-shoulder__Movement | -0.6768168 | ID_6 |
| 21days | r-shoulder__Movement | 0.09110983 | ID_7 |
| 21days | r-shoulder__Movement | -0.3405079 | ID_8 |
| 21days | r-shoulder__Movement | 0.2599305 | ID_9 |
| 21days | r-shoulder__Movement | -0.3577997 | ID_10 |
| 21days | r-shoulder__Movement | -0.4206061 | ID_11 |
| 0days | r-tail-base__Average_Height | -0.3279288 | ID_1 |
| 0days | r-tail-base__Average_Height | 0.18965672 | ID_2 |
| 0days | r-tail-base__Average_Height | -0.1291052 | ID_3 |
| 0days | r-tail-base__Average_Height | -0.1928172 | ID_4 |
| 0days | r-tail-base__Average_Height | 0.51735276 | ID_5 |
| 0days | r-tail-base__Average_Height | 1.24942018 | ID_6 |
| 0days | r-tail-base__Average_Height | -0.2173085 | ID_7 |
| 0days | r-tail-base__Average_Height | -0.2346401 | ID_8 |
| 0days | r-tail-base__Average_Height | -0.7510641 | ID_9 |
| 0days | r-tail-base__Average_Height | 0.12910519 | ID_10 |
| 0days | r-tail-base__Average_Height | 0.91701353 | ID_11 |
| 0days | r-tail-base__Average_Height | 0.62503555 | ID_12 |
| 3days | r-tail-base__Average_Height | -0.0605289 | ID_13 |
| 3days | r-tail-base__Average_Height | -0.0085418 | ID_1 |
| 3days | r-tail-base__Average_Height | -0.1436656 | ID_2 |
| 3days | r-tail-base__Average_Height | -0.0564892 | ID_3 |
| 3days | r-tail-base__Average_Height | -0.5335647 | ID_4 |
| 3days | r-tail-base__Average_Height | -0.3954393 | ID_5 |
| 3days | r-tail-base__Average_Height | 0.6222008 | ID_6 |
| 3days | r-tail-base__Average_Height | 0.44764986 | ID_7 |
| 7days | r-tail-base__Average_Height | 0.48413936 | ID_8 |
| 7days | r-tail-base__Average_Height | -0.7542402 | ID_1 |
| 7days | r-tail-base__Average_Height | -0.6463946 | ID_2 |
| 7days | r-tail-base__Average_Height | -0.4015599 | ID_3 |
| 7days | r-tail-base__Average_Height | -0.0265957 | ID_4 |
| 7days | r-tail-base__Average_Height | -0.2390397 | ID_5 |
| 7days | r-tail-base__Average_Height | 0.23780745 | ID_6 |
| 7days | r-tail-base__Average_Height | -0.3404965 | ID_7 |
| 7days | r-tail-base__Average_Height | -0.2709525 | ID_8 |
| 14days | r-tail-base__Average_Height | 0.16090946 | ID_9 |
| 14days | r-tail-base__Average_Height | -0.2433202 | ID_1 |
| 14days | r-tail-base__Average_Height | -0.1217874 | ID_2 |
| 14days | r-tail-base__Average_Height | -0.404408 | ID_3 |
| 14days | r-tail-base__Average_Height | -0.1276667 | ID_4 |
| 14days | r-tail-base__Average_Height | 0.48894002 | ID_5 |
| 14days | r-tail-base__Average_Height | -0.5521597 | ID_6 |
| 14days | r-tail-base__Average_Height | -0.6625228 | ID_7 |
| 14days | r-tail-base__Average_Height | 0.42708776 | ID_8 |
| 14days | r-tail-base__Average_Height | 0.52358889 | ID_9 |
| 14days | r-tail-base__Average_Height | -0.5269455 | ID_10 |
| 21days | r-tail-base__Average_Height | -0.1884373 | ID_11 |
| 21days | r-tail-base__Average_Height | -0.3999281 | ID_1 |

| | | | |
|---|---|---|---|
| 21days | r-tail-base__Average_Height | -0.1141487 | ID_2 |
| 21days | r-tail-base__Average_Height | -0.3506993 | ID_3 |
| 21days | r-tail-base__Average_Height | -0.4600231 | ID_4 |
| 21days | r-tail-base__Average_Height | -0.0594046 | ID_5 |
| 21days | r-tail-base__Average_Height | -0.6471636 | ID_6 |
| 21days | r-tail-base__Average_Height | -0.1401804 | ID_7 |
| 21days | r-tail-base__Average_Height | -0.074647 | ID_8 |
| 21days | r-tail-base__Average_Height | 0.25237762 | ID_9 |
| 0days | r-tail-base__Movement | -1.4884328 | ID_10 |
| 0days | r-tail-base__Movement | 0.6628308 | ID_1 |
| 0days | r-tail-base__Movement | -0.0280488 | ID_2 |
| 0days | r-tail-base__Movement | -0.7519261 | ID_3 |
| 0days | r-tail-base__Movement | 0.6841327 | ID_4 |
| 0days | r-tail-base__Movement | 1.21929734 | ID_5 |
| 0days | r-tail-base__Movement | -0.1554949 | ID_6 |
| 0days | r-tail-base__Movement | 0.02804877 | ID_7 |
| 0days | r-tail-base__Movement | -1.3594903 | ID_8 |
| 0days | r-tail-base__Movement | -0.4168364 | ID_9 |
| 0days | r-tail-base__Movement | 0.3103583 | ID_10 |
| 0days | r-tail-base__Movement | 0.44898195 | ID_11 |
| 3days | r-tail-base__Movement | -0.5096249 | ID_12 |
| 3days | r-tail-base__Movement | -0.2337252 | ID_13 |
| 3days | r-tail-base__Movement | 0.24137003 | ID_1 |
| 3days | r-tail-base__Movement | -0.2597581 | ID_2 |
| 3days | r-tail-base__Movement | -1.2060889 | ID_3 |
| 3days | r-tail-base__Movement | -0.4682711 | ID_4 |
| 3days | r-tail-base__Movement | 0.66638495 | ID_5 |
| 3days | r-tail-base__Movement | 0.66566255 | ID_6 |
| 7days | r-tail-base__Movement | 0.39704532 | ID_7 |
| 7days | r-tail-base__Movement | -1.146498 | ID_8 |
| 7days | r-tail-base__Movement | -1.4238269 | ID_1 |
| 7days | r-tail-base__Movement | -0.3049662 | ID_2 |
| 7days | r-tail-base__Movement | -0.6296294 | ID_3 |
| 7days | r-tail-base__Movement | -0.976006 | ID_4 |
| 7days | r-tail-base__Movement | -0.3438036 | ID_5 |
| 7days | r-tail-base__Movement | -1.4581196 | ID_6 |
| 7days | r-tail-base__Movement | -0.133536 | ID_7 |
| 14days | r-tail-base__Movement | 2.08222264 | ID_8 |
| 14days | r-tail-base__Movement | -0.4948627 | ID_9 |
| 14days | r-tail-base__Movement | -0.2319818 | ID_1 |
| 14days | r-tail-base__Movement | -1.3286843 | ID_2 |
| 14days | r-tail-base__Movement | -0.8492673 | ID_3 |
| 14days | r-tail-base__Movement | 0.62519524 | ID_4 |
| 14days | r-tail-base__Movement | -1.6729081 | ID_5 |
| 14days | r-tail-base__Movement | -1.5562913 | ID_6 |
| 14days | r-tail-base__Movement | 0.54171191 | ID_7 |
| 14days | r-tail-base__Movement | 0.09801549 | ID_8 |
| 14days | r-tail-base__Movement | -0.3923154 | ID_9 |
| 21days | r-tail-base__Movement | -0.5128268 | ID_10 |
| 21days | r-tail-base__Movement | -1.0002567 | ID_11 |
| 21days | r-tail-base__Movement | -0.7168314 | ID_1 |
| 21days | r-tail-base__Movement | -1.4626938 | ID_2 |
| 21days | r-tail-base__Movement | -1.5081911 | ID_3 |
| 21days | r-tail-base__Movement | -0.3173735 | ID_4 |
| 21days | r-tail-base__Movement | -0.6899591 | ID_5 |
| 21days | r-tail-base__Movement | -1.1208162 | ID_6 |

| | | | |
|---|---|---|---|
| 21days | r-tail-base__Movement | -1.1293678 | ID_7 |
| 21days | r-tail-base__Movement | -0.0105076 | ID_8 |
| 0days | r-wrist__Average_Height | 0.26295344 | ID_1 |
| 0days | r-wrist__Average_Height | 0.51838424 | ID_2 |
| 0days | r-wrist__Average_Height | 0.03283572 | ID_3 |
| 0days | r-wrist__Average_Height | 0 | ID_4 |
| 0days | r-wrist__Average_Height | -0.3560535 | ID_5 |
| 0days | r-wrist__Average_Height | -0.1617122 | ID_6 |
| 0days | r-wrist__Average_Height | -0.0015835 | ID_7 |
| 0days | r-wrist__Average_Height | 0.38365351 | ID_8 |
| 0days | r-wrist__Average_Height | -0.2755916 | ID_9 |
| 0days | r-wrist__Average_Height | -0.216229 | ID_10 |
| 0days | r-wrist__Average_Height | -0.5535379 | ID_11 |
| 0days | r-wrist__Average_Height | 0.06851116 | ID_12 |
| 0days | r-wrist__Average_Height | 0.23325182 | ID_13 |
| 3days | r-wrist__Average_Height | -0.87 | ID_1 |
| 3days | r-wrist__Average_Height | -0.6058212 | ID_2 |
| 3days | r-wrist__Average_Height | -0.4749152 | ID_3 |
| 3days | r-wrist__Average_Height | -0.7838648 | ID_4 |
| 3days | r-wrist__Average_Height | -0.4865431 | ID_5 |
| 3days | r-wrist__Average_Height | -0.2179645 | ID_6 |
| 3days | r-wrist__Average_Height | -0.5395759 | ID_7 |
| 3days | r-wrist__Average_Height | -0.3081241 | ID_8 |
| 7days | r-wrist__Average_Height | -0.2724075 | ID_1 |
| 7days | r-wrist__Average_Height | 0.52607768 | ID_2 |
| 7days | r-wrist__Average_Height | -0.3460832 | ID_3 |
| 7days | r-wrist__Average_Height | 0.28358721 | ID_4 |
| 7days | r-wrist__Average_Height | -0.228636 | ID_5 |
| 7days | r-wrist__Average_Height | 0.71822456 | ID_6 |
| 7days | r-wrist__Average_Height | -0.7131964 | ID_7 |
| 7days | r-wrist__Average_Height | -0.6053384 | ID_8 |
| 7days | r-wrist__Average_Height | -0.6369777 | ID_9 |
| 14days | r-wrist__Average_Height | -0.428482 | ID_1 |
| 14days | r-wrist__Average_Height | -0.2108008 | ID_2 |
| 14days | r-wrist__Average_Height | 0.49135368 | ID_3 |
| 14days | r-wrist__Average_Height | 0.30058789 | ID_4 |
| 14days | r-wrist__Average_Height | 0.38267076 | ID_5 |
| 14days | r-wrist__Average_Height | -0.9187281 | ID_6 |
| 14days | r-wrist__Average_Height | 0.2700295 | ID_7 |
| 14days | r-wrist__Average_Height | 0.45310704 | ID_8 |
| 14days | r-wrist__Average_Height | -0.5165896 | ID_9 |
| 14days | r-wrist__Average_Height | 0.139375 | ID_10 |
| 14days | r-wrist__Average_Height | -0.6044063 | ID_11 |
| 21days | r-wrist__Average_Height | -0.0892213 | ID_1 |
| 21days | r-wrist__Average_Height | 0.21994043 | ID_2 |
| 21days | r-wrist__Average_Height | -0.2067468 | ID_3 |
| 21days | r-wrist__Average_Height | 0.54698761 | ID_4 |
| 21days | r-wrist__Average_Height | -0.5284613 | ID_5 |
| 21days | r-wrist__Average_Height | 0.43272774 | ID_6 |
| 21days | r-wrist__Average_Height | -0.5033973 | ID_7 |
| 21days | r-wrist__Average_Height | 0.10798609 | ID_8 |
| 21days | r-wrist__Average_Height | 0.56595993 | ID_9 |
| 21days | r-wrist__Average_Height | 0.32389897 | ID_10 |
| 0days | r-wrist__Movement | 0.1908363 | ID_1 |
| 0days | r-wrist__Movement | 0.81883435 | ID_2 |
| 0days | r-wrist__Movement | -0.1474982 | ID_3 |

| | | | |
|---|---|---|---|
| 0days | r-wrist__Movement | 0.84177678 | ID_4 |
| 0days | r-wrist__Movement | 0.18972796 | ID_5 |
| 0days | r-wrist__Movement | -0.2184829 | ID_6 |
| 0days | r-wrist__Movement | -0.058932 | ID_7 |
| 0days | r-wrist__Movement | 1.2874895 | ID_8 |
| 0days | r-wrist__Movement | -0.0486132 | ID_9 |
| 0days | r-wrist__Movement | -0.2187028 | ID_10 |
| 0days | r-wrist__Movement | -0.47457 | ID_11 |
| 0days | r-wrist__Movement | 0 | ID_12 |
| 0days | r-wrist__Movement | 0.38107663 | ID_13 |
| 3days | r-wrist__Movement | -1.1042429 | ID_1 |
| 3days | r-wrist__Movement | -1.0237118 | ID_2 |
| 3days | r-wrist__Movement | -0.9050514 | ID_3 |
| 3days | r-wrist__Movement | -1.1928219 | ID_4 |
| 3days | r-wrist__Movement | 0.2480288 | ID_5 |
| 3days | r-wrist__Movement | -0.5360732 | ID_6 |
| 3days | r-wrist__Movement | -0.2693848 | ID_7 |
| 3days | r-wrist__Movement | -0.4260094 | ID_8 |
| 7days | r-wrist__Movement | -0.0994617 | ID_1 |
| 7days | r-wrist__Movement | 0.65778584 | ID_2 |
| 7days | r-wrist__Movement | -0.5010031 | ID_3 |
| 7days | r-wrist__Movement | 0.5203005 | ID_4 |
| 7days | r-wrist__Movement | -0.3331101 | ID_5 |
| 7days | r-wrist__Movement | 1.50076289 | ID_6 |
| 7days | r-wrist__Movement | -0.859766 | ID_7 |
| 7days | r-wrist__Movement | -0.7879348 | ID_8 |
| 7days | r-wrist__Movement | -0.7187565 | ID_9 |
| 14days | r-wrist__Movement | -0.8010218 | ID_1 |
| 14days | r-wrist__Movement | 0.11010138 | ID_2 |
| 14days | r-wrist__Movement | 0.50357344 | ID_3 |
| 14days | r-wrist__Movement | 0.2559005 | ID_4 |
| 14days | r-wrist__Movement | 0.81963793 | ID_5 |
| 14days | r-wrist__Movement | -1.1598687 | ID_6 |
| 14days | r-wrist__Movement | 0.64383236 | ID_7 |
| 14days | r-wrist__Movement | 0.53176811 | ID_8 |
| 14days | r-wrist__Movement | -0.2621248 | ID_9 |
| 14days | r-wrist__Movement | 0.34417921 | ID_10 |
| 14days | r-wrist__Movement | -0.5241281 | ID_11 |
| 21days | r-wrist__Movement | -0.1508607 | ID_1 |
| 21days | r-wrist__Movement | 0.52496405 | ID_2 |
| 21days | r-wrist__Movement | -0.332752 | ID_3 |
| 21days | r-wrist__Movement | 0.88027179 | ID_4 |
| 21days | r-wrist__Movement | -0.5483315 | ID_5 |
| 21days | r-wrist__Movement | 0.62183994 | ID_6 |
| 21days | r-wrist__Movement | -0.5415581 | ID_7 |
| 21days | r-wrist__Movement | 0.04978707 | ID_8 |
| 21days | r-wrist__Movement | 1.03831121 | ID_9 |
| 21days | r-wrist__Movement | 1.07516934 | ID_10 |

**Suppl. Table 20: Summary of side perspective height analysis**

| day | Measure | Value_Mean | Value_SD |
|---|---|---|---|
| 0days | l-back-ankle__Average_Heig | 0 | 0.67205514 |
| 0days | l-back-ankle__Movement | 0 | 1.39839106 |
| 0days | l-back-toe__Average_Height | 0 | 0.30972082 |
| 0days | l-back-toe__Movement | 0 | 1.13500976 |
| 0days | l-elbow__Average_Height | 0 | 0.2953487 |
| 0days | l-elbow__Movement | 0 | 0.56956062 |
| 0days | l-front-toe-tip__Average_Hei | 0 | 0.21328618 |
| 0days | l-front-toe-tip__Movement | 0 | 0.68814422 |
| 0days | l-head__Average_Height | 0 | 0.69609999 |
| 0days | l-head__Movement | 0 | 1.78519764 |
| 0days | l-hip__Average_Height | 0 | 0.27585357 |
| 0days | l-hip__Movement | 0 | 0.58467604 |
| 0days | l-iliac-crest__Average_Height | 0 | 0.29740806 |
| 0days | l-iliac-crest__Movement | 0 | 0.77458719 |
| 0days | l-shoulder__Average_Height | 0 | 0.34751774 |
| 0days | l-shoulder__Movement | 0 | 0.78403194 |
| 0days | l-tail-base__Average_Height | 0 | 0.36446301 |
| 0days | l-tail-base__Movement | 0 | 1.02050403 |
| 0days | l-wrist__Average_Height | 0 | 0.4812395 |
| 0days | l-wrist__Movement | 0 | 0.56512497 |
| 0days | r-back-ankle__Average_Heig | 0 | 0.42151241 |
| 0days | r-back-ankle__Movement | 0 | 0.81877275 |
| 0days | r-back-toe__Average_Height | 0 | 0.15521927 |
| 0days | r-back-toe__Movement | 0 | 0.61251167 |
| 0days | r-elbow__Average_Height | 0 | 0.27137972 |
| 0days | r-elbow__Movement | 0 | 0.58583293 |
| 0days | r-front-toe-tip__Average_Hei | 0 | 0.21423905 |
| 0days | r-front-toe-tip__Movement | 0 | 0.50975445 |
| 0days | r-head__Average_Height | 0 | 0.29488311 |
| 0days | r-head__Movement | 0 | 0.89707013 |
| 0days | r-hip__Average_Height | 0 | 0.27823373 |
| 0days | r-hip__Movement | 0 | 0.73561941 |
| 0days | r-iliac-crest__Average_Heigh | 0 | 0.24053368 |
| 0days | r-iliac-crest__Movement | 0 | 0.77555514 |
| 0days | r-shoulder__Average_Height | 0 | 0.30191406 |
| 0days | r-shoulder__Movement | 0 | 0.62044276 |
| 0days | r-tail-base__Average_Height | 0 | 0.57825888 |
| 0days | r-tail-base__Movement | 0 | 0.82356179 |
| 0days | r-wrist__Average_Height | 0 | 0.30645097 |
| 0days | r-wrist__Movement | 0 | 0.50869838 |
| 3days | l-back-ankle__Average_Heig | 0.02570948 | 0.58695474 |
| 3days | l-back-ankle__Movement | 0.01975225 | 1.19471529 |
| 3days | l-back-toe__Average_Height | -0.2515432 | 0.09761713 |
| 3days | l-back-toe__Movement | -1.1891297 | 0.30131302 |
| 3days | l-elbow__Average_Height | -0.1817734 | 0.11139445 |
| 3days | l-elbow__Movement | -0.6664027 | 0.23348492 |
| 3days | l-front-toe-tip__Average_Hei | -0.4894445 | 0.15474541 |
| 3days | l-front-toe-tip__Movement | -0.7720142 | 0.32233553 |
| 3days | l-head__Average_Height | -0.6188926 | 0.2477646 |
| 3days | l-head__Movement | -2.4197983 | 0.85032677 |
| 3days | l-hip__Average_Height | -0.3319208 | 0.26338908 |
| 3days | l-hip__Movement | -0.4688948 | 0.70446915 |
| 3days | l-iliac-crest__Average_Height | -0.1741278 | 0.2886531 |
| 3days | l-iliac-crest__Movement | -0.6873641 | 0.75218416 |

| | | | |
|---|---|---|---|
| 3days | l-shoulder__Average_Height | -0.2549553 | 0.1938969 |
| 3days | l-shoulder__Movement | -0.7711989 | 0.23496572 |
| 3days | l-tail-base__Average_Height | -0.1843617 | 0.50454738 |
| 3days | l-tail-base__Movement | -0.8913054 | 0.90303613 |
| 3days | l-wrist__Average_Height | -0.6604331 | 0.33195927 |
| 3days | l-wrist__Movement | -0.8620946 | 0.59232428 |
| 3days | r-back-ankle__Average_Heig | 0.29535306 | 0.54584935 |
| 3days | r-back-ankle__Movement | -0.2481358 | 1.08787015 |
| 3days | r-back-toe__Average_Height | -0.1167776 | 0.14477382 |
| 3days | r-back-toe__Movement | -0.2128761 | 0.65892732 |
| 3days | r-elbow__Average_Height | -0.4631945 | 0.25532092 |
| 3days | r-elbow__Movement | -0.9819639 | 0.52538211 |
| 3days | r-front-toe-tip__Average_Hei | -0.2258158 | 0.16613749 |
| 3days | r-front-toe-tip__Movement | -0.1309558 | 0.22474753 |
| 3days | r-head__Average_Height | -0.5023801 | 0.6731405 |
| 3days | r-head__Movement | -1.3000743 | 1.40052281 |
| 3days | r-hip__Average_Height | -0.3189425 | 0.27016954 |
| 3days | r-hip__Movement | -0.5717939 | 0.7964572 |
| 3days | r-iliac-crest__Average_Heigh | -0.0896488 | 0.19038016 |
| 3days | r-iliac-crest__Movement | -0.3541893 | 0.5509961 |
| 3days | r-shoulder__Average_Height | -0.039671 | 0.32845559 |
| 3days | r-shoulder__Movement | -0.1246607 | 0.55625091 |
| 3days | r-tail-base__Average_Height | -0.058509 | 0.38805286 |
| 3days | r-tail-base__Movement | -0.2467417 | 0.63794498 |
| 3days | r-wrist__Average_Height | -0.5130595 | 0.21945775 |
| 3days | r-wrist__Movement | -0.7205623 | 0.49571998 |
| 7days | l-back-ankle__Average_Heig | -0.351378 | 0.37982559 |
| 7days | l-back-ankle__Movement | -0.6315821 | 0.54145176 |
| 7days | l-back-toe__Average_Height | -0.18197 | 0.10763682 |
| 7days | l-back-toe__Movement | -1.1917231 | 0.4414276 |
| 7days | l-elbow__Average_Height | -0.1196973 | 0.19817834 |
| 7days | l-elbow__Movement | -0.2017046 | 0.64264963 |
| 7days | l-front-toe-tip__Average_Hei | -0.3830309 | 0.06593579 |
| 7days | l-front-toe-tip__Movement | -0.7220486 | 0.16284231 |
| 7days | l-head__Average_Height | -0.7840083 | 0.58142185 |
| 7days | l-head__Movement | -2.0484046 | 1.41819698 |
| 7days | l-hip__Average_Height | -0.2643041 | 0.28978815 |
| 7days | l-hip__Movement | -0.3605221 | 0.49323238 |
| 7days | l-iliac-crest__Average_Heigh | -0.2711173 | 0.29193128 |
| 7days | l-iliac-crest__Movement | -0.5854092 | 0.65743408 |
| 7days | l-shoulder__Average_Height | -0.2049188 | 0.15827927 |
| 7days | l-shoulder__Movement | -0.7485486 | 0.19805054 |
| 7days | l-tail-base__Average_Height | -0.19543 | 0.30844884 |
| 7days | l-tail-base__Movement | -0.5314102 | 0.79321631 |
| 7days | l-wrist__Average_Height | -0.7732756 | 0.24734098 |
| 7days | l-wrist__Movement | -1.0666631 | 0.46789948 |
| 7days | r-back-ankle__Average_Heig | 0.36533421 | 0.38194934 |
| 7days | r-back-ankle__Movement | 0.05341691 | 0.98345023 |
| 7days | r-back-toe__Average_Height | -0.0422763 | 0.32740692 |
| 7days | r-back-toe__Movement | 0.57444742 | 0.54023539 |
| 7days | r-elbow__Average_Height | -0.269897 | 0.17864471 |
| 7days | r-elbow__Movement | -0.9367159 | 0.215844 |
| 7days | r-front-toe-tip__Average_Hei | -0.1284495 | 0.28579316 |
| 7days | r-front-toe-tip__Movement | -0.1118996 | 0.55254655 |
| 7days | r-head__Average_Height | -0.2459201 | 0.57873624 |
| 7days | r-head__Movement | -1.0167984 | 1.4092157 |

| | | | |
|---|---|---|---|
| 7days | r-hip__Average_Height | -0.0476098 | 0.19670667 |
| 7days | r-hip__Movement | -0.076238 | 0.42739124 |
| 7days | r-iliac-crest__Average_Heigh | 0.03775729 | 0.34700319 |
| 7days | r-iliac-crest__Movement | -0.1402113 | 0.77575065 |
| 7days | r-shoulder__Average_Height | -0.0876859 | 0.11653689 |
| 7days | r-shoulder__Movement | -0.3843292 | 0.19162907 |
| 7days | r-tail-base__Average_Height | -0.2709525 | 0.39692806 |
| 7days | r-tail-base__Movement | -0.6296294 | 0.63025073 |
| 7days | r-wrist__Average_Height | -0.2724075 | 0.52672929 |
| 7days | r-wrist__Movement | -0.3331101 | 0.80322153 |
| 14days | l-back-ankle__Average_Heig | 0.0320646 | 0.6093304 |
| 14days | l-back-ankle__Movement | 0.27062048 | 0.84411346 |
| 14days | l-back-toe__Average_Height | -0.1521897 | 0.16525996 |
| 14days | l-back-toe__Movement | -0.8431799 | 0.82663937 |
| 14days | l-elbow__Average_Height | 0.17608252 | 0.3304861 |
| 14days | l-elbow__Movement | 0.70404172 | 0.74357192 |
| 14days | l-front-toe-tip__Average_Hei | -0.2986062 | 0.26680589 |
| 14days | l-front-toe-tip__Movement | -0.419892 | 0.58181483 |
| 14days | l-head__Average_Height | -0.1329957 | 0.62624392 |
| 14days | l-head__Movement | -0.9574556 | 1.68229377 |
| 14days | l-hip__Average_Height | -0.4010188 | 0.35154962 |
| 14days | l-hip__Movement | -0.0778198 | 0.70516224 |
| 14days | l-iliac-crest__Average_Height | -0.4039169 | 0.28894924 |
| 14days | l-iliac-crest__Movement | -0.6846737 | 0.79258113 |
| 14days | l-shoulder__Average_Height | 0.02657494 | 0.18503801 |
| 14days | l-shoulder__Movement | -0.1772168 | 0.41194747 |
| 14days | l-tail-base__Average_Height | -0.3294469 | 0.65252075 |
| 14days | l-tail-base__Movement | -0.9008344 | 1.13319296 |
| 14days | l-wrist__Average_Height | -0.7906645 | 0.4105328 |
| 14days | l-wrist__Movement | -0.7988941 | 0.54820836 |
| 14days | r-back-ankle__Average_Heig | 0.70032853 | 0.4892584 |
| 14days | r-back-ankle__Movement | 0.57465742 | 1.0550547 |
| 14days | r-back-toe__Average_Height | 0.14321988 | 0.31110989 |
| 14days | r-back-toe__Movement | 0.51693385 | 0.87934271 |
| 14days | r-elbow__Average_Height | -0.2750497 | 0.20660772 |
| 14days | r-elbow__Movement | -0.6878739 | 0.47170411 |
| 14days | r-front-toe-tip__Average_Hei | 0.03896691 | 0.2579201 |
| 14days | r-front-toe-tip__Movement | 0.19486483 | 0.59646224 |
| 14days | r-head__Average_Height | 0.00655582 | 0.68196822 |
| 14days | r-head__Movement | -0.4262264 | 1.52949285 |
| 14days | r-hip__Average_Height | -0.0463459 | 0.27163519 |
| 14days | r-hip__Movement | 0.08961341 | 0.5628902 |
| 14days | r-iliac-crest__Average_Heigh | 0.13260314 | 0.33662786 |
| 14days | r-iliac-crest__Movement | 0.39869722 | 0.65654526 |
| 14days | r-shoulder__Average_Height | -0.016972 | 0.17646677 |
| 14days | r-shoulder__Movement | -0.1576145 | 0.39330664 |
| 14days | r-tail-base__Average_Height | -0.1276667 | 0.43576028 |
| 14days | r-tail-base__Movement | -0.3923154 | 1.10773143 |
| 14days | r-wrist__Average_Height | 0.139375 | 0.49423445 |
| 14days | r-wrist__Movement | 0.2559005 | 0.64262869 |
| 21days | l-back-ankle__Average_Heig | -0.2302505 | 0.8024245 |
| 21days | l-back-ankle__Movement | -0.7434575 | 1.19399032 |
| 21days | l-back-toe__Average_Height | -0.0553157 | 0.16828268 |
| 21days | l-back-toe__Movement | -0.5940266 | 0.6753578 |
| 21days | l-elbow__Average_Height | 0.18487568 | 0.41207686 |
| 21days | l-elbow__Movement | 0.26455313 | 0.96353144 |

| | | | |
|---|---|---|---|
| 21days | l-front-toe-tip__Average_Hei | -0.0893582 | 0.26423018 |
| 21days | l-front-toe-tip__Movement | -0.1606363 | 0.57479182 |
| 21days | l-head__Average_Height | -0.3836985 | 0.66208614 |
| 21days | l-head__Movement | -1.8546309 | 1.604357 |
| 21days | l-hip__Average_Height | -0.1809973 | 0.26610034 |
| 21days | l-hip__Movement | -0.1865542 | 0.59149771 |
| 21days | l-iliac-crest__Average_Height | -0.0731095 | 0.33413492 |
| 21days | l-iliac-crest__Movement | -0.3395875 | 0.55901705 |
| 21days | l-shoulder__Average_Height | -0.0736804 | 0.18763304 |
| 21days | l-shoulder__Movement | -0.4397853 | 0.25248021 |
| 21days | l-tail-base__Average_Height | -0.2371947 | 0.25386117 |
| 21days | l-tail-base__Movement | -0.9806249 | 0.53855857 |
| 21days | l-wrist__Average_Height | -0.3176237 | 0.38897663 |
| 21days | l-wrist__Movement | 0.06593108 | 0.73611936 |
| 21days | r-back-ankle__Average_Heig | 0.42509925 | 0.43154865 |
| 21days | r-back-ankle__Movement | 0.73807505 | 0.99192076 |
| 21days | r-back-toe__Average_Height | 0.14850844 | 0.36265549 |
| 21days | r-back-toe__Movement | 0.49927316 | 0.99247557 |
| 21days | r-elbow__Average_Height | -0.0764663 | 0.35030255 |
| 21days | r-elbow__Movement | -0.4391015 | 0.8225804 |
| 21days | r-front-toe-tip__Average_Hei | -0.0184253 | 0.32348955 |
| 21days | r-front-toe-tip__Movement | 0.00602683 | 0.55799092 |
| 21days | r-head__Average_Height | -0.1934474 | 0.36130061 |
| 21days | r-head__Movement | -0.6266862 | 0.79998934 |
| 21days | r-hip__Average_Height | -0.1058788 | 0.35112317 |
| 21days | r-hip__Movement | -0.1093872 | 0.70830282 |
| 21days | r-iliac-crest__Average_Heigh | 0.01160277 | 0.38496129 |
| 21days | r-iliac-crest__Movement | -0.1016065 | 0.78793804 |
| 21days | r-shoulder__Average_Height | -0.2018968 | 0.22436402 |
| 21days | r-shoulder__Movement | -0.3803541 | 0.33118472 |
| 21days | r-tail-base__Average_Height | -0.1643088 | 0.25345077 |
| 21days | r-tail-base__Movement | -0.858544 | 0.48624961 |
| 21days | r-wrist__Average_Height | 0.16396326 | 0.405443 |
| 21days | r-wrist__Movement | 0.28737556 | 0.64259529 |

**Suppl. Table 21: Statistical test of bside perspective height analysis**

| Measure | group1 | group2 | p.adj | sig | Test |
|---|---|---|---|---|---|
| l-wrist__Movement | 0days | 7days | 0.00033674 | *** | Repeat ANOVA + PH test |
| l-wrist__Movement | 0days | 3days | 0.00034229 | *** | Repeat ANOVA + PH test |
| l-back-toe__Movement | 0days | 7days | 0.00086311 | *** | Repeat ANOVA + PH test |
| l-back-toe__Movement | 0days | 3days | 0.00086311 | *** | Repeat ANOVA + PH test |
| l-front-toe-tip__Movement | 0days | 7days | 0.00281907 | ** | Repeat ANOVA + PH test |
| l-front-toe-tip__Movement | 0days | 3days | 0.00662191 | ** | Repeat ANOVA + PH test |
| r-head__Movement | 0days | 3days | 0.00870331 | ** | Repeat ANOVA + PH test |
| r-back-ankle__Movement | 0days | 14days | 0.00887464 | ** | Repeat ANOVA + PH test |
| l-wrist__Movement | 0days | 14days | 0.0103341 | * | Repeat ANOVA + PH test |
| r-wrist__Movement | 0days | 3days | 0.01154712 | * | Repeat ANOVA + PH test |
| l-back-ankle__Movement | 0days | 21days | 0.01242591 | * | Repeat ANOVA + PH test |
| r-elbow__Movement | 0days | 7days | 0.01630839 | * | Repeat ANOVA + PH test |
| r-tail-base__Movement | 0days | 21days | 0.01656914 | * | Repeat ANOVA + PH test |
| l-shoulder__Movement | 0days | 7days | 0.01905166 | * | Repeat ANOVA + PH test |
| l-back-toe__Movement | 0days | 14days | 0.02112799 | * | Repeat ANOVA + PH test |
| l-back-toe__Movement | 0days | 21days | 0.02431522 | * | Repeat ANOVA + PH test |
| r-back-toe__Movement | 0days | 21days | 0.03658615 | * | Repeat ANOVA + PH test |
| l-wrist__Average_Height | 0days | 14days | 0.03960501 | * | Repeat ANOVA + PH test |
| l-wrist__Average_Height | 0days | 3days | 0.04154142 | * | Repeat ANOVA + PH test |
| l-shoulder__Movement | 0days | 3days | 0.04981773 | * | Repeat ANOVA + PH test |
| l-wrist__Average_Height | 0days | 7days | 0.05797498 | * | Repeat ANOVA + PH test |
| r-elbow__Movement | 0days | 3days | 0.05872624 | * | Repeat ANOVA + PH test |
| l-back-ankle__Movement | 0days | 3days | 0.06160991 | * | Repeat ANOVA + PH test |
| l-back-ankle__Movement | 0days | 7days | 0.06160991 | * | Repeat ANOVA + PH test |
| l-elbow__Movement | 0days | 3days | 0.0680736 | * | Repeat ANOVA + PH test |
| r-head__Movement | 0days | 7days | 0.07106673 | * | Repeat ANOVA + PH test |
| l-iliac-crest__Movement | 0days | 14days | 0.07607329 | * | Repeat ANOVA + PH test |
| r-head__Movement | 0days | 21days | 0.08018946 | * | Repeat ANOVA + PH test |
| r-tail-base__Movement | 0days | 7days | 0.09549484 | * | Repeat ANOVA + PH test |
| l-shoulder__Movement | 0days | 21days | 0.09593472 | * | Repeat ANOVA + PH test |
| r-elbow__Movement | 0days | 14days | 0.10261614 | ns | Repeat ANOVA + PH test |
| r-back-ankle__Movement | 0days | 21days | 0.1363321 | ns | Repeat ANOVA + PH test |
| l-front-toe-tip__Movement | 0days | 14days | 0.14456596 | ns | Repeat ANOVA + PH test |
| l-hip__Movement | 0days | 7days | 0.14519584 | ns | Repeat ANOVA + PH test |
| l-hip__Movement | 0days | 3days | 0.14519584 | ns | Repeat ANOVA + PH test |
| l-hip__Movement | 0days | 14days | 0.14519584 | ns | Repeat ANOVA + PH test |
| l-hip__Movement | 0days | 21days | 0.14519584 | ns | Repeat ANOVA + PH test |
| l-front-toe-tip__Movement | 0days | 21days | 0.16410928 | ns | Repeat ANOVA + PH test |
| r-shoulder__Movement | 0days | 7days | 0.16842294 | ns | Repeat ANOVA + PH test |
| l-elbow__Movement | 0days | 21days | 0.18648537 | ns | Repeat ANOVA + PH test |
| r-shoulder__Movement | 0days | 21days | 0.18735186 | ns | Repeat ANOVA + PH test |
| r-wrist__Average_Height | 0days | 3days | 0.24632881 | ns | Repeat ANOVA + PH test |
| l-shoulder__Movement | 0days | 14days | 0.26841028 | ns | Repeat ANOVA + PH test |
| l-tail-base__Movement | 0days | 21days | 0.28857549 | ns | Repeat ANOVA + PH test |
| r-head__Movement | 0days | 14days | 0.29530385 | ns | Repeat ANOVA + PH test |
| l-iliac-crest__Movement | 0days | 7days | 0.3239988 | ns | Repeat ANOVA + PH test |
| l-iliac-crest__Movement | 0days | 3days | 0.3239988 | ns | Repeat ANOVA + PH test |
| l-iliac-crest__Movement | 0days | 21days | 0.3239988 | ns | Repeat ANOVA + PH test |
| r-shoulder__Movement | 0days | 14days | 0.37581512 | ns | Repeat ANOVA + PH test |
| r-shoulder__Movement | 0days | 3days | 0.37581512 | ns | Repeat ANOVA + PH test |
| l-elbow__Movement | 0days | 14days | 0.38853879 | ns | Repeat ANOVA + PH test |
| r-back-toe__Movement | 0days | 14days | 0.40695018 | ns | Repeat ANOVA + PH test |
| r-back-toe__Movement | 0days | 7days | 0.40778698 | ns | Repeat ANOVA + PH test |
| l-elbow__Movement | 0days | 7days | 0.4104934 | ns | Repeat ANOVA + PH test |

| | | | | | |
|---|---|---|---|---|---|
| l-wrist__Average_Height | 0days | 21days | 0.4227289 | ns | Repeat ANOVA + PH test |
| l-back-ankle__Movement | 0days | 14days | 0.51051272 | ns | Repeat ANOVA + PH test |
| r-iliac-crest__Movement | 0days | 3days | 0.51755507 | ns | Repeat ANOVA + PH test |
| r-hip__Movement | 0days | 3days | 0.51909376 | ns | Repeat ANOVA + PH test |
| r-elbow__Movement | 0days | 21days | 0.54997212 | ns | Repeat ANOVA + PH test |
| l-front-toe-tip__Average_He | 0days | 3days | 0.62671622 | ns | Repeat ANOVA + PH test |
| r-tail-base__Average_Height | 0days | 21days | 0.70378738 | ns | Repeat ANOVA + PH test |
| r-tail-base__Average_Height | 0days | 7days | 0.70378738 | ns | Repeat ANOVA + PH test |
| r-tail-base__Average_Height | 0days | 14days | 0.71642155 | ns | Repeat ANOVA + PH test |
| r-tail-base__Average_Height | 0days | 3days | 0.71642155 | ns | Repeat ANOVA + PH test |
| l-wrist__Movement | 0days | 21days | 0.72937234 | ns | Repeat ANOVA + PH test |
| l-front-toe-tip__Average_He | 0days | 7days | 0.74647315 | ns | Repeat ANOVA + PH test |
| l-front-toe-tip__Average_He | 0days | 14days | 0.74647315 | ns | Repeat ANOVA + PH test |
| l-front-toe-tip__Average_He | 0days | 21days | 0.74647315 | ns | Repeat ANOVA + PH test |
| r-back-ankle__Average_Hei | 0days | 14days | 0.80579192 | ns | Repeat ANOVA + PH test |
| r-tail-base__Movement | 0days | 14days | 0.81479353 | ns | Repeat ANOVA + PH test |
| r-tail-base__Movement | 0days | 3days | 0.8150049 | ns | Repeat ANOVA + PH test |
| r-back-toe__Movement | 0days | 3days | 0.89626413 | ns | Repeat ANOVA + PH test |
| l-iliac-crest__Average_Heigh | 0days | 14days | 0.9075294 | ns | Repeat ANOVA + PH test |
| l-tail-base__Movement | 0days | 14days | 0.92793154 | ns | Repeat ANOVA + PH test |
| l-back-ankle__Average_Hei | 0days | 7days | 1 | ns | Repeat ANOVA + PH test |
| l-back-ankle__Average_Hei | 0days | 21days | 1 | ns | Repeat ANOVA + PH test |
| l-back-ankle__Average_Hei | 0days | 14days | 1 | ns | Repeat ANOVA + PH test |
| l-back-ankle__Average_Hei | 0days | 3days | 1 | ns | Repeat ANOVA + PH test |
| l-back-toe__Average_Heigh | 0days | 3days | 1 | ns | Repeat ANOVA + PH test |
| l-back-toe__Average_Heigh | 0days | 7days | 1 | ns | Repeat ANOVA + PH test |
| l-back-toe__Average_Heigh | 0days | 14days | 1 | ns | Repeat ANOVA + PH test |
| l-back-toe__Average_Heigh | 0days | 21days | 1 | ns | Repeat ANOVA + PH test |
| l-elbow__Average_Height | 0days | 21days | 1 | ns | Repeat ANOVA + PH test |
| l-elbow__Average_Height | 0days | 14days | 1 | ns | Repeat ANOVA + PH test |
| l-elbow__Average_Height | 0days | 3days | 1 | ns | Repeat ANOVA + PH test |
| l-elbow__Average_Height | 0days | 7days | 1 | ns | Repeat ANOVA + PH test |
| l-hip__Average_Height | 0days | 14days | 1 | ns | Repeat ANOVA + PH test |
| l-hip__Average_Height | 0days | 7days | 1 | ns | Repeat ANOVA + PH test |
| l-hip__Average_Height | 0days | 3days | 1 | ns | Repeat ANOVA + PH test |
| l-hip__Average_Height | 0days | 21days | 1 | ns | Repeat ANOVA + PH test |
| l-iliac-crest__Average_Heigh | 0days | 7days | 1 | ns | Repeat ANOVA + PH test |
| l-iliac-crest__Average_Heigh | 0days | 3days | 1 | ns | Repeat ANOVA + PH test |
| l-iliac-crest__Average_Heigh | 0days | 21days | 1 | ns | Repeat ANOVA + PH test |
| l-shoulder__Average_Heigh | 0days | 7days | 1 | ns | Repeat ANOVA + PH test |
| l-shoulder__Average_Heigh | 0days | 3days | 1 | ns | Repeat ANOVA + PH test |
| l-shoulder__Average_Heigh | 0days | 21days | 1 | ns | Repeat ANOVA + PH test |
| l-shoulder__Average_Heigh | 0days | 14days | 1 | ns | Repeat ANOVA + PH test |
| l-tail-base__Average_Height | 0days | 21days | 1 | ns | Repeat ANOVA + PH test |
| l-tail-base__Average_Height | 0days | 7days | 1 | ns | Repeat ANOVA + PH test |
| l-tail-base__Average_Height | 0days | 3days | 1 | ns | Repeat ANOVA + PH test |
| l-tail-base__Average_Height | 0days | 14days | 1 | ns | Repeat ANOVA + PH test |
| l-tail-base__Movement | 0days | 7days | 1 | ns | Repeat ANOVA + PH test |
| l-tail-base__Movement | 0days | 3days | 1 | ns | Repeat ANOVA + PH test |
| r-back-ankle__Average_Hei | 0days | 21days | 1 | ns | Repeat ANOVA + PH test |
| r-back-ankle__Average_Hei | 0days | 7days | 1 | ns | Repeat ANOVA + PH test |
| r-back-ankle__Average_Hei | 0days | 3days | 1 | ns | Repeat ANOVA + PH test |
| r-back-ankle__Movement | 0days | 7days | 1 | ns | Repeat ANOVA + PH test |
| r-back-ankle__Movement | 0days | 3days | 1 | ns | Repeat ANOVA + PH test |
| r-back-toe__Average_Heigh | 0days | 21days | 1 | ns | Repeat ANOVA + PH test |
| r-back-toe__Average_Heigh | 0days | 14days | 1 | ns | Repeat ANOVA + PH test |

| | | | | | |
|---|---|---|---|---|---|
| r-back-toe__Average_Heigh | 0days | 7days | 1 | ns | Repeat ANOVA + PH test |
| r-back-toe__Average_Heigh | 0days | 3days | 1 | ns | Repeat ANOVA + PH test |
| r-elbow__Average_Height | 0days | 3days | 1 | ns | Repeat ANOVA + PH test |
| r-elbow__Average_Height | 0days | 14days | 1 | ns | Repeat ANOVA + PH test |
| r-elbow__Average_Height | 0days | 7days | 1 | ns | Repeat ANOVA + PH test |
| r-elbow__Average_Height | 0days | 21days | 1 | ns | Repeat ANOVA + PH test |
| r-front-toe-tip__Average_He | 0days | 3days | 1 | ns | Repeat ANOVA + PH test |
| r-front-toe-tip__Average_He | 0days | 7days | 1 | ns | Repeat ANOVA + PH test |
| r-front-toe-tip__Average_He | 0days | 14days | 1 | ns | Repeat ANOVA + PH test |
| r-front-toe-tip__Average_He | 0days | 21days | 1 | ns | Repeat ANOVA + PH test |
| r-front-toe-tip__Movement | 0days | 3days | 1 | ns | Repeat ANOVA + PH test |
| r-front-toe-tip__Movement | 0days | 21days | 1 | ns | Repeat ANOVA + PH test |
| r-front-toe-tip__Movement | 0days | 14days | 1 | ns | Repeat ANOVA + PH test |
| r-front-toe-tip__Movement | 0days | 7days | 1 | ns | Repeat ANOVA + PH test |
| r-head__Average_Height | 0days | 3days | 1 | ns | Repeat ANOVA + PH test |
| r-head__Average_Height | 0days | 7days | 1 | ns | Repeat ANOVA + PH test |
| r-head__Average_Height | 0days | 14days | 1 | ns | Repeat ANOVA + PH test |
| r-head__Average_Height | 0days | 21days | 1 | ns | Repeat ANOVA + PH test |
| r-hip__Average_Height | 0days | 3days | 1 | ns | Repeat ANOVA + PH test |
| r-hip__Average_Height | 0days | 7days | 1 | ns | Repeat ANOVA + PH test |
| r-hip__Average_Height | 0days | 14days | 1 | ns | Repeat ANOVA + PH test |
| r-hip__Average_Height | 0days | 21days | 1 | ns | Repeat ANOVA + PH test |
| r-hip__Movement | 0days | 14days | 1 | ns | Repeat ANOVA + PH test |
| r-hip__Movement | 0days | 7days | 1 | ns | Repeat ANOVA + PH test |
| r-hip__Movement | 0days | 21days | 1 | ns | Repeat ANOVA + PH test |
| r-iliac-crest__Average_Heigl | 0days | 3days | 1 | ns | Repeat ANOVA + PH test |
| r-iliac-crest__Average_Heigl | 0days | 14days | 1 | ns | Repeat ANOVA + PH test |
| r-iliac-crest__Average_Heigl | 0days | 7days | 1 | ns | Repeat ANOVA + PH test |
| r-iliac-crest__Average_Heigl | 0days | 21days | 1 | ns | Repeat ANOVA + PH test |
| r-iliac-crest__Movement | 0days | 14days | 1 | ns | Repeat ANOVA + PH test |
| r-iliac-crest__Movement | 0days | 21days | 1 | ns | Repeat ANOVA + PH test |
| r-iliac-crest__Movement | 0days | 7days | 1 | ns | Repeat ANOVA + PH test |
| r-shoulder__Average_Heigh | 0days | 21days | 1 | ns | Repeat ANOVA + PH test |
| r-shoulder__Average_Heigh | 0days | 7days | 1 | ns | Repeat ANOVA + PH test |
| r-shoulder__Average_Heigh | 0days | 14days | 1 | ns | Repeat ANOVA + PH test |
| r-shoulder__Average_Heigh | 0days | 3days | 1 | ns | Repeat ANOVA + PH test |
| r-wrist__Average_Height | 0days | 7days | 1 | ns | Repeat ANOVA + PH test |
| r-wrist__Average_Height | 0days | 21days | 1 | ns | Repeat ANOVA + PH test |
| r-wrist__Average_Height | 0days | 14days | 1 | ns | Repeat ANOVA + PH test |
| r-wrist__Movement | 0days | 7days | 1 | ns | Repeat ANOVA + PH test |
| r-wrist__Movement | 0days | 14days | 1 | ns | Repeat ANOVA + PH test |
| r-wrist__Movement | 0days | 21days | 1 | ns | Repeat ANOVA + PH test |

**Suppl. Table 22: Raw data side perspective horizontal analysis**

| day | ID | Measure | Length, mm |
|---|---|---|---|
| 0days | ID_1 | left__back__average | -0.665732881 |
| 0days | ID_2 | left__back__average | -2.43937031 |
| 0days | ID_3 | left__back__average | -3.165219533 |
| 0days | ID_4 | left__back__average | 2.780764825 |
| 0days | ID_5 | left__back__average | -1.016136842 |
| 0days | ID_6 | left__back__average | -0.11607343 |
| 0days | ID_7 | left__back__average | 0.331419497 |
| 0days | ID_8 | left__back__average | 0.998726988 |
| 0days | ID_9 | left__back__average | 1.029317169 |
| 0days | ID_10 | left__back__average | 1.439472752 |
| 0days | ID_11 | left__back__average | -6.288500711 |
| 0days | ID_12 | left__back__average | 0.790370464 |
| 0days | ID_13 | left__back__average | 0 |
| 3days | ID_1 | left__back__average | -3.252123236 |
| 3days | ID_2 | left__back__average | -0.693573233 |
| 3days | ID_3 | left__back__average | -1.321642182 |
| 3days | ID_4 | left__back__average | -4.446968622 |
| 3days | ID_5 | left__back__average | -5.077335534 |
| 3days | ID_6 | left__back__average | -2.322387965 |
| 3days | ID_7 | left__back__average | 0.183361776 |
| 3days | ID_8 | left__back__average | -0.502326708 |
| 7days | ID_1 | left__back__average | -2.16032667 |
| 7days | ID_2 | left__back__average | -2.015743425 |
| 7days | ID_3 | left__back__average | -2.928351493 |
| 7days | ID_4 | left__back__average | -2.23863132 |
| 7days | ID_5 | left__back__average | -2.862697583 |
| 7days | ID_6 | left__back__average | -2.712748448 |
| 7days | ID_7 | left__back__average | -3.986746149 |
| 7days | ID_8 | left__back__average | -1.923224411 |
| 7days | ID_9 | left__back__average | -2.317728529 |
| 14days | ID_1 | left__back__average | 0.323925364 |
| 14days | ID_2 | left__back__average | -3.950857748 |
| 14days | ID_3 | left__back__average | -2.796805228 |
| 14days | ID_4 | left__back__average | -0.687508374 |
| 14days | ID_5 | left__back__average | -0.947685938 |
| 14days | ID_6 | left__back__average | -7.996255861 |
| 14days | ID_7 | left__back__average | -1.453731628 |
| 14days | ID_8 | left__back__average | -1.463413908 |
| 14days | ID_9 | left__back__average | -0.913276722 |
| 14days | ID_10 | left__back__average | -4.226148943 |
| 14days | ID_11 | left__back__average | 1.900302369 |
| 21days | ID_1 | left__back__average | -0.672312122 |
| 21days | ID_2 | left__back__average | -0.248371865 |
| 21days | ID_3 | left__back__average | -3.100045223 |
| 21days | ID_4 | left__back__average | -0.890662024 |
| 21days | ID_5 | left__back__average | 1.142253382 |

| | | | |
|---|---|---|---|
| 21days | ID_6 | left__back__average | 0.988501652 |
| 21days | ID_7 | left__back__average | 0.482267541 |
| 21days | ID_8 | left__back__average | 1.844449648 |
| 21days | ID_9 | left__back__average | 1.094912192 |
| 21days | ID_10 | left__back__average | -0.679299503 |
| 0days | ID_1 | left__back__median | -1.674089125 |
| 0days | ID_2 | left__back__median | -4.439980208 |
| 0days | ID_3 | left__back__median | -5.078559266 |
| 0days | ID_4 | left__back__median | 2.833925522 |
| 0days | ID_5 | left__back__median | -2.598521964 |
| 0days | ID_6 | left__back__median | -0.140126659 |
| 0days | ID_7 | left__back__median | 0 |
| 0days | ID_8 | left__back__median | 0.999616945 |
| 0days | ID_9 | left__back__median | 1.562406868 |
| 0days | ID_10 | left__back__median | 2.202879524 |
| 0days | ID_11 | left__back__median | -8.200006562 |
| 0days | ID_12 | left__back__median | 0.828872722 |
| 0days | ID_13 | left__back__median | 0.040724184 |
| 3days | ID_1 | left__back__median | -6.2651764 |
| 3days | ID_2 | left__back__median | -2.219129741 |
| 3days | ID_3 | left__back__median | -3.693438635 |
| 3days | ID_4 | left__back__median | -7.473039746 |
| 3days | ID_5 | left__back__median | -6.403805071 |
| 3days | ID_6 | left__back__median | -4.467193939 |
| 3days | ID_7 | left__back__median | 1.544503241 |
| 3days | ID_8 | left__back__median | -0.534412942 |
| 7days | ID_1 | left__back__median | -3.80758827 |
| 7days | ID_2 | left__back__median | -3.348074382 |
| 7days | ID_3 | left__back__median | -3.855174598 |
| 7days | ID_4 | left__back__median | -1.834017477 |
| 7days | ID_5 | left__back__median | -3.759343996 |
| 7days | ID_6 | left__back__median | -3.443448227 |
| 7days | ID_7 | left__back__median | -4.111624234 |
| 7days | ID_8 | left__back__median | -1.317366042 |
| 7days | ID_9 | left__back__median | -1.844135333 |
| 14days | ID_1 | left__back__median | -0.695586301 |
| 14days | ID_2 | left__back__median | -6.229323465 |
| 14days | ID_3 | left__back__median | -4.933527567 |
| 14days | ID_4 | left__back__median | -0.795866341 |
| 14days | ID_5 | left__back__median | -1.176232972 |
| 14days | ID_6 | left__back__median | -12.74506704 |
| 14days | ID_7 | left__back__median | -1.659057825 |
| 14days | ID_8 | left__back__median | -1.623890045 |
| 14days | ID_9 | left__back__median | -3.03993928 |
| 14days | ID_10 | left__back__median | -4.904743153 |
| 14days | ID_11 | left__back__median | 2.441005261 |
| 21days | ID_1 | left__back__median | -2.050191019 |
| 21days | ID_2 | left__back__median | 0.448892446 |

| | | | |
|---|---|---|---|
| 21days | ID_3 | left__back__median | -4.139947609 |
| 21days | ID_4 | left__back__median | -1.527488306 |
| 21days | ID_5 | left__back__median | 1.802578924 |
| 21days | ID_6 | left__back__median | 0.815885025 |
| 21days | ID_7 | left__back__median | 1.306126126 |
| 21days | ID_8 | left__back__median | 1.568666375 |
| 21days | ID_9 | left__back__median | 0.326482619 |
| 21days | ID_10 | left__back__median | -1.795462783 |
| 0days | ID_1 | left__back__protraction | -1.007182427 |
| 0days | ID_2 | left__back__protraction | -3.653931951 |
| 0days | ID_3 | left__back__protraction | -2.141409021 |
| 0days | ID_4 | left__back__protraction | 0.576821813 |
| 0days | ID_5 | left__back__protraction | 0.309006776 |
| 0days | ID_6 | left__back__protraction | 1.158577959 |
| 0days | ID_7 | left__back__protraction | 0.596753465 |
| 0days | ID_8 | left__back__protraction | -1.254364972 |
| 0days | ID_9 | left__back__protraction | 0.226353945 |
| 0days | ID_10 | left__back__protraction | 0 |
| 0days | ID_11 | left__back__protraction | -4.147008126 |
| 0days | ID_12 | left__back__protraction | 0.754228211 |
| 0days | ID_13 | left__back__protraction | -0.60652709 |
| 3days | ID_1 | left__back__protraction | -1.274293944 |
| 3days | ID_2 | left__back__protraction | 0.096048212 |
| 3days | ID_3 | left__back__protraction | -1.865729097 |
| 3days | ID_4 | left__back__protraction | -1.979965267 |
| 3days | ID_5 | left__back__protraction | -5.552383843 |
| 3days | ID_6 | left__back__protraction | -1.237684327 |
| 3days | ID_7 | left__back__protraction | -1.105148234 |
| 3days | ID_8 | left__back__protraction | -0.442272116 |
| 7days | ID_1 | left__back__protraction | -2.661670395 |
| 7days | ID_2 | left__back__protraction | -3.245417491 |
| 7days | ID_3 | left__back__protraction | -1.545360848 |
| 7days | ID_4 | left__back__protraction | -2.447104324 |
| 7days | ID_5 | left__back__protraction | -2.507040057 |
| 7days | ID_6 | left__back__protraction | -2.07934681 |
| 7days | ID_7 | left__back__protraction | -0.816228153 |
| 7days | ID_8 | left__back__protraction | -1.485682354 |
| 7days | ID_9 | left__back__protraction | -1.402951927 |
| 14days | ID_1 | left__back__protraction | -0.324161835 |
| 14days | ID_2 | left__back__protraction | -1.57424086 |
| 14days | ID_3 | left__back__protraction | -2.527734181 |
| 14days | ID_4 | left__back__protraction | -1.709245047 |
| 14days | ID_5 | left__back__protraction | -1.607006233 |
| 14days | ID_6 | left__back__protraction | -3.121501111 |
| 14days | ID_7 | left__back__protraction | -0.924197633 |
| 14days | ID_8 | left__back__protraction | -1.643220046 |
| 14days | ID_9 | left__back__protraction | -0.153902842 |
| 14days | ID_10 | left__back__protraction | -3.418664461 |

| | | | |
|---|---|---|---|
| 14days | ID_11 | left__back__protraction | 2.046542141 |
| 21days | ID_1 | left__back__protraction | -0.24156792 |
| 21days | ID_2 | left__back__protraction | 1.626731032 |
| 21days | ID_3 | left__back__protraction | 0.851509272 |
| 21days | ID_4 | left__back__protraction | 0.064025612 |
| 21days | ID_5 | left__back__protraction | 1.261124624 |
| 21days | ID_6 | left__back__protraction | 0.445258176 |
| 21days | ID_7 | left__back__protraction | 0.348099895 |
| 21days | ID_8 | left__back__protraction | 0.664082467 |
| 21days | ID_9 | left__back__protraction | 0.599460844 |
| 21days | ID_10 | left__back__protraction | 0.506442497 |
| 0days | ID_8 | left__back__retraction | 1.210140798 |
| 0days | ID_9 | left__back__retraction | 0.02219099 |
| 0days | ID_10 | left__back__retraction | -1.162018988 |
| 0days | ID_8 | left__back__retraction | 3.017222757 |
| 0days | ID_9 | left__back__retraction | -1.27637061 |
| 0days | ID_10 | left__back__retraction | -1.538501377 |
| 0days | ID_1 | left__back__retraction | 0.12559081 |
| 0days | ID_2 | left__back__retraction | 3.889002621 |
| 0days | ID_3 | left__back__retraction | 1.010955045 |
| 0days | ID_4 | left__back__retraction | -0.341514206 |
| 0days | ID_5 | left__back__retraction | -8.110801699 |
| 0days | ID_6 | left__back__retraction | 0 |
| 0days | ID_7 | left__back__retraction | -0.57797182 |
| 3days | ID_8 | left__back__retraction | -2.819717048 |
| 3days | ID_9 | left__back__retraction | -0.135272303 |
| 3days | ID_10 | left__back__retraction | 0.460051372 |
| 3days | ID_11 | left__back__retraction | -2.545966881 |
| 3days | ID_12 | left__back__retraction | -2.757772999 |
| 3days | ID_13 | left__back__retraction | -1.293005832 |
| 3days | ID_1 | left__back__retraction | -1.836321935 |
| 3days | ID_2 | left__back__retraction | 1.001731981 |
| 7days | ID_3 | left__back__retraction | 0.13659226 |
| 7days | ID_4 | left__back__retraction | 0.549676978 |
| 7days | ID_5 | left__back__retraction | -3.194304685 |
| 7days | ID_6 | left__back__retraction | -2.038100464 |
| 7days | ID_7 | left__back__retraction | -2.368477429 |
| 7days | ID_8 | left__back__retraction | -2.694002504 |
| 7days | ID_1 | left__back__retraction | -6.256429814 |
| 7days | ID_2 | left__back__retraction | -4.775949884 |
| 7days | ID_3 | left__back__retraction | -1.625930648 |
| 14days | ID_4 | left__back__retraction | 1.87976914 |
| 14days | ID_5 | left__back__retraction | -2.587458199 |
| 14days | ID_6 | left__back__retraction | -0.883775699 |
| 14days | ID_7 | left__back__retraction | -1.009640645 |
| 14days | ID_8 | left__back__retraction | -1.44399568 |
| 14days | ID_9 | left__back__retraction | -8.1193916 |
| 14days | ID_1 | left__back__retraction | -2.842384321 |

| | | | |
|---|---|---|---|
| 14days | ID_2 | left__back__retraction | -2.638135661 |
| 14days | ID_3 | left__back__retraction | 0.80479594 |
| 14days | ID_4 | left__back__retraction | -5.799396098 |
| 14days | ID_5 | left__back__retraction | -0.789313388 |
| 21days | ID_6 | left__back__retraction | 0.243836979 |
| 21days | ID_7 | left__back__retraction | -2.424044339 |
| 21days | ID_8 | left__back__retraction | -6.781224612 |
| 21days | ID_9 | left__back__retraction | -0.216298896 |
| 21days | ID_10 | left__back__retraction | -0.419762593 |
| 21days | ID_11 | left__back__retraction | -0.177530285 |
| 21days | ID_1 | left__back__retraction | -0.789324573 |
| 21days | ID_2 | left__back__retraction | 1.585602295 |
| 21days | ID_3 | left__back__retraction | 2.569213531 |
| 21days | ID_4 | left__back__retraction | -0.980244494 |
| 0days | ID_1 | left__front__average | -0.879607339 |
| 0days | ID_2 | left__front__average | -0.709980082 |
| 0days | ID_3 | left__front__average | 0 |
| 0days | ID_4 | left__front__average | 0.504144505 |
| 0days | ID_5 | left__front__average | -1.40961481 |
| 0days | ID_6 | left__front__average | -0.527251359 |
| 0days | ID_7 | left__front__average | 1.072535671 |
| 0days | ID_8 | left__front__average | 0.448437723 |
| 0days | ID_9 | left__front__average | 0.181167374 |
| 0days | ID_10 | left__front__average | -0.005719772 |
| 0days | ID_11 | left__front__average | -3.404842295 |
| 0days | ID_12 | left__front__average | 0.316464062 |
| 0days | ID_13 | left__front__average | 1.607013021 |
| 3days | ID_1 | left__front__average | -0.907181554 |
| 3days | ID_2 | left__front__average | -0.401869236 |
| 3days | ID_3 | left__front__average | 0.335036403 |
| 3days | ID_4 | left__front__average | -2.658869037 |
| 3days | ID_5 | left__front__average | -2.227689565 |
| 3days | ID_6 | left__front__average | -0.721798769 |
| 3days | ID_7 | left__front__average | 0.585521939 |
| 3days | ID_8 | left__front__average | -0.258398494 |
| 7days | ID_1 | left__front__average | 0.025801317 |
| 7days | ID_2 | left__front__average | -1.214567938 |
| 7days | ID_3 | left__front__average | -1.002909718 |
| 7days | ID_4 | left__front__average | -1.739412223 |
| 7days | ID_5 | left__front__average | 0.549012538 |
| 7days | ID_6 | left__front__average | -3.108332693 |
| 7days | ID_7 | left__front__average | -1.714345392 |
| 7days | ID_8 | left__front__average | 0.232114186 |
| 7days | ID_9 | left__front__average | -0.193090236 |
| 14days | ID_1 | left__front__average | 1.910567496 |
| 14days | ID_2 | left__front__average | 0.082685496 |
| 14days | ID_3 | left__front__average | -1.241697683 |
| 14days | ID_4 | left__front__average | -1.110237164 |

| | | | |
|---|---|---|---|
| 14days | ID_5 | left__front__average | -0.247411429 |
| 14days | ID_6 | left__front__average | -1.310406232 |
| 14days | ID_7 | left__front__average | -1.554612921 |
| 14days | ID_8 | left__front__average | -1.724099018 |
| 14days | ID_9 | left__front__average | 0.254252606 |
| 14days | ID_10 | left__front__average | 0.237044329 |
| 14days | ID_11 | left__front__average | 0.660205908 |
| 21days | ID_1 | left__front__average | -0.540968367 |
| 21days | ID_2 | left__front__average | -0.860216841 |
| 21days | ID_3 | left__front__average | -1.483103708 |
| 21days | ID_4 | left__front__average | 0.362900494 |
| 21days | ID_5 | left__front__average | -0.284639455 |
| 21days | ID_6 | left__front__average | -3.403529073 |
| 21days | ID_7 | left__front__average | 0.910377542 |
| 21days | ID_8 | left__front__average | -0.930189315 |
| 21days | ID_9 | left__front__average | -1.665121787 |
| 21days | ID_10 | left__front__average | 0.083718823 |
| 0days | ID_1 | left__front__median | -0.850321644 |
| 0days | ID_2 | left__front__median | 0 |
| 0days | ID_3 | left__front__median | 0.281198903 |
| 0days | ID_4 | left__front__median | 0.15545647 |
| 0days | ID_5 | left__front__median | -1.272624197 |
| 0days | ID_6 | left__front__median | -0.915424142 |
| 0days | ID_7 | left__front__median | 0.927347784 |
| 0days | ID_8 | left__front__median | 1.163047102 |
| 0days | ID_9 | left__front__median | -1.034528229 |
| 0days | ID_10 | left__front__median | -0.029419574 |
| 0days | ID_11 | left__front__median | -4.107992567 |
| 0days | ID_12 | left__front__median | 0.239413114 |
| 0days | ID_13 | left__front__median | 2.592267089 |
| 3days | ID_1 | left__front__median | -0.958318096 |
| 3days | ID_2 | left__front__median | -1.267064368 |
| 3days | ID_3 | left__front__median | -1.021439728 |
| 3days | ID_4 | left__front__median | -2.699232424 |
| 3days | ID_5 | left__front__median | -2.323193601 |
| 3days | ID_6 | left__front__median | -1.015288122 |
| 3days | ID_7 | left__front__median | 0.995621417 |
| 3days | ID_8 | left__front__median | -0.428999883 |
| 7days | ID_1 | left__front__median | -0.332171623 |
| 7days | ID_2 | left__front__median | -0.318934555 |
| 7days | ID_3 | left__front__median | -0.968282335 |
| 7days | ID_4 | left__front__median | -0.770697385 |
| 7days | ID_5 | left__front__median | 0.724418545 |
| 7days | ID_6 | left__front__median | -3.629760179 |
| 7days | ID_7 | left__front__median | -1.833860047 |
| 7days | ID_8 | left__front__median | 0.629543459 |
| 7days | ID_9 | left__front__median | 1.712253759 |
| 14days | ID_1 | left__front__median | 2.191838668 |

| | | | |
|---|---|---|---|
| 14days | ID_2 | left__front__median | -0.120205915 |
| 14days | ID_3 | left__front__median | -0.697275341 |
| 14days | ID_4 | left__front__median | -1.74039099 |
| 14days | ID_5 | left__front__median | 0.386450839 |
| 14days | ID_6 | left__front__median | -1.765467357 |
| 14days | ID_7 | left__front__median | -1.946032421 |
| 14days | ID_8 | left__front__median | -1.83085407 |
| 14days | ID_9 | left__front__median | 0.234359659 |
| 14days | ID_10 | left__front__median | 0.771166045 |
| 14days | ID_11 | left__front__median | 0.782629212 |
| 21days | ID_1 | left__front__median | -0.75497561 |
| 21days | ID_2 | left__front__median | -1.1884572 |
| 21days | ID_3 | left__front__median | -1.419486097 |
| 21days | ID_4 | left__front__median | -0.349997282 |
| 21days | ID_5 | left__front__median | -0.828709842 |
| 21days | ID_6 | left__front__median | -4.917535076 |
| 21days | ID_7 | left__front__median | 2.072451757 |
| 21days | ID_8 | left__front__median | -0.071091841 |
| 21days | ID_9 | left__front__median | -2.686558747 |
| 21days | ID_10 | left__front__median | 0.053536251 |
| 0days | ID_1 | left__front__protraction | -0.431356953 |
| 0days | ID_2 | left__front__protraction | -2.219015636 |
| 0days | ID_3 | left__front__protraction | -0.255317951 |
| 0days | ID_4 | left__front__protraction | 0 |
| 0days | ID_5 | left__front__protraction | 0.094228582 |
| 0days | ID_6 | left__front__protraction | 1.451605376 |
| 0days | ID_7 | left__front__protraction | 1.151527795 |
| 0days | ID_8 | left__front__protraction | -1.429568036 |
| 0days | ID_9 | left__front__protraction | 1.495405382 |
| 0days | ID_10 | left__front__protraction | -0.122580362 |
| 0days | ID_11 | left__front__protraction | -1.54287188 |
| 0days | ID_12 | left__front__protraction | 0.769433412 |
| 0days | ID_13 | left__front__protraction | 1.832993513 |
| 3days | ID_1 | left__front__protraction | 0.909095638 |
| 3days | ID_2 | left__front__protraction | 1.372744342 |
| 3days | ID_3 | left__front__protraction | 1.438993147 |
| 3days | ID_4 | left__front__protraction | -2.392774837 |
| 3days | ID_5 | left__front__protraction | -1.703263852 |
| 3days | ID_6 | left__front__protraction | -0.216873025 |
| 3days | ID_7 | left__front__protraction | 0.550482385 |
| 3days | ID_8 | left__front__protraction | 0.289721795 |
| 7days | ID_1 | left__front__protraction | 0.172296956 |
| 7days | ID_2 | left__front__protraction | -1.681786091 |
| 7days | ID_3 | left__front__protraction | 0.198160508 |
| 7days | ID_4 | left__front__protraction | -0.937126705 |
| 7days | ID_5 | left__front__protraction | -0.315310416 |
| 7days | ID_6 | left__front__protraction | -1.431788271 |
| 7days | ID_7 | left__front__protraction | 1.069687936 |

| | | | |
|---|---|---|---|
| 7days | ID_8 | left__front__protraction | 0.72263386 |
| 7days | ID_9 | left__front__protraction | -0.315511619 |
| 14days | ID_1 | left__front__protraction | 0.154797216 |
| 14days | ID_2 | left__front__protraction | 1.708096434 |
| 14days | ID_3 | left__front__protraction | -2.185781336 |
| 14days | ID_4 | left__front__protraction | 0.065733988 |
| 14days | ID_5 | left__front__protraction | 0.617927104 |
| 14days | ID_6 | left__front__protraction | 0.325474586 |
| 14days | ID_7 | left__front__protraction | 0.869700545 |
| 14days | ID_8 | left__front__protraction | 1.07357953 |
| 14days | ID_9 | left__front__protraction | 0.432166648 |
| 14days | ID_10 | left__front__protraction | 1.361641015 |
| 14days | ID_11 | left__front__protraction | 1.624899738 |
| 21days | ID_1 | left__front__protraction | -0.984004383 |
| 21days | ID_2 | left__front__protraction | 1.04709775 |
| 21days | ID_3 | left__front__protraction | 1.55130306 |
| 21days | ID_4 | left__front__protraction | 1.78538358 |
| 21days | ID_5 | left__front__protraction | 1.463509028 |
| 21days | ID_6 | left__front__protraction | -1.253657421 |
| 21days | ID_7 | left__front__protraction | 1.223119841 |
| 21days | ID_8 | left__front__protraction | 0.735382676 |
| 21days | ID_9 | left__front__protraction | 0.537097742 |
| 21days | ID_10 | left__front__protraction | 1.588416205 |
| 0days | ID_1 | left__front__retraction | -0.449184075 |
| 0days | ID_2 | left__front__retraction | 0.032254181 |
| 0days | ID_3 | left__front__retraction | 0.243039985 |
| 0days | ID_4 | left__front__retraction | 0.394707929 |
| 0days | ID_5 | left__front__retraction | -3.266687878 |
| 0days | ID_6 | left__front__retraction | -2.29195909 |
| 0days | ID_7 | left__front__retraction | 0 |
| 0days | ID_8 | left__front__retraction | 2.031415473 |
| 0days | ID_9 | left__front__retraction | -0.332777796 |
| 0days | ID_10 | left__front__retraction | 0.089530468 |
| 0days | ID_11 | left__front__retraction | -4.599697686 |
| 0days | ID_12 | left__front__retraction | -1.017194709 |
| 0days | ID_13 | left__front__retraction | 0.094253216 |
| 3days | ID_1 | left__front__retraction | -3.503634286 |
| 3days | ID_2 | left__front__retraction | -1.644096679 |
| 3days | ID_3 | left__front__retraction | 0.518083081 |
| 3days | ID_4 | left__front__retraction | -3.413815571 |
| 3days | ID_5 | left__front__retraction | -2.765435548 |
| 3days | ID_6 | left__front__retraction | -1.516642635 |
| 3days | ID_7 | left__front__retraction | -0.328488286 |
| 3days | ID_8 | left__front__retraction | -1.219683644 |
| 7days | ID_1 | left__front__retraction | -0.311802376 |
| 7days | ID_2 | left__front__retraction | -0.796664338 |
| 7days | ID_3 | left__front__retraction | -2.146841945 |
| 7days | ID_4 | left__front__retraction | -2.380134239 |

| | | | |
|---|---|---|---|
| 7days | ID_5 | left__front__retraction | -0.074984075 |
| 7days | ID_6 | left__front__retraction | -4.494462827 |
| 7days | ID_7 | left__front__retraction | -5.45622217 |
| 7days | ID_8 | left__front__retraction | -0.965980117 |
| 7days | ID_9 | left__front__retraction | -2.086637506 |
| 14days | ID_1 | left__front__retraction | 2.140659653 |
| 14days | ID_2 | left__front__retraction | -1.73669188 |
| 14days | ID_3 | left__front__retraction | -0.430918492 |
| 14days | ID_4 | left__front__retraction | -1.647172503 |
| 14days | ID_5 | left__front__retraction | -2.652616405 |
| 14days | ID_6 | left__front__retraction | -2.955103925 |
| 14days | ID_7 | left__front__retraction | -2.613589569 |
| 14days | ID_8 | left__front__retraction | -4.162630955 |
| 14days | ID_9 | left__front__retraction | -0.28831661 |
| 14days | ID_10 | left__front__retraction | 0.024024928 |
| 14days | ID_11 | left__front__retraction | -1.671537507 |
| 21days | ID_1 | left__front__retraction | -0.2012184 |
| 21days | ID_2 | left__front__retraction | -2.696259895 |
| 21days | ID_3 | left__front__retraction | -5.184760652 |
| 21days | ID_4 | left__front__retraction | -0.499028704 |
| 21days | ID_5 | left__front__retraction | -0.968420668 |
| 21days | ID_6 | left__front__retraction | -3.540820983 |
| 21days | ID_7 | left__front__retraction | -2.347366179 |
| 21days | ID_8 | left__front__retraction | -2.202341262 |
| 21days | ID_9 | left__front__retraction | -1.636486244 |
| 21days | ID_10 | left__front__retraction | -1.354557853 |
| 0days | ID_1 | right__back__average | -1.220707321 |
| 0days | ID_2 | right__back__average | 0 |
| 0days | ID_3 | right__back__average | -2.597076385 |
| 0days | ID_4 | right__back__average | -0.108458564 |
| 0days | ID_5 | right__back__average | -1.584014298 |
| 0days | ID_6 | right__back__average | -0.16124635 |
| 0days | ID_7 | right__back__average | 2.241302811 |
| 0days | ID_8 | right__back__average | 5.920478421 |
| 0days | ID_9 | right__back__average | 0.89439213 |
| 0days | ID_10 | right__back__average | 2.723608993 |
| 0days | ID_11 | right__back__average | -2.645229343 |
| 0days | ID_12 | right__back__average | 1.305643589 |
| 0days | ID_13 | right__back__average | 0.691280553 |
| 3days | ID_1 | right__back__average | -4.532717748 |
| 3days | ID_2 | right__back__average | -3.159389207 |
| 3days | ID_3 | right__back__average | -3.361433109 |
| 3days | ID_4 | right__back__average | -2.602608926 |
| 3days | ID_5 | right__back__average | -1.691631756 |
| 3days | ID_6 | right__back__average | -1.028984944 |
| 3days | ID_7 | right__back__average | -0.954060279 |
| 3days | ID_8 | right__back__average | 1.085135535 |
| 7days | ID_1 | right__back__average | -0.944070368 |

| | | | |
|---|---|---|---|
| 7days | ID_2 | right__back__average | 0.427358733 |
| 7days | ID_3 | right__back__average | -0.487637607 |
| 7days | ID_4 | right__back__average | 2.39815611 |
| 7days | ID_5 | right__back__average | 0.701435722 |
| 7days | ID_6 | right__back__average | 2.165795863 |
| 7days | ID_7 | right__back__average | -2.410576257 |
| 7days | ID_8 | right__back__average | 0.774297871 |
| 7days | ID_9 | right__back__average | -0.245413339 |
| 14days | ID_1 | right__back__average | -1.060872574 |
| 14days | ID_2 | right__back__average | -2.184781052 |
| 14days | ID_3 | right__back__average | 1.943203888 |
| 14days | ID_4 | right__back__average | 2.568507429 |
| 14days | ID_5 | right__back__average | 0.612213986 |
| 14days | ID_6 | right__back__average | -3.290772364 |
| 14days | ID_7 | right__back__average | -0.006108082 |
| 14days | ID_8 | right__back__average | -0.037379197 |
| 14days | ID_9 | right__back__average | 1.848193881 |
| 14days | ID_10 | right__back__average | 2.666578077 |
| 14days | ID_11 | right__back__average | 2.549472541 |
| 21days | ID_1 | right__back__average | -0.78674041 |
| 21days | ID_2 | right__back__average | 0.931724259 |
| 21days | ID_3 | right__back__average | 0.975036331 |
| 21days | ID_4 | right__back__average | 2.303445472 |
| 21days | ID_5 | right__back__average | 1.324666077 |
| 21days | ID_6 | right__back__average | 2.533424963 |
| 21days | ID_7 | right__back__average | 3.006823535 |
| 21days | ID_8 | right__back__average | 0.58944103 |
| 21days | ID_9 | right__back__average | 1.727858666 |
| 21days | ID_10 | right__back__average | 1.771106012 |
| 0days | ID_1 | right__back__median | -1.830371539 |
| 0days | ID_2 | right__back__median | -1.312863594 |
| 0days | ID_3 | right__back__median | -3.319817877 |
| 0days | ID_4 | right__back__median | -0.469532213 |
| 0days | ID_5 | right__back__median | -1.245295153 |
| 0days | ID_6 | right__back__median | 0 |
| 0days | ID_7 | right__back__median | 2.680109103 |
| 0days | ID_8 | right__back__median | 6.497710752 |
| 0days | ID_9 | right__back__median | 0.790934626 |
| 0days | ID_10 | right__back__median | 2.968405315 |
| 0days | ID_11 | right__back__median | -3.894430257 |
| 0days | ID_12 | right__back__median | 0.788309133 |
| 0days | ID_13 | right__back__median | 0.520071148 |
| 3days | ID_1 | right__back__median | -6.962627438 |
| 3days | ID_2 | right__back__median | -4.178970217 |
| 3days | ID_3 | right__back__median | -4.405582381 |
| 3days | ID_4 | right__back__median | -3.12863867 |
| 3days | ID_5 | right__back__median | -3.626008271 |
| 3days | ID_6 | right__back__median | -2.320210661 |

| | | | |
|---|---|---|---|
| 3days | ID_7 | right__back__median | -0.367657838 |
| 3days | ID_8 | right__back__median | 2.149825638 |
| 7days | ID_1 | right__back__median | -0.520040288 |
| 7days | ID_2 | right__back__median | 0.56389815 |
| 7days | ID_3 | right__back__median | -0.903462467 |
| 7days | ID_4 | right__back__median | 2.806679172 |
| 7days | ID_5 | right__back__median | 0.640917201 |
| 7days | ID_6 | right__back__median | 4.375360868 |
| 7days | ID_7 | right__back__median | 0.258314772 |
| 7days | ID_8 | right__back__median | 1.74841411 |
| 7days | ID_9 | right__back__median | 1.949187482 |
| 14days | ID_1 | right__back__median | -1.336873786 |
| 14days | ID_2 | right__back__median | -3.613745458 |
| 14days | ID_3 | right__back__median | 2.698669552 |
| 14days | ID_4 | right__back__median | 3.631003844 |
| 14days | ID_5 | right__back__median | 2.291049403 |
| 14days | ID_6 | right__back__median | -4.093526801 |
| 14days | ID_7 | right__back__median | 2.034673739 |
| 14days | ID_8 | right__back__median | 1.972429748 |
| 14days | ID_9 | right__back__median | 2.663750211 |
| 14days | ID_10 | right__back__median | 4.397476978 |
| 14days | ID_11 | right__back__median | 2.572437652 |
| 21days | ID_1 | right__back__median | -0.792845149 |
| 21days | ID_2 | right__back__median | 0.734144003 |
| 21days | ID_3 | right__back__median | 0.862819433 |
| 21days | ID_4 | right__back__median | 4.263139462 |
| 21days | ID_5 | right__back__median | 2.315868584 |
| 21days | ID_6 | right__back__median | 3.80544907 |
| 21days | ID_7 | right__back__median | 3.04927329 |
| 21days | ID_8 | right__back__median | 2.234963897 |
| 21days | ID_9 | right__back__median | 3.362991192 |
| 21days | ID_10 | right__back__median | 1.786609545 |
| 0days | ID_1 | right__back__protraction | -1.421076867 |
| 0days | ID_2 | right__back__protraction | -0.500102876 |
| 0days | ID_3 | right__back__protraction | -2.286100223 |
| 0days | ID_4 | right__back__protraction | 0 |
| 0days | ID_5 | right__back__protraction | -0.220407759 |
| 0days | ID_6 | right__back__protraction | 0.881491698 |
| 0days | ID_7 | right__back__protraction | 2.694013764 |
| 0days | ID_8 | right__back__protraction | 2.882976574 |
| 0days | ID_9 | right__back__protraction | -0.369787074 |
| 0days | ID_10 | right__back__protraction | 3.616231241 |
| 0days | ID_11 | right__back__protraction | -0.380579943 |
| 0days | ID_12 | right__back__protraction | 1.778413854 |
| 0days | ID_13 | right__back__protraction | 1.553195225 |
| 3days | ID_1 | right__back__protraction | -2.588404577 |
| 3days | ID_2 | right__back__protraction | -3.125251707 |
| 3days | ID_3 | right__back__protraction | -4.376133131 |

| | | | |
|---|---|---|---|
| 3days | ID_4 | right__back__protraction | -2.631863487 |
| 3days | ID_5 | right__back__protraction | 0.606737611 |
| 3days | ID_6 | right__back__protraction | -0.779698286 |
| 3days | ID_7 | right__back__protraction | 0.891683823 |
| 3days | ID_8 | right__back__protraction | 2.392053499 |
| 7days | ID_1 | right__back__protraction | -1.466488182 |
| 7days | ID_2 | right__back__protraction | 0.32045122 |
| 7days | ID_3 | right__back__protraction | -0.011322923 |
| 7days | ID_4 | right__back__protraction | 2.701702383 |
| 7days | ID_5 | right__back__protraction | 1.246791091 |
| 7days | ID_6 | right__back__protraction | 3.823594072 |
| 7days | ID_7 | right__back__protraction | -0.242853724 |
| 7days | ID_8 | right__back__protraction | 1.711197726 |
| 7days | ID_9 | right__back__protraction | 0.846442875 |
| 14days | ID_1 | right__back__protraction | -1.938068344 |
| 14days | ID_2 | right__back__protraction | -1.865193 |
| 14days | ID_3 | right__back__protraction | 1.435711494 |
| 14days | ID_4 | right__back__protraction | 2.083280655 |
| 14days | ID_5 | right__back__protraction | 1.251088937 |
| 14days | ID_6 | right__back__protraction | -0.011532146 |
| 14days | ID_7 | right__back__protraction | 0.883791713 |
| 14days | ID_8 | right__back__protraction | 1.507829887 |
| 14days | ID_9 | right__back__protraction | 3.836127713 |
| 14days | ID_10 | right__back__protraction | 2.288746274 |
| 14days | ID_11 | right__back__protraction | 1.804490211 |
| 21days | ID_1 | right__back__protraction | 0.275301351 |
| 21days | ID_2 | right__back__protraction | 2.943302486 |
| 21days | ID_3 | right__back__protraction | 3.205936847 |
| 21days | ID_4 | right__back__protraction | 3.620898837 |
| 21days | ID_5 | right__back__protraction | 3.365270743 |
| 21days | ID_6 | right__back__protraction | 2.508306541 |
| 21days | ID_7 | right__back__protraction | 3.022708311 |
| 21days | ID_8 | right__back__protraction | 2.153404345 |
| 21days | ID_9 | right__back__protraction | 1.172660773 |
| 21days | ID_10 | right__back__protraction | 2.371177014 |
| 0days | ID_1 | right__back__retraction | 0 |
| 0days | ID_2 | right__back__retraction | 1.416050016 |
| 0days | ID_3 | right__back__retraction | -1.957660531 |
| 0days | ID_4 | right__back__retraction | -1.984040499 |
| 0days | ID_5 | right__back__retraction | -3.312804411 |
| 0days | ID_6 | right__back__retraction | -1.960314377 |
| 0days | ID_7 | right__back__retraction | 1.678681237 |
| 0days | ID_8 | right__back__retraction | 7.073698638 |
| 0days | ID_9 | right__back__retraction | 0.849269889 |
| 0days | ID_10 | right__back__retraction | 0.812162463 |
| 0days | ID_11 | right__back__retraction | -2.742840295 |
| 0days | ID_12 | right__back__retraction | 1.100014994 |
| 0days | ID_13 | right__back__retraction | -0.055768991 |

| | | | |
|---|---|---|---|
| 3days | ID_1 | right__back__retraction | -3.686972257 |
| 3days | ID_2 | right__back__retraction | -2.869935908 |
| 3days | ID_3 | right__back__retraction | -0.972749326 |
| 3days | ID_4 | right__back__retraction | -2.255567001 |
| 3days | ID_5 | right__back__retraction | -2.232078302 |
| 3days | ID_6 | right__back__retraction | -0.681708899 |
| 3days | ID_7 | right__back__retraction | -5.12461956 |
| 3days | ID_8 | right__back__retraction | -1.350775104 |
| 7days | ID_1 | right__back__retraction | -0.58724218 |
| 7days | ID_2 | right__back__retraction | 0.787624248 |
| 7days | ID_3 | right__back__retraction | -0.183581407 |
| 7days | ID_4 | right__back__retraction | 1.530287351 |
| 7days | ID_5 | right__back__retraction | -0.393741324 |
| 7days | ID_6 | right__back__retraction | -2.962342971 |
| 7days | ID_7 | right__back__retraction | -5.638722507 |
| 7days | ID_8 | right__back__retraction | -2.386434933 |
| 7days | ID_9 | right__back__retraction | -5.308127703 |
| 14days | ID_1 | right__back__retraction | -0.207466585 |
| 14days | ID_2 | right__back__retraction | -2.561717823 |
| 14days | ID_3 | right__back__retraction | 1.112462825 |
| 14days | ID_4 | right__back__retraction | 1.014349941 |
| 14days | ID_5 | right__back__retraction | -3.386940034 |
| 14days | ID_6 | right__back__retraction | -6.122774698 |
| 14days | ID_7 | right__back__retraction | -4.125879163 |
| 14days | ID_8 | right__back__retraction | -3.498621284 |
| 14days | ID_9 | right__back__retraction | -1.131703461 |
| 14days | ID_10 | right__back__retraction | -0.959433361 |
| 14days | ID_11 | right__back__retraction | 1.274775143 |
| 21days | ID_1 | right__back__retraction | 0.054960709 |
| 21days | ID_2 | right__back__retraction | -0.958421751 |
| 21days | ID_3 | right__back__retraction | -1.723538194 |
| 21days | ID_4 | right__back__retraction | -0.415421186 |
| 21days | ID_5 | right__back__retraction | -1.936323889 |
| 21days | ID_6 | right__back__retraction | -0.430320046 |
| 21days | ID_7 | right__back__retraction | 1.544883197 |
| 21days | ID_8 | right__back__retraction | -2.939745193 |
| 21days | ID_9 | right__back__retraction | -1.126855587 |
| 21days | ID_10 | right__back__retraction | 0.324875024 |
| 0days | ID_1 | right__front__average | -0.414956268 |
| 0days | ID_2 | right__front__average | -0.129620843 |
| 0days | ID_3 | right__front__average | 0.021914097 |
| 0days | ID_4 | right__front__average | 0.504541113 |
| 0days | ID_5 | right__front__average | -1.859685875 |
| 0days | ID_6 | right__front__average | -0.652199165 |
| 0days | ID_7 | right__front__average | 1.289240568 |
| 0days | ID_8 | right__front__average | 1.10557979 |
| 0days | ID_9 | right__front__average | 0.415070213 |
| 0days | ID_10 | right__front__average | -3.017275615 |

| | | | |
|---|---|---|---|
| 0days | ID_11 | right__front__average | -0.70525656 |
| 0days | ID_12 | right__front__average | 0 |
| 0days | ID_13 | right__front__average | 0.936034643 |
| 3days | ID_1 | right__front__average | -1.847480913 |
| 3days | ID_2 | right__front__average | -1.006830148 |
| 3days | ID_3 | right__front__average | -0.519916629 |
| 3days | ID_4 | right__front__average | -2.375509092 |
| 3days | ID_5 | right__front__average | -1.133696988 |
| 3days | ID_6 | right__front__average | -1.479573856 |
| 3days | ID_7 | right__front__average | -1.795672612 |
| 3days | ID_8 | right__front__average | 0.225308201 |
| 7days | ID_1 | right__front__average | 0.03472778 |
| 7days | ID_2 | right__front__average | 0.042579331 |
| 7days | ID_3 | right__front__average | -2.374681633 |
| 7days | ID_4 | right__front__average | -0.817392494 |
| 7days | ID_5 | right__front__average | -2.633270102 |
| 7days | ID_6 | right__front__average | -1.599808714 |
| 7days | ID_7 | right__front__average | -1.862385246 |
| 7days | ID_8 | right__front__average | -2.367818189 |
| 7days | ID_9 | right__front__average | -1.35271496 |
| 14days | ID_1 | right__front__average | 1.3860377 |
| 14days | ID_2 | right__front__average | -2.395939268 |
| 14days | ID_3 | right__front__average | 0.305387048 |
| 14days | ID_4 | right__front__average | -1.726435765 |
| 14days | ID_5 | right__front__average | -2.023496358 |
| 14days | ID_6 | right__front__average | -3.476992686 |
| 14days | ID_7 | right__front__average | -1.318233544 |
| 14days | ID_8 | right__front__average | -1.365925182 |
| 14days | ID_9 | right__front__average | -0.455525058 |
| 14days | ID_10 | right__front__average | -0.109342186 |
| 14days | ID_11 | right__front__average | 0.568691943 |
| 21days | ID_1 | right__front__average | -0.16368815 |
| 21days | ID_2 | right__front__average | -3.077911527 |
| 21days | ID_3 | right__front__average | -3.423564686 |
| 21days | ID_4 | right__front__average | 0.697291126 |
| 21days | ID_5 | right__front__average | -3.322927206 |
| 21days | ID_6 | right__front__average | -0.251843726 |
| 21days | ID_7 | right__front__average | -1.335368763 |
| 21days | ID_8 | right__front__average | 0.031217492 |
| 21days | ID_9 | right__front__average | -0.198973876 |
| 21days | ID_10 | right__front__average | -0.798304955 |
| 0days | ID_1 | right__front__median | -0.168571973 |
| 0days | ID_2 | right__front__median | -0.826783694 |
| 0days | ID_3 | right__front__median | -0.385938075 |
| 0days | ID_4 | right__front__median | 0.431279483 |
| 0days | ID_5 | right__front__median | -1.101756862 |
| 0days | ID_6 | right__front__median | 0.5611322 |
| 0days | ID_7 | right__front__median | 1.315754519 |

| | | | |
|---|---|---|---|
| 0days | ID_8 | right__front__median | 0.366439149 |
| 0days | ID_9 | right__front__median | 1.503179119 |
| 0days | ID_10 | right__front__median | -2.93875279 |
| 0days | ID_11 | right__front__median | -1.669213039 |
| 0days | ID_12 | right__front__median | 0 |
| 0days | ID_13 | right__front__median | 1.570389115 |
| 3days | ID_1 | right__front__median | -2.416762596 |
| 3days | ID_2 | right__front__median | -1.391173985 |
| 3days | ID_3 | right__front__median | -0.744887037 |
| 3days | ID_4 | right__front__median | -3.806978384 |
| 3days | ID_5 | right__front__median | -2.725821269 |
| 3days | ID_6 | right__front__median | -3.085073207 |
| 3days | ID_7 | right__front__median | -1.90808633 |
| 3days | ID_8 | right__front__median | 1.319839772 |
| 7days | ID_1 | right__front__median | 0.017742105 |
| 7days | ID_2 | right__front__median | -0.909505354 |
| 7days | ID_3 | right__front__median | -3.417498827 |
| 7days | ID_4 | right__front__median | -1.868867582 |
| 7days | ID_5 | right__front__median | -4.705078082 |
| 7days | ID_6 | right__front__median | -3.174683055 |
| 7days | ID_7 | right__front__median | -2.608414205 |
| 7days | ID_8 | right__front__median | -3.375109376 |
| 7days | ID_9 | right__front__median | -0.989122911 |
| 14days | ID_1 | right__front__median | 1.42538104 |
| 14days | ID_2 | right__front__median | -4.247891766 |
| 14days | ID_3 | right__front__median | -0.431757111 |
| 14days | ID_4 | right__front__median | -2.746476822 |
| 14days | ID_5 | right__front__median | -2.517615182 |
| 14days | ID_6 | right__front__median | -4.421366903 |
| 14days | ID_7 | right__front__median | -2.217291703 |
| 14days | ID_8 | right__front__median | -2.385538486 |
| 14days | ID_9 | right__front__median | -0.035296915 |
| 14days | ID_10 | right__front__median | 0.673202659 |
| 14days | ID_11 | right__front__median | -0.211709649 |
| 21days | ID_1 | right__front__median | -0.234139268 |
| 21days | ID_2 | right__front__median | -4.916247242 |
| 21days | ID_3 | right__front__median | -4.778671159 |
| 21days | ID_4 | right__front__median | 0.80778347 |
| 21days | ID_5 | right__front__median | -5.618115113 |
| 21days | ID_6 | right__front__median | -0.706095552 |
| 21days | ID_7 | right__front__median | -1.9772455 |
| 21days | ID_8 | right__front__median | 0.147007758 |
| 21days | ID_9 | right__front__median | 0.13535427 |
| 21days | ID_10 | right__front__median | -1.132010508 |
| 0days | ID_1 | right__front__protraction | -1.512247084 |
| 0days | ID_2 | right__front__protraction | 0 |
| 0days | ID_3 | right__front__protraction | -0.329057487 |
| 0days | ID_4 | right__front__protraction | 0.189957542 |

| | | | |
|---|---|---|---|
| 0days | ID_5 | right__front__protraction | -1.519248755 |
| 0days | ID_6 | right__front__protraction | -0.757627436 |
| 0days | ID_7 | right__front__protraction | 1.766036297 |
| 0days | ID_8 | right__front__protraction | 0.089914395 |
| 0days | ID_9 | right__front__protraction | -0.782273392 |
| 0days | ID_10 | right__front__protraction | -1.977281229 |
| 0days | ID_11 | right__front__protraction | 0.124241544 |
| 0days | ID_12 | right__front__protraction | 0.812229566 |
| 0days | ID_13 | right__front__protraction | 1.385868305 |
| 3days | ID_1 | right__front__protraction | -0.012704658 |
| 3days | ID_2 | right__front__protraction | -0.154354055 |
| 3days | ID_3 | right__front__protraction | -0.407589008 |
| 3days | ID_4 | right__front__protraction | -0.755500501 |
| 3days | ID_5 | right__front__protraction | 0.164153271 |
| 3days | ID_6 | right__front__protraction | -0.439168329 |
| 3days | ID_7 | right__front__protraction | -0.94982165 |
| 3days | ID_8 | right__front__protraction | -1.08244552 |
| 7days | ID_1 | right__front__protraction | -0.976820236 |
| 7days | ID_2 | right__front__protraction | 0.183013131 |
| 7days | ID_3 | right__front__protraction | -1.375000616 |
| 7days | ID_4 | right__front__protraction | 0.225423235 |
| 7days | ID_5 | right__front__protraction | -1.557079569 |
| 7days | ID_6 | right__front__protraction | 0.996728966 |
| 7days | ID_7 | right__front__protraction | -1.045804756 |
| 7days | ID_8 | right__front__protraction | 0.608472236 |
| 7days | ID_9 | right__front__protraction | -1.928753953 |
| 14days | ID_1 | right__front__protraction | -0.236254125 |
| 14days | ID_2 | right__front__protraction | -1.52548254 |
| 14days | ID_3 | right__front__protraction | 0.279243327 |
| 14days | ID_4 | right__front__protraction | 0.412871035 |
| 14days | ID_5 | right__front__protraction | -0.935036379 |
| 14days | ID_6 | right__front__protraction | -1.142357874 |
| 14days | ID_7 | right__front__protraction | 1.62375621 |
| 14days | ID_8 | right__front__protraction | 0.960321015 |
| 14days | ID_9 | right__front__protraction | -1.384088641 |
| 14days | ID_10 | right__front__protraction | 1.544552256 |
| 14days | ID_11 | right__front__protraction | 2.948871686 |
| 21days | ID_1 | right__front__protraction | -0.4977671 |
| 21days | ID_2 | right__front__protraction | -1.257713203 |
| 21days | ID_3 | right__front__protraction | -0.595169044 |
| 21days | ID_4 | right__front__protraction | 1.832973045 |
| 21days | ID_5 | right__front__protraction | -0.476463466 |
| 21days | ID_6 | right__front__protraction | 0.927586227 |
| 21days | ID_7 | right__front__protraction | -0.590027022 |
| 21days | ID_8 | right__front__protraction | 1.68678415 |
| 21days | ID_9 | right__front__protraction | 0.159135913 |
| 21days | ID_10 | right__front__protraction | 1.020922517 |
| 0days | ID_1 | right__front__retraction | -0.93066662 |

| | | | |
|---|---|---|---|
| 0days | ID_2 | right__front__retraction | 0.44705624 |
| 0days | ID_3 | right__front__retraction | -0.60513673 |
| 0days | ID_4 | right__front__retraction | 0.58771033 |
| 0days | ID_5 | right__front__retraction | -3.259730399 |
| 0days | ID_6 | right__front__retraction | -2.044834264 |
| 0days | ID_7 | right__front__retraction | 1.03871289 |
| 0days | ID_8 | right__front__retraction | 1.992417761 |
| 0days | ID_9 | right__front__retraction | 0 |
| 0days | ID_10 | right__front__retraction | -2.937163791 |
| 0days | ID_11 | right__front__retraction | 0.203480725 |
| 0days | ID_12 | right__front__retraction | -0.215854572 |
| 0days | ID_13 | right__front__retraction | 0.294718757 |
| 3days | ID_1 | right__front__retraction | -2.038213048 |
| 3days | ID_2 | right__front__retraction | -0.407148309 |
| 3days | ID_3 | right__front__retraction | 0.750361345 |
| 3days | ID_4 | right__front__retraction | -2.18971475 |
| 3days | ID_5 | right__front__retraction | -1.678337059 |
| 3days | ID_6 | right__front__retraction | -0.37815693 |
| 3days | ID_7 | right__front__retraction | -2.078306624 |
| 3days | ID_8 | right__front__retraction | 1.173070963 |
| 7days | ID_1 | right__front__retraction | 1.836241544 |
| 7days | ID_2 | right__front__retraction | 0.752301178 |
| 7days | ID_3 | right__front__retraction | -1.538627901 |
| 7days | ID_4 | right__front__retraction | -1.651799997 |
| 7days | ID_5 | right__front__retraction | -1.623444265 |
| 7days | ID_6 | right__front__retraction | -4.187294494 |
| 7days | ID_7 | right__front__retraction | -0.550974868 |
| 7days | ID_8 | right__front__retraction | -3.731654882 |
| 7days | ID_9 | right__front__retraction | -0.706718797 |
| 14days | ID_1 | right__front__retraction | 2.816010627 |
| 14days | ID_2 | right__front__retraction | -1.488736399 |
| 14days | ID_3 | right__front__retraction | 0.471879453 |
| 14days | ID_4 | right__front__retraction | -2.837391993 |
| 14days | ID_5 | right__front__retraction | -3.58770493 |
| 14days | ID_6 | right__front__retraction | -4.199274051 |
| 14days | ID_7 | right__front__retraction | -2.868433603 |
| 14days | ID_8 | right__front__retraction | -2.479819801 |
| 14days | ID_9 | right__front__retraction | 0.870701604 |
| 14days | ID_10 | right__front__retraction | -2.344768186 |
| 14days | ID_11 | right__front__retraction | -0.964389038 |
| 21days | ID_1 | right__front__retraction | 0.92750539 |
| 21days | ID_2 | right__front__retraction | -3.021517049 |
| 21days | ID_3 | right__front__retraction | -4.12117692 |
| 21days | ID_4 | right__front__retraction | -0.895782426 |
| 21days | ID_5 | right__front__retraction | -3.664846613 |
| 21days | ID_6 | right__front__retraction | -1.429468384 |
| 21days | ID_7 | right__front__retraction | -1.27661404 |
| 21days | ID_8 | right__front__retraction | -1.132579016 |

| | | | |
|---|---|---|---|
| 21days | ID_9 | right__front__retraction | -1.847145473 |
| 21days | ID_10 | right__front__retraction | -1.66798754 |

**Suppl. Table 23: Summary of side perspective horizontal analysis**

| day | Measure | Mean | SD |
|---|---|---|---|
| 0days | left__back__ | 0 | 2.354541209 |
| 0days | left__back__ | 0 | 3.227139569 |
| 0days | left__back__ | 0 | 1.69170068 |
| 0days | left__back__ | 0 | 2.846657111 |
| 0days | left__front__ | 0 | 1.253349506 |
| 0days | left__front__ | 0 | 1.570427129 |
| 0days | left__front__ | 0 | 1.264781286 |
| 0days | left__front__ | 0 | 1.743007692 |
| 0days | right__back_ | 0 | 2.344531176 |
| 0days | right__back_ | 0 | 2.77491306 |
| 0days | right__back_ | 0 | 1.770511725 |
| 0days | right__back_ | 0 | 2.700298056 |
| 0days | right__front_ | 0 | 1.203423662 |
| 0days | right__front_ | 0 | 1.305458505 |
| 0days | right__front_ | 0 | 1.12076202 |
| 0days | right__front_ | 0 | 1.5320941 |
| 3days | left__back__ | -1.822015073 | 1.929090297 |
| 3days | left__back__ | -4.080316287 | 3.126680302 |
| 3days | left__back__ | -1.255989136 | 1.71114298 |
| 3days | left__back__ | -1.564663883 | 1.511813249 |
| 3days | left__front__ | -0.561834002 | 1.144458372 |
| 3days | left__front__ | -1.018363925 | 1.131125044 |
| 3days | left__front__ | 0.42010209 | 1.405823096 |
| 3days | left__front__ | -1.580369657 | 1.433498932 |
| 3days | right__back_ | -2.147120341 | 1.757349752 |
| 3days | right__back_ | -3.377323471 | 2.757335366 |
| 3days | right__back_ | -1.684051431 | 2.344941218 |
| 3days | right__back_ | -2.243822651 | 1.485169903 |
| 3days | right__front_ | -1.306635422 | 0.824263826 |
| 3days | right__front_ | -2.162424463 | 1.600281066 |
| 3days | right__front_ | -0.423378668 | 0.447325136 |
| 3days | right__front_ | -1.042742684 | 1.33491689 |
| 7days | left__back__ | -2.317728529 | 0.643078789 |
| 7days | left__back__ | -3.443448227 | 1.062367573 |
| 7days | left__back__ | -2.07934681 | 0.764617288 |
| 7days | left__back__ | -2.368477429 | 2.152067839 |
| 7days | left__front__ | -1.002909718 | 1.17748234 |
| 7days | left__front__ | -0.332171623 | 1.564770365 |
| 7days | left__front__ | -0.315310416 | 0.934484209 |
| 7days | left__front__ | -2.086637506 | 1.84924638 |
| 7days | right__back_ | 0.427358733 | 1.503626524 |
| 7days | right__back_ | 0.640917201 | 1.677492813 |
| 7days | right__back_ | 0.846442875 | 1.603551536 |
| 7days | right__back_ | -0.58724218 | 2.560789847 |
| 7days | right__front_ | -1.599808714 | 1.008113666 |

| | | | |
|---|---|---|---|
| 7days | right__front_ | -2.608414205 | 1.510248011 |
| 7days | right__front_ | -0.976820236 | 1.054519984 |
| 7days | right__front_ | -1.538627901 | 1.919825549 |
| 14days | left__back__ | -1.453731628 | 2.656271657 |
| 14days | left__back__ | -1.659057825 | 3.988066508 |
| 14days | left__back__ | -1.607006233 | 1.527604957 |
| 14days | left__back__ | -1.44399568 | 2.82983814 |
| 14days | left__front__ | -0.247411429 | 1.125586933 |
| 14days | left__front__ | -0.120205915 | 1.368714832 |
| 14days | left__front__ | 0.617927104 | 1.07125209 |
| 14days | left__front__ | -1.671537507 | 1.733422223 |
| 14days | right__back_ | 0.612213986 | 2.040695014 |
| 14days | right__back_ | 2.291049403 | 2.872608367 |
| 14days | right__back_ | 1.435711494 | 1.729372995 |
| 14days | right__back_ | -1.131703461 | 2.439141909 |
| 14days | right__front_ | -1.318233544 | 1.44321152 |
| 14days | right__front_ | -2.217291703 | 1.953600325 |
| 14days | right__front_ | 0.279243327 | 1.441428049 |
| 14days | right__front_ | -2.344768186 | 2.130770982 |
| 21days | left__back__ | 0.116947838 | 1.433482162 |
| 21days | left__back__ | 0.387687533 | 1.950397104 |
| 21days | left__back__ | 0.55295167 | 0.542495908 |
| 21days | left__back__ | -0.318030745 | 2.525711836 |
| 21days | left__front__ | -0.700592604 | 1.220282587 |
| 21days | left__front__ | -0.791842726 | 1.835661276 |
| 21days | left__front__ | 1.135108796 | 1.069862785 |
| 21days | left__front__ | -1.919413753 | 1.499065916 |
| 21days | right__back_ | 1.526262371 | 1.093320778 |
| 21days | right__back_ | 2.275416241 | 1.562512757 |
| 21days | right__back_ | 2.725804514 | 1.043927813 |
| 21days | right__back_ | -0.694370898 | 1.274467572 |
| 21days | right__front_ | -0.52507434 | 1.536426947 |
| 21days | right__front_ | -0.91905303 | 2.394837165 |
| 21days | right__front_ | -0.158663776 | 1.075149523 |
| 21days | right__front_ | -1.548727962 | 1.471551016 |

**Suppl. Table 24: Statistical test of bside perspective horizontal analysis**

| Measur | group1 | group2 | p.adj | sig | Test* |
|---|---|---|---|---|---|
| right__ba | 0days | 3days | 0.001853114 | ** | Repeat ANOVA + PH test |
| left__bac | 0days | 3days | 0.008909729 | ** | Repeat ANOVA + PH test |
| right__ba | 0days | 3days | 0.016891748 | * | Repeat ANOVA + PH test |
| left__bac | 0days | 14days | 0.017744158 | * | Repeat ANOVA + PH test |
| right__ba | 0days | 3days | 0.017941213 | * | Repeat ANOVA + PH test |
| right__fr | 0days | 7days | 0.029066542 | * | Repeat ANOVA + PH test |
| left__bac | 0days | 7days | 0.034047886 | * | Repeat ANOVA + PH test |
| left__bac | 0days | 7days | 0.034277897 | * | Repeat ANOVA + PH test |
| right__ba | 0days | 21days | 0.040234352 | * | Repeat ANOVA + PH test |
| left__bac | 0days | 7days | 0.048678422 | * | Repeat ANOVA + PH test |
| left__bac | 0days | 14days | 0.057031105 | * | Repeat ANOVA + PH test |
| right__ba | 0days | 7days | 0.075050598 | * | Repeat ANOVA + PH test |
| right__ba | 0days | 14days | 0.075050598 | * | Repeat ANOVA + PH test |
| right__ba | 0days | 21days | 0.092891088 | * | Repeat ANOVA + PH test |
| right__fr | 0days | 3days | 0.097701518 | * | Repeat ANOVA + PH test |
| right__fr | 0days | 14days | 0.097701518 | * | Repeat ANOVA + PH test |
| right__fr | 0days | 21days | 0.097701518 | * | Repeat ANOVA + PH test |
| right__ba | 0days | 3days | 0.099814864 | * | Repeat ANOVA + PH test |
| left__bac | 0days | 3days | 0.148248421 | ns | Repeat ANOVA + PH test |
| left__bac | 0days | 14days | 0.148248421 | ns | Repeat ANOVA + PH test |
| right__ba | 0days | 21days | 0.302434764 | ns | Repeat ANOVA + PH test |
| right__fr | 0days | 21days | 0.334110592 | ns | Repeat ANOVA + PH test |
| left__fror | 0days | 7days | 0.361568073 | ns | Repeat ANOVA + PH test |
| left__fror | 0days | 21days | 0.361568073 | ns | Repeat ANOVA + PH test |
| left__bac | 0days | 21days | 0.367291339 | ns | Repeat ANOVA + PH test |
| right__ba | 0days | 7days | 0.375270127 | ns | Repeat ANOVA + PH test |
| right__ba | 0days | 14days | 0.375270127 | ns | Repeat ANOVA + PH test |
| left__bac | 0days | 7days | 0.406980531 | ns | Repeat ANOVA + PH test |
| left__bac | 0days | 21days | 0.406980531 | ns | Repeat ANOVA + PH test |
| left__fror | 0days | 3days | 0.457508352 | ns | Repeat ANOVA + PH test |
| left__fror | 0days | 14days | 0.457508352 | ns | Repeat ANOVA + PH test |
| right__fr | 0days | 14days | 0.492138568 | ns | Repeat ANOVA + PH test |
| left__bac | 0days | 3days | 0.526400626 | ns | Repeat ANOVA + PH test |
| left__bac | 0days | 14days | 0.526400626 | ns | Repeat ANOVA + PH test |
| left__bac | 0days | 3days | 0.536024904 | ns | Repeat ANOVA + PH test |
| right__fr | 0days | 7days | 0.53778735 | ns | Repeat ANOVA + PH test |
| left__bac | 0days | 21days | 0.549397538 | ns | Repeat ANOVA + PH test |
| left__bac | 0days | 21days | 0.574923688 | ns | Repeat ANOVA + PH test |
| right__fr | 0days | 3days | 0.613412544 | ns | Repeat ANOVA + PH test |
| right__fr | 0days | 7days | 0.613412544 | ns | Repeat ANOVA + PH test |
| right__ba | 0days | 21days | 0.620197338 | ns | Repeat ANOVA + PH test |
| right__fr | 0days | 3days | 0.655804103 | ns | Repeat ANOVA + PH test |
| right__fr | 0days | 14days | 0.655804103 | ns | Repeat ANOVA + PH test |
| right__fr | 0days | 21days | 0.655804103 | ns | Repeat ANOVA + PH test |
| left__fror | 0days | 3days | 1 | ns | Repeat ANOVA + PH test |

| | | | | | |
|---|---|---|---|---|---|
| left__fror | 0days | 7days | 1 | ns | Repeat ANOVA + PH test |
| left__fror | 0days | 14days | 1 | ns | Repeat ANOVA + PH test |
| left__fror | 0days | 21days | 1 | ns | Repeat ANOVA + PH test |
| left__fror | 0days | 3days | 1 | ns | Repeat ANOVA + PH test |
| left__fror | 0days | 7days | 1 | ns | Repeat ANOVA + PH test |
| left__fror | 0days | 14days | 1 | ns | Repeat ANOVA + PH test |
| left__fror | 0days | 21days | 1 | ns | Repeat ANOVA + PH test |
| left__fror | 0days | 3days | 1 | ns | Repeat ANOVA + PH test |
| left__fror | 0days | 7days | 1 | ns | Repeat ANOVA + PH test |
| left__fror | 0days | 14days | 1 | ns | Repeat ANOVA + PH test |
| left__fror | 0days | 21days | 1 | ns | Repeat ANOVA + PH test |
| right__ba | 0days | 7days | 1 | ns | Repeat ANOVA + PH test |
| right__ba | 0days | 14days | 1 | ns | Repeat ANOVA + PH test |
| right__ba | 0days | 7days | 1 | ns | Repeat ANOVA + PH test |
| right__ba | 0days | 14days | 1 | ns | Repeat ANOVA + PH test |
| right__fr | 0days | 3days | 1 | ns | Repeat ANOVA + PH test |
| right__fr | 0days | 7days | 1 | ns | Repeat ANOVA + PH test |
| right__fr | 0days | 14days | 1 | ns | Repeat ANOVA + PH test |
| right__fr | 0days | 21days | 1 | ns | Repeat ANOVA + PH test |

**Suppl. Table 25: Raw data side perspective horizontal angles analysis**

| day | ID | Measure | Angle, ° |
|---|---|---|---|
| 0days | ID_1 | avg_Angle__left__back__hip_ankle_toe | 114.1670397 |
| 0days | ID_2 | avg_Angle__left__back__hip_ankle_toe | 120.1753257 |
| 0days | ID_3 | avg_Angle__left__back__hip_ankle_toe | 102.0163458 |
| 0days | ID_4 | avg_Angle__left__back__hip_ankle_toe | 111.8714042 |
| 0days | ID_5 | avg_Angle__left__back__hip_ankle_toe | 109.0708804 |
| 0days | ID_6 | avg_Angle__left__back__hip_ankle_toe | 130.7947873 |
| 0days | ID_7 | avg_Angle__left__back__hip_ankle_toe | 117.6747812 |
| 0days | ID_8 | avg_Angle__left__back__hip_ankle_toe | 130.0793998 |
| 0days | ID_9 | avg_Angle__left__back__hip_ankle_toe | 116.1412727 |
| 0days | ID_10 | avg_Angle__left__back__hip_ankle_toe | 117.8603151 |
| 0days | ID_11 | avg_Angle__left__back__hip_ankle_toe | 128.3921617 |
| 0days | ID_12 | avg_Angle__left__back__hip_ankle_toe | 125.9447653 |
| 0days | ID_13 | avg_Angle__left__back__hip_ankle_toe | 124.2589827 |
| 3days | ID_1 | avg_Angle__left__back__hip_ankle_toe | 117.7780998 |
| 3days | ID_2 | avg_Angle__left__back__hip_ankle_toe | 109.0184176 |
| 3days | ID_3 | avg_Angle__left__back__hip_ankle_toe | 106.8257991 |
| 3days | ID_4 | avg_Angle__left__back__hip_ankle_toe | 118.9717829 |
| 3days | ID_5 | avg_Angle__left__back__hip_ankle_toe | 121.5482061 |
| 3days | ID_6 | avg_Angle__left__back__hip_ankle_toe | 120.6246118 |
| 3days | ID_7 | avg_Angle__left__back__hip_ankle_toe | 104.2305844 |
| 3days | ID_8 | avg_Angle__left__back__hip_ankle_toe | 127.5396905 |
| 7days | ID_1 | avg_Angle__left__back__hip_ankle_toe | 109.4231579 |
| 7days | ID_2 | avg_Angle__left__back__hip_ankle_toe | 106.9625165 |
| 7days | ID_3 | avg_Angle__left__back__hip_ankle_toe | 109.1608043 |
| 7days | ID_4 | avg_Angle__left__back__hip_ankle_toe | 122.5989211 |
| 7days | ID_5 | avg_Angle__left__back__hip_ankle_toe | 109.2314524 |
| 7days | ID_6 | avg_Angle__left__back__hip_ankle_toe | 122.5098544 |
| 7days | ID_7 | avg_Angle__left__back__hip_ankle_toe | 107.4999364 |
| 7days | ID_8 | avg_Angle__left__back__hip_ankle_toe | 118.1545642 |
| 7days | ID_9 | avg_Angle__left__back__hip_ankle_toe | 111.0962299 |
| 14days | ID_1 | avg_Angle__left__back__hip_ankle_toe | 105.6096997 |
| 14days | ID_2 | avg_Angle__left__back__hip_ankle_toe | 106.5926892 |
| 14days | ID_3 | avg_Angle__left__back__hip_ankle_toe | 116.7620652 |
| 14days | ID_4 | avg_Angle__left__back__hip_ankle_toe | 107.810918 |
| 14days | ID_5 | avg_Angle__left__back__hip_ankle_toe | 107.584073 |
| 14days | ID_6 | avg_Angle__left__back__hip_ankle_toe | 109.015013 |
| 14days | ID_7 | avg_Angle__left__back__hip_ankle_toe | 121.4844352 |
| 14days | ID_8 | avg_Angle__left__back__hip_ankle_toe | 132.0615907 |
| 14days | ID_9 | avg_Angle__left__back__hip_ankle_toe | 118.3741866 |
| 14days | ID_10 | avg_Angle__left__back__hip_ankle_toe | 116.3048084 |
| 14days | ID_11 | avg_Angle__left__back__hip_ankle_toe | 116.3657691 |
| 21days | ID_1 | avg_Angle__left__back__hip_ankle_toe | 121.3778167 |
| 21days | ID_2 | avg_Angle__left__back__hip_ankle_toe | 120.9414171 |
| 21days | ID_3 | avg_Angle__left__back__hip_ankle_toe | 117.2057509 |
| 21days | ID_4 | avg_Angle__left__back__hip_ankle_toe | 109.018637 |
| 21days | ID_5 | avg_Angle__left__back__hip_ankle_toe | 110.2264305 |

| | | | |
|---|---|---|---|
| 21days | ID_6 | avg_Angle__left__back__hip_ankle_toe | 110.245508 |
| 21days | ID_7 | avg_Angle__left__back__hip_ankle_toe | 115.0626434 |
| 21days | ID_8 | avg_Angle__left__back__hip_ankle_toe | 132.4976511 |
| 21days | ID_9 | avg_Angle__left__back__hip_ankle_toe | 125.3111799 |
| 21days | ID_10 | avg_Angle__left__back__hip_ankle_toe | 130.1877934 |
| 0days | ID_1 | avg_Angle__left__back__iliac_hip_ankle | 135.0505341 |
| 0days | ID_2 | avg_Angle__left__back__iliac_hip_ankle | 135.5793202 |
| 0days | ID_3 | avg_Angle__left__back__iliac_hip_ankle | 144.4318321 |
| 0days | ID_4 | avg_Angle__left__back__iliac_hip_ankle | 135.7760854 |
| 0days | ID_5 | avg_Angle__left__back__iliac_hip_ankle | 133.371106 |
| 0days | ID_6 | avg_Angle__left__back__iliac_hip_ankle | 138.2339967 |
| 0days | ID_7 | avg_Angle__left__back__iliac_hip_ankle | 142.3933013 |
| 0days | ID_8 | avg_Angle__left__back__iliac_hip_ankle | 143.1065826 |
| 0days | ID_9 | avg_Angle__left__back__iliac_hip_ankle | 131.2486022 |
| 0days | ID_10 | avg_Angle__left__back__iliac_hip_ankle | 141.9465582 |
| 0days | ID_11 | avg_Angle__left__back__iliac_hip_ankle | 155.9406815 |
| 0days | ID_12 | avg_Angle__left__back__iliac_hip_ankle | 127.003597 |
| 0days | ID_13 | avg_Angle__left__back__iliac_hip_ankle | 130.4798667 |
| 3days | ID_1 | avg_Angle__left__back__iliac_hip_ankle | 131.4994529 |
| 3days | ID_2 | avg_Angle__left__back__iliac_hip_ankle | 137.002001 |
| 3days | ID_3 | avg_Angle__left__back__iliac_hip_ankle | 170.6036914 |
| 3days | ID_4 | avg_Angle__left__back__iliac_hip_ankle | 139.9021649 |
| 3days | ID_5 | avg_Angle__left__back__iliac_hip_ankle | 145.1201929 |
| 3days | ID_6 | avg_Angle__left__back__iliac_hip_ankle | 163.3106646 |
| 3days | ID_7 | avg_Angle__left__back__iliac_hip_ankle | 157.2863115 |
| 3days | ID_8 | avg_Angle__left__back__iliac_hip_ankle | 145.5540685 |
| 7days | ID_1 | avg_Angle__left__back__iliac_hip_ankle | 148.901809 |
| 7days | ID_2 | avg_Angle__left__back__iliac_hip_ankle | 138.5188622 |
| 7days | ID_3 | avg_Angle__left__back__iliac_hip_ankle | 141.9063018 |
| 7days | ID_4 | avg_Angle__left__back__iliac_hip_ankle | 145.9886662 |
| 7days | ID_5 | avg_Angle__left__back__iliac_hip_ankle | 148.6831878 |
| 7days | ID_6 | avg_Angle__left__back__iliac_hip_ankle | 145.1924703 |
| 7days | ID_7 | avg_Angle__left__back__iliac_hip_ankle | 146.1507297 |
| 7days | ID_8 | avg_Angle__left__back__iliac_hip_ankle | 147.2430243 |
| 7days | ID_9 | avg_Angle__left__back__iliac_hip_ankle | 139.1611706 |
| 14days | ID_1 | avg_Angle__left__back__iliac_hip_ankle | 139.9935782 |
| 14days | ID_2 | avg_Angle__left__back__iliac_hip_ankle | 138.3982793 |
| 14days | ID_3 | avg_Angle__left__back__iliac_hip_ankle | 171.7831316 |
| 14days | ID_4 | avg_Angle__left__back__iliac_hip_ankle | 136.8134784 |
| 14days | ID_5 | avg_Angle__left__back__iliac_hip_ankle | 137.8729391 |
| 14days | ID_6 | avg_Angle__left__back__iliac_hip_ankle | 144.0741428 |
| 14days | ID_7 | avg_Angle__left__back__iliac_hip_ankle | 138.5934962 |
| 14days | ID_8 | avg_Angle__left__back__iliac_hip_ankle | 147.2877546 |
| 14days | ID_9 | avg_Angle__left__back__iliac_hip_ankle | 150.289629 |
| 14days | ID_10 | avg_Angle__left__back__iliac_hip_ankle | 143.6005744 |
| 14days | ID_11 | avg_Angle__left__back__iliac_hip_ankle | 127.0132765 |
| 21days | ID_1 | avg_Angle__left__back__iliac_hip_ankle | 145.752007 |
| 21days | ID_2 | avg_Angle__left__back__iliac_hip_ankle | 134.3066689 |

| | | | |
|---|---|---|---|
| 21days | ID_3 | avg_Angle__left__back__iliac_hip_ankle | 131.6968323 |
| 21days | ID_4 | avg_Angle__left__back__iliac_hip_ankle | 134.7369823 |
| 21days | ID_5 | avg_Angle__left__back__iliac_hip_ankle | 132.8104309 |
| 21days | ID_6 | avg_Angle__left__back__iliac_hip_ankle | 138.8536897 |
| 21days | ID_7 | avg_Angle__left__back__iliac_hip_ankle | 138.2547258 |
| 21days | ID_8 | avg_Angle__left__back__iliac_hip_ankle | 135.402751 |
| 21days | ID_9 | avg_Angle__left__back__iliac_hip_ankle | 137.5116714 |
| 21days | ID_10 | avg_Angle__left__back__iliac_hip_ankle | 138.3122277 |
| 0days | ID_1 | avg_Angle__left__front__elbow_wrist_toetip | 144.2891192 |
| 0days | ID_2 | avg_Angle__left__front__elbow_wrist_toetip | 151.1045287 |
| 0days | ID_3 | avg_Angle__left__front__elbow_wrist_toetip | 141.0231684 |
| 0days | ID_4 | avg_Angle__left__front__elbow_wrist_toetip | 140.8208984 |
| 0days | ID_5 | avg_Angle__left__front__elbow_wrist_toetip | 149.9373826 |
| 0days | ID_6 | avg_Angle__left__front__elbow_wrist_toetip | 144.0921415 |
| 0days | ID_7 | avg_Angle__left__front__elbow_wrist_toetip | 142.3731221 |
| 0days | ID_8 | avg_Angle__left__front__elbow_wrist_toetip | 146.789637 |
| 0days | ID_9 | avg_Angle__left__front__elbow_wrist_toetip | 145.8770071 |
| 0days | ID_10 | avg_Angle__left__front__elbow_wrist_toetip | 140.9206528 |
| 0days | ID_11 | avg_Angle__left__front__elbow_wrist_toetip | 130.8326697 |
| 0days | ID_12 | avg_Angle__left__front__elbow_wrist_toetip | 149.3871516 |
| 0days | ID_13 | avg_Angle__left__front__elbow_wrist_toetip | 165.2511709 |
| 3days | ID_1 | avg_Angle__left__front__elbow_wrist_toetip | 146.8133437 |
| 3days | ID_2 | avg_Angle__left__front__elbow_wrist_toetip | 139.6066276 |
| 3days | ID_3 | avg_Angle__left__front__elbow_wrist_toetip | 137.106981 |
| 3days | ID_4 | avg_Angle__left__front__elbow_wrist_toetip | 134.1458412 |
| 3days | ID_5 | avg_Angle__left__front__elbow_wrist_toetip | 148.8868582 |
| 3days | ID_6 | avg_Angle__left__front__elbow_wrist_toetip | 121.7303035 |
| 3days | ID_7 | avg_Angle__left__front__elbow_wrist_toetip | 131.7931426 |
| 3days | ID_8 | avg_Angle__left__front__elbow_wrist_toetip | 137.0905538 |
| 7days | ID_1 | avg_Angle__left__front__elbow_wrist_toetip | 139.3027937 |
| 7days | ID_2 | avg_Angle__left__front__elbow_wrist_toetip | 152.1742211 |
| 7days | ID_3 | avg_Angle__left__front__elbow_wrist_toetip | 148.1289414 |
| 7days | ID_4 | avg_Angle__left__front__elbow_wrist_toetip | 142.5887226 |
| 7days | ID_5 | avg_Angle__left__front__elbow_wrist_toetip | 133.3288446 |
| 7days | ID_6 | avg_Angle__left__front__elbow_wrist_toetip | 139.6149882 |
| 7days | ID_7 | avg_Angle__left__front__elbow_wrist_toetip | 136.1857868 |
| 7days | ID_8 | avg_Angle__left__front__elbow_wrist_toetip | 153.7552099 |
| 7days | ID_9 | avg_Angle__left__front__elbow_wrist_toetip | 135.4245809 |
| 14days | ID_1 | avg_Angle__left__front__elbow_wrist_toetip | 121.5699832 |
| 14days | ID_2 | avg_Angle__left__front__elbow_wrist_toetip | 125.4997733 |
| 14days | ID_3 | avg_Angle__left__front__elbow_wrist_toetip | 130.359543 |
| 14days | ID_4 | avg_Angle__left__front__elbow_wrist_toetip | 136.1223004 |
| 14days | ID_5 | avg_Angle__left__front__elbow_wrist_toetip | 132.2485372 |
| 14days | ID_6 | avg_Angle__left__front__elbow_wrist_toetip | 137.4785996 |
| 14days | ID_7 | avg_Angle__left__front__elbow_wrist_toetip | 149.7871871 |
| 14days | ID_8 | avg_Angle__left__front__elbow_wrist_toetip | 149.0367611 |
| 14days | ID_9 | avg_Angle__left__front__elbow_wrist_toetip | 132.8074463 |
| 14days | ID_10 | avg_Angle__left__front__elbow_wrist_toetip | 133.3898257 |

| | | | |
|---|---|---|---|
| 14days | ID_11 | avg_Angle__left__front__elbow_wrist_toetip | 135.5794195 |
| 21days | ID_1 | avg_Angle__left__front__elbow_wrist_toetip | 134.826873 |
| 21days | ID_2 | avg_Angle__left__front__elbow_wrist_toetip | 124.349755 |
| 21days | ID_3 | avg_Angle__left__front__elbow_wrist_toetip | 148.4783237 |
| 21days | ID_4 | avg_Angle__left__front__elbow_wrist_toetip | 121.3859247 |
| 21days | ID_5 | avg_Angle__left__front__elbow_wrist_toetip | 151.7144917 |
| 21days | ID_6 | avg_Angle__left__front__elbow_wrist_toetip | 127.5613641 |
| 21days | ID_7 | avg_Angle__left__front__elbow_wrist_toetip | 126.4471065 |
| 21days | ID_8 | avg_Angle__left__front__elbow_wrist_toetip | 141.4164273 |
| 21days | ID_9 | avg_Angle__left__front__elbow_wrist_toetip | 144.2206726 |
| 21days | ID_10 | avg_Angle__left__front__elbow_wrist_toetip | 145.2322498 |
| 0days | ID_1 | avg_Angle__left__front__shoulder_elbow_wrist | 118.090206 |
| 0days | ID_2 | avg_Angle__left__front__shoulder_elbow_wrist | 115.5624276 |
| 0days | ID_3 | avg_Angle__left__front__shoulder_elbow_wrist | 137.0553127 |
| 0days | ID_4 | avg_Angle__left__front__shoulder_elbow_wrist | 121.8632327 |
| 0days | ID_5 | avg_Angle__left__front__shoulder_elbow_wrist | 115.7192397 |
| 0days | ID_6 | avg_Angle__left__front__shoulder_elbow_wrist | 120.8249607 |
| 0days | ID_7 | avg_Angle__left__front__shoulder_elbow_wrist | 118.5762272 |
| 0days | ID_8 | avg_Angle__left__front__shoulder_elbow_wrist | 118.7520815 |
| 0days | ID_9 | avg_Angle__left__front__shoulder_elbow_wrist | 116.9937495 |
| 0days | ID_10 | avg_Angle__left__front__shoulder_elbow_wrist | 118.5028724 |
| 0days | ID_11 | avg_Angle__left__front__shoulder_elbow_wrist | 129.1210117 |
| 0days | ID_12 | avg_Angle__left__front__shoulder_elbow_wrist | 114.4313034 |
| 0days | ID_13 | avg_Angle__left__front__shoulder_elbow_wrist | 121.3388142 |
| 3days | ID_1 | avg_Angle__left__front__shoulder_elbow_wrist | 124.0229817 |
| 3days | ID_2 | avg_Angle__left__front__shoulder_elbow_wrist | 126.3040495 |
| 3days | ID_3 | avg_Angle__left__front__shoulder_elbow_wrist | 118.4023969 |
| 3days | ID_4 | avg_Angle__left__front__shoulder_elbow_wrist | 124.0874927 |
| 3days | ID_5 | avg_Angle__left__front__shoulder_elbow_wrist | 123.6507557 |
| 3days | ID_6 | avg_Angle__left__front__shoulder_elbow_wrist | 127.5112357 |
| 3days | ID_7 | avg_Angle__left__front__shoulder_elbow_wrist | 121.8592033 |
| 3days | ID_8 | avg_Angle__left__front__shoulder_elbow_wrist | 121.9268735 |
| 7days | ID_1 | avg_Angle__left__front__shoulder_elbow_wrist | 126.7409911 |
| 7days | ID_2 | avg_Angle__left__front__shoulder_elbow_wrist | 108.3339656 |
| 7days | ID_3 | avg_Angle__left__front__shoulder_elbow_wrist | 124.1091533 |
| 7days | ID_4 | avg_Angle__left__front__shoulder_elbow_wrist | 122.2584768 |
| 7days | ID_5 | avg_Angle__left__front__shoulder_elbow_wrist | 127.9885279 |
| 7days | ID_6 | avg_Angle__left__front__shoulder_elbow_wrist | 117.9327268 |
| 7days | ID_7 | avg_Angle__left__front__shoulder_elbow_wrist | 123.8393014 |
| 7days | ID_8 | avg_Angle__left__front__shoulder_elbow_wrist | 116.0393596 |
| 7days | ID_9 | avg_Angle__left__front__shoulder_elbow_wrist | 121.0498651 |
| 14days | ID_1 | avg_Angle__left__front__shoulder_elbow_wrist | 136.2185214 |
| 14days | ID_2 | avg_Angle__left__front__shoulder_elbow_wrist | 138.3342963 |
| 14days | ID_3 | avg_Angle__left__front__shoulder_elbow_wrist | 124.3993537 |
| 14days | ID_4 | avg_Angle__left__front__shoulder_elbow_wrist | 118.1238956 |
| 14days | ID_5 | avg_Angle__left__front__shoulder_elbow_wrist | 119.8397393 |
| 14days | ID_6 | avg_Angle__left__front__shoulder_elbow_wrist | 123.1079442 |
| 14days | ID_7 | avg_Angle__left__front__shoulder_elbow_wrist | 114.9154838 |

| | | | |
|---|---|---|---|
| 14days | ID_8 | avg_Angle__left__front__shoulder_elbow_wrist | 112.221325 |
| 14days | ID_9 | avg_Angle__left__front__shoulder_elbow_wrist | 134.8577193 |
| 14days | ID_10 | avg_Angle__left__front__shoulder_elbow_wrist | 131.066123 |
| 14days | ID_11 | avg_Angle__left__front__shoulder_elbow_wrist | 128.6653096 |
| 21days | ID_1 | avg_Angle__left__front__shoulder_elbow_wrist | 118.5018086 |
| 21days | ID_2 | avg_Angle__left__front__shoulder_elbow_wrist | 134.9192923 |
| 21days | ID_3 | avg_Angle__left__front__shoulder_elbow_wrist | 119.6870298 |
| 21days | ID_4 | avg_Angle__left__front__shoulder_elbow_wrist | 133.1994913 |
| 21days | ID_5 | avg_Angle__left__front__shoulder_elbow_wrist | 124.0967548 |
| 21days | ID_6 | avg_Angle__left__front__shoulder_elbow_wrist | 138.6167141 |
| 21days | ID_7 | avg_Angle__left__front__shoulder_elbow_wrist | 129.9839995 |
| 21days | ID_8 | avg_Angle__left__front__shoulder_elbow_wrist | 117.7658956 |
| 21days | ID_9 | avg_Angle__left__front__shoulder_elbow_wrist | 110.2205771 |
| 21days | ID_10 | avg_Angle__left__front__shoulder_elbow_wrist | 119.8867237 |
| 0days | ID_1 | avg_Angle__right__back__hip_ankle_toe | 114.5769635 |
| 0days | ID_2 | avg_Angle__right__back__hip_ankle_toe | 122.2705118 |
| 0days | ID_3 | avg_Angle__right__back__hip_ankle_toe | 103.3324766 |
| 0days | ID_4 | avg_Angle__right__back__hip_ankle_toe | 112.8440145 |
| 0days | ID_5 | avg_Angle__right__back__hip_ankle_toe | 110.2712448 |
| 0days | ID_6 | avg_Angle__right__back__hip_ankle_toe | 121.8546027 |
| 0days | ID_7 | avg_Angle__right__back__hip_ankle_toe | 126.7599113 |
| 0days | ID_8 | avg_Angle__right__back__hip_ankle_toe | 121.4244838 |
| 0days | ID_9 | avg_Angle__right__back__hip_ankle_toe | 113.8967985 |
| 0days | ID_10 | avg_Angle__right__back__hip_ankle_toe | 110.1801702 |
| 0days | ID_11 | avg_Angle__right__back__hip_ankle_toe | 108.940755 |
| 0days | ID_12 | avg_Angle__right__back__hip_ankle_toe | 120.5667082 |
| 0days | ID_13 | avg_Angle__right__back__hip_ankle_toe | 119.9883246 |
| 3days | ID_1 | avg_Angle__right__back__hip_ankle_toe | 106.8085309 |
| 3days | ID_2 | avg_Angle__right__back__hip_ankle_toe | 103.6828796 |
| 3days | ID_3 | avg_Angle__right__back__hip_ankle_toe | 102.5763069 |
| 3days | ID_4 | avg_Angle__right__back__hip_ankle_toe | 107.0847789 |
| 3days | ID_5 | avg_Angle__right__back__hip_ankle_toe | 115.7164427 |
| 3days | ID_6 | avg_Angle__right__back__hip_ankle_toe | 101.30363 |
| 3days | ID_7 | avg_Angle__right__back__hip_ankle_toe | 115.9324685 |
| 3days | ID_8 | avg_Angle__right__back__hip_ankle_toe | 108.7504831 |
| 7days | ID_1 | avg_Angle__right__back__hip_ankle_toe | 105.1805545 |
| 7days | ID_2 | avg_Angle__right__back__hip_ankle_toe | 104.6534542 |
| 7days | ID_3 | avg_Angle__right__back__hip_ankle_toe | 106.6916234 |
| 7days | ID_4 | avg_Angle__right__back__hip_ankle_toe | 116.8184409 |
| 7days | ID_5 | avg_Angle__right__back__hip_ankle_toe | 122.1769937 |
| 7days | ID_6 | avg_Angle__right__back__hip_ankle_toe | 113.2392476 |
| 7days | ID_7 | avg_Angle__right__back__hip_ankle_toe | 122.0734282 |
| 7days | ID_8 | avg_Angle__right__back__hip_ankle_toe | 112.2490091 |
| 7days | ID_9 | avg_Angle__right__back__hip_ankle_toe | 115.0983191 |
| 14days | ID_1 | avg_Angle__right__back__hip_ankle_toe | 111.8480481 |
| 14days | ID_2 | avg_Angle__right__back__hip_ankle_toe | 110.8127988 |
| 14days | ID_3 | avg_Angle__right__back__hip_ankle_toe | 106.8557434 |
| 14days | ID_4 | avg_Angle__right__back__hip_ankle_toe | 111.8824012 |

| | | | |
|---|---|---|---|
| 14days | ID_5 | avg_Angle__right__back__hip_ankle_toe | 111.6102234 |
| 14days | ID_6 | avg_Angle__right__back__hip_ankle_toe | 106.1085687 |
| 14days | ID_7 | avg_Angle__right__back__hip_ankle_toe | 120.6597897 |
| 14days | ID_8 | avg_Angle__right__back__hip_ankle_toe | 123.1715971 |
| 14days | ID_9 | avg_Angle__right__back__hip_ankle_toe | 122.5984961 |
| 14days | ID_10 | avg_Angle__right__back__hip_ankle_toe | 115.5226258 |
| 14days | ID_11 | avg_Angle__right__back__hip_ankle_toe | 106.8107176 |
| 21days | ID_1 | avg_Angle__right__back__hip_ankle_toe | 115.293123 |
| 21days | ID_2 | avg_Angle__right__back__hip_ankle_toe | 124.3191046 |
| 21days | ID_3 | avg_Angle__right__back__hip_ankle_toe | 109.0419382 |
| 21days | ID_4 | avg_Angle__right__back__hip_ankle_toe | 109.2447171 |
| 21days | ID_5 | avg_Angle__right__back__hip_ankle_toe | 108.6220688 |
| 21days | ID_6 | avg_Angle__right__back__hip_ankle_toe | 113.6744101 |
| 21days | ID_7 | avg_Angle__right__back__hip_ankle_toe | 115.6251443 |
| 21days | ID_8 | avg_Angle__right__back__hip_ankle_toe | 113.520347 |
| 21days | ID_9 | avg_Angle__right__back__hip_ankle_toe | 117.1446924 |
| 21days | ID_10 | avg_Angle__right__back__hip_ankle_toe | 124.3530475 |
| 0days | ID_1 | avg_Angle__right__back__iliac_hip_ankle | 139.9973957 |
| 0days | ID_2 | avg_Angle__right__back__iliac_hip_ankle | 132.2151988 |
| 0days | ID_3 | avg_Angle__right__back__iliac_hip_ankle | 132.1324056 |
| 0days | ID_4 | avg_Angle__right__back__iliac_hip_ankle | 138.7875944 |
| 0days | ID_5 | avg_Angle__right__back__iliac_hip_ankle | 135.5329007 |
| 0days | ID_6 | avg_Angle__right__back__iliac_hip_ankle | 143.9298157 |
| 0days | ID_7 | avg_Angle__right__back__iliac_hip_ankle | 140.863859 |
| 0days | ID_8 | avg_Angle__right__back__iliac_hip_ankle | 144.9916763 |
| 0days | ID_9 | avg_Angle__right__back__iliac_hip_ankle | 138.9638264 |
| 0days | ID_10 | avg_Angle__right__back__iliac_hip_ankle | 139.7879085 |
| 0days | ID_11 | avg_Angle__right__back__iliac_hip_ankle | 145.3395096 |
| 0days | ID_12 | avg_Angle__right__back__iliac_hip_ankle | 136.4507223 |
| 0days | ID_13 | avg_Angle__right__back__iliac_hip_ankle | 135.6055551 |
| 3days | ID_1 | avg_Angle__right__back__iliac_hip_ankle | 145.0867908 |
| 3days | ID_2 | avg_Angle__right__back__iliac_hip_ankle | 138.6000723 |
| 3days | ID_3 | avg_Angle__right__back__iliac_hip_ankle | 158.7954739 |
| 3days | ID_4 | avg_Angle__right__back__iliac_hip_ankle | 152.3592993 |
| 3days | ID_5 | avg_Angle__right__back__iliac_hip_ankle | 152.4397201 |
| 3days | ID_6 | avg_Angle__right__back__iliac_hip_ankle | 157.2265762 |
| 3days | ID_7 | avg_Angle__right__back__iliac_hip_ankle | 158.5761679 |
| 3days | ID_8 | avg_Angle__right__back__iliac_hip_ankle | 146.042992 |
| 7days | ID_1 | avg_Angle__right__back__iliac_hip_ankle | 134.5954576 |
| 7days | ID_2 | avg_Angle__right__back__iliac_hip_ankle | 134.4924278 |
| 7days | ID_3 | avg_Angle__right__back__iliac_hip_ankle | 139.6648178 |
| 7days | ID_4 | avg_Angle__right__back__iliac_hip_ankle | 143.5968235 |
| 7days | ID_5 | avg_Angle__right__back__iliac_hip_ankle | 142.1148327 |
| 7days | ID_6 | avg_Angle__right__back__iliac_hip_ankle | 140.4937131 |
| 7days | ID_7 | avg_Angle__right__back__iliac_hip_ankle | 133.3226465 |
| 7days | ID_8 | avg_Angle__right__back__iliac_hip_ankle | 139.603305 |
| 7days | ID_9 | avg_Angle__right__back__iliac_hip_ankle | 130.7913714 |
| 14days | ID_1 | avg_Angle__right__back__iliac_hip_ankle | 136.2968398 |

| | | | |
|---|---|---|---|
| 14days | ID_2 | avg_Angle__right__back__iliac_hip_ankle | 136.3452176 |
| 14days | ID_3 | avg_Angle__right__back__iliac_hip_ankle | 145.6442664 |
| 14days | ID_4 | avg_Angle__right__back__iliac_hip_ankle | 136.4898882 |
| 14days | ID_5 | avg_Angle__right__back__iliac_hip_ankle | 135.8930869 |
| 14days | ID_6 | avg_Angle__right__back__iliac_hip_ankle | 134.8852585 |
| 14days | ID_7 | avg_Angle__right__back__iliac_hip_ankle | 146.1150925 |
| 14days | ID_8 | avg_Angle__right__back__iliac_hip_ankle | 136.7045303 |
| 14days | ID_9 | avg_Angle__right__back__iliac_hip_ankle | 135.7985651 |
| 14days | ID_10 | avg_Angle__right__back__iliac_hip_ankle | 126.191667 |
| 14days | ID_11 | avg_Angle__right__back__iliac_hip_ankle | 133.6199036 |
| 21days | ID_1 | avg_Angle__right__back__iliac_hip_ankle | 130.3417223 |
| 21days | ID_2 | avg_Angle__right__back__iliac_hip_ankle | 134.8075987 |
| 21days | ID_3 | avg_Angle__right__back__iliac_hip_ankle | 132.5816702 |
| 21days | ID_4 | avg_Angle__right__back__iliac_hip_ankle | 134.2766531 |
| 21days | ID_5 | avg_Angle__right__back__iliac_hip_ankle | 137.0984114 |
| 21days | ID_6 | avg_Angle__right__back__iliac_hip_ankle | 139.8085174 |
| 21days | ID_7 | avg_Angle__right__back__iliac_hip_ankle | 131.4147009 |
| 21days | ID_8 | avg_Angle__right__back__iliac_hip_ankle | 142.0365838 |
| 21days | ID_9 | avg_Angle__right__back__iliac_hip_ankle | 133.996252 |
| 21days | ID_10 | avg_Angle__right__back__iliac_hip_ankle | 132.5078168 |
| 0days | ID_1 | avg_Angle__right__front__elbow_wrist_toetip | 147.4149331 |
| 0days | ID_2 | avg_Angle__right__front__elbow_wrist_toetip | 144.1783268 |
| 0days | ID_3 | avg_Angle__right__front__elbow_wrist_toetip | 144.5530737 |
| 0days | ID_4 | avg_Angle__right__front__elbow_wrist_toetip | 148.8118398 |
| 0days | ID_5 | avg_Angle__right__front__elbow_wrist_toetip | 137.4973971 |
| 0days | ID_6 | avg_Angle__right__front__elbow_wrist_toetip | 143.5859217 |
| 0days | ID_7 | avg_Angle__right__front__elbow_wrist_toetip | 142.3324726 |
| 0days | ID_8 | avg_Angle__right__front__elbow_wrist_toetip | 149.4534672 |
| 0days | ID_9 | avg_Angle__right__front__elbow_wrist_toetip | 140.2925521 |
| 0days | ID_10 | avg_Angle__right__front__elbow_wrist_toetip | 139.6580109 |
| 0days | ID_11 | avg_Angle__right__front__elbow_wrist_toetip | 141.1675697 |
| 0days | ID_12 | avg_Angle__right__front__elbow_wrist_toetip | 138.9766468 |
| 0days | ID_13 | avg_Angle__right__front__elbow_wrist_toetip | 137.7000768 |
| 3days | ID_1 | avg_Angle__right__front__elbow_wrist_toetip | 131.7338568 |
| 3days | ID_2 | avg_Angle__right__front__elbow_wrist_toetip | 140.388301 |
| 3days | ID_3 | avg_Angle__right__front__elbow_wrist_toetip | 130.3957893 |
| 3days | ID_4 | avg_Angle__right__front__elbow_wrist_toetip | 134.4634459 |
| 3days | ID_5 | avg_Angle__right__front__elbow_wrist_toetip | 129.943651 |
| 3days | ID_6 | avg_Angle__right__front__elbow_wrist_toetip | 121.2009802 |
| 3days | ID_7 | avg_Angle__right__front__elbow_wrist_toetip | 128.0118273 |
| 3days | ID_8 | avg_Angle__right__front__elbow_wrist_toetip | 129.0815659 |
| 7days | ID_1 | avg_Angle__right__front__elbow_wrist_toetip | 126.7751667 |
| 7days | ID_2 | avg_Angle__right__front__elbow_wrist_toetip | 123.717096 |
| 7days | ID_3 | avg_Angle__right__front__elbow_wrist_toetip | 128.4997988 |
| 7days | ID_4 | avg_Angle__right__front__elbow_wrist_toetip | 136.6072312 |
| 7days | ID_5 | avg_Angle__right__front__elbow_wrist_toetip | 151.5066857 |
| 7days | ID_6 | avg_Angle__right__front__elbow_wrist_toetip | 132.9994875 |
| 7days | ID_7 | avg_Angle__right__front__elbow_wrist_toetip | 144.6194409 |

| | | | |
|---|---|---|---|
| 7days | ID_8 | avg_Angle__right__front__elbow_wrist_toetip | 122.4405135 |
| 7days | ID_9 | avg_Angle__right__front__elbow_wrist_toetip | 134.7909266 |
| 14days | ID_1 | avg_Angle__right__front__elbow_wrist_toetip | 131.1095135 |
| 14days | ID_2 | avg_Angle__right__front__elbow_wrist_toetip | 139.8249006 |
| 14days | ID_3 | avg_Angle__right__front__elbow_wrist_toetip | 118.1638702 |
| 14days | ID_4 | avg_Angle__right__front__elbow_wrist_toetip | 137.4691995 |
| 14days | ID_5 | avg_Angle__right__front__elbow_wrist_toetip | 133.8326746 |
| 14days | ID_6 | avg_Angle__right__front__elbow_wrist_toetip | 141.5698108 |
| 14days | ID_7 | avg_Angle__right__front__elbow_wrist_toetip | 143.4577418 |
| 14days | ID_8 | avg_Angle__right__front__elbow_wrist_toetip | 128.863032 |
| 14days | ID_9 | avg_Angle__right__front__elbow_wrist_toetip | 149.2347708 |
| 14days | ID_10 | avg_Angle__right__front__elbow_wrist_toetip | 143.3999389 |
| 14days | ID_11 | avg_Angle__right__front__elbow_wrist_toetip | 133.0763005 |
| 21days | ID_1 | avg_Angle__right__front__elbow_wrist_toetip | 123.6987701 |
| 21days | ID_2 | avg_Angle__right__front__elbow_wrist_toetip | 144.9612586 |
| 21days | ID_3 | avg_Angle__right__front__elbow_wrist_toetip | 127.0699557 |
| 21days | ID_4 | avg_Angle__right__front__elbow_wrist_toetip | 142.9652855 |
| 21days | ID_5 | avg_Angle__right__front__elbow_wrist_toetip | 125.3413558 |
| 21days | ID_6 | avg_Angle__right__front__elbow_wrist_toetip | 146.6411507 |
| 21days | ID_7 | avg_Angle__right__front__elbow_wrist_toetip | 150.4408418 |
| 21days | ID_8 | avg_Angle__right__front__elbow_wrist_toetip | 136.7602109 |
| 21days | ID_9 | avg_Angle__right__front__elbow_wrist_toetip | 126.0278079 |
| 21days | ID_10 | avg_Angle__right__front__elbow_wrist_toetip | 129.9179322 |
| 0days | ID_1 | avg_Angle__right__front__shoulder_elbow_wrist | 113.2772849 |
| 0days | ID_2 | avg_Angle__right__front__shoulder_elbow_wrist | 126.8813728 |
| 0days | ID_3 | avg_Angle__right__front__shoulder_elbow_wrist | 125.4043527 |
| 0days | ID_4 | avg_Angle__right__front__shoulder_elbow_wrist | 119.6655822 |
| 0days | ID_5 | avg_Angle__right__front__shoulder_elbow_wrist | 116.8499557 |
| 0days | ID_6 | avg_Angle__right__front__shoulder_elbow_wrist | 127.2919935 |
| 0days | ID_7 | avg_Angle__right__front__shoulder_elbow_wrist | 120.9959559 |
| 0days | ID_8 | avg_Angle__right__front__shoulder_elbow_wrist | 115.0245043 |
| 0days | ID_9 | avg_Angle__right__front__shoulder_elbow_wrist | 134.3973256 |
| 0days | ID_10 | avg_Angle__right__front__shoulder_elbow_wrist | 119.6953038 |
| 0days | ID_11 | avg_Angle__right__front__shoulder_elbow_wrist | 116.0094194 |
| 0days | ID_12 | avg_Angle__right__front__shoulder_elbow_wrist | 123.6923289 |
| 0days | ID_13 | avg_Angle__right__front__shoulder_elbow_wrist | 131.2565151 |
| 3days | ID_1 | avg_Angle__right__front__shoulder_elbow_wrist | 126.3122977 |
| 3days | ID_2 | avg_Angle__right__front__shoulder_elbow_wrist | 126.139292 |
| 3days | ID_3 | avg_Angle__right__front__shoulder_elbow_wrist | 122.6527397 |
| 3days | ID_4 | avg_Angle__right__front__shoulder_elbow_wrist | 130.1594753 |
| 3days | ID_5 | avg_Angle__right__front__shoulder_elbow_wrist | 134.1475915 |
| 3days | ID_6 | avg_Angle__right__front__shoulder_elbow_wrist | 128.6257931 |
| 3days | ID_7 | avg_Angle__right__front__shoulder_elbow_wrist | 128.3984498 |
| 3days | ID_8 | avg_Angle__right__front__shoulder_elbow_wrist | 133.7257222 |
| 7days | ID_1 | avg_Angle__right__front__shoulder_elbow_wrist | 118.7624668 |
| 7days | ID_2 | avg_Angle__right__front__shoulder_elbow_wrist | 126.7617471 |
| 7days | ID_3 | avg_Angle__right__front__shoulder_elbow_wrist | 133.1418351 |
| 7days | ID_4 | avg_Angle__right__front__shoulder_elbow_wrist | 125.7375807 |

| | | | |
|---|---|---|---|
| 7days | ID_5 | avg_Angle__right__front__shoulder_elbow_wrist | 126.1182715 |
| 7days | ID_6 | avg_Angle__right__front__shoulder_elbow_wrist | 123.7695133 |
| 7days | ID_7 | avg_Angle__right__front__shoulder_elbow_wrist | 114.7837253 |
| 7days | ID_8 | avg_Angle__right__front__shoulder_elbow_wrist | 132.59376 |
| 7days | ID_9 | avg_Angle__right__front__shoulder_elbow_wrist | 114.3278074 |
| 14days | ID_1 | avg_Angle__right__front__shoulder_elbow_wrist | 129.8586273 |
| 14days | ID_2 | avg_Angle__right__front__shoulder_elbow_wrist | 126.3839048 |
| 14days | ID_3 | avg_Angle__right__front__shoulder_elbow_wrist | 125.23928 |
| 14days | ID_4 | avg_Angle__right__front__shoulder_elbow_wrist | 121.9049694 |
| 14days | ID_5 | avg_Angle__right__front__shoulder_elbow_wrist | 123.0158817 |
| 14days | ID_6 | avg_Angle__right__front__shoulder_elbow_wrist | 124.9294654 |
| 14days | ID_7 | avg_Angle__right__front__shoulder_elbow_wrist | 130.5094206 |
| 14days | ID_8 | avg_Angle__right__front__shoulder_elbow_wrist | 124.4738622 |
| 14days | ID_9 | avg_Angle__right__front__shoulder_elbow_wrist | 116.8330872 |
| 14days | ID_10 | avg_Angle__right__front__shoulder_elbow_wrist | 127.9545522 |
| 14days | ID_11 | avg_Angle__right__front__shoulder_elbow_wrist | 139.1968385 |
| 21days | ID_1 | avg_Angle__right__front__shoulder_elbow_wrist | 128.3816376 |
| 21days | ID_2 | avg_Angle__right__front__shoulder_elbow_wrist | 112.0888589 |
| 21days | ID_3 | avg_Angle__right__front__shoulder_elbow_wrist | 119.7675932 |
| 21days | ID_4 | avg_Angle__right__front__shoulder_elbow_wrist | 121.3687758 |
| 21days | ID_5 | avg_Angle__right__front__shoulder_elbow_wrist | 136.0642249 |
| 21days | ID_6 | avg_Angle__right__front__shoulder_elbow_wrist | 122.9247985 |
| 21days | ID_7 | avg_Angle__right__front__shoulder_elbow_wrist | 119.0599052 |
| 21days | ID_8 | avg_Angle__right__front__shoulder_elbow_wrist | 128.6807356 |
| 21days | ID_9 | avg_Angle__right__front__shoulder_elbow_wrist | 135.3323797 |
| 21days | ID_10 | avg_Angle__right__front__shoulder_elbow_wrist | 137.6440652 |
| 0days | ID_1 | max__left__back__hip_ankle_toe | 91.72732898 |
| 0days | ID_2 | max__left__back__hip_ankle_toe | 103.415943 |
| 0days | ID_3 | max__left__back__hip_ankle_toe | 86.94896634 |
| 0days | ID_4 | max__left__back__hip_ankle_toe | 94.89330671 |
| 0days | ID_5 | max__left__back__hip_ankle_toe | 92.53341159 |
| 0days | ID_6 | max__left__back__hip_ankle_toe | 117.3569744 |
| 0days | ID_7 | max__left__back__hip_ankle_toe | 104.3963876 |
| 0days | ID_8 | max__left__back__hip_ankle_toe | 117.5357266 |
| 0days | ID_9 | max__left__back__hip_ankle_toe | 99.03623939 |
| 0days | ID_10 | max__left__back__hip_ankle_toe | 106.4866912 |
| 0days | ID_11 | max__left__back__hip_ankle_toe | 103.7768113 |
| 0days | ID_12 | max__left__back__hip_ankle_toe | 112.5409555 |
| 0days | ID_13 | max__left__back__hip_ankle_toe | 110.1309223 |
| 3days | ID_1 | max__left__back__hip_ankle_toe | 101.6614086 |
| 3days | ID_2 | max__left__back__hip_ankle_toe | 91.52570474 |
| 3days | ID_3 | max__left__back__hip_ankle_toe | 50.62944087 |
| 3days | ID_4 | max__left__back__hip_ankle_toe | 105.2300481 |
| 3days | ID_5 | max__left__back__hip_ankle_toe | 107.9986038 |
| 3days | ID_6 | max__left__back__hip_ankle_toe | 64.51797797 |
| 3days | ID_7 | max__left__back__hip_ankle_toe | 69.17091801 |
| 3days | ID_8 | max__left__back__hip_ankle_toe | 113.7393019 |
| 7days | ID_1 | max__left__back__hip_ankle_toe | 98.52715467 |

| | | | |
|---|---|---|---|
| 7days | ID_2 | max__left__back__hip_ankle_toe | 92.8590913 |
| 7days | ID_3 | max__left__back__hip_ankle_toe | 93.78106375 |
| 7days | ID_4 | max__left__back__hip_ankle_toe | 109.6313984 |
| 7days | ID_5 | max__left__back__hip_ankle_toe | 91.86109296 |
| 7days | ID_6 | max__left__back__hip_ankle_toe | 107.0616571 |
| 7days | ID_7 | max__left__back__hip_ankle_toe | 89.9443564 |
| 7days | ID_8 | max__left__back__hip_ankle_toe | 100.1166307 |
| 7days | ID_9 | max__left__back__hip_ankle_toe | 100.9042805 |
| 14days | ID_1 | max__left__back__hip_ankle_toe | 83.07903389 |
| 14days | ID_2 | max__left__back__hip_ankle_toe | 96.32599253 |
| 14days | ID_3 | max__left__back__hip_ankle_toe | 101.6801311 |
| 14days | ID_4 | max__left__back__hip_ankle_toe | 94.9717 |
| 14days | ID_5 | max__left__back__hip_ankle_toe | 92.57461962 |
| 14days | ID_6 | max__left__back__hip_ankle_toe | 94.93041555 |
| 14days | ID_7 | max__left__back__hip_ankle_toe | 103.0470832 |
| 14days | ID_8 | max__left__back__hip_ankle_toe | 99.14942948 |
| 14days | ID_9 | max__left__back__hip_ankle_toe | 99.2078674 |
| 14days | ID_10 | max__left__back__hip_ankle_toe | 102.8291154 |
| 14days | ID_11 | max__left__back__hip_ankle_toe | 101.9907584 |
| 21days | ID_1 | max__left__back__hip_ankle_toe | 110.083241 |
| 21days | ID_2 | max__left__back__hip_ankle_toe | 109.9518523 |
| 21days | ID_3 | max__left__back__hip_ankle_toe | 109.1246646 |
| 21days | ID_4 | max__left__back__hip_ankle_toe | 94.69368837 |
| 21days | ID_5 | max__left__back__hip_ankle_toe | 97.25821168 |
| 21days | ID_6 | max__left__back__hip_ankle_toe | 84.79785403 |
| 21days | ID_7 | max__left__back__hip_ankle_toe | 104.4087642 |
| 21days | ID_8 | max__left__back__hip_ankle_toe | 115.0738421 |
| 21days | ID_9 | max__left__back__hip_ankle_toe | 108.9990528 |
| 21days | ID_10 | max__left__back__hip_ankle_toe | 120.5220217 |
| 0days | ID_1 | max__left__back__iliac_hip_ankle | 115.3714243 |
| 0days | ID_2 | max__left__back__iliac_hip_ankle | 117.372634 |
| 0days | ID_3 | max__left__back__iliac_hip_ankle | 126.0277972 |
| 0days | ID_4 | max__left__back__iliac_hip_ankle | 115.9285448 |
| 0days | ID_5 | max__left__back__iliac_hip_ankle | 119.2465194 |
| 0days | ID_6 | max__left__back__iliac_hip_ankle | 117.4722069 |
| 0days | ID_7 | max__left__back__iliac_hip_ankle | 117.9216707 |
| 0days | ID_8 | max__left__back__iliac_hip_ankle | 119.4815222 |
| 0days | ID_9 | max__left__back__iliac_hip_ankle | 113.882047 |
| 0days | ID_10 | max__left__back__iliac_hip_ankle | 110.8659699 |
| 0days | ID_11 | max__left__back__iliac_hip_ankle | 113.6981679 |
| 0days | ID_12 | max__left__back__iliac_hip_ankle | 107.4834909 |
| 0days | ID_13 | max__left__back__iliac_hip_ankle | 111.6424051 |
| 3days | ID_1 | max__left__back__iliac_hip_ankle | 115.6355542 |
| 3days | ID_2 | max__left__back__iliac_hip_ankle | 119.2463184 |
| 3days | ID_3 | max__left__back__iliac_hip_ankle | 137.8699229 |
| 3days | ID_4 | max__left__back__iliac_hip_ankle | 112.5517247 |
| 3days | ID_5 | max__left__back__iliac_hip_ankle | 118.6173548 |
| 3days | ID_6 | max__left__back__iliac_hip_ankle | 138.6060308 |

| | | | |
|---|---|---|---|
| 3days | ID_7 | max__left__back__iliac_hip_ankle | 134.0105199 |
| 3days | ID_8 | max__left__back__iliac_hip_ankle | 114.0096058 |
| 7days | ID_1 | max__left__back__iliac_hip_ankle | 111.5305125 |
| 7days | ID_2 | max__left__back__iliac_hip_ankle | 117.4604646 |
| 7days | ID_3 | max__left__back__iliac_hip_ankle | 119.3846507 |
| 7days | ID_4 | max__left__back__iliac_hip_ankle | 119.5076853 |
| 7days | ID_5 | max__left__back__iliac_hip_ankle | 126.3379295 |
| 7days | ID_6 | max__left__back__iliac_hip_ankle | 111.7489541 |
| 7days | ID_7 | max__left__back__iliac_hip_ankle | 117.2496673 |
| 7days | ID_8 | max__left__back__iliac_hip_ankle | 118.186402 |
| 7days | ID_9 | max__left__back__iliac_hip_ankle | 112.3142177 |
| 14days | ID_1 | max__left__back__iliac_hip_ankle | 116.937847 |
| 14days | ID_2 | max__left__back__iliac_hip_ankle | 115.1037084 |
| 14days | ID_3 | max__left__back__iliac_hip_ankle | 115.4921607 |
| 14days | ID_4 | max__left__back__iliac_hip_ankle | 114.6037582 |
| 14days | ID_5 | max__left__back__iliac_hip_ankle | 114.0522261 |
| 14days | ID_6 | max__left__back__iliac_hip_ankle | 117.8522634 |
| 14days | ID_7 | max__left__back__iliac_hip_ankle | 114.5243285 |
| 14days | ID_8 | max__left__back__iliac_hip_ankle | 118.9617234 |
| 14days | ID_9 | max__left__back__iliac_hip_ankle | 126.5048296 |
| 14days | ID_10 | max__left__back__iliac_hip_ankle | 117.2050801 |
| 14days | ID_11 | max__left__back__iliac_hip_ankle | 111.5723757 |
| 21days | ID_1 | max__left__back__iliac_hip_ankle | 109.5141684 |
| 21days | ID_2 | max__left__back__iliac_hip_ankle | 112.4554472 |
| 21days | ID_3 | max__left__back__iliac_hip_ankle | 110.4079356 |
| 21days | ID_4 | max__left__back__iliac_hip_ankle | 107.3789559 |
| 21days | ID_5 | max__left__back__iliac_hip_ankle | 116.0243885 |
| 21days | ID_6 | max__left__back__iliac_hip_ankle | 115.9531074 |
| 21days | ID_7 | max__left__back__iliac_hip_ankle | 115.1965799 |
| 21days | ID_8 | max__left__back__iliac_hip_ankle | 107.166235 |
| 21days | ID_9 | max__left__back__iliac_hip_ankle | 106.5565077 |
| 21days | ID_10 | max__left__back__iliac_hip_ankle | 118.5481452 |
| 0days | ID_1 | max__left__front__elbow_wrist_toetip | 109.1394446 |
| 0days | ID_2 | max__left__front__elbow_wrist_toetip | 115.0539533 |
| 0days | ID_3 | max__left__front__elbow_wrist_toetip | 117.5839455 |
| 0days | ID_4 | max__left__front__elbow_wrist_toetip | 112.1229604 |
| 0days | ID_5 | max__left__front__elbow_wrist_toetip | 110.4515599 |
| 0days | ID_6 | max__left__front__elbow_wrist_toetip | 111.7664072 |
| 0days | ID_7 | max__left__front__elbow_wrist_toetip | 109.6098502 |
| 0days | ID_8 | max__left__front__elbow_wrist_toetip | 114.7311377 |
| 0days | ID_9 | max__left__front__elbow_wrist_toetip | 117.5478112 |
| 0days | ID_10 | max__left__front__elbow_wrist_toetip | 106.8709378 |
| 0days | ID_11 | max__left__front__elbow_wrist_toetip | 96.15509457 |
| 0days | ID_12 | max__left__front__elbow_wrist_toetip | 115.4105023 |
| 0days | ID_13 | max__left__front__elbow_wrist_toetip | 114.237395 |
| 3days | ID_1 | max__left__front__elbow_wrist_toetip | 107.7074702 |
| 3days | ID_2 | max__left__front__elbow_wrist_toetip | 103.5673784 |
| 3days | ID_3 | max__left__front__elbow_wrist_toetip | 88.77930008 |

| | | | |
|---|---|---|---|
| 3days | ID_4 | max__left__front__elbow_wrist_toetip | 101.5046599 |
| 3days | ID_5 | max__left__front__elbow_wrist_toetip | 111.288825 |
| 3days | ID_6 | max__left__front__elbow_wrist_toetip | 86.92063815 |
| 3days | ID_7 | max__left__front__elbow_wrist_toetip | 88.82415752 |
| 3days | ID_8 | max__left__front__elbow_wrist_toetip | 106.0736816 |
| 7days | ID_1 | max__left__front__elbow_wrist_toetip | 89.66225482 |
| 7days | ID_2 | max__left__front__elbow_wrist_toetip | 99.98182634 |
| 7days | ID_3 | max__left__front__elbow_wrist_toetip | 101.1681076 |
| 7days | ID_4 | max__left__front__elbow_wrist_toetip | 109.8275479 |
| 7days | ID_5 | max__left__front__elbow_wrist_toetip | 99.6299151 |
| 7days | ID_6 | max__left__front__elbow_wrist_toetip | 108.8489702 |
| 7days | ID_7 | max__left__front__elbow_wrist_toetip | 89.39504579 |
| 7days | ID_8 | max__left__front__elbow_wrist_toetip | 110.7440502 |
| 7days | ID_9 | max__left__front__elbow_wrist_toetip | 93.27650942 |
| 14days | ID_1 | max__left__front__elbow_wrist_toetip | 89.50101884 |
| 14days | ID_2 | max__left__front__elbow_wrist_toetip | 92.82770689 |
| 14days | ID_3 | max__left__front__elbow_wrist_toetip | 74.48369071 |
| 14days | ID_4 | max__left__front__elbow_wrist_toetip | 93.15068127 |
| 14days | ID_5 | max__left__front__elbow_wrist_toetip | 92.3054446 |
| 14days | ID_6 | max__left__front__elbow_wrist_toetip | 103.1200699 |
| 14days | ID_7 | max__left__front__elbow_wrist_toetip | 111.6858828 |
| 14days | ID_8 | max__left__front__elbow_wrist_toetip | 114.2046072 |
| 14days | ID_9 | max__left__front__elbow_wrist_toetip | 103.6076443 |
| 14days | ID_10 | max__left__front__elbow_wrist_toetip | 93.22818414 |
| 14days | ID_11 | max__left__front__elbow_wrist_toetip | 102.6981546 |
| 21days | ID_1 | max__left__front__elbow_wrist_toetip | 106.3413062 |
| 21days | ID_2 | max__left__front__elbow_wrist_toetip | 89.01349576 |
| 21days | ID_3 | max__left__front__elbow_wrist_toetip | 108.3413232 |
| 21days | ID_4 | max__left__front__elbow_wrist_toetip | 91.94315286 |
| 21days | ID_5 | max__left__front__elbow_wrist_toetip | 104.1403033 |
| 21days | ID_6 | max__left__front__elbow_wrist_toetip | 92.71269221 |
| 21days | ID_7 | max__left__front__elbow_wrist_toetip | 93.69806701 |
| 21days | ID_8 | max__left__front__elbow_wrist_toetip | 104.3914548 |
| 21days | ID_9 | max__left__front__elbow_wrist_toetip | 112.6317496 |
| 21days | ID_10 | max__left__front__elbow_wrist_toetip | 108.3729804 |
| 0days | ID_1 | max__left__front__shoulder_elbow_wrist | 84.30849219 |
| 0days | ID_2 | max__left__front__shoulder_elbow_wrist | 86.62929107 |
| 0days | ID_3 | max__left__front__shoulder_elbow_wrist | 114.6183676 |
| 0days | ID_4 | max__left__front__shoulder_elbow_wrist | 88.00997075 |
| 0days | ID_5 | max__left__front__shoulder_elbow_wrist | 84.53577601 |
| 0days | ID_6 | max__left__front__shoulder_elbow_wrist | 92.35080845 |
| 0days | ID_7 | max__left__front__shoulder_elbow_wrist | 95.63154558 |
| 0days | ID_8 | max__left__front__shoulder_elbow_wrist | 88.54447942 |
| 0days | ID_9 | max__left__front__shoulder_elbow_wrist | 89.92213854 |
| 0days | ID_10 | max__left__front__shoulder_elbow_wrist | 88.9051869 |
| 0days | ID_11 | max__left__front__shoulder_elbow_wrist | 89.83989865 |
| 0days | ID_12 | max__left__front__shoulder_elbow_wrist | 88.97322094 |
| 0days | ID_13 | max__left__front__shoulder_elbow_wrist | 88.75442785 |

| | | | |
|---|---|---|---|
| 3days | ID_1 | max__left__front__shoulder_elbow_wrist | 87.90184741 |
| 3days | ID_2 | max__left__front__shoulder_elbow_wrist | 93.36346025 |
| 3days | ID_3 | max__left__front__shoulder_elbow_wrist | 84.58947978 |
| 3days | ID_4 | max__left__front__shoulder_elbow_wrist | 90.75839225 |
| 3days | ID_5 | max__left__front__shoulder_elbow_wrist | 94.3020503 |
| 3days | ID_6 | max__left__front__shoulder_elbow_wrist | 90.77141301 |
| 3days | ID_7 | max__left__front__shoulder_elbow_wrist | 102.7489449 |
| 3days | ID_8 | max__left__front__shoulder_elbow_wrist | 90.32943105 |
| 7days | ID_1 | max__left__front__shoulder_elbow_wrist | 85.3607612 |
| 7days | ID_2 | max__left__front__shoulder_elbow_wrist | 81.52548133 |
| 7days | ID_3 | max__left__front__shoulder_elbow_wrist | 90.66974203 |
| 7days | ID_4 | max__left__front__shoulder_elbow_wrist | 93.92620484 |
| 7days | ID_5 | max__left__front__shoulder_elbow_wrist | 97.96018927 |
| 7days | ID_6 | max__left__front__shoulder_elbow_wrist | 85.74692944 |
| 7days | ID_7 | max__left__front__shoulder_elbow_wrist | 94.25048263 |
| 7days | ID_8 | max__left__front__shoulder_elbow_wrist | 88.3251087 |
| 7days | ID_9 | max__left__front__shoulder_elbow_wrist | 98.89196048 |
| 14days | ID_1 | max__left__front__shoulder_elbow_wrist | 101.2726377 |
| 14days | ID_2 | max__left__front__shoulder_elbow_wrist | 102.3203625 |
| 14days | ID_3 | max__left__front__shoulder_elbow_wrist | 88.79720305 |
| 14days | ID_4 | max__left__front__shoulder_elbow_wrist | 91.51977852 |
| 14days | ID_5 | max__left__front__shoulder_elbow_wrist | 95.31226494 |
| 14days | ID_6 | max__left__front__shoulder_elbow_wrist | 89.89538177 |
| 14days | ID_7 | max__left__front__shoulder_elbow_wrist | 83.54411405 |
| 14days | ID_8 | max__left__front__shoulder_elbow_wrist | 80.75655096 |
| 14days | ID_9 | max__left__front__shoulder_elbow_wrist | 103.6051296 |
| 14days | ID_10 | max__left__front__shoulder_elbow_wrist | 100.5696788 |
| 14days | ID_11 | max__left__front__shoulder_elbow_wrist | 87.70256677 |
| 21days | ID_1 | max__left__front__shoulder_elbow_wrist | 83.46066055 |
| 21days | ID_2 | max__left__front__shoulder_elbow_wrist | 96.88246865 |
| 21days | ID_3 | max__left__front__shoulder_elbow_wrist | 83.30474903 |
| 21days | ID_4 | max__left__front__shoulder_elbow_wrist | 101.5530932 |
| 21days | ID_5 | max__left__front__shoulder_elbow_wrist | 90.38312205 |
| 21days | ID_6 | max__left__front__shoulder_elbow_wrist | 105.9018232 |
| 21days | ID_7 | max__left__front__shoulder_elbow_wrist | 97.570341 |
| 21days | ID_8 | max__left__front__shoulder_elbow_wrist | 87.07982696 |
| 21days | ID_9 | max__left__front__shoulder_elbow_wrist | 80.401434 |
| 21days | ID_10 | max__left__front__shoulder_elbow_wrist | 86.63374923 |
| 0days | ID_1 | max__right__back__hip_ankle_toe | 98.09740996 |
| 0days | ID_2 | max__right__back__hip_ankle_toe | 107.1280945 |
| 0days | ID_3 | max__right__back__hip_ankle_toe | 91.38798124 |
| 0days | ID_4 | max__right__back__hip_ankle_toe | 89.79730061 |
| 0days | ID_5 | max__right__back__hip_ankle_toe | 92.44508567 |
| 0days | ID_6 | max__right__back__hip_ankle_toe | 107.5178033 |
| 0days | ID_7 | max__right__back__hip_ankle_toe | 109.9635415 |
| 0days | ID_8 | max__right__back__hip_ankle_toe | 101.4994907 |
| 0days | ID_9 | max__right__back__hip_ankle_toe | 96.96533204 |
| 0days | ID_10 | max__right__back__hip_ankle_toe | 98.60137476 |

| | | | |
|---|---|---|---|
| 0days | ID_11 | max__right__back__hip_ankle_toe | 99.60377412 |
| 0days | ID_12 | max__right__back__hip_ankle_toe | 108.8444601 |
| 0days | ID_13 | max__right__back__hip_ankle_toe | 108.9121 |
| 3days | ID_1 | max__right__back__hip_ankle_toe | 89.9122847 |
| 3days | ID_2 | max__right__back__hip_ankle_toe | 83.29081951 |
| 3days | ID_3 | max__right__back__hip_ankle_toe | 71.29628097 |
| 3days | ID_4 | max__right__back__hip_ankle_toe | 92.12811429 |
| 3days | ID_5 | max__right__back__hip_ankle_toe | 98.33487308 |
| 3days | ID_6 | max__right__back__hip_ankle_toe | 56.93142735 |
| 3days | ID_7 | max__right__back__hip_ankle_toe | 79.44998109 |
| 3days | ID_8 | max__right__back__hip_ankle_toe | 87.01503671 |
| 7days | ID_1 | max__right__back__hip_ankle_toe | 91.26276361 |
| 7days | ID_2 | max__right__back__hip_ankle_toe | 87.81608679 |
| 7days | ID_3 | max__right__back__hip_ankle_toe | 91.70726729 |
| 7days | ID_4 | max__right__back__hip_ankle_toe | 98.27124061 |
| 7days | ID_5 | max__right__back__hip_ankle_toe | 106.57002 |
| 7days | ID_6 | max__right__back__hip_ankle_toe | 97.42542199 |
| 7days | ID_7 | max__right__back__hip_ankle_toe | 104.8321287 |
| 7days | ID_8 | max__right__back__hip_ankle_toe | 93.23755083 |
| 7days | ID_9 | max__right__back__hip_ankle_toe | 100.0327779 |
| 14days | ID_1 | max__right__back__hip_ankle_toe | 90.50324738 |
| 14days | ID_2 | max__right__back__hip_ankle_toe | 90.68399971 |
| 14days | ID_3 | max__right__back__hip_ankle_toe | 93.96665162 |
| 14days | ID_4 | max__right__back__hip_ankle_toe | 99.00391277 |
| 14days | ID_5 | max__right__back__hip_ankle_toe | 99.47641336 |
| 14days | ID_6 | max__right__back__hip_ankle_toe | 84.58263895 |
| 14days | ID_7 | max__right__back__hip_ankle_toe | 102.3983019 |
| 14days | ID_8 | max__right__back__hip_ankle_toe | 109.8576089 |
| 14days | ID_9 | max__right__back__hip_ankle_toe | 102.7613329 |
| 14days | ID_10 | max__right__back__hip_ankle_toe | 103.8333401 |
| 14days | ID_11 | max__right__back__hip_ankle_toe | 93.45188218 |
| 21days | ID_1 | max__right__back__hip_ankle_toe | 101.6829115 |
| 21days | ID_2 | max__right__back__hip_ankle_toe | 95.87819758 |
| 21days | ID_3 | max__right__back__hip_ankle_toe | 96.80301542 |
| 21days | ID_4 | max__right__back__hip_ankle_toe | 92.78694543 |
| 21days | ID_5 | max__right__back__hip_ankle_toe | 95.79613646 |
| 21days | ID_6 | max__right__back__hip_ankle_toe | 85.8144255 |
| 21days | ID_7 | max__right__back__hip_ankle_toe | 102.0594265 |
| 21days | ID_8 | max__right__back__hip_ankle_toe | 98.60975728 |
| 21days | ID_9 | max__right__back__hip_ankle_toe | 103.0829294 |
| 21days | ID_10 | max__right__back__hip_ankle_toe | 107.7974327 |
| 0days | ID_1 | max__right__back__iliac_hip_ankle | 112.0990845 |
| 0days | ID_2 | max__right__back__iliac_hip_ankle | 108.7072396 |
| 0days | ID_3 | max__right__back__iliac_hip_ankle | 119.5516735 |
| 0days | ID_4 | max__right__back__iliac_hip_ankle | 120.3724578 |
| 0days | ID_5 | max__right__back__iliac_hip_ankle | 114.6983898 |
| 0days | ID_6 | max__right__back__iliac_hip_ankle | 119.9536656 |
| 0days | ID_7 | max__right__back__iliac_hip_ankle | 118.0108086 |

| | | | |
|---|---|---|---|
| 0days | ID_8 | max__right__back__iliac_hip_ankle | 116.2949838 |
| 0days | ID_9 | max__right__back__iliac_hip_ankle | 117.0934756 |
| 0days | ID_10 | max__right__back__iliac_hip_ankle | 115.2232001 |
| 0days | ID_11 | max__right__back__iliac_hip_ankle | 110.0888078 |
| 0days | ID_12 | max__right__back__iliac_hip_ankle | 113.8600195 |
| 0days | ID_13 | max__right__back__iliac_hip_ankle | 111.0195792 |
| 3days | ID_1 | max__right__back__iliac_hip_ankle | 115.5956252 |
| 3days | ID_2 | max__right__back__iliac_hip_ankle | 120.4523037 |
| 3days | ID_3 | max__right__back__iliac_hip_ankle | 124.113007 |
| 3days | ID_4 | max__right__back__iliac_hip_ankle | 127.5679809 |
| 3days | ID_5 | max__right__back__iliac_hip_ankle | 128.0599037 |
| 3days | ID_6 | max__right__back__iliac_hip_ankle | 125.0521204 |
| 3days | ID_7 | max__right__back__iliac_hip_ankle | 116.140444 |
| 3days | ID_8 | max__right__back__iliac_hip_ankle | 127.9030432 |
| 7days | ID_1 | max__right__back__iliac_hip_ankle | 104.8485708 |
| 7days | ID_2 | max__right__back__iliac_hip_ankle | 112.7933078 |
| 7days | ID_3 | max__right__back__iliac_hip_ankle | 113.5135571 |
| 7days | ID_4 | max__right__back__iliac_hip_ankle | 121.1774945 |
| 7days | ID_5 | max__right__back__iliac_hip_ankle | 118.1878291 |
| 7days | ID_6 | max__right__back__iliac_hip_ankle | 115.6278345 |
| 7days | ID_7 | max__right__back__iliac_hip_ankle | 106.1001075 |
| 7days | ID_8 | max__right__back__iliac_hip_ankle | 115.2921169 |
| 7days | ID_9 | max__right__back__iliac_hip_ankle | 111.2469024 |
| 14days | ID_1 | max__right__back__iliac_hip_ankle | 113.3525857 |
| 14days | ID_2 | max__right__back__iliac_hip_ankle | 119.1741782 |
| 14days | ID_3 | max__right__back__iliac_hip_ankle | 101.7573607 |
| 14days | ID_4 | max__right__back__iliac_hip_ankle | 113.7868875 |
| 14days | ID_5 | max__right__back__iliac_hip_ankle | 115.1471597 |
| 14days | ID_6 | max__right__back__iliac_hip_ankle | 115.6366594 |
| 14days | ID_7 | max__right__back__iliac_hip_ankle | 123.2281414 |
| 14days | ID_8 | max__right__back__iliac_hip_ankle | 106.4285606 |
| 14days | ID_9 | max__right__back__iliac_hip_ankle | 112.7082898 |
| 14days | ID_10 | max__right__back__iliac_hip_ankle | 110.6225826 |
| 14days | ID_11 | max__right__back__iliac_hip_ankle | 110.02597 |
| 21days | ID_1 | max__right__back__iliac_hip_ankle | 108.6244495 |
| 21days | ID_2 | max__right__back__iliac_hip_ankle | 107.0100279 |
| 21days | ID_3 | max__right__back__iliac_hip_ankle | 111.9578181 |
| 21days | ID_4 | max__right__back__iliac_hip_ankle | 115.4965786 |
| 21days | ID_5 | max__right__back__iliac_hip_ankle | 121.9876146 |
| 21days | ID_6 | max__right__back__iliac_hip_ankle | 118.4272161 |
| 21days | ID_7 | max__right__back__iliac_hip_ankle | 121.6298435 |
| 21days | ID_8 | max__right__back__iliac_hip_ankle | 116.6923442 |
| 21days | ID_9 | max__right__back__iliac_hip_ankle | 108.3975982 |
| 21days | ID_10 | max__right__back__iliac_hip_ankle | 107.3501603 |
| 0days | ID_1 | max__right__front__elbow_wrist_toetip | 101.5465915 |
| 0days | ID_2 | max__right__front__elbow_wrist_toetip | 107.5680155 |
| 0days | ID_3 | max__right__front__elbow_wrist_toetip | 116.2310477 |
| 0days | ID_4 | max__right__front__elbow_wrist_toetip | 114.1121752 |

| | | | |
|---|---|---|---|
| 0days | ID_5 | max__right__front__elbow_wrist_toetip | 96.68553235 |
| 0days | ID_6 | max__right__front__elbow_wrist_toetip | 109.2646057 |
| 0days | ID_7 | max__right__front__elbow_wrist_toetip | 107.061423 |
| 0days | ID_8 | max__right__front__elbow_wrist_toetip | 113.1218669 |
| 0days | ID_9 | max__right__front__elbow_wrist_toetip | 115.7744057 |
| 0days | ID_10 | max__right__front__elbow_wrist_toetip | 88.75873999 |
| 0days | ID_11 | max__right__front__elbow_wrist_toetip | 86.57834199 |
| 0days | ID_12 | max__right__front__elbow_wrist_toetip | 96.54917803 |
| 0days | ID_13 | max__right__front__elbow_wrist_toetip | 104.4515338 |
| 3days | ID_1 | max__right__front__elbow_wrist_toetip | 98.54918166 |
| 3days | ID_2 | max__right__front__elbow_wrist_toetip | 99.12041589 |
| 3days | ID_3 | max__right__front__elbow_wrist_toetip | 82.8112308 |
| 3days | ID_4 | max__right__front__elbow_wrist_toetip | 92.36834177 |
| 3days | ID_5 | max__right__front__elbow_wrist_toetip | 96.04177349 |
| 3days | ID_6 | max__right__front__elbow_wrist_toetip | 82.71672841 |
| 3days | ID_7 | max__right__front__elbow_wrist_toetip | 92.41954918 |
| 3days | ID_8 | max__right__front__elbow_wrist_toetip | 100.5168292 |
| 7days | ID_1 | max__right__front__elbow_wrist_toetip | 82.28177006 |
| 7days | ID_2 | max__right__front__elbow_wrist_toetip | 77.81775026 |
| 7days | ID_3 | max__right__front__elbow_wrist_toetip | 96.65081218 |
| 7days | ID_4 | max__right__front__elbow_wrist_toetip | 100.5536034 |
| 7days | ID_5 | max__right__front__elbow_wrist_toetip | 109.0033982 |
| 7days | ID_6 | max__right__front__elbow_wrist_toetip | 93.83449558 |
| 7days | ID_7 | max__right__front__elbow_wrist_toetip | 107.8572129 |
| 7days | ID_8 | max__right__front__elbow_wrist_toetip | 91.21495971 |
| 7days | ID_9 | max__right__front__elbow_wrist_toetip | 105.9094185 |
| 14days | ID_1 | max__right__front__elbow_wrist_toetip | 100.277387 |
| 14days | ID_2 | max__right__front__elbow_wrist_toetip | 103.7613447 |
| 14days | ID_3 | max__right__front__elbow_wrist_toetip | 68.21320466 |
| 14days | ID_4 | max__right__front__elbow_wrist_toetip | 101.5115169 |
| 14days | ID_5 | max__right__front__elbow_wrist_toetip | 98.65636688 |
| 14days | ID_6 | max__right__front__elbow_wrist_toetip | 99.92760486 |
| 14days | ID_7 | max__right__front__elbow_wrist_toetip | 112.1650696 |
| 14days | ID_8 | max__right__front__elbow_wrist_toetip | 93.97238681 |
| 14days | ID_9 | max__right__front__elbow_wrist_toetip | 109.3471815 |
| 14days | ID_10 | max__right__front__elbow_wrist_toetip | 104.1243676 |
| 14days | ID_11 | max__right__front__elbow_wrist_toetip | 96.04925953 |
| 21days | ID_1 | max__right__front__elbow_wrist_toetip | 84.35084646 |
| 21days | ID_2 | max__right__front__elbow_wrist_toetip | 107.2040119 |
| 21days | ID_3 | max__right__front__elbow_wrist_toetip | 89.27738323 |
| 21days | ID_4 | max__right__front__elbow_wrist_toetip | 108.8625753 |
| 21days | ID_5 | max__right__front__elbow_wrist_toetip | 92.95017037 |
| 21days | ID_6 | max__right__front__elbow_wrist_toetip | 107.0054988 |
| 21days | ID_7 | max__right__front__elbow_wrist_toetip | 110.2696775 |
| 21days | ID_8 | max__right__front__elbow_wrist_toetip | 105.3672396 |
| 21days | ID_9 | max__right__front__elbow_wrist_toetip | 93.63894942 |
| 21days | ID_10 | max__right__front__elbow_wrist_toetip | 105.2930821 |
| 0days | ID_1 | max__right__front__shoulder_elbow_wrist | 92.78129787 |

| | | | |
|---|---|---|---|
| 0days | ID_2 | max__right__front__shoulder_elbow_wrist | 97.63453396 |
| 0days | ID_3 | max__right__front__shoulder_elbow_wrist | 97.41524957 |
| 0days | ID_4 | max__right__front__shoulder_elbow_wrist | 100.5802426 |
| 0days | ID_5 | max__right__front__shoulder_elbow_wrist | 93.68928193 |
| 0days | ID_6 | max__right__front__shoulder_elbow_wrist | 104.9450912 |
| 0days | ID_7 | max__right__front__shoulder_elbow_wrist | 86.43819261 |
| 0days | ID_8 | max__right__front__shoulder_elbow_wrist | 96.18107262 |
| 0days | ID_9 | max__right__front__shoulder_elbow_wrist | 104.4692796 |
| 0days | ID_10 | max__right__front__shoulder_elbow_wrist | 98.35556161 |
| 0days | ID_11 | max__right__front__shoulder_elbow_wrist | 98.79397534 |
| 0days | ID_12 | max__right__front__shoulder_elbow_wrist | 95.66104579 |
| 0days | ID_13 | max__right__front__shoulder_elbow_wrist | 102.9404682 |
| 3days | ID_1 | max__right__front__shoulder_elbow_wrist | 99.92582509 |
| 3days | ID_2 | max__right__front__shoulder_elbow_wrist | 106.6574159 |
| 3days | ID_3 | max__right__front__shoulder_elbow_wrist | 95.97237495 |
| 3days | ID_4 | max__right__front__shoulder_elbow_wrist | 101.77282 |
| 3days | ID_5 | max__right__front__shoulder_elbow_wrist | 105.8384717 |
| 3days | ID_6 | max__right__front__shoulder_elbow_wrist | 99.98260491 |
| 3days | ID_7 | max__right__front__shoulder_elbow_wrist | 82.91539274 |
| 3days | ID_8 | max__right__front__shoulder_elbow_wrist | 95.89536911 |
| 7days | ID_1 | max__right__front__shoulder_elbow_wrist | 91.74715648 |
| 7days | ID_2 | max__right__front__shoulder_elbow_wrist | 99.7368484 |
| 7days | ID_3 | max__right__front__shoulder_elbow_wrist | 104.1167479 |
| 7days | ID_4 | max__right__front__shoulder_elbow_wrist | 100.4532928 |
| 7days | ID_5 | max__right__front__shoulder_elbow_wrist | 88.8668416 |
| 7days | ID_6 | max__right__front__shoulder_elbow_wrist | 99.89928787 |
| 7days | ID_7 | max__right__front__shoulder_elbow_wrist | 84.19134761 |
| 7days | ID_8 | max__right__front__shoulder_elbow_wrist | 104.7109843 |
| 7days | ID_9 | max__right__front__shoulder_elbow_wrist | 85.80438216 |
| 14days | ID_1 | max__right__front__shoulder_elbow_wrist | 98.13646818 |
| 14days | ID_2 | max__right__front__shoulder_elbow_wrist | 92.27651552 |
| 14days | ID_3 | max__right__front__shoulder_elbow_wrist | 95.8072057 |
| 14days | ID_4 | max__right__front__shoulder_elbow_wrist | 79.09497643 |
| 14days | ID_5 | max__right__front__shoulder_elbow_wrist | 77.0886452 |
| 14days | ID_6 | max__right__front__shoulder_elbow_wrist | 101.5909612 |
| 14days | ID_7 | max__right__front__shoulder_elbow_wrist | 103.7774518 |
| 14days | ID_8 | max__right__front__shoulder_elbow_wrist | 101.623618 |
| 14days | ID_9 | max__right__front__shoulder_elbow_wrist | 89.51785551 |
| 14days | ID_10 | max__right__front__shoulder_elbow_wrist | 93.877852 |
| 14days | ID_11 | max__right__front__shoulder_elbow_wrist | 109.7220009 |
| 21days | ID_1 | max__right__front__shoulder_elbow_wrist | 96.22340739 |
| 21days | ID_2 | max__right__front__shoulder_elbow_wrist | 83.83375309 |
| 21days | ID_3 | max__right__front__shoulder_elbow_wrist | 91.94362043 |
| 21days | ID_4 | max__right__front__shoulder_elbow_wrist | 83.9856519 |
| 21days | ID_5 | max__right__front__shoulder_elbow_wrist | 106.0741007 |
| 21days | ID_6 | max__right__front__shoulder_elbow_wrist | 86.65861773 |
| 21days | ID_7 | max__right__front__shoulder_elbow_wrist | 90.2682807 |
| 21days | ID_8 | max__right__front__shoulder_elbow_wrist | 101.844095 |

| | | | |
|---|---|---|---|
| 21days | ID_9 | max__right__front__shoulder_elbow_wrist | 97.15777285 |
| 21days | ID_10 | max__right__front__shoulder_elbow_wrist | 96.1437489 |
| 0days | ID_1 | min__left__back__hip_ankle_toe | 147.0412207 |
| 0days | ID_2 | min__left__back__hip_ankle_toe | 145.978783 |
| 0days | ID_3 | min__left__back__hip_ankle_toe | 120.1431936 |
| 0days | ID_4 | min__left__back__hip_ankle_toe | 139.3240292 |
| 0days | ID_5 | min__left__back__hip_ankle_toe | 139.2348709 |
| 0days | ID_6 | min__left__back__hip_ankle_toe | 149.1721694 |
| 0days | ID_7 | min__left__back__hip_ankle_toe | 138.6449589 |
| 0days | ID_8 | min__left__back__hip_ankle_toe | 150.3085489 |
| 0days | ID_9 | min__left__back__hip_ankle_toe | 132.692295 |
| 0days | ID_10 | min__left__back__hip_ankle_toe | 140.0301931 |
| 0days | ID_11 | min__left__back__hip_ankle_toe | 155.1288184 |
| 0days | ID_12 | min__left__back__hip_ankle_toe | 147.809568 |
| 0days | ID_13 | min__left__back__hip_ankle_toe | 151.6148064 |
| 3days | ID_1 | min__left__back__hip_ankle_toe | 145.7085498 |
| 3days | ID_2 | min__left__back__hip_ankle_toe | 132.4933488 |
| 3days | ID_3 | min__left__back__hip_ankle_toe | 136.9114195 |
| 3days | ID_4 | min__left__back__hip_ankle_toe | 143.8136867 |
| 3days | ID_5 | min__left__back__hip_ankle_toe | 149.3277455 |
| 3days | ID_6 | min__left__back__hip_ankle_toe | 148.5041428 |
| 3days | ID_7 | min__left__back__hip_ankle_toe | 121.7562151 |
| 3days | ID_8 | min__left__back__hip_ankle_toe | 151.7270164 |
| 7days | ID_1 | min__left__back__hip_ankle_toe | 143.9241946 |
| 7days | ID_2 | min__left__back__hip_ankle_toe | 151.217144 |
| 7days | ID_3 | min__left__back__hip_ankle_toe | 138.5545709 |
| 7days | ID_4 | min__left__back__hip_ankle_toe | 149.3910072 |
| 7days | ID_5 | min__left__back__hip_ankle_toe | 137.2258423 |
| 7days | ID_6 | min__left__back__hip_ankle_toe | 151.2679264 |
| 7days | ID_7 | min__left__back__hip_ankle_toe | 130.0495142 |
| 7days | ID_8 | min__left__back__hip_ankle_toe | 151.5681176 |
| 7days | ID_9 | min__left__back__hip_ankle_toe | 135.8193939 |
| 14days | ID_1 | min__left__back__hip_ankle_toe | 123.3397087 |
| 14days | ID_2 | min__left__back__hip_ankle_toe | 125.2653653 |
| 14days | ID_3 | min__left__back__hip_ankle_toe | 148.2992027 |
| 14days | ID_4 | min__left__back__hip_ankle_toe | 125.0830137 |
| 14days | ID_5 | min__left__back__hip_ankle_toe | 123.9229151 |
| 14days | ID_6 | min__left__back__hip_ankle_toe | 133.9349623 |
| 14days | ID_7 | min__left__back__hip_ankle_toe | 146.3442794 |
| 14days | ID_8 | min__left__back__hip_ankle_toe | 152.5210301 |
| 14days | ID_9 | min__left__back__hip_ankle_toe | 139.9180621 |
| 14days | ID_10 | min__left__back__hip_ankle_toe | 134.163788 |
| 14days | ID_11 | min__left__back__hip_ankle_toe | 150.7651374 |
| 21days | ID_1 | min__left__back__hip_ankle_toe | 147.0323842 |
| 21days | ID_2 | min__left__back__hip_ankle_toe | 142.5949188 |
| 21days | ID_3 | min__left__back__hip_ankle_toe | 139.6556164 |
| 21days | ID_4 | min__left__back__hip_ankle_toe | 126.163333 |
| 21days | ID_5 | min__left__back__hip_ankle_toe | 138.2883367 |

| | | | |
|---|---|---|---|
| 21days | ID_6 | min__left__back__hip_ankle_toe | 126.9477694 |
| 21days | ID_7 | min__left__back__hip_ankle_toe | 130.5556982 |
| 21days | ID_8 | min__left__back__hip_ankle_toe | 151.9466522 |
| 21days | ID_9 | min__left__back__hip_ankle_toe | 149.1913322 |
| 21days | ID_10 | min__left__back__hip_ankle_toe | 145.8561784 |
| 0days | ID_1 | min__left__back__iliac_hip_ankle | 164.6210857 |
| 0days | ID_2 | min__left__back__iliac_hip_ankle | 163.6421312 |
| 0days | ID_3 | min__left__back__iliac_hip_ankle | 165.8535725 |
| 0days | ID_4 | min__left__back__iliac_hip_ankle | 163.5440885 |
| 0days | ID_5 | min__left__back__iliac_hip_ankle | 168.5864692 |
| 0days | ID_6 | min__left__back__iliac_hip_ankle | 165.7297628 |
| 0days | ID_7 | min__left__back__iliac_hip_ankle | 163.6402368 |
| 0days | ID_8 | min__left__back__iliac_hip_ankle | 170.6709701 |
| 0days | ID_9 | min__left__back__iliac_hip_ankle | 155.8743763 |
| 0days | ID_10 | min__left__back__iliac_hip_ankle | 170.6238285 |
| 0days | ID_11 | min__left__back__iliac_hip_ankle | 178.8308243 |
| 0days | ID_12 | min__left__back__iliac_hip_ankle | 161.273217 |
| 0days | ID_13 | min__left__back__iliac_hip_ankle | 161.2293732 |
| 3days | ID_1 | min__left__back__iliac_hip_ankle | 171.7858811 |
| 3days | ID_2 | min__left__back__iliac_hip_ankle | 168.7570161 |
| 3days | ID_3 | min__left__back__iliac_hip_ankle | 184.1085158 |
| 3days | ID_4 | min__left__back__iliac_hip_ankle | 168.6934658 |
| 3days | ID_5 | min__left__back__iliac_hip_ankle | 166.8368813 |
| 3days | ID_6 | min__left__back__iliac_hip_ankle | 176.4547118 |
| 3days | ID_7 | min__left__back__iliac_hip_ankle | 176.0680471 |
| 3days | ID_8 | min__left__back__iliac_hip_ankle | 174.815604 |
| 7days | ID_1 | min__left__back__iliac_hip_ankle | 179.8671498 |
| 7days | ID_2 | min__left__back__iliac_hip_ankle | 177.4249699 |
| 7days | ID_3 | min__left__back__iliac_hip_ankle | 173.041342 |
| 7days | ID_4 | min__left__back__iliac_hip_ankle | 167.9842361 |
| 7days | ID_5 | min__left__back__iliac_hip_ankle | 167.4127881 |
| 7days | ID_6 | min__left__back__iliac_hip_ankle | 176.5534784 |
| 7days | ID_7 | min__left__back__iliac_hip_ankle | 169.0594087 |
| 7days | ID_8 | min__left__back__iliac_hip_ankle | 173.0691147 |
| 7days | ID_9 | min__left__back__iliac_hip_ankle | 169.5393214 |
| 14days | ID_1 | min__left__back__iliac_hip_ankle | 170.028497 |
| 14days | ID_2 | min__left__back__iliac_hip_ankle | 173.4257797 |
| 14days | ID_3 | min__left__back__iliac_hip_ankle | 185.9926747 |
| 14days | ID_4 | min__left__back__iliac_hip_ankle | 171.4087567 |
| 14days | ID_5 | min__left__back__iliac_hip_ankle | 173.3833956 |
| 14days | ID_6 | min__left__back__iliac_hip_ankle | 166.4887985 |
| 14days | ID_7 | min__left__back__iliac_hip_ankle | 164.1716742 |
| 14days | ID_8 | min__left__back__iliac_hip_ankle | 170.9004482 |
| 14days | ID_9 | min__left__back__iliac_hip_ankle | 170.788648 |
| 14days | ID_10 | min__left__back__iliac_hip_ankle | 175.9657905 |
| 14days | ID_11 | min__left__back__iliac_hip_ankle | 167.2471454 |
| 21days | ID_1 | min__left__back__iliac_hip_ankle | 173.9259527 |
| 21days | ID_2 | min__left__back__iliac_hip_ankle | 166.0741409 |

| | | | |
|---|---|---|---|
| 21days | ID_3 | min__left__back__iliac_hip_ankle | 171.9912105 |
| 21days | ID_4 | min__left__back__iliac_hip_ankle | 167.556084 |
| 21days | ID_5 | min__left__back__iliac_hip_ankle | 168.0364067 |
| 21days | ID_6 | min__left__back__iliac_hip_ankle | 163.2132321 |
| 21days | ID_7 | min__left__back__iliac_hip_ankle | 162.280375 |
| 21days | ID_8 | min__left__back__iliac_hip_ankle | 166.7748447 |
| 21days | ID_9 | min__left__back__iliac_hip_ankle | 167.0039304 |
| 21days | ID_10 | min__left__back__iliac_hip_ankle | 166.1222 |
| 0days | ID_1 | min__left__front__elbow_wrist_toetip | 190.6963804 |
| 0days | ID_2 | min__left__front__elbow_wrist_toetip | 184.949143 |
| 0days | ID_3 | min__left__front__elbow_wrist_toetip | 166.4973578 |
| 0days | ID_4 | min__left__front__elbow_wrist_toetip | 186.4425396 |
| 0days | ID_5 | min__left__front__elbow_wrist_toetip | 183.1381513 |
| 0days | ID_6 | min__left__front__elbow_wrist_toetip | 189.2530872 |
| 0days | ID_7 | min__left__front__elbow_wrist_toetip | 174.9726881 |
| 0days | ID_8 | min__left__front__elbow_wrist_toetip | 186.6502383 |
| 0days | ID_9 | min__left__front__elbow_wrist_toetip | 179.6208628 |
| 0days | ID_10 | min__left__front__elbow_wrist_toetip | 175.4246851 |
| 0days | ID_11 | min__left__front__elbow_wrist_toetip | 177.9955273 |
| 0days | ID_12 | min__left__front__elbow_wrist_toetip | 185.889003 |
| 0days | ID_13 | min__left__front__elbow_wrist_toetip | 199.5761567 |
| 3days | ID_1 | min__left__front__elbow_wrist_toetip | 182.773857 |
| 3days | ID_2 | min__left__front__elbow_wrist_toetip | 178.1527578 |
| 3days | ID_3 | min__left__front__elbow_wrist_toetip | 167.2368804 |
| 3days | ID_4 | min__left__front__elbow_wrist_toetip | 168.1945912 |
| 3days | ID_5 | min__left__front__elbow_wrist_toetip | 185.2736523 |
| 3days | ID_6 | min__left__front__elbow_wrist_toetip | 162.7330485 |
| 3days | ID_7 | min__left__front__elbow_wrist_toetip | 165.3021763 |
| 3days | ID_8 | min__left__front__elbow_wrist_toetip | 181.0537716 |
| 7days | ID_1 | min__left__front__elbow_wrist_toetip | 167.6104371 |
| 7days | ID_2 | min__left__front__elbow_wrist_toetip | 178.8125769 |
| 7days | ID_3 | min__left__front__elbow_wrist_toetip | 182.3836035 |
| 7days | ID_4 | min__left__front__elbow_wrist_toetip | 186.3958372 |
| 7days | ID_5 | min__left__front__elbow_wrist_toetip | 165.5964541 |
| 7days | ID_6 | min__left__front__elbow_wrist_toetip | 186.8359075 |
| 7days | ID_7 | min__left__front__elbow_wrist_toetip | 175.577155 |
| 7days | ID_8 | min__left__front__elbow_wrist_toetip | 180.1725916 |
| 7days | ID_9 | min__left__front__elbow_wrist_toetip | 167.2945748 |
| 14days | ID_1 | min__left__front__elbow_wrist_toetip | 157.5877504 |
| 14days | ID_2 | min__left__front__elbow_wrist_toetip | 161.0129519 |
| 14days | ID_3 | min__left__front__elbow_wrist_toetip | 166.1516207 |
| 14days | ID_4 | min__left__front__elbow_wrist_toetip | 162.1183818 |
| 14days | ID_5 | min__left__front__elbow_wrist_toetip | 158.9915776 |
| 14days | ID_6 | min__left__front__elbow_wrist_toetip | 179.6207762 |
| 14days | ID_7 | min__left__front__elbow_wrist_toetip | 182.464194 |
| 14days | ID_8 | min__left__front__elbow_wrist_toetip | 192.8429763 |
| 14days | ID_9 | min__left__front__elbow_wrist_toetip | 158.2858851 |
| 14days | ID_10 | min__left__front__elbow_wrist_toetip | 158.505806 |

| | | | |
|---|---|---|---|
| 14days | ID_11 | min__left__front__elbow_wrist_toetip | 169.6498049 |
| 21days | ID_1 | min__left__front__elbow_wrist_toetip | 173.5109729 |
| 21days | ID_2 | min__left__front__elbow_wrist_toetip | 155.099008 |
| 21days | ID_3 | min__left__front__elbow_wrist_toetip | 194.0440285 |
| 21days | ID_4 | min__left__front__elbow_wrist_toetip | 157.2325801 |
| 21days | ID_5 | min__left__front__elbow_wrist_toetip | 181.7846292 |
| 21days | ID_6 | min__left__front__elbow_wrist_toetip | 155.1752468 |
| 21days | ID_7 | min__left__front__elbow_wrist_toetip | 162.8179563 |
| 21days | ID_8 | min__left__front__elbow_wrist_toetip | 188.0455322 |
| 21days | ID_9 | min__left__front__elbow_wrist_toetip | 186.4701767 |
| 21days | ID_10 | min__left__front__elbow_wrist_toetip | 189.4067468 |
| 0days | ID_1 | min__left__front__shoulder_elbow_wrist | 159.4007057 |
| 0days | ID_2 | min__left__front__shoulder_elbow_wrist | 159.0398387 |
| 0days | ID_3 | min__left__front__shoulder_elbow_wrist | 169.0967875 |
| 0days | ID_4 | min__left__front__shoulder_elbow_wrist | 159.1771127 |
| 0days | ID_5 | min__left__front__shoulder_elbow_wrist | 160.6846703 |
| 0days | ID_6 | min__left__front__shoulder_elbow_wrist | 168.3004372 |
| 0days | ID_7 | min__left__front__shoulder_elbow_wrist | 154.3385826 |
| 0days | ID_8 | min__left__front__shoulder_elbow_wrist | 158.5790006 |
| 0days | ID_9 | min__left__front__shoulder_elbow_wrist | 157.3644562 |
| 0days | ID_10 | min__left__front__shoulder_elbow_wrist | 157.2528986 |
| 0days | ID_11 | min__left__front__shoulder_elbow_wrist | 178.1307668 |
| 0days | ID_12 | min__left__front__shoulder_elbow_wrist | 160.2604429 |
| 0days | ID_13 | min__left__front__shoulder_elbow_wrist | 161.6050236 |
| 3days | ID_1 | min__left__front__shoulder_elbow_wrist | 160.3175702 |
| 3days | ID_2 | min__left__front__shoulder_elbow_wrist | 168.6898251 |
| 3days | ID_3 | min__left__front__shoulder_elbow_wrist | 174.4881478 |
| 3days | ID_4 | min__left__front__shoulder_elbow_wrist | 171.0912211 |
| 3days | ID_5 | min__left__front__shoulder_elbow_wrist | 170.3889173 |
| 3days | ID_6 | min__left__front__shoulder_elbow_wrist | 173.6895015 |
| 3days | ID_7 | min__left__front__shoulder_elbow_wrist | 168.9631246 |
| 3days | ID_8 | min__left__front__shoulder_elbow_wrist | 163.0139214 |
| 7days | ID_1 | min__left__front__shoulder_elbow_wrist | 179.27291 |
| 7days | ID_2 | min__left__front__shoulder_elbow_wrist | 161.1427312 |
| 7days | ID_3 | min__left__front__shoulder_elbow_wrist | 163.5644764 |
| 7days | ID_4 | min__left__front__shoulder_elbow_wrist | 164.8556868 |
| 7days | ID_5 | min__left__front__shoulder_elbow_wrist | 174.7093452 |
| 7days | ID_6 | min__left__front__shoulder_elbow_wrist | 165.2443637 |
| 7days | ID_7 | min__left__front__shoulder_elbow_wrist | 172.8900061 |
| 7days | ID_8 | min__left__front__shoulder_elbow_wrist | 162.6271194 |
| 7days | ID_9 | min__left__front__shoulder_elbow_wrist | 175.5311295 |
| 14days | ID_1 | min__left__front__shoulder_elbow_wrist | 176.4368758 |
| 14days | ID_2 | min__left__front__shoulder_elbow_wrist | 175.7049537 |
| 14days | ID_3 | min__left__front__shoulder_elbow_wrist | 179.6366167 |
| 14days | ID_4 | min__left__front__shoulder_elbow_wrist | 169.305756 |
| 14days | ID_5 | min__left__front__shoulder_elbow_wrist | 173.8749867 |
| 14days | ID_6 | min__left__front__shoulder_elbow_wrist | 165.832295 |
| 14days | ID_7 | min__left__front__shoulder_elbow_wrist | 156.5270369 |

| | | | |
|---|---|---|---|
| 14days | ID_8 | min__left__front__shoulder_elbow_wrist | 160.4084698 |
| 14days | ID_9 | min__left__front__shoulder_elbow_wrist | 176.1660423 |
| 14days | ID_10 | min__left__front__shoulder_elbow_wrist | 171.5972762 |
| 14days | ID_11 | min__left__front__shoulder_elbow_wrist | 164.6457876 |
| 21days | ID_1 | min__left__front__shoulder_elbow_wrist | 175.6742946 |
| 21days | ID_2 | min__left__front__shoulder_elbow_wrist | 168.382931 |
| 21days | ID_3 | min__left__front__shoulder_elbow_wrist | 156.4274699 |
| 21days | ID_4 | min__left__front__shoulder_elbow_wrist | 173.549149 |
| 21days | ID_5 | min__left__front__shoulder_elbow_wrist | 170.8004995 |
| 21days | ID_6 | min__left__front__shoulder_elbow_wrist | 180.6563487 |
| 21days | ID_7 | min__left__front__shoulder_elbow_wrist | 169.3810777 |
| 21days | ID_8 | min__left__front__shoulder_elbow_wrist | 159.6027987 |
| 21days | ID_9 | min__left__front__shoulder_elbow_wrist | 162.9043 |
| 21days | ID_10 | min__left__front__shoulder_elbow_wrist | 161.3284819 |
| 0days | ID_1 | min__right__back__hip_ankle_toe | 137.566607 |
| 0days | ID_2 | min__right__back__hip_ankle_toe | 144.436746 |
| 0days | ID_3 | min__right__back__hip_ankle_toe | 118.2785234 |
| 0days | ID_4 | min__right__back__hip_ankle_toe | 127.8559332 |
| 0days | ID_5 | min__right__back__hip_ankle_toe | 124.2762428 |
| 0days | ID_6 | min__right__back__hip_ankle_toe | 145.8065366 |
| 0days | ID_7 | min__right__back__hip_ankle_toe | 145.7821659 |
| 0days | ID_8 | min__right__back__hip_ankle_toe | 152.1876887 |
| 0days | ID_9 | min__right__back__hip_ankle_toe | 139.9501295 |
| 0days | ID_10 | min__right__back__hip_ankle_toe | 127.748874 |
| 0days | ID_11 | min__right__back__hip_ankle_toe | 126.9095843 |
| 0days | ID_12 | min__right__back__hip_ankle_toe | 138.013383 |
| 0days | ID_13 | min__right__back__hip_ankle_toe | 155.2297325 |
| 3days | ID_1 | min__right__back__hip_ankle_toe | 132.0886164 |
| 3days | ID_2 | min__right__back__hip_ankle_toe | 125.5735534 |
| 3days | ID_3 | min__right__back__hip_ankle_toe | 116.0563943 |
| 3days | ID_4 | min__right__back__hip_ankle_toe | 127.6385145 |
| 3days | ID_5 | min__right__back__hip_ankle_toe | 136.7274602 |
| 3days | ID_6 | min__right__back__hip_ankle_toe | 120.4866862 |
| 3days | ID_7 | min__right__back__hip_ankle_toe | 144.2561887 |
| 3days | ID_8 | min__right__back__hip_ankle_toe | 132.637952 |
| 7days | ID_1 | min__right__back__hip_ankle_toe | 126.2641575 |
| 7days | ID_2 | min__right__back__hip_ankle_toe | 122.3565674 |
| 7days | ID_3 | min__right__back__hip_ankle_toe | 119.9246068 |
| 7days | ID_4 | min__right__back__hip_ankle_toe | 137.2615942 |
| 7days | ID_5 | min__right__back__hip_ankle_toe | 146.8664482 |
| 7days | ID_6 | min__right__back__hip_ankle_toe | 134.900534 |
| 7days | ID_7 | min__right__back__hip_ankle_toe | 155.1670414 |
| 7days | ID_8 | min__right__back__hip_ankle_toe | 134.4494994 |
| 7days | ID_9 | min__right__back__hip_ankle_toe | 152.1363313 |
| 14days | ID_1 | min__right__back__hip_ankle_toe | 139.1109345 |
| 14days | ID_2 | min__right__back__hip_ankle_toe | 151.4425914 |
| 14days | ID_3 | min__right__back__hip_ankle_toe | 126.8615427 |
| 14days | ID_4 | min__right__back__hip_ankle_toe | 147.9064403 |

| | | | |
|---|---|---|---|
| 14days | ID_5 | min__right__back__hip_ankle_toe | 148.7383313 |
| 14days | ID_6 | min__right__back__hip_ankle_toe | 124.8138493 |
| 14days | ID_7 | min__right__back__hip_ankle_toe | 140.035204 |
| 14days | ID_8 | min__right__back__hip_ankle_toe | 269.3681928 |
| 14days | ID_9 | min__right__back__hip_ankle_toe | 145.3913629 |
| 14days | ID_10 | min__right__back__hip_ankle_toe | 143.9447768 |
| 14days | ID_11 | min__right__back__hip_ankle_toe | 128.1831943 |
| 21days | ID_1 | min__right__back__hip_ankle_toe | 141.3016094 |
| 21days | ID_2 | min__right__back__hip_ankle_toe | 150.3090076 |
| 21days | ID_3 | min__right__back__hip_ankle_toe | 126.6448869 |
| 21days | ID_4 | min__right__back__hip_ankle_toe | 147.2348186 |
| 21days | ID_5 | min__right__back__hip_ankle_toe | 129.7379315 |
| 21days | ID_6 | min__right__back__hip_ankle_toe | 155.0685307 |
| 21days | ID_7 | min__right__back__hip_ankle_toe | 145.6272112 |
| 21days | ID_8 | min__right__back__hip_ankle_toe | 140.070641 |
| 21days | ID_9 | min__right__back__hip_ankle_toe | 139.5810897 |
| 21days | ID_10 | min__right__back__hip_ankle_toe | 141.8285164 |
| 0days | ID_1 | min__right__back__iliac_hip_ankle | 165.2021713 |
| 0days | ID_2 | min__right__back__iliac_hip_ankle | 157.1508637 |
| 0days | ID_3 | min__right__back__iliac_hip_ankle | 149.9793554 |
| 0days | ID_4 | min__right__back__iliac_hip_ankle | 164.5641548 |
| 0days | ID_5 | min__right__back__iliac_hip_ankle | 164.3973745 |
| 0days | ID_6 | min__right__back__iliac_hip_ankle | 163.0924444 |
| 0days | ID_7 | min__right__back__iliac_hip_ankle | 162.1550654 |
| 0days | ID_8 | min__right__back__iliac_hip_ankle | 160.7031448 |
| 0days | ID_9 | min__right__back__iliac_hip_ankle | 171.5267003 |
| 0days | ID_10 | min__right__back__iliac_hip_ankle | 175.4941822 |
| 0days | ID_11 | min__right__back__iliac_hip_ankle | 172.1369545 |
| 0days | ID_12 | min__right__back__iliac_hip_ankle | 162.7766401 |
| 0days | ID_13 | min__right__back__iliac_hip_ankle | 166.4335421 |
| 3days | ID_1 | min__right__back__iliac_hip_ankle | 184.5324493 |
| 3days | ID_2 | min__right__back__iliac_hip_ankle | 170.2128867 |
| 3days | ID_3 | min__right__back__iliac_hip_ankle | 177.9152088 |
| 3days | ID_4 | min__right__back__iliac_hip_ankle | 175.8283244 |
| 3days | ID_5 | min__right__back__iliac_hip_ankle | 166.5011262 |
| 3days | ID_6 | min__right__back__iliac_hip_ankle | 176.135997 |
| 3days | ID_7 | min__right__back__iliac_hip_ankle | 176.8437227 |
| 3days | ID_8 | min__right__back__iliac_hip_ankle | 166.1998313 |
| 7days | ID_1 | min__right__back__iliac_hip_ankle | 179.0141384 |
| 7days | ID_2 | min__right__back__iliac_hip_ankle | 174.9241482 |
| 7days | ID_3 | min__right__back__iliac_hip_ankle | 186.82825 |
| 7days | ID_4 | min__right__back__iliac_hip_ankle | 165.2977495 |
| 7days | ID_5 | min__right__back__iliac_hip_ankle | 170.3762344 |
| 7days | ID_6 | min__right__back__iliac_hip_ankle | 161.1185885 |
| 7days | ID_7 | min__right__back__iliac_hip_ankle | 161.6724177 |
| 7days | ID_8 | min__right__back__iliac_hip_ankle | 163.4165674 |
| 7days | ID_9 | min__right__back__iliac_hip_ankle | 170.4908928 |
| 14days | ID_1 | min__right__back__iliac_hip_ankle | 164.4972315 |

| | | | |
|---|---|---|---|
| 14days | ID_2 | min__right__back__iliac_hip_ankle | 171.1924813 |
| 14days | ID_3 | min__right__back__iliac_hip_ankle | 181.4079536 |
| 14days | ID_4 | min__right__back__iliac_hip_ankle | 175.177989 |
| 14days | ID_5 | min__right__back__iliac_hip_ankle | 175.5167888 |
| 14days | ID_6 | min__right__back__iliac_hip_ankle | 165.4213545 |
| 14days | ID_7 | min__right__back__iliac_hip_ankle | 165.6178041 |
| 14days | ID_8 | min__right__back__iliac_hip_ankle | 161.8336829 |
| 14days | ID_9 | min__right__back__iliac_hip_ankle | 157.739733 |
| 14days | ID_10 | min__right__back__iliac_hip_ankle | 168.4328513 |
| 14days | ID_11 | min__right__back__iliac_hip_ankle | 162.4237469 |
| 21days | ID_1 | min__right__back__iliac_hip_ankle | 164.9995803 |
| 21days | ID_2 | min__right__back__iliac_hip_ankle | 166.820456 |
| 21days | ID_3 | min__right__back__iliac_hip_ankle | 166.6415109 |
| 21days | ID_4 | min__right__back__iliac_hip_ankle | 169.8237028 |
| 21days | ID_5 | min__right__back__iliac_hip_ankle | 164.0759398 |
| 21days | ID_6 | min__right__back__iliac_hip_ankle | 170.1621808 |
| 21days | ID_7 | min__right__back__iliac_hip_ankle | 167.8193744 |
| 21days | ID_8 | min__right__back__iliac_hip_ankle | 163.2236519 |
| 21days | ID_9 | min__right__back__iliac_hip_ankle | 165.9912849 |
| 21days | ID_10 | min__right__back__iliac_hip_ankle | 160.0504547 |
| 0days | ID_1 | min__right__front__elbow_wrist_toetip | 170.0339887 |
| 0days | ID_2 | min__right__front__elbow_wrist_toetip | 166.890535 |
| 0days | ID_3 | min__right__front__elbow_wrist_toetip | 186.9580726 |
| 0days | ID_4 | min__right__front__elbow_wrist_toetip | 170.0988295 |
| 0days | ID_5 | min__right__front__elbow_wrist_toetip | 189.7303798 |
| 0days | ID_6 | min__right__front__elbow_wrist_toetip | 167.2936008 |
| 0days | ID_7 | min__right__front__elbow_wrist_toetip | 184.0455259 |
| 0days | ID_8 | min__right__front__elbow_wrist_toetip | 179.692531 |
| 0days | ID_9 | min__right__front__elbow_wrist_toetip | 167.26677 |
| 0days | ID_10 | min__right__front__elbow_wrist_toetip | 174.2524409 |
| 0days | ID_11 | min__right__front__elbow_wrist_toetip | 169.205196 |
| 0days | ID_12 | min__right__front__elbow_wrist_toetip | 168.0820087 |
| 0days | ID_13 | min__right__front__elbow_wrist_toetip | 162.0875121 |
| 3days | ID_1 | min__right__front__elbow_wrist_toetip | 167.2473128 |
| 3days | ID_2 | min__right__front__elbow_wrist_toetip | 166.1233948 |
| 3days | ID_3 | min__right__front__elbow_wrist_toetip | 165.790433 |
| 3days | ID_4 | min__right__front__elbow_wrist_toetip | 167.2423151 |
| 3days | ID_5 | min__right__front__elbow_wrist_toetip | 163.0803778 |
| 3days | ID_6 | min__right__front__elbow_wrist_toetip | 166.4967531 |
| 3days | ID_7 | min__right__front__elbow_wrist_toetip | 166.8737621 |
| 3days | ID_8 | min__right__front__elbow_wrist_toetip | 168.8544368 |
| 7days | ID_1 | min__right__front__elbow_wrist_toetip | 164.5990524 |
| 7days | ID_2 | min__right__front__elbow_wrist_toetip | 160.0227807 |
| 7days | ID_3 | min__right__front__elbow_wrist_toetip | 161.5073815 |
| 7days | ID_4 | min__right__front__elbow_wrist_toetip | 169.8251834 |
| 7days | ID_5 | min__right__front__elbow_wrist_toetip | 185.6517438 |
| 7days | ID_6 | min__right__front__elbow_wrist_toetip | 174.5290069 |
| 7days | ID_7 | min__right__front__elbow_wrist_toetip | 191.3853582 |

| | | | |
|---|---|---|---|
| 7days | ID_8 | min__right__front__elbow_wrist_toetip | 170.9888852 |
| 7days | ID_9 | min__right__front__elbow_wrist_toetip | 183.5572218 |
| 14days | ID_1 | min__right__front__elbow_wrist_toetip | 171.1387231 |
| 14days | ID_2 | min__right__front__elbow_wrist_toetip | 175.0013657 |
| 14days | ID_3 | min__right__front__elbow_wrist_toetip | 158.9738122 |
| 14days | ID_4 | min__right__front__elbow_wrist_toetip | 172.0452112 |
| 14days | ID_5 | min__right__front__elbow_wrist_toetip | 172.3047269 |
| 14days | ID_6 | min__right__front__elbow_wrist_toetip | 170.7034391 |
| 14days | ID_7 | min__right__front__elbow_wrist_toetip | 165.0177694 |
| 14days | ID_8 | min__right__front__elbow_wrist_toetip | 159.43749 |
| 14days | ID_9 | min__right__front__elbow_wrist_toetip | 188.2935208 |
| 14days | ID_10 | min__right__front__elbow_wrist_toetip | 169.8941946 |
| 14days | ID_11 | min__right__front__elbow_wrist_toetip | 160.6890114 |
| 21days | ID_1 | min__right__front__elbow_wrist_toetip | 167.8940692 |
| 21days | ID_2 | min__right__front__elbow_wrist_toetip | 189.3744234 |
| 21days | ID_3 | min__right__front__elbow_wrist_toetip | 168.1499567 |
| 21days | ID_4 | min__right__front__elbow_wrist_toetip | 176.5429955 |
| 21days | ID_5 | min__right__front__elbow_wrist_toetip | 158.7068834 |
| 21days | ID_6 | min__right__front__elbow_wrist_toetip | 180.7328527 |
| 21days | ID_7 | min__right__front__elbow_wrist_toetip | 182.5258043 |
| 21days | ID_8 | min__right__front__elbow_wrist_toetip | 164.3850086 |
| 21days | ID_9 | min__right__front__elbow_wrist_toetip | 177.4029136 |
| 21days | ID_10 | min__right__front__elbow_wrist_toetip | 167.3078975 |
| 0days | ID_1 | min__right__front__shoulder_elbow_wrist | 167.5594076 |
| 0days | ID_2 | min__right__front__shoulder_elbow_wrist | 165.4411877 |
| 0days | ID_3 | min__right__front__shoulder_elbow_wrist | 151.7757242 |
| 0days | ID_4 | min__right__front__shoulder_elbow_wrist | 161.2687298 |
| 0days | ID_5 | min__right__front__shoulder_elbow_wrist | 172.4710485 |
| 0days | ID_6 | min__right__front__shoulder_elbow_wrist | 172.0836837 |
| 0days | ID_7 | min__right__front__shoulder_elbow_wrist | 159.4804423 |
| 0days | ID_8 | min__right__front__shoulder_elbow_wrist | 158.3577267 |
| 0days | ID_9 | min__right__front__shoulder_elbow_wrist | 180.7920302 |
| 0days | ID_10 | min__right__front__shoulder_elbow_wrist | 176.5775949 |
| 0days | ID_11 | min__right__front__shoulder_elbow_wrist | 166.2460127 |
| 0days | ID_12 | min__right__front__shoulder_elbow_wrist | 173.117989 |
| 0days | ID_13 | min__right__front__shoulder_elbow_wrist | 177.5191236 |
| 3days | ID_1 | min__right__front__shoulder_elbow_wrist | 171.9144366 |
| 3days | ID_2 | min__right__front__shoulder_elbow_wrist | 162.5037427 |
| 3days | ID_3 | min__right__front__shoulder_elbow_wrist | 171.4350699 |
| 3days | ID_4 | min__right__front__shoulder_elbow_wrist | 174.6256185 |
| 3days | ID_5 | min__right__front__shoulder_elbow_wrist | 175.6781465 |
| 3days | ID_6 | min__right__front__shoulder_elbow_wrist | 179.0694642 |
| 3days | ID_7 | min__right__front__shoulder_elbow_wrist | 162.7436089 |
| 3days | ID_8 | min__right__front__shoulder_elbow_wrist | 172.4220438 |
| 7days | ID_1 | min__right__front__shoulder_elbow_wrist | 163.5977541 |
| 7days | ID_2 | min__right__front__shoulder_elbow_wrist | 183.8641945 |
| 7days | ID_3 | min__right__front__shoulder_elbow_wrist | 165.8388769 |
| 7days | ID_4 | min__right__front__shoulder_elbow_wrist | 163.5583693 |

| | | | |
|---|---|---|---|
| 7days | ID_5 | min__right__front__shoulder_elbow_wrist | 163.9256761 |
| 7days | ID_6 | min__right__front__shoulder_elbow_wrist | 175.0210027 |
| 7days | ID_7 | min__right__front__shoulder_elbow_wrist | 157.5719481 |
| 7days | ID_8 | min__right__front__shoulder_elbow_wrist | 180.2347552 |
| 7days | ID_9 | min__right__front__shoulder_elbow_wrist | 167.8046076 |
| 14days | ID_1 | min__right__front__shoulder_elbow_wrist | 170.4457194 |
| 14days | ID_2 | min__right__front__shoulder_elbow_wrist | 175.2760213 |
| 14days | ID_3 | min__right__front__shoulder_elbow_wrist | 187.446569 |
| 14days | ID_4 | min__right__front__shoulder_elbow_wrist | 165.9493805 |
| 14days | ID_5 | min__right__front__shoulder_elbow_wrist | 163.8274377 |
| 14days | ID_6 | min__right__front__shoulder_elbow_wrist | 163.0330161 |
| 14days | ID_7 | min__right__front__shoulder_elbow_wrist | 173.0932763 |
| 14days | ID_8 | min__right__front__shoulder_elbow_wrist | 177.0766709 |
| 14days | ID_9 | min__right__front__shoulder_elbow_wrist | 154.6229821 |
| 14days | ID_10 | min__right__front__shoulder_elbow_wrist | 174.7191183 |
| 14days | ID_11 | min__right__front__shoulder_elbow_wrist | 180.0441825 |
| 21days | ID_1 | min__right__front__shoulder_elbow_wrist | 181.0484775 |
| 21days | ID_2 | min__right__front__shoulder_elbow_wrist | 160.619939 |
| 21days | ID_3 | min__right__front__shoulder_elbow_wrist | 170.9794967 |
| 21days | ID_4 | min__right__front__shoulder_elbow_wrist | 167.0312112 |
| 21days | ID_5 | min__right__front__shoulder_elbow_wrist | 177.8685191 |
| 21days | ID_6 | min__right__front__shoulder_elbow_wrist | 166.3579638 |
| 21days | ID_7 | min__right__front__shoulder_elbow_wrist | 162.4931435 |
| 21days | ID_8 | min__right__front__shoulder_elbow_wrist | 168.1769528 |
| 21days | ID_9 | min__right__front__shoulder_elbow_wrist | 174.1670074 |
| 21days | ID_10 | min__right__front__shoulder_elbow_wrist | 169.8967262 |

**Suppl. Table 26: Summary of side perspective horizontal angles analysis**

| Day | Measure | Mean | SD |
|---|---|---|---|
| 0days | avg_Angle__left__back__hip_ankle_toe | 119.111343 | 8.65844201 |
| 0days | avg_Angle__left__back__iliac_hip_ankle | 138.043236 | 7.57301528 |
| 0days | avg_Angle__left__front__elbow_wrist_toetip | 145.592204 | 7.90188404 |
| 0days | avg_Angle__left__front__shoulder_elbow_wrist | 120.525495 | 6.21804693 |
| 0days | avg_Angle__right__back__hip_ankle_toe | 115.91592 | 6.76303026 |
| 0days | avg_Angle__right__back__iliac_hip_ankle | 138.815259 | 4.3619968 |
| 0days | avg_Angle__right__front__elbow_wrist_toetip | 142.740176 | 4.03154536 |
| 0days | avg_Angle__right__front__shoulder_elbow_wrist | 122.341684 | 6.47474687 |
| 0days | max__left__back__hip_ankle_toe | 103.136897 | 9.78782328 |
| 0days | max__left__back__iliac_hip_ankle | 115.876492 | 4.64422425 |
| 0days | max__left__front__elbow_wrist_toetip | 111.590846 | 5.6784448 |
| 0days | max__left__front__shoulder_elbow_wrist | 90.8479695 | 7.73242322 |
| 0days | max__right__back__hip_ankle_toe | 100.827981 | 7.11727027 |
| 0days | max__right__back__iliac_hip_ankle | 115.151799 | 3.8553309 |
| 0days | max__right__front__elbow_wrist_toetip | 104.438727 | 9.88292302 |
| 0days | max__right__front__shoulder_elbow_wrist | 97.6834841 | 5.07734702 |
| 0days | min__left__back__hip_ankle_toe | 142.85565 | 9.3320989 |
| 0days | min__left__back__iliac_hip_ankle | 165.701534 | 5.62838471 |
| 0days | min__left__front__elbow_wrist_toetip | 183.161986 | 8.3702189 |
| 0days | min__left__front__shoulder_elbow_wrist | 161.786979 | 6.40634549 |
| 0days | min__right__back__hip_ankle_toe | 137.234011 | 11.4226258 |
| 0days | min__right__back__iliac_hip_ankle | 164.277892 | 6.59481822 |
| 0days | min__right__front__elbow_wrist_toetip | 173.510569 | 8.74332525 |
| 0days | min__right__front__shoulder_elbow_wrist | 167.899285 | 8.53651527 |
| 3days | avg_Angle__left__back__hip_ankle_toe | 115.817149 | 8.18087786 |
| 3days | avg_Angle__left__back__iliac_hip_ankle | 148.784818 | 13.6262678 |
| 3days | avg_Angle__left__front__elbow_wrist_toetip | 137.146706 | 8.55788941 |
| 3days | avg_Angle__left__front__shoulder_elbow_wrist | 123.470624 | 2.82184551 |
| 3days | avg_Angle__right__back__hip_ankle_toe | 107.73194 | 5.57556526 |
| 3days | avg_Angle__right__back__iliac_hip_ankle | 151.140887 | 7.30944456 |
| 3days | avg_Angle__right__front__elbow_wrist_toetip | 130.652427 | 5.47486517 |
| 3days | avg_Angle__right__front__shoulder_elbow_wrist | 128.77017 | 3.89044541 |
| 3days | max__left__back__hip_ankle_toe | 88.0591755 | 23.4851485 |
| 3days | max__left__back__iliac_hip_ankle | 123.818379 | 11.0716034 |
| 3days | max__left__front__elbow_wrist_toetip | 99.3332639 | 9.68828038 |
| 3days | max__left__front__shoulder_elbow_wrist | 91.8456274 | 5.34602804 |
| 3days | max__right__back__hip_ankle_toe | 82.2948522 | 13.1346753 |
| 3days | max__right__back__iliac_hip_ankle | 123.110554 | 5.13016493 |
| 3days | max__right__front__elbow_wrist_toetip | 93.0680063 | 7.01288227 |
| 3days | max__right__front__shoulder_elbow_wrist | 98.6200343 | 7.47640818 |
| 3days | min__left__back__hip_ankle_toe | 141.280266 | 10.2091984 |
| 3days | min__left__back__iliac_hip_ankle | 173.440015 | 5.63790546 |
| 3days | min__left__front__elbow_wrist_toetip | 173.840092 | 8.88748092 |
| 3days | min__left__front__shoulder_elbow_wrist | 168.830279 | 4.92077719 |
| 3days | min__right__back__hip_ankle_toe | 129.433171 | 9.00274326 |
| 3days | min__right__back__iliac_hip_ankle | 174.271193 | 6.2459468 |

| | | | |
|---|---|---|---|
| 3days | min__right__front__elbow_wrist_toetip | 166.463598 | 1.65275189 |
| 3days | min__right__front__shoulder_elbow_wrist | 171.299016 | 5.88986056 |
| 7days | avg_Angle__left__back__hip_ankle_toe | 112.959715 | 6.33633335 |
| 7days | avg_Angle__left__back__iliac_hip_ankle | 144.638469 | 3.88348317 |
| 7days | avg_Angle__left__front__elbow_wrist_toetip | 142.278232 | 7.45072884 |
| 7days | avg_Angle__left__front__shoulder_elbow_wrist | 120.921374 | 6.07764475 |
| 7days | avg_Angle__right__back__hip_ankle_toe | 113.13123 | 6.67643384 |
| 7days | avg_Angle__right__back__iliac_hip_ankle | 137.630599 | 4.42047671 |
| 7days | avg_Angle__right__front__elbow_wrist_toetip | 133.550705 | 9.66743436 |
| 7days | avg_Angle__right__front__shoulder_elbow_wrist | 123.999634 | 6.87872875 |
| 7days | max__left__back__hip_ankle_toe | 98.2985251 | 6.8611388 |
| 7days | max__left__back__iliac_hip_ankle | 117.080054 | 4.74659798 |
| 7days | max__left__front__elbow_wrist_toetip | 100.281581 | 8.33402003 |
| 7days | max__left__front__shoulder_elbow_wrist | 90.7396511 | 5.97570012 |
| 7days | max__right__back__hip_ankle_toe | 96.7950286 | 6.35387056 |
| 7days | max__right__back__iliac_hip_ankle | 113.198636 | 5.27995414 |
| 7days | max__right__front__elbow_wrist_toetip | 96.1248245 | 11.0412133 |
| 7days | max__right__front__shoulder_elbow_wrist | 95.5029877 | 7.91283351 |
| 7days | min__left__back__hip_ankle_toe | 143.22419 | 8.08484991 |
| 7days | min__left__back__iliac_hip_ankle | 172.661312 | 4.50303778 |
| 7days | min__left__front__elbow_wrist_toetip | 176.742126 | 8.22486184 |
| 7days | min__left__front__shoulder_elbow_wrist | 168.870863 | 6.69834259 |
| 7days | min__right__back__hip_ankle_toe | 136.591864 | 12.6848151 |
| 7days | min__right__back__iliac_hip_ankle | 170.348776 | 8.6722293 |
| 7days | min__right__front__elbow_wrist_toetip | 173.562957 | 11.147965 |
| 7days | min__right__front__shoulder_elbow_wrist | 169.046354 | 8.73186362 |
| 14days | avg_Angle__left__back__hip_ankle_toe | 114.360477 | 8.05024414 |
| 14days | avg_Angle__left__back__iliac_hip_ankle | 143.247298 | 11.2720928 |
| 14days | avg_Angle__left__front__elbow_wrist_toetip | 134.898125 | 8.5465791 |
| 14days | avg_Angle__left__front__shoulder_elbow_wrist | 125.61361 | 8.88687637 |
| 14days | avg_Angle__right__back__hip_ankle_toe | 113.443728 | 6.25543739 |
| 14days | avg_Angle__right__back__iliac_hip_ankle | 136.725847 | 5.41883874 |
| 14days | avg_Angle__right__front__elbow_wrist_toetip | 136.363796 | 8.59137686 |
| 14days | avg_Angle__right__front__shoulder_elbow_wrist | 126.390899 | 5.70462638 |
| 14days | max__left__back__hip_ankle_toe | 97.2532861 | 5.90232909 |
| 14days | max__left__back__iliac_hip_ankle | 116.619118 | 3.86336044 |
| 14days | max__left__front__elbow_wrist_toetip | 97.3466441 | 11.2055642 |
| 14days | max__left__front__shoulder_elbow_wrist | 93.2086971 | 7.91715736 |
| 14days | max__right__back__hip_ankle_toe | 97.3199391 | 7.35833156 |
| 14days | max__right__back__iliac_hip_ankle | 112.897125 | 5.82312227 |
| 14days | max__right__front__elbow_wrist_toetip | 98.9096082 | 11.4889138 |
| 14days | max__right__front__shoulder_elbow_wrist | 94.7739591 | 10.017674 |
| 14days | min__left__back__hip_ankle_toe | 136.687042 | 11.4349251 |
| 14days | min__left__back__iliac_hip_ankle | 171.800146 | 5.7981734 |
| 14days | min__left__front__elbow_wrist_toetip | 167.930157 | 11.9373203 |
| 14days | min__left__front__shoulder_elbow_wrist | 170.012372 | 7.36523572 |
| 14days | min__right__back__hip_ankle_toe | 151.436038 | 40.1940476 |
| 14days | min__right__back__iliac_hip_ankle | 168.114692 | 7.0499522 |

| | | | |
|---|---|---|---|
| 14days | min__right__front__elbow_wrist_toetip | 169.409024 | 8.43672161 |
| 14days | min__right__front__shoulder_elbow_wrist | 171.412216 | 9.12569363 |
| 21days | avg_Angle__left__back__hip_ankle_toe | 119.207483 | 8.35426229 |
| 21days | avg_Angle__left__back__iliac_hip_ankle | 136.763799 | 3.99494863 |
| 21days | avg_Angle__left__front__elbow_wrist_toetip | 136.563319 | 11.0291336 |
| 21days | avg_Angle__left__front__shoulder_elbow_wrist | 124.687829 | 9.08652152 |
| 21days | avg_Angle__right__back__hip_ankle_toe | 115.083859 | 5.70163392 |
| 21days | avg_Angle__right__back__iliac_hip_ankle | 134.886993 | 3.72815045 |
| 21days | avg_Angle__right__front__elbow_wrist_toetip | 135.382457 | 10.1569051 |
| 21days | avg_Angle__right__front__shoulder_elbow_wrist | 126.131297 | 8.48583852 |
| 21days | max__left__back__hip_ankle_toe | 105.491319 | 10.5303718 |
| 21days | max__left__back__iliac_hip_ankle | 111.920147 | 4.32213586 |
| 21days | max__left__front__elbow_wrist_toetip | 101.158653 | 8.43863629 |
| 21days | max__left__front__shoulder_elbow_wrist | 91.3171268 | 8.64998268 |
| 21days | max__right__back__hip_ankle_toe | 98.0311178 | 6.13448138 |
| 21days | max__right__back__iliac_hip_ankle | 113.757365 | 5.85045605 |
| 21days | max__right__front__elbow_wrist_toetip | 100.421943 | 9.36935615 |
| 21days | max__right__front__shoulder_elbow_wrist | 93.4133049 | 7.44664291 |
| 21days | min__left__back__hip_ankle_toe | 139.823222 | 9.25821405 |
| 21days | min__left__back__iliac_hip_ankle | 167.297838 | 3.52247117 |
| 21days | min__left__front__elbow_wrist_toetip | 174.358688 | 15.5251392 |
| 21days | min__left__front__shoulder_elbow_wrist | 167.870735 | 7.70650868 |
| 21days | min__right__back__hip_ankle_toe | 141.740424 | 8.66628733 |
| 21days | min__right__back__iliac_hip_ankle | 165.960814 | 3.05745037 |
| 21days | min__right__front__elbow_wrist_toetip | 173.302281 | 9.4879407 |
| 21days | min__right__front__shoulder_elbow_wrist | 169.863944 | 6.42798948 |

**Suppl. Table 27: Statistical test of bside perspective horizontal angle analysis**

| Measure | group1 | group2 | p.adj | sig | Test* |
|---|---|---|---|---|---|
| avg_Angle__left__back__hip_ankle_toe | 0days | 3days | 1 | ns | Rep ANOVA with PH test |
| avg_Angle__left__back__hip_ankle_toe | 0days | 7days | 1 | ns | Rep ANOVA with PH test |
| avg_Angle__left__back__hip_ankle_toe | 0days | 14days | 1 | ns | Rep ANOVA with PH test |
| avg_Angle__left__back__hip_ankle_toe | 0days | 21days | 1 | ns | Rep ANOVA with PH test |
| avg_Angle__left__back__iliac_hip_ankle | 0days | 3days | 0.78 | ns | Rep ANOVA with PH test |
| avg_Angle__left__back__iliac_hip_ankle | 0days | 7days | 1 | ns | Rep ANOVA with PH test |
| avg_Angle__left__back__iliac_hip_ankle | 0days | 14days | 1 | ns | Rep ANOVA with PH test |
| avg_Angle__left__back__iliac_hip_ankle | 0days | 21days | 1 | ns | Rep ANOVA with PH test |
| avg_Angle__left__front__ellbow_wrist_toetip | 0days | 3days | 0.734 | ns | Rep ANOVA with PH test |
| avg_Angle__left__front__ellbow_wrist_toetip | 0days | 7days | 0.734 | ns | Rep ANOVA with PH test |
| avg_Angle__left__front__ellbow_wrist_toetip | 0days | 14days | 0.628 | ns | Rep ANOVA with PH test |
| avg_Angle__left__front__ellbow_wrist_toetip | 0days | 21days | 0.734 | ns | Rep ANOVA with PH test |
| avg_Angle__left__front__shoulder_ellbow_wris | 0days | 3days | 1 | ns | Rep ANOVA with PH test |
| avg_Angle__left__front__shoulder_ellbow_wris | 0days | 7days | 1 | ns | Rep ANOVA with PH test |
| avg_Angle__left__front__shoulder_ellbow_wris | 0days | 14days | 1 | ns | Rep ANOVA with PH test |
| avg_Angle__left__front__shoulder_ellbow_wris | 0days | 21days | 1 | ns | Rep ANOVA with PH test |
| avg_Angle__right__back__hip_ankle_toe | 0days | 3days | 1 | ns | Rep ANOVA with PH test |
| avg_Angle__right__back__hip_ankle_toe | 0days | 7days | 1 | ns | Rep ANOVA with PH test |
| avg_Angle__right__back__hip_ankle_toe | 0days | 14days | 1 | ns | Rep ANOVA with PH test |
| avg_Angle__right__back__hip_ankle_toe | 0days | 21days | 1 | ns | Rep ANOVA with PH test |
| avg_Angle__right__back__iliac_hip_ankle | 0days | 3days | 0.548 | ns | Rep ANOVA with PH test |
| avg_Angle__right__back__iliac_hip_ankle | 0days | 7days | 1 | ns | Rep ANOVA with PH test |
| avg_Angle__right__back__iliac_hip_ankle | 0days | 14days | 1 | ns | Rep ANOVA with PH test |
| avg_Angle__right__back__iliac_hip_ankle | 0days | 21days | 1 | ns | Rep ANOVA with PH test |
| avg_Angle__right__front__ellbow_wrist_toetip | 0days | 3days | 0.579 | ns | Rep ANOVA with PH test |
| avg_Angle__right__front__ellbow_wrist_toetip | 0days | 7days | 0.752 | ns | Rep ANOVA with PH test |
| avg_Angle__right__front__ellbow_wrist_toetip | 0days | 14days | 0.752 | ns | Rep ANOVA with PH test |
| avg_Angle__right__front__ellbow_wrist_toetip | 0days | 21days | 0.752 | ns | Rep ANOVA with PH test |
| avg_Angle__right__front__shoulder_ellbow_wr | 0days | 3days | 1 | ns | Rep ANOVA with PH test |
| avg_Angle__right__front__shoulder_ellbow_wr | 0days | 7days | 1 | ns | Rep ANOVA with PH test |
| avg_Angle__right__front__shoulder_ellbow_wr | 0days | 14days | 1 | ns | Rep ANOVA with PH test |
| avg_Angle__right__front__shoulder_ellbow_wr | 0days | 21days | 1 | ns | Rep ANOVA with PH test |
| max__left__back__hip_ankle_toe | 0days | 3days | 0.276 | ns | Rep ANOVA with PH test |
| max__left__back__hip_ankle_toe | 0days | 7days | 1 | ns | Rep ANOVA with PH test |
| max__left__back__hip_ankle_toe | 0days | 14days | 1 | ns | Rep ANOVA with PH test |
| max__left__back__hip_ankle_toe | 0days | 21days | 1 | ns | Rep ANOVA with PH test |
| max__left__back__iliac_hip_ankle | 0days | 3days | 1 | ns | Rep ANOVA with PH test |
| max__left__back__iliac_hip_ankle | 0days | 7days | 1 | ns | Rep ANOVA with PH test |
| max__left__back__iliac_hip_ankle | 0days | 14days | 1 | ns | Rep ANOVA with PH test |
| max__left__back__iliac_hip_ankle | 0days | 21days | 1 | ns | Rep ANOVA with PH test |
| max__left__front__ellbow_wrist_toetip | 0days | 3days | 0.418 | ns | Rep ANOVA with PH test |
| max__left__front__ellbow_wrist_toetip | 0days | 7days | 0.418 | ns | Rep ANOVA with PH test |
| max__left__front__ellbow_wrist_toetip | 0days | 14days | 0.238 | ns | Rep ANOVA with PH test |
| max__left__front__ellbow_wrist_toetip | 0days | 21days | 0.418 | ns | Rep ANOVA with PH test |
| max__left__front__shoulder_ellbow_wrist | 0days | 3days | 1 | ns | Rep ANOVA with PH test |
| max__left__front__shoulder_ellbow_wrist | 0days | 7days | 1 | ns | Rep ANOVA with PH test |

| | | | | | |
|---|---|---|---|---|---|
| max__left__front__shoulder_ellbow_wrist | 0days | 14days | 1 | ns | Rep ANOVA with PH test |
| max__left__front__shoulder_ellbow_wrist | 0days | 21days | 1 | ns | Rep ANOVA with PH test |
| max__right__back__hip_ankle_toe | 0days | 3days | 0.102 | ns | Rep ANOVA with PH test |
| max__right__back__hip_ankle_toe | 0days | 7days | 1 | ns | Rep ANOVA with PH test |
| max__right__back__hip_ankle_toe | 0days | 14days | 1 | ns | Rep ANOVA with PH test |
| max__right__back__hip_ankle_toe | 0days | 21days | 1 | ns | Rep ANOVA with PH test |
| max__right__back__iliac_hip_ankle | 0days | 3days | 1 | ns | Rep ANOVA with PH test |
| max__right__back__iliac_hip_ankle | 0days | 7days | 1 | ns | Rep ANOVA with PH test |
| max__right__back__iliac_hip_ankle | 0days | 14days | 1 | ns | Rep ANOVA with PH test |
| max__right__back__iliac_hip_ankle | 0days | 21days | 1 | ns | Rep ANOVA with PH test |
| max__right__front__ellbow_wrist_toetip | 0days | 3days | 0.68 | ns | Rep ANOVA with PH test |
| max__right__front__ellbow_wrist_toetip | 0days | 7days | 0.896 | ns | Rep ANOVA with PH test |
| max__right__front__ellbow_wrist_toetip | 0days | 14days | 0.929 | ns | Rep ANOVA with PH test |
| max__right__front__ellbow_wrist_toetip | 0days | 21days | 0.929 | ns | Rep ANOVA with PH test |
| max__right__front__shoulder_ellbow_wrist | 0days | 3days | 1 | ns | Rep ANOVA with PH test |
| max__right__front__shoulder_ellbow_wrist | 0days | 7days | 1 | ns | Rep ANOVA with PH test |
| max__right__front__shoulder_ellbow_wrist | 0days | 14days | 1 | ns | Rep ANOVA with PH test |
| max__right__front__shoulder_ellbow_wrist | 0days | 21days | 1 | ns | Rep ANOVA with PH test |
| min__left__back__hip_ankle_toe | 0days | 3days | 1 | ns | Rep ANOVA with PH test |
| min__left__back__hip_ankle_toe | 0days | 7days | 1 | ns | Rep ANOVA with PH test |
| min__left__back__hip_ankle_toe | 0days | 14days | 1 | ns | Rep ANOVA with PH test |
| min__left__back__hip_ankle_toe | 0days | 21days | 1 | ns | Rep ANOVA with PH test |
| min__left__back__iliac_hip_ankle | 0days | 3days | 1 | ns | Rep ANOVA with PH test |
| min__left__back__iliac_hip_ankle | 0days | 7days | 1 | ns | Rep ANOVA with PH test |
| min__left__back__iliac_hip_ankle | 0days | 14days | 1 | ns | Rep ANOVA with PH test |
| min__left__back__iliac_hip_ankle | 0days | 21days | 1 | ns | Rep ANOVA with PH test |
| min__left__front__ellbow_wrist_toetip | 0days | 3days | 0.769 | ns | Rep ANOVA with PH test |
| min__left__front__ellbow_wrist_toetip | 0days | 7days | 0.769 | ns | Rep ANOVA with PH test |
| min__left__front__ellbow_wrist_toetip | 0days | 14days | 0.176 | ns | Rep ANOVA with PH test |
| min__left__front__ellbow_wrist_toetip | 0days | 21days | 0.769 | ns | Rep ANOVA with PH test |
| min__left__front__shoulder_ellbow_wrist | 0days | 3days | 1 | ns | Rep ANOVA with PH test |
| min__left__front__shoulder_ellbow_wrist | 0days | 7days | 1 | ns | Rep ANOVA with PH test |
| min__left__front__shoulder_ellbow_wrist | 0days | 14days | 1 | ns | Rep ANOVA with PH test |
| min__left__front__shoulder_ellbow_wrist | 0days | 21days | 1 | ns | Rep ANOVA with PH test |
| min__right__back__hip_ankle_toe | 0days | 3days | 1 | ns | Rep ANOVA with PH test |
| min__right__back__hip_ankle_toe | 0days | 7days | 1 | ns | Rep ANOVA with PH test |
| min__right__back__hip_ankle_toe | 0days | 14days | 0.241 | ns | Rep ANOVA with PH test |
| min__right__back__hip_ankle_toe | 0days | 21days | 1 | ns | Rep ANOVA with PH test |
| min__right__back__iliac_hip_ankle | 0days | 3days | 0.912 | ns | Rep ANOVA with PH test |
| min__right__back__iliac_hip_ankle | 0days | 7days | 1 | ns | Rep ANOVA with PH test |
| min__right__back__iliac_hip_ankle | 0days | 14days | 1 | ns | Rep ANOVA with PH test |
| min__right__back__iliac_hip_ankle | 0days | 21days | 1 | ns | Rep ANOVA with PH test |
| min__right__front__ellbow_wrist_toetip | 0days | 3days | 1 | ns | Rep ANOVA with PH test |
| min__right__front__ellbow_wrist_toetip | 0days | 7days | 1 | ns | Rep ANOVA with PH test |
| min__right__front__ellbow_wrist_toetip | 0days | 14days | 1 | ns | Rep ANOVA with PH test |
| min__right__front__ellbow_wrist_toetip | 0days | 21days | 1 | ns | Rep ANOVA with PH test |
| min__right__front__shoulder_ellbow_wrist | 0days | 3days | 1 | ns | Rep ANOVA with PH test |
| min__right__front__shoulder_ellbow_wrist | 0days | 7days | 1 | ns | Rep ANOVA with PH test |

| | | | | | |
|---|---|---|---|---|---|
| min__right__front__shoulder_ellbow_wrist | 0days | 14days | 1 | ns | Rep ANOVA with PH test |
| min__right__front__shoulder_ellbow_wrist | 0days | 21days | 1 | ns | Rep ANOVA with PH test |

**Suppl. Table 28: Parameter importance based on random forest classification baseline, 3, 7, 14, 21 dpi**

| Ranking | Importance (Gini) | Name Parameter | Groups |
|---|---|---|---|
| 1 | 1.404140601 | l-back-toe__Average_Height | baseline, 3,7,14,21 dpi |
| 2 | 1.098874678 | left__back__protraction | baseline, 3,7,14,21 dpi |
| 3 | 1.007898844 | l-front-toe-tip__Movement | baseline, 3,7,14,21 dpi |
| 4 | 0.958852155 | right__back__median | baseline, 3,7,14,21 dpi |
| 5 | 0.94089747 | right__back__movement_per_step | baseline, 3,7,14,21 dpi |
| 6 | 0.925255406 | left__front__protraction | baseline, 3,7,14,21 dpi |
| 7 | 0.922614498 | left__back__average | baseline, 3,7,14,21 dpi |
| 8 | 0.888160511 | l-wrist__Average_Height | baseline, 3,7,14,21 dpi |
| 9 | 0.860963463 | l-back-ankle__Movement | baseline, 3,7,14,21 dpi |
| 10 | 0.844080488 | right__back__protraction | baseline, 3,7,14,21 dpi |
| 11 | 0.73386252 | l-wrist__Movement | baseline, 3,7,14,21 dpi |
| 12 | 0.725433395 | right__front__protraction | baseline, 3,7,14,21 dpi |
| 13 | 0.718656082 | FRBL_RATIO | baseline, 3,7,14,21 dpi |
| 14 | 0.682319128 | l-back-toe__Movement | baseline, 3,7,14,21 dpi |
| 15 | 0.639498284 | right__back__average | baseline, 3,7,14,21 dpi |
| 16 | 0.622255378 | left__front__retraction | baseline, 3,7,14,21 dpi |
| 17 | 0.610304695 | r-wrist__Movement | baseline, 3,7,14,21 dpi |
| 18 | 0.604568189 | left__back__retraction | baseline, 3,7,14,21 dpi |
| 19 | 0.60391649 | right__back__seconds | baseline, 3,7,14,21 dpi |
| 20 | 0.573511732 | r-wrist__Average_Height | baseline, 3,7,14,21 dpi |

**Suppl. Table 29: Parameter importance based on random forest classification baseline and 3 dpi**

| Ranking | Importance (Gini) | Name Parameter | Groups |
|---|---|---|---|
| 1 | 0.343571429 | l-hip__Average_Height | baseline, 3 dpi |
| 2 | 0.236904762 | r-front-toe-tip__Average_Height | baseline, 3 dpi |
| 3 | 0.221428571 | l-hip__Movement | baseline, 3 dpi |
| 4 | 0.215714286 | right__back__median | baseline, 3 dpi |
| 5 | 0.210952381 | right__back__seconds | baseline, 3 dpi |
| 6 | 0.187142857 | r-wrist__Average_Height | baseline, 3 dpi |
| 7 | 0.157142857 | r-hip__Average_Height | baseline, 3 dpi |
| 8 | 0.15 | r-hip__Movement | baseline, 3 dpi |
| 9 | 0.145714286 | right__back__average | baseline, 3 dpi |
| 10 | 0.139285714 | FRBL_RATIO | baseline, 3 dpi |
| 11 | 0.105714286 | l-front-toe-tip__Movement | baseline, 3 dpi |
| 12 | 0.102857143 | left__front__seconds | baseline, 3 dpi |
| 13 | 0.10047619 | left__back__average | baseline, 3 dpi |
| 14 | 0.100238095 | l-back-toe__Average_Height | baseline, 3 dpi |
| 15 | 0.098571429 | r-wrist__Movement | baseline, 3 dpi |
| 16 | 0.098571429 | right__front__average | baseline, 3 dpi |
| 17 | 0.097142857 | l-front-toe-tip__Average_Height | baseline, 3 dpi |
| 18 | 0.096785714 | left__back__seconds | baseline, 3 dpi |
| 19 | 0.078095238 | l-back-toe__Movement | baseline, 3 dpi |
| 20 | 0.066666667 | l-wrist__Movement | baseline, 3 dpi |

**Suppl. Table 30: Parameter importance based on random forest classification baseline and 21 dpi**

| Ranking | Importance (Gini) | Name Parameter | Groups |
|---|---|---|---|
| 1 | 0.698500309 | left__front__retraction | baseline, 21 dpi |
| 2 | 0.328743558 | left__front__median | baseline, 21 dpi |
| 3 | 0.242116574 | right__front__seconds | baseline, 21 dpi |
| 4 | 0.217580396 | l-tail-base__Average_Height | baseline, 21 dpi |
| 5 | 0.171238404 | l-tail-base__Movement | baseline, 21 dpi |
| 6 | 0.166821274 | right__front__retraction | baseline, 21 dpi |
| 7 | 0.166770254 | right__front__average | baseline, 21 dpi |
| 8 | 0.147867965 | r-back-ankle__Movement | baseline, 21 dpi |
| 9 | 0.14038961 | left__front__average | baseline, 21 dpi |
| 10 | 0.138056586 | left__front__protraction | baseline, 21 dpi |
| 11 | 0.136403319 | l-front-toe-tip__Movement | baseline, 21 dpi |
| 12 | 0.122979283 | right__back__seconds | baseline, 21 dpi |
| 13 | 0.122492785 | left__front__seconds | baseline, 21 dpi |
| 14 | 0.118863636 | r-iliac-crest__Average_Height | baseline, 21 dpi |
| 15 | 0.1180906 | left__front__retraction | baseline, 21 dpi |
| 16 | 0.109763451 | l-hip__Average_Height | baseline, 21 dpi |
| 17 | 0.10025974 | left__back__seconds | baseline, 21 dpi |
| 18 | 0.098092146 | left__back__median | baseline, 21 dpi |
| 19 | 0.093073593 | avg_stance_time | baseline, 21 dpi |
| 20 | 0.088682746 | avg_swing_time | baseline, 21 dpi |

**Suppl. Table 31: Error rate between intact and injured mice during ladder rung test**

| Measure | Day | median | mean | sd |
|---|---|---|---|---|
| l-back-toe | intact | 0 | 5.07111936 | 7.338168353 |
| l-back-toe | stroked | 12.5 | 14.3187065 | 13.7133233 |
| l-front-toe-tip | intact | 0 | 8.54711306 | 11.48099774 |
| l-front-toe-tip | stroked | 14.2857143 | 15.5634395 | 13.1484941 |
| r-back-toe | intact | 0 | 1.66624501 | 4.431341378 |
| r-back-toe | stroked | 0 | 6.90764791 | 10.88975155 |
| r-front-toe-tip | intact | 0 | 5.77959198 | 7.420014048 |
| r-front-toe-tip | stroked | 0 | 6.99085418 | 9.900592938 |

**Suppl. Table 32: Statistics for error rate between intact and injured mice during ladder rung test**

| Measure | group1 | group2 | p.adj | sig | Test* |
|---|---|---|---|---|---|
| l-back-toe | intact | stroked | 3.31E-05 | **** | Dunn Test |
| l-front-toe-tip | intact | stroked | 0.00092729 | *** | Dunn Test |
| r-back-toe | intact | stroked | 0.00097815 | *** | Dunn Test |
| r-front-toe-tip | intact | stroked | 0.77167211 | ns | Dunn Test |

**Suppl. Table 33: Error rate in a time course during ladder rung test**

| Measure | Day | median | mean | sd |
|---|---|---|---|---|
| l-front-toe-tip | 0days | 0 | 9.07653292 | 11.59493899 |
| l-front-toe-tip | 3days | 14.2857143 | 14.8750304 | 12.4414783 |
| l-front-toe-tip | 7days | 16.6666667 | 19.0929903 | 14.82716005 |
| l-front-toe-tip | 14days | 12.5 | 16.6249612 | 17.05103827 |
| l-front-toe-tip | 21days | 12.5 | 14.4451062 | 15.43371401 |
| l-back-toe | 0days | 0 | 5.22108844 | 7.162169313 |
| l-back-toe | 3days | 12.5 | 14.6640165 | 13.73748383 |
| l-back-toe | 7days | 10 | 12.2643206 | 15.0284742 |
| l-back-toe | 14days | 9.09090909 | 10.4341769 | 11.42036526 |
| l-back-toe | 21days | 0 | 6.88517964 | 10.68409771 |
| r-back-toe | 0days | 0 | 1.75754611 | 4.534867628 |
| r-back-toe | 3days | 0 | 6.95922814 | 10.89317723 |
| r-back-toe | 7days | 0 | 3.69699123 | 8.373892334 |
| r-back-toe | 14days | 0 | 4.01435574 | 7.569811652 |
| r-back-toe | 21days | 0 | 4.31710495 | 8.36361438 |
| r-front-toe-tip | 0days | 0 | 4.77024959 | 6.140608234 |
| r-front-toe-tip | 3days | 0 | 7.29295811 | 10.08561303 |
| r-front-toe-tip | 7days | 0 | 3.77267426 | 7.092408012 |
| r-front-toe-tip | 14days | 0 | 8.01656476 | 10.54868099 |
| r-front-toe-tip | 21days | 0 | 5.61199553 | 11.92702074 |

**Suppl. Table 34:Statistics throughout a time course during ladder rung test**

| parameter | group1 | group2 | p.adj | p.adj.signif | Test* |
|---|---|---|---|---|---|
| l-back-toe | 0days | 3days | 1.73E-05 | **** | Repeated ANOVA with post hoc test |
| l-back-toe | 0days | 7days | 0.00067035 | *** | Repeated ANOVA with post hoc test |
| l-back-toe | 0days | 14days | 0.01370947 | * | Repeated ANOVA with post hoc test |
| l-back-toe | 0days | 21days | 0.39747262 | ns | Repeated ANOVA with post hoc test |
| l-front-toe-tip | 0days | 3days | 0.01174657 | * | Repeated ANOVA with post hoc test |
| l-front-toe-tip | 0days | 7days | 8.95E-07 | **** | Repeated ANOVA with post hoc test |
| l-front-toe-tip | 0days | 14days | 0.00025968 | *** | Repeated ANOVA with post hoc test |
| l-front-toe-tip | 0days | 21days | 0.01174657 | * | Repeated ANOVA with post hoc test |
| r-back-toe | 0days | 3days | 0.04348641 | * | Repeated ANOVA with post hoc test |
| r-back-toe | 0days | 7days | 0.52810778 | ns | Repeated ANOVA with post hoc test |
| r-back-toe | 0days | 14days | 0.52810778 | ns | Repeated ANOVA with post hoc test |
| r-back-toe | 0days | 21days | 0.52810778 | ns | Repeated ANOVA with post hoc test |
| r-front-toe-tip | 0days | 3days | 0.65330132 | ns | Repeated ANOVA with post hoc test |
| r-front-toe-tip | 0days | 7days | 1 | ns | Repeated ANOVA with post hoc test |
| r-front-toe-tip | 0days | 14days | 0.34823814 | ns | Repeated ANOVA with post hoc test |
| r-front-toe-tip | 0days | 21days | 1 | ns | Repeated ANOVA with post hoc test |

**Suppl. Table 35: Raw data during conventional tests**

| Behavior test | Score | Parameter | Mouse_ID | Day |
|---|---|---|---|---|
| ladder rung | 0.013924 | FL | 1 | baseline |
| ladder rung | 0.034935 | FL | 2 | baseline |
| ladder rung | 0.054539 | FL | 3 | baseline |
| ladder rung | 0.03522 | FL | 4 | baseline |
| ladder rung | 0.04778985 | FL | 5 | baseline |
| ladder rung | 0.04300255 | FL | 6 | baseline |
| ladder rung | 0.04352911 | FL | 7 | baseline |
| ladder rung | 0.04876631 | FL | 8 | baseline |
| ladder rung | 0.04440803 | FL | 9 | baseline |
| ladder rung | 0.06496047 | FL | 10 | baseline |
| ladder rung | 0.06659583 | FL | 11 | baseline |
| ladder rung | 0.04204352 | FL | 12 | baseline |
| ladder rung | 0.184202 | FL | 1 | 3 |
| ladder rung | 0.249242 | FL | 2 | 3 |
| ladder rung | 0.21934 | FL | 3 | 3 |
| ladder rung | 0.2294 | FL | 4 | 3 |
| ladder rung | 0.2885575 | FL | 5 | 3 |
| ladder rung | 0.2779464 | FL | 6 | 3 |
| ladder rung | 0.2447014 | FL | 7 | 3 |
| ladder rung | 0.2359469 | FL | 8 | 3 |
| ladder rung | 0.2811619 | FL | 9 | 3 |
| ladder rung | 0.1750913 | FL | 10 | 3 |
| ladder rung | 0.2881345 | FL | 11 | 3 |
| ladder rung | 0.2974798 | FL | 12 | 3 |
| ladder rung | 0.1924652 | FL | 1 | 7 |
| ladder rung | 0.1758639 | FL | 2 | 7 |
| ladder rung | 0.234539 | FL | 3 | 7 |
| ladder rung | 0.2678063 | FL | 4 | 7 |
| ladder rung | 0.1683676 | FL | 5 | 7 |
| ladder rung | 0.1737422 | FL | 6 | 7 |
| ladder rung | 0.2573249 | FL | 7 | 7 |
| ladder rung | 0.2104021 | FL | 8 | 7 |
| ladder rung | 0.2819787 | FL | 9 | 7 |
| ladder rung | 0.2490475 | FL | 10 | 7 |
| ladder rung | 0.2944221 | FL | 11 | 7 |
| ladder rung | 0.1558081 | FL | 12 | 7 |
| ladder rung | 0.2152912 | FL | 1 | 14 |
| ladder rung | 0.2604266 | FL | 2 | 14 |
| ladder rung | 0.1762194 | FL | 3 | 14 |
| ladder rung | 0.2038705 | FL | 4 | 14 |
| ladder rung | 0.2518033 | FL | 5 | 14 |
| ladder rung | 0.1612487 | FL | 6 | 14 |
| ladder rung | 0.2015572 | FL | 7 | 14 |
| ladder rung | 0.2098318 | FL | 8 | 14 |
| ladder rung | 0.1715338 | FL | 9 | 14 |
| ladder rung | 0.1924448 | FL | 10 | 14 |
| ladder rung | 0.1960025 | FL | 11 | 14 |
| ladder rung | 0.1385358 | FL | 12 | 14 |
| ladder rung | 0.11456 | FL | 1 | 21 |
| ladder rung | 0.14304 | FL | 2 | 21 |

| | | | | |
|---|---|---|---|---|
| ladder rung | 0.12314 | FL | 3 | 21 |
| ladder rung | 0.110394 | FL | 4 | 21 |
| ladder rung | 0.1703651 | FL | 5 | 21 |
| ladder rung | 0.2166202 | FL | 6 | 21 |
| ladder rung | 0.1061859 | FL | 7 | 21 |
| ladder rung | 0.1828388 | FL | 8 | 21 |
| ladder rung | 0.1609431 | FL | 9 | 21 |
| ladder rung | 0.2353897 | FL | 10 | 21 |
| ladder rung | 0.1891668 | FL | 11 | 21 |
| ladder rung | 0.2170997 | FL | 12 | 21 |
| ladder rung | 0.0437 | ladder rung | 1 | baseline |
| ladder rung | 0.066394 | ladder rung | 2 | baseline |
| ladder rung | 0.02452 | ladder rung | 3 | baseline |
| ladder rung | 0.0195602 | ladder rung | 4 | baseline |
| ladder rung | 0.03563 | ladder rung | 5 | baseline |
| ladder rung | 0.02546 | ladder rung | 6 | baseline |
| ladder rung | 0.013825 | ladder rung | 7 | baseline |
| ladder rung | 0.03994745 | ladder rung | 8 | baseline |
| ladder rung | 0.01769902 | ladder rung | 9 | baseline |
| ladder rung | 0.0375225 | ladder rung | 10 | baseline |
| ladder rung | 0.02544737 | ladder rung | 11 | baseline |
| ladder rung | 0.05332957 | ladder rung | 12 | baseline |
| ladder rung | 0.209953 | ladder rung | 1 | 3 |
| ladder rung | 0.1724 | ladder rung | 2 | 3 |
| ladder rung | 0.15355 | ladder rung | 3 | 3 |
| ladder rung | 0.13385 | ladder rung | 4 | 3 |
| ladder rung | 0.25836 | ladder rung | 5 | 3 |
| ladder rung | 0.1836 | ladder rung | 6 | 3 |
| ladder rung | 0.16754 | ladder rung | 7 | 3 |
| ladder rung | 0.1636239 | ladder rung | 8 | 3 |
| ladder rung | 0.2400068 | ladder rung | 9 | 3 |
| ladder rung | 0.1358733 | ladder rung | 10 | 3 |
| ladder rung | 0.157042 | ladder rung | 11 | 3 |
| ladder rung | 0.09770476 | ladder rung | 12 | 3 |
| ladder rung | 0.18425 | ladder rung | 1 | 7 |
| ladder rung | 0.14425 | ladder rung | 2 | 7 |
| ladder rung | 0.1856 | ladder rung | 3 | 7 |
| ladder rung | 0.17285 | ladder rung | 4 | 7 |
| ladder rung | 0.24258 | ladder rung | 5 | 7 |
| ladder rung | 0.143536 | ladder rung | 6 | 7 |
| ladder rung | 0.15342 | ladder rung | 7 | 7 |
| ladder rung | 0.2172824 | ladder rung | 8 | 7 |
| ladder rung | 0.1547405 | ladder rung | 9 | 7 |
| ladder rung | 0.199411 | ladder rung | 10 | 7 |
| ladder rung | 0.1812098 | ladder rung | 11 | 7 |
| ladder rung | 0.2150024 | ladder rung | 12 | 7 |
| ladder rung | 0.10353 | ladder rung | 1 | 14 |
| ladder rung | 0.11429 | ladder rung | 2 | 14 |
| ladder rung | 0.1434835 | ladder rung | 3 | 14 |
| ladder rung | 0.114935 | ladder rung | 4 | 14 |
| ladder rung | 0.1842 | ladder rung | 5 | 14 |
| ladder rung | 0.094852 | ladder rung | 6 | 14 |

| | | | | |
|---|---|---|---|---|
| ladder rung | 0.14356 | ladder rung | 7 | 14 |
| ladder rung | 0.201433 | ladder rung | 8 | 14 |
| ladder rung | 0.1061027 | ladder rung | 9 | 14 |
| ladder rung | 0.1566955 | ladder rung | 10 | 14 |
| ladder rung | 0.1511585 | ladder rung | 11 | 14 |
| ladder rung | 0.1738028 | ladder rung | 12 | 14 |
| ladder rung | 0.0535 | ladder rung | 1 | 21 |
| ladder rung | 0.0245 | ladder rung | 2 | 21 |
| ladder rung | 0.13285 | ladder rung | 3 | 21 |
| ladder rung | 0.18646 | ladder rung | 4 | 21 |
| ladder rung | 0.09435 | ladder rung | 5 | 21 |
| ladder rung | 0.0843536 | ladder rung | 6 | 21 |
| ladder rung | 0.0483535 | ladder rung | 7 | 21 |
| ladder rung | 0.04401902 | ladder rung | 8 | 21 |
| ladder rung | 0.1562648 | ladder rung | 9 | 21 |
| ladder rung | 0.15005045 | ladder rung | 10 | 21 |
| ladder rung | 0.05684677 | ladder rung | 11 | 21 |
| ladder rung | 0.09030958 | ladder rung | 12 | 21 |
| Neurolog Score | 0 | Neurolog Score | 1 | baseline |
| Neurolog Score | 0 | Neurolog Score | 2 | baseline |
| Neurolog Score | 0.5 | Neurolog Score | 3 | baseline |
| Neurolog Score | 0 | Neurolog Score | 4 | baseline |
| Neurolog Score | 0 | Neurolog Score | 5 | baseline |
| Neurolog Score | 0 | Neurolog Score | 6 | baseline |
| Neurolog Score | 0 | Neurolog Score | 7 | baseline |
| Neurolog Score | 3.5 | Neurolog Score | 1 | 0 |
| Neurolog Score | 4 | Neurolog Score | 2 | 0 |
| Neurolog Score | 3.5 | Neurolog Score | 3 | 0 |
| Neurolog Score | 4.5 | Neurolog Score | 4 | 0 |
| Neurolog Score | 4 | Neurolog Score | 5 | 0 |
| Neurolog Score | 4 | Neurolog Score | 6 | 0 |
| Neurolog Score | 4 | Neurolog Score | 7 | 0 |
| Neurolog Score | 1.5 | Neurolog Score | 1 | 3 |
| Neurolog Score | 2 | Neurolog Score | 2 | 3 |
| Neurolog Score | 1.5 | Neurolog Score | 3 | 3 |
| Neurolog Score | 2 | Neurolog Score | 4 | 3 |
| Neurolog Score | 1.5 | Neurolog Score | 5 | 3 |
| Neurolog Score | 2 | Neurolog Score | 6 | 3 |
| Neurolog Score | 1.5 | Neurolog Score | 7 | 3 |
| Neurolog Score | 1.5 | Neurolog Score | 1 | 7 |
| Neurolog Score | 0 | Neurolog Score | 2 | 7 |
| Neurolog Score | 0.5 | Neurolog Score | 3 | 7 |
| Neurolog Score | 0 | Neurolog Score | 4 | 7 |
| Neurolog Score | 1 | Neurolog Score | 5 | 7 |
| Neurolog Score | 1 | Neurolog Score | 6 | 7 |
| Neurolog Score | 1 | Neurolog Score | 7 | 7 |
| Neurolog Score | 0 | Neurolog Score | 1 | 14 |
| Neurolog Score | 0 | Neurolog Score | 2 | 14 |
| Neurolog Score | 0.5 | Neurolog Score | 3 | 14 |
| Neurolog Score | 0.5 | Neurolog Score | 4 | 14 |
| Neurolog Score | 0.5 | Neurolog Score | 5 | 14 |
| Neurolog Score | 0.5 | Neurolog Score | 6 | 14 |

| | | | | |
|---|---|---|---|---|
| Neurolog Score | 0.5 | Neurolog Score | 7 | 14 |
| Neurolog Score | 0 | Neurolog Score | 1 | 21 |
| Neurolog Score | 0 | Neurolog Score | 2 | 21 |
| Neurolog Score | 0 | Neurolog Score | 3 | 21 |
| Neurolog Score | 0.5 | Neurolog Score | 4 | 21 |
| Neurolog Score | 0.5 | Neurolog Score | 5 | 21 |
| Neurolog Score | 0 | Neurolog Score | 6 | 21 |
| Neurolog Score | 0.5 | Neurolog Score | 7 | 21 |
| Cylinder | 0.125 | Dragging | 1 | baseline |
| Cylinder | 0.090909 | Dragging | 2 | baseline |
| Cylinder | 0 | Dragging | 3 | baseline |
| Cylinder | 0 | Dragging | 4 | baseline |
| Cylinder | 0 | Dragging | 5 | baseline |
| Cylinder | 0.0625 | Dragging | 6 | baseline |
| Cylinder | 0 | Dragging | 7 | baseline |
| Cylinder | 0 | Dragging | 8 | baseline |
| Cylinder | 0 | Dragging | 9 | baseline |
| Cylinder | 0.03125 | Dragging | 10 | baseline |
| Cylinder | 0 | Dragging | 11 | baseline |
| Cylinder | 0 | Dragging | 12 | baseline |
| Cylinder | 0.333333 | Dragging | 1 | 3 |
| Cylinder | 0.342857 | Dragging | 2 | 3 |
| Cylinder | 0.215385 | Dragging | 3 | 3 |
| Cylinder | 0.310526 | Dragging | 4 | 3 |
| Cylinder | 0.45 | Dragging | 5 | 3 |
| Cylinder | 0.46 | Dragging | 6 | 3 |
| Cylinder | 0.357143 | Dragging | 7 | 3 |
| Cylinder | 0.333333 | Dragging | 8 | 3 |
| Cylinder | 0.2142857 | Dragging | 9 | 3 |
| Cylinder | 0.125 | Dragging | 10 | 3 |
| Cylinder | 0.5 | Dragging | 11 | 3 |
| Cylinder | 0.2173913 | Dragging | 12 | 3 |
| Cylinder | 0.28 | Dragging | 1 | 7 |
| Cylinder | 0.367 | Dragging | 2 | 7 |
| Cylinder | 0.254 | Dragging | 3 | 7 |
| Cylinder | 0.2842 | Dragging | 4 | 7 |
| Cylinder | 0.28 | Dragging | 5 | 7 |
| Cylinder | 0.33 | Dragging | 6 | 7 |
| Cylinder | 0.615385 | Dragging | 7 | 7 |
| Cylinder | 0.35 | Dragging | 8 | 7 |
| Cylinder | 0.353 | Dragging | 9 | 7 |
| Cylinder | 0.2453 | Dragging | 10 | 7 |
| Cylinder | 0.368 | Dragging | 11 | 7 |
| Cylinder | 0.3524 | Dragging | 12 | 7 |
| Cylinder | 0.2 | Dragging | 1 | 14 |
| Cylinder | 0.29 | Dragging | 2 | 14 |
| Cylinder | 0.22 | Dragging | 3 | 14 |
| Cylinder | 0.2 | Dragging | 4 | 14 |
| Cylinder | 0.2 | Dragging | 5 | 14 |
| Cylinder | 0.57 | Dragging | 6 | 14 |
| Cylinder | 0.4 | Dragging | 7 | 14 |
| Cylinder | 0.36 | Dragging | 8 | 14 |

| | | | | |
|---|---|---|---|---|
| Cylinder | 0.15 | Dragging | 9 | 14 |
| Cylinder | 0.356 | Dragging | 10 | 14 |
| Cylinder | 0.294 | Dragging | 11 | 14 |
| Cylinder | 0.2256 | Dragging | 12 | 14 |
| Cylinder | 0.15 | Dragging | 1 | 21 |
| Cylinder | 0.166 | Dragging | 2 | 21 |
| Cylinder | 0.25 | Dragging | 3 | 21 |
| Cylinder | 0.25 | Dragging | 4 | 21 |
| Cylinder | 0.4 | Dragging | 5 | 21 |
| Cylinder | 0.3 | Dragging | 6 | 21 |
| Cylinder | 0.291667 | Dragging | 7 | 21 |
| Cylinder | 0.381818 | Dragging | 8 | 21 |
| Cylinder | 0.3333333 | Dragging | 9 | 21 |
| Cylinder | 0.25 | Dragging | 10 | 21 |
| Cylinder | 0.5 | Dragging | 11 | 21 |
| Cylinder | 0.4 | Dragging | 12 | 21 |
| Cylinder | 0.5 | Asymmetry | 1 | baseline |
| Cylinder | 0.3 | Asymmetry | 2 | baseline |
| Cylinder | 0.4 | Asymmetry | 3 | baseline |
| Cylinder | 0.7 | Asymmetry | 4 | baseline |
| Cylinder | 0.25 | Asymmetry | 5 | baseline |
| Cylinder | 0 | Asymmetry | 6 | baseline |
| Cylinder | 0.146 | Asymmetry | 7 | baseline |
| Cylinder | 0.14 | Asymmetry | 8 | baseline |
| Cylinder | 0.27 | Asymmetry | 9 | baseline |
| Cylinder | 0.43 | Asymmetry | 10 | baseline |
| Cylinder | 0 | Asymmetry | 11 | baseline |
| Cylinder | 0.090909 | Asymmetry | 12 | baseline |
| Cylinder | 0.5 | Asymmetry | 1 | 3 |
| Cylinder | 0.85 | Asymmetry | 2 | 3 |
| Cylinder | 0.9 | Asymmetry | 3 | 3 |
| Cylinder | 0.6 | Asymmetry | 4 | 3 |
| Cylinder | 0.4744 | Asymmetry | 5 | 3 |
| Cylinder | 0.64674 | Asymmetry | 6 | 3 |
| Cylinder | 0.67 | Asymmetry | 7 | 3 |
| Cylinder | 0.79 | Asymmetry | 8 | 3 |
| Cylinder | 0.77 | Asymmetry | 9 | 3 |
| Cylinder | 0.85 | Asymmetry | 10 | 3 |
| Cylinder | 0.667 | Asymmetry | 11 | 3 |
| Cylinder | 0.37 | Asymmetry | 12 | 3 |
| Cylinder | 0.736 | Asymmetry | 1 | 7 |
| Cylinder | 0.665 | Asymmetry | 2 | 7 |
| Cylinder | 0.6755 | Asymmetry | 3 | 7 |
| Cylinder | 0.7055 | Asymmetry | 4 | 7 |
| Cylinder | 0.68 | Asymmetry | 5 | 7 |
| Cylinder | 0.57 | Asymmetry | 6 | 7 |
| Cylinder | 0.68 | Asymmetry | 7 | 7 |
| Cylinder | 0.7 | Asymmetry | 8 | 7 |
| Cylinder | 0.574 | Asymmetry | 9 | 7 |
| Cylinder | 0.79 | Asymmetry | 10 | 7 |
| Cylinder | 0.4 | Asymmetry | 11 | 7 |
| Cylinder | 0.4577 | Asymmetry | 12 | 7 |

| | | | | |
|---|---|---|---|---|
| Cylinder | 0.75 | Asymmetry | 1 | 14 |
| Cylinder | 0.67 | Asymmetry | 2 | 14 |
| Cylinder | 0.766 | Asymmetry | 3 | 14 |
| Cylinder | 0.799 | Asymmetry | 4 | 14 |
| Cylinder | 0.55 | Asymmetry | 5 | 14 |
| Cylinder | 0.56 | Asymmetry | 6 | 14 |
| Cylinder | 0.77 | Asymmetry | 7 | 14 |
| Cylinder | 0.55 | Asymmetry | 8 | 14 |
| Cylinder | 0.842 | Asymmetry | 9 | 14 |
| Cylinder | 0.95 | Asymmetry | 10 | 14 |
| Cylinder | 0.5 | Asymmetry | 11 | 14 |
| Cylinder | 0.855 | Asymmetry | 12 | 14 |
| Cylinder | 0.9 | Asymmetry | 1 | 21 |
| Cylinder | 0.75 | Asymmetry | 2 | 21 |
| Cylinder | 0.9 | Asymmetry | 3 | 21 |
| Cylinder | 0.82307692 | Asymmetry | 4 | 21 |
| Cylinder | 0.58 | Asymmetry | 5 | 21 |
| Cylinder | 0.5 | Asymmetry | 6 | 21 |
| Cylinder | 0.56 | Asymmetry | 7 | 21 |
| Cylinder | 0.7 | Asymmetry | 8 | 21 |
| Cylinder | 0.8 | Asymmetry | 9 | 21 |
| Cylinder | 0.9 | Asymmetry | 10 | 21 |
| Cylinder | 0.35 | Asymmetry | 11 | 21 |
| Cylinder | 0.76 | Asymmetry | 12 | 21 |
| Rotarod | 0.00689655 | Rotarod_Lat | 1 | baseline |
| Rotarod | 0.00655022 | Rotarod_Lat | 2 | baseline |
| Rotarod | 0.00562852 | Rotarod_Lat | 3 | baseline |
| Rotarod | 0.00993377 | Rotarod_Lat | 4 | baseline |
| Rotarod | 0.01132075 | Rotarod_Lat | 5 | baseline |
| Rotarod | 0.00689655 | Rotarod_Lat | 6 | baseline |
| Rotarod | 0.00806452 | Rotarod_Lat | 7 | baseline |
| Rotarod | 0.015625 | Rotarod_Lat | 8 | baseline |
| Rotarod | 0.01345291 | Rotarod_Lat | 9 | baseline |
| Rotarod | 0.00849858 | Rotarod_Lat | 10 | baseline |
| Rotarod | 0.01003344 | Rotarod_Lat | 11 | baseline |
| Rotarod | 0.01079137 | Rotarod_Lat | 12 | baseline |
| Rotarod | 0.01041667 | Rotarod_Lat | 1 | 3 |
| Rotarod | 0.00898204 | Rotarod_Lat | 2 | 3 |
| Rotarod | 0.04761905 | Rotarod_Lat | 3 | 3 |
| Rotarod | 0.01315789 | Rotarod_Lat | 4 | 3 |
| Rotarod | 0.01376147 | Rotarod_Lat | 5 | 3 |
| Rotarod | 0.01204819 | Rotarod_Lat | 6 | 3 |
| Rotarod | 0.00828729 | Rotarod_Lat | 7 | 3 |
| Rotarod | 0.02362205 | Rotarod_Lat | 8 | 3 |
| Rotarod | 0.01538462 | Rotarod_Lat | 9 | 3 |
| Rotarod | 0.00898204 | Rotarod_Lat | 10 | 3 |
| Rotarod | 0.02912621 | Rotarod_Lat | 11 | 3 |
| Rotarod | 0.02857143 | Rotarod_Lat | 12 | 3 |
| Rotarod | 0.01158301 | Rotarod_Lat | 1 | 7 |
| Rotarod | 0.01195219 | Rotarod_Lat | 2 | 7 |
| Rotarod | 0.00704225 | Rotarod_Lat | 3 | 7 |
| Rotarod | 0.01530612 | Rotarod_Lat | 4 | 7 |

| | | | | |
|---|---|---|---|---|
| Rotarod | 0.01219512 | Rotarod_Lat | 5 | 7 |
| Rotarod | 0.01010101 | Rotarod_Lat | 6 | 7 |
| Rotarod | 0.0078329 | Rotarod_Lat | 7 | 7 |
| Rotarod | 0.01158301 | Rotarod_Lat | 8 | 7 |
| Rotarod | 0.00707547 | Rotarod_Lat | 9 | 7 |
| Rotarod | 0.00977199 | Rotarod_Lat | 10 | 7 |
| Rotarod | 0.00900901 | Rotarod_Lat | 11 | 7 |
| Rotarod | 0.01578947 | Rotarod_Lat | 12 | 7 |
| Rotarod | 0.01075269 | Rotarod_Lat | 1 | 14 |
| Rotarod | 0.00923077 | Rotarod_Lat | 2 | 14 |
| Rotarod | 0.00573614 | Rotarod_Lat | 3 | 14 |
| Rotarod | 0.01145038 | Rotarod_Lat | 4 | 14 |
| Rotarod | 0.00815217 | Rotarod_Lat | 5 | 14 |
| Rotarod | 0.00683371 | Rotarod_Lat | 6 | 14 |
| Rotarod | 0.00674157 | Rotarod_Lat | 7 | 14 |
| Rotarod | 0.01578947 | Rotarod_Lat | 8 | 14 |
| Rotarod | 0.02173913 | Rotarod_Lat | 9 | 14 |
| Rotarod | 0.0096463 | Rotarod_Lat | 10 | 14 |
| Rotarod | 0.00869565 | Rotarod_Lat | 11 | 14 |
| Rotarod | 0.01578947 | Rotarod_Lat | 12 | 14 |
| Rotarod | 0.01016949 | Rotarod_Lat | 1 | 21 |
| Rotarod | 0.00466563 | Rotarod_Lat | 2 | 21 |
| Rotarod | 0.00717703 | Rotarod_Lat | 3 | 21 |
| Rotarod | 0.00806452 | Rotarod_Lat | 4 | 21 |
| Rotarod | 0.00717703 | Rotarod_Lat | 5 | 21 |
| Rotarod | 0.00895522 | Rotarod_Lat | 6 | 21 |
| Rotarod | 0.00791557 | Rotarod_Lat | 7 | 21 |
| Rotarod | 0.00777202 | Rotarod_Lat | 8 | 21 |
| Rotarod | 0.01045296 | Rotarod_Lat | 9 | 21 |
| Rotarod | 0.00714286 | Rotarod_Lat | 10 | 21 |
| Rotarod | 0.01045296 | Rotarod_Lat | 11 | 21 |
| Rotarod | 0.01428571 | Rotarod_Lat | 12 | 21 |
| Single Pellet Grasping | 0 | DragDrop | 1 | baseline |
| Single Pellet Grasping | 10 | DragDrop | 1 | 4 |
| Single Pellet Grasping | 8 | DragDrop | 1 | 7 |
| Single Pellet Grasping | 14 | DragDrop | 1 | 14 |
| Single Pellet Grasping | 8 | DragDrop | 1 | 21 |
| Single Pellet Grasping | 11 | DragDrop | 1 | 28 |
| Single Pellet Grasping | 0 | DragDrop | 2 | baseline |
| Single Pellet Grasping | 21 | DragDrop | 2 | 4 |
| Single Pellet Grasping | 17 | DragDrop | 2 | 7 |
| Single Pellet Grasping | 18 | DragDrop | 2 | 14 |
| Single Pellet Grasping | 16 | DragDrop | 2 | 21 |
| Single Pellet Grasping | 18 | DragDrop | 2 | 28 |
| Single Pellet Grasping | 0 | DragDrop | 3 | baseline |
| Single Pellet Grasping | 20 | DragDrop | 3 | 4 |
| Single Pellet Grasping | 20 | DragDrop | 3 | 7 |
| Single Pellet Grasping | 14 | DragDrop | 3 | 14 |
| Single Pellet Grasping | 16 | DragDrop | 3 | 21 |
| Single Pellet Grasping | 18 | DragDrop | 3 | 28 |
| Single Pellet Grasping | 0 | DragDrop | 4 | baseline |
| Single Pellet Grasping | 20 | DragDrop | 4 | 4 |

| | | | | |
|---|---|---|---|---|
| Single Pellet Grasping | 17 | DragDrop | 4 | 7 |
| Single Pellet Grasping | 13 | DragDrop | 4 | 14 |
| Single Pellet Grasping | NA | DragDrop | 4 | 21 |
| Single Pellet Grasping | NA | DragDrop | 4 | 28 |
| Single Pellet Grasping | 0 | DragDrop | 5 | baseline |
| Single Pellet Grasping | 18 | DragDrop | 5 | 4 |
| Single Pellet Grasping | 15 | DragDrop | 5 | 7 |
| Single Pellet Grasping | 19 | DragDrop | 5 | 14 |
| Single Pellet Grasping | 17 | DragDrop | 5 | 21 |
| Single Pellet Grasping | 12 | DragDrop | 5 | 28 |
| Single Pellet Grasping | 0 | DragDrop | 6 | baseline |
| Single Pellet Grasping | 16 | DragDrop | 6 | 4 |
| Single Pellet Grasping | 19 | DragDrop | 6 | 7 |
| Single Pellet Grasping | 16 | DragDrop | 6 | 14 |
| Single Pellet Grasping | 17 | DragDrop | 6 | 21 |
| Single Pellet Grasping | 16 | DragDrop | 6 | 28 |
| Single Pellet Grasping | 23.33333 | Fail | 1 | baseline |
| Single Pellet Grasping | 40 | Fail | 1 | 4 |
| Single Pellet Grasping | 16.66667 | Fail | 1 | 7 |
| Single Pellet Grasping | 26.66667 | Fail | 1 | 14 |
| Single Pellet Grasping | 20 | Fail | 1 | 21 |
| Single Pellet Grasping | 18.33333 | Fail | 1 | 28 |
| Single Pellet Grasping | 30 | Fail | 2 | baseline |
| Single Pellet Grasping | 48.33333 | Fail | 2 | 4 |
| Single Pellet Grasping | 35 | Fail | 2 | 7 |
| Single Pellet Grasping | 33.33333 | Fail | 2 | 14 |
| Single Pellet Grasping | 43.33333 | Fail | 2 | 21 |
| Single Pellet Grasping | 33.33333 | Fail | 2 | 28 |
| Single Pellet Grasping | 50 | Fail | 3 | baseline |
| Single Pellet Grasping | 43.47826 | Fail | 3 | 4 |
| Single Pellet Grasping | 43.33333 | Fail | 3 | 7 |
| Single Pellet Grasping | 23.33333 | Fail | 3 | 14 |
| Single Pellet Grasping | 30 | Fail | 3 | 21 |
| Single Pellet Grasping | 30 | Fail | 3 | 28 |
| Single Pellet Grasping | 20 | Fail | 4 | baseline |
| Single Pellet Grasping | 33.33333 | Fail | 4 | 4 |
| Single Pellet Grasping | 30.35714 | Fail | 4 | 7 |
| Single Pellet Grasping | 25 | Fail | 4 | 14 |
| Single Pellet Grasping | NA | Fail | 4 | 21 |
| Single Pellet Grasping | NA | Fail | 4 | 28 |
| Single Pellet Grasping | 20 | Fail | 5 | baseline |
| Single Pellet Grasping | 52 | Fail | 5 | 4 |
| Single Pellet Grasping | 35 | Fail | 5 | 7 |
| Single Pellet Grasping | 38.33333 | Fail | 5 | 14 |
| Single Pellet Grasping | 41.66667 | Fail | 5 | 21 |
| Single Pellet Grasping | 22.22222 | Fail | 5 | 28 |
| Single Pellet Grasping | 36.66667 | Fail | 6 | baseline |
| Single Pellet Grasping | 40 | Fail | 6 | 4 |
| Single Pellet Grasping | 45 | Fail | 6 | 7 |
| Single Pellet Grasping | 30 | Fail | 6 | 14 |
| Single Pellet Grasping | 35 | Fail | 6 | 21 |
| Single Pellet Grasping | 33.33333 | Fail | 6 | 28 |

**Suppl. Table 36: Summary of conventional tests**

| Behavior test | Parameter | Day | Value | Median | sd |
|---|---|---|---|---|---|
| Cylinder | Dragging | baseline | 0.02580492 | 0 | 0.043465278 |
| Cylinder | Dragging | 3 | 0.3216045 | 0.333333 | 0.113574086 |
| Cylinder | Dragging | 7 | 0.33994042 | 0.34 | |
| Cylinder | Dragging | 14 | 0.2888 | 0.2578 | 0.11782193 |
| Cylinder | Dragging | 21 | 0.30606819 | 0.2958335 | 0.102420194 |
| Cylinder | Asymmetry | baseline | 0.26890908 | 0.26 | 0.211431335 |
| Cylinder | Asymmetry | 3 | 0.67401167 | 0.6685 | 0.166382983 |
| Cylinder | Asymmetry | 7 | 0.63614167 | 0.67775 | 0.114654481 |
| Cylinder | Asymmetry | 14 | 0.7135 | 0.758 | 0.145089001 |
| Cylinder | Asymmetry | 21 | 0.71025641 | 0.755 | 0.177478706 |
| ladder rung | FL | baseline | 0.04497614 | 0.04396857 | 0.013998926 |
| ladder rung | FL | 3 | 0.24760031 | 0.2469717 | 0.040909479 |
| ladder rung | FL | 7 | 0.22181397 | 0.22247055 | 0.048428787 |
| ladder rung | FL | 14 | 0.19823047 | 0.19877985 | 0.034868174 |
| ladder rung | FL | 21 | 0.16414528 | 0.1656541 | 0.045218736 |
| ladder rung | ladder rung | baseline | 0.03358626 | 0.030545 | 0.015616676 |
| ladder rung | ladder rung | 3 | 0.17279198 | 0.16558195 | 0.045312186 |
| ladder rung | ladder rung | 7 | 0.18284434 | 0.1827299 | 0.031428618 |
| ladder rung | ladder rung | 14 | 0.14067025 | 0.14352175 | 0.034509077 |
| ladder rung | ladder rung | 21 | 0.09348814 | 0.08733159 | 0.051885317 |
| Neurolog Score | Neurolog Scc | baseline | 0.07142857 | 0 | 0.188982237 |
| Neurolog Score | Neurolog Scc | 0 | 3.92857143 | 4 | 0.34503278 |
| Neurolog Score | Neurolog Scc | 3 | 1.71428571 | 1.5 | 0.267261242 |
| Neurolog Score | Neurolog Scc | 7 | 0.71428571 | 1 | 0.56694671 |
| Neurolog Score | Neurolog Scc | 14 | 0.35714286 | 0.5 | 0.243975018 |
| Neurolog Score | Neurolog Scc | 21 | 0.21428571 | 0 | 0.267261242 |
| Rotarod | Rotarod_Lat | baseline | 0.00947435 | 0.00921618 | 0.002998449 |
| Rotarod | Rotarod_Lat | 3 | 0.01832991 | 0.01345968 | 0.011837166 |
| Rotarod | Rotarod_Lat | 7 | 0.01077013 | 0.01084201 | 0.002880008 |
| Rotarod | Rotarod_Lat | 14 | 0.01087979 | 0.00943854 | 0.004698944 |
| Rotarod | Rotarod_Lat | 21 | 0.00868592 | 0.00799004 | 0.002426897 |
| Single Pellet Grasping | Fail | baseline | 30 | 26.666665 | 11.73787867 |
| Single Pellet Grasping | Fail | 4 | 42.8574867 | 41.73913 | 6.648244732 |
| Single Pellet Grasping | Fail | 7 | 34.22619 | 35 | 10.23248587 |
| Single Pellet Grasping | Fail | 14 | 29.4444433 | 28.333335 | 5.643743373 |
| Single Pellet Grasping | Fail | 21 | 34 | 35 | 9.472181346 |
| Single Pellet Grasping | Fail | 28 | 27.444442 | 30 | 6.822258514 |
| Single Pellet Grasping | DragDrop | baseline | 0 | 0 | 0 |
| Single Pellet Grasping | DragDrop | 4 | 17.5 | 19 | 4.086563348 |
| Single Pellet Grasping | DragDrop | 7 | 16 | 17 | 4.289522118 |
| Single Pellet Grasping | DragDrop | 14 | 15.6666667 | 15 | 2.422120283 |
| Single Pellet Grasping | DragDrop | 21 | 14.8 | 16 | 3.834057903 |
| Single Pellet Grasping | DragDrop | 28 | 15 | 16 | 3.31662479 |

**Suppl. Table 37: Statistical test of conventional tests**

| Behavior test | Parameter | group1 | group2 | p.adj.signif | Stat test |
|---|---|---|---|---|---|
| Neurolog Score | Neurolog Scc | baseline | 0 | **** | Repeat. ANOVA with PH |
| Neurolog Score | Neurolog Scc | baseline | 3 | **** | Repeat. ANOVA with PH |
| Neurolog Score | Neurolog Scc | baseline | 7 | * | Repeat. ANOVA with PH |
| Neurolog Score | Neurolog Scc | baseline | 14 | ns | Repeat. ANOVA with PH |
| Neurolog Score | Neurolog Scc | baseline | 21 | ns | Repeat. ANOVA with PH |
| Rotarod | Rotarod_Lat | baseline | 3 | ** | Repeat. ANOVA with PH |
| Rotarod | Rotarod_Lat | baseline | 7 | ns | Repeat. ANOVA with PH |
| Rotarod | Rotarod_Lat | baseline | 14 | ns | Repeat. ANOVA with PH |
| Rotarod | Rotarod_Lat | baseline | 21 | ns | Repeat. ANOVA with PH |
| Cylinder | Dragging | baseline | 3 | **** | Repeat. ANOVA with PH |
| Cylinder | Dragging | baseline | 7 | **** | Repeat. ANOVA with PH |
| Cylinder | Dragging | baseline | 14 | **** | Repeat. ANOVA with PH |
| Cylinder | Dragging | baseline | 21 | **** | Repeat. ANOVA with PH |
| Cylinder | Asymmetry | baseline | 3 | **** | Repeat. ANOVA with PH |
| Cylinder | Asymmetry | baseline | 7 | **** | Repeat. ANOVA with PH |
| Cylinder | Asymmetry | baseline | 14 | **** | Repeat. ANOVA with PH |
| Cylinder | Asymmetry | baseline | 21 | **** | Repeat. ANOVA with PH |
| ladder rung | FL | baseline | 3 | **** | Repeat. ANOVA with PH |
| ladder rung | FL | baseline | 7 | **** | Repeat. ANOVA with PH |
| ladder rung | FL | baseline | 14 | **** | Repeat. ANOVA with PH |
| ladder rung | FL | baseline | 21 | **** | Repeat. ANOVA with PH |
| ladder rung | ladder rung | baseline | 3 | **** | Repeat. ANOVA with PH |
| ladder rung | ladder rung | baseline | 7 | **** | Repeat. ANOVA with PH |
| ladder rung | ladder rung | baseline | 14 | **** | Repeat. ANOVA with PH |
| ladder rung | ladder rung | baseline | 21 | ** | Repeat. ANOVA with PH |
| Single Pellet Grasping | Fail | baseline | 4 | ns | Repeat. ANOVA with PH |
| Single Pellet Grasping | Fail | baseline | 7 | ns | Repeat. ANOVA with PH |
| Single Pellet Grasping | Fail | baseline | 14 | ns | Repeat. ANOVA with PH |
| Single Pellet Grasping | Fail | baseline | 21 | ns | Repeat. ANOVA with PH |
| Single Pellet Grasping | Fail | baseline | 28 | ns | Repeat. ANOVA with PH |
| Single Pellet Grasping | DragDrop | baseline | 4 | **** | Repeat. ANOVA with PH |
| Single Pellet Grasping | DragDrop | baseline | 7 | **** | Repeat. ANOVA with PH |
| Single Pellet Grasping | DragDrop | baseline | 14 | **** | Repeat. ANOVA with PH |
| Single Pellet Grasping | DragDrop | baseline | 21 | **** | Repeat. ANOVA with PH |
| Single Pellet Grasping | DragDrop | baseline | 28 | **** | Repeat. ANOVA with PH |
